# An Unbiased Proteomic Platform for ATE1-based Arginylation Profiling

**DOI:** 10.1101/2024.06.01.596974

**Authors:** Zongtao Lin, Yixuan Xie, Joanna Gongora, Xingyu Liu, Emily Zahn, Bibhuti Bhusana Palai, Daniel H. Ramirez, Richard M. Searfoss, Francisca N. Vitorino, Rashmi Karki, Geoffrey P. Dann, Chenfeng Zhao, Xian Han, Brittany MacTaggart, Xin Lan, Dechen Fu, Lina Greenberg, Yi Zhang, Kory J. Lavine, Michael J. Greenberg, Dongwen Lv, Anna Kashina, Benjamin A. Garcia

**Affiliations:** Department of Biochemistry and Molecular Biophysics, Washington University in St. Louis, St. Louis, MO 63110; Department of Biochemistry and Biophysics, University of Pennsylvania, Philadelphia, PA, 19104; McKelvey School of Engineering, Washington University in St. Louis, St. Louis, MO 63110; School of Veterinary Medicine, University of Pennsylvania, Philadelphia, PA 19104; Department of Biochemistry, Case Western Reserve University, Cleveland, OH 44106; Department of Medicine, Washington University School of Medicine, St. Louis, MO, USA; Department of Biochemistry and Structural Biology, University of Texas Health Science Center at San Antonio, San Antonio, TX 78229

**Keywords:** arginylation, arginyltransferase, proteomics, posttranslational modification, protein profiling

## Abstract

Protein arginylation is an essential posttranslational modification (PTM) catalyzed by arginyl-tRNA-protein transferase 1 (ATE1) in mammalian systems. Arginylation features a post-translational conjugation of an arginyl to a protein, making it extremely challenging to differentiate from translational arginine residues with the same mass in a protein sequence. Here we present a general ATE1-based arginylation profiling platform for the unbiased discovery of arginylation substrates and their precise modification sites. This method integrates isotopic arginine labeling into an ATE1 assay utilizing biological lysates (*ex vivo*) rather than live cells, thus eliminating translational bias derived from the ribosomal activity and enabling *bona fide* arginylation identification using isotopic features. The method has been successfully applied to an array of peptide, protein, cell, patient, and animal tissue samples using 20 µg sample input, with 235 unique arginylation sites revealed from human proteomes. Representative sites were validated and followed up for their biological functions. The developed platform is globally applicable to the aforementioned sample types and therefore paves the way for functional studies of this difficult-to-characterize protein modification.

## Introduction

As a critical posttranslational modification (PTM), arginylation is catalyzed by arginyltransferase (ATE1) which is the only known enzyme capable of installing arginylation in mammalian systems^1^. The absence of arginylation in ATE1 knockout (KO) models resulted in embryonic lethality due to heart defects in cardiovascular development and angiogenic remodeling^1^. Specifically, ATE1KO caused thinned myocardium with immature septa, non-separation of the aorta and pulmonary artery, resulting in defects in cardiac contractility, myofibril dysfunction and eventually embryonic death^1^, demonstrating the essential nature of arginylation^1^. Tissue-specific knockdown (KD) or deletion (KO) of arginylation resulted in impaired myosin phosphorylation and thrombus formation^2^, elevated myocardial fibrosis and progressive heart failure^3^, cardiomyocyte hypertrophy^4^, and many other symptoms^5,6^. At the molecular level, the arginylation field has long tried to understand the biological roles of arginylation in cardiovascular-associated proteins including β-actin (Asp3 arginylation) regulating cytoskeleton and cell motility^7,8^, calreticulin (CALR, Asp18 arginylation) regulating stress granules in a Ca^2+^-dependent manner^9,10^, and RGS4/5/16 (tri-oxidized Cys2 arginylation) acting as nitric oxide and oxygen sensors^11,12^. Many of these studies focused on the implications of shortened half-lives of arginylated proteins involved in cardiovascular biology^1,7-12^.

Meanwhile, many studies suggested that arginylation is also important for certain proteins to function properly (noncanonical roles) beyond its canonical role in the *N*-degron pathway^13,14^. It is reported that arginylation of β-amyloid guides proper α-helical shape preventing misfolding and aggregation^15^. Arginylation of α-synuclein (α-syn) facilitates brain health by preventing neurodegeneration^16,17^. Arginylation of calreticulin promotes its association with stress granules^18,19^. Decreased arginylation of nuclear proteins results in smaller nucleus size and architecture^20^. In addition, arginylation has been detected on histone proteins^20,21^, potentially facilitating their interaction with DNA through the positively charged Arg residue installed post-translationally. Other prominent arginylated examples are chaperone HSPA5^22^, BRCA1^23^, PDI^24^, and CDC6^25^. It is becoming clear that protein arginylation has many biological functions yet to be determined in regulating physiological processes. However, only a handful of proteins have been validated for their arginylation sites so far, and the discovery of arginylation substrates and their modification sites is becoming a bottleneck for the study of arginylation biology.

Identification of arginylation has been inherently challenging because both ATE1 and ribosomes use the same arginine (Arg or R) source, arginyl-tRNA, to add an arginyl group to a protein, yielding posttranslational and translational addition respectively of arginyl with the same mass shift (+156 Da)^13^ (**Fig. 1a**). This makes unbiased differentiation of posttranslational arginylation and translational arginine residues extremely difficult when both ATE1 and the ribosome are active (*i*.*e*. when cells are alive). In addition, arginylated proteins may go through the Arg/*N*-degron pathway for rapid ubiquitin-mediated degradation, decreasing the endogenous arginylation level for detection^13,25^. Furthermore, arginylation mostly happens on the protein *N*-termini after proteolytic cleavage, which is poorly understood for individual proteins in the whole proteome^13^. Additional confusion comes from trypsin miscleavage resulting in peptides starting with an R residue in bottom-up proteomics. Disregarding those challenges, efforts have been made aiming to systematically identify arginylation.

**Figure 1.**
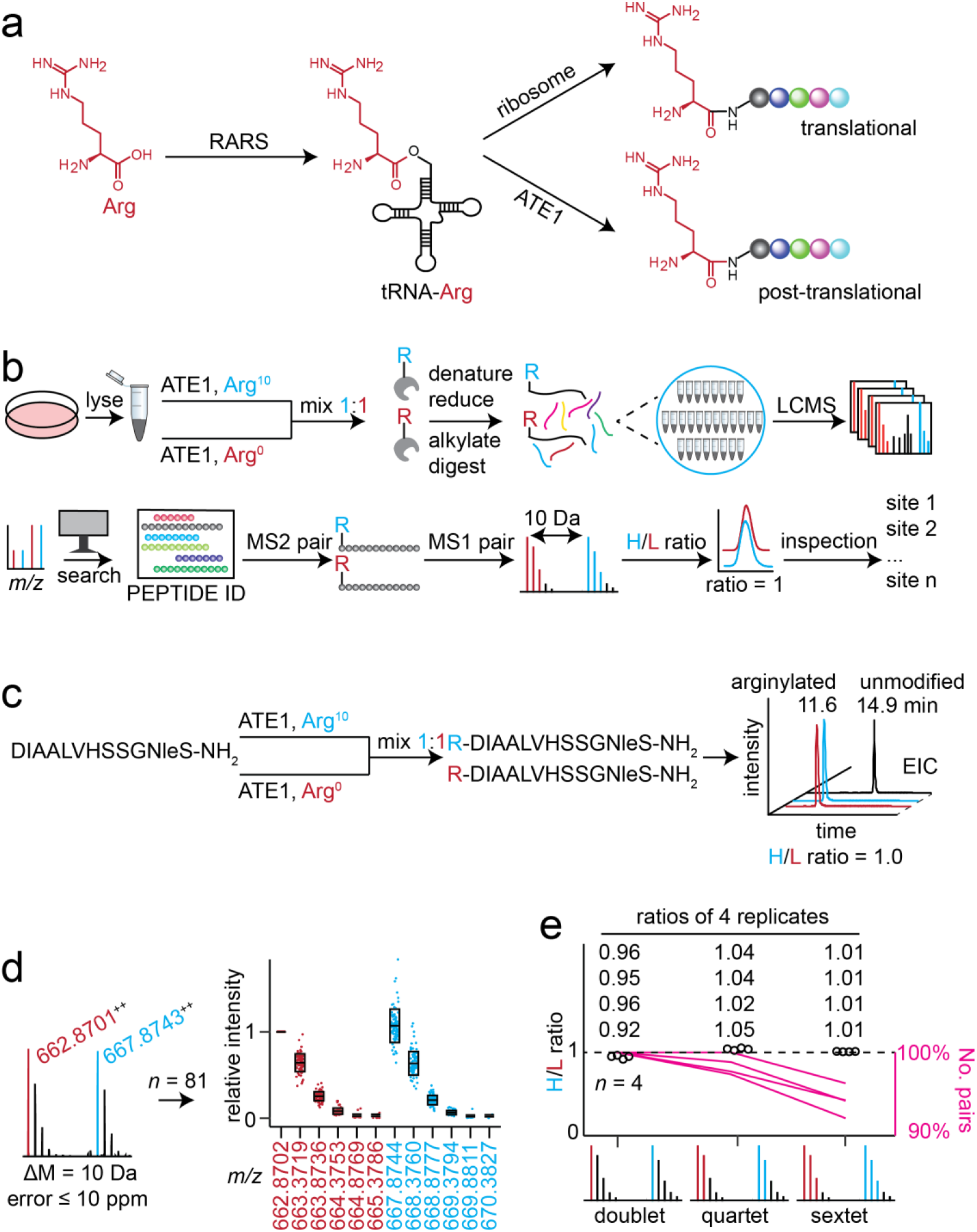
Isotopic arginine labeling and detection strategy for ATE1 substrates and arginylation sites. **a**, scheme of arginyl installation onto proteins by ribosomal synthesis (translational) and ATE1 (post-translational). RARS, arginine-tRNA ligase. ATE1, arginyl-tRNA-protein transferase 1. **b**, arginylation profiling platform for arginylation site and substrate discovery from biological samples. Lysate is labeled by isotopic arginine molecules (Arg^10^ and Arg^0^) respectively using ATE1 assay, mixed, and digested. Peptides are fractionated and analyzed by mass spectrometry in data-dependent acquisition (DDA) mode. Proteomics data is searched to produce peptide identifications (IDs), among which peptide pairs modified by heavy and light Arg are further evaluated for MS1 isotopic features. **c**, isotopic arginylation of a peptide ATE1 substrate. EICs were extracted using monoisotopic peaks based on calculated *m/z* values. EIC in black indicates chromatography of the unmodified peptide. **d**, isotopic feature in MS1 spectra and their summary. The first monoisotopic peak in each MS1 scan is set at 1 for normalization. Relative intensities of other isotopic peaks are displayed. 10-ppm error is set for all MS1 isotopic peaks. **e**, ratio summary of MS1 pairs in 4 replicates (*n* = 4) using doublet, quartet, and sextet peaks. Detailed ratios from the replicates are provided. The numbers of pairs in quartet and sextet are normalized to the numbers of pairs in doublet from respective replicate. EIC: extracted ion chromatogram. Nle: norleucine.

One early approach to screening arginylation substrates involved adding sub-divisions of a complete cDNA library to a transcription-translation-degradation system, which assumes that arginylated substrate is prone to the *N*-degron pathway^26^. A few arginylation substrates (*e*.*g*., BiP and PDI) have been identified from the whole proteome through the combination of ATE1 assay *ex vivo* and [^3^H]arginine autoradiography^24^. The autoradiography was also used for further identification of individual proteins (*e*.*g*., CALR^10^ and α-syn^16^) as ATE1 substrates. Notably, these methods heavily relied on mass spectrometry to reveal the arginylation sites on the labeled (*ex vivo*) proteins. In contrast, proteomic profiling was attempted to identify the endogenous (*in vivo*) arginylation sites from animal tissues using anti-arginylation antibody enrichment and proteomics. Such an approach enabled the first proteomics-based analysis to identify 43 plausible arginylated proteins^21^. Similarly, two later studies identified 19 and 15 proteins potentially modified by arginylation on their side-chain Asp and Glu residues^27,28^, a different ATE1 catalysis mechanism from canonical *N-*term arginylation. However, most of those sites have little overlap with existing confirmed proteins/sites from *ex vivo* experiments and were largely postulated from search algorithms without additional validation. Therefore, there is still a huge unmet need for the creation of an unbiased profiling method for the arginylation field at this stage.

Inspired by unbiased labeling using isotopic Arg^24^ and the screening power of mass spectrometry-based proteomics^21^, here we present a profiling technology for the unbiased discovery of ATE1 substrates and arginylation sites from complex whole proteomes (**Fig. 1b**). We first established the workflow using peptide and proteome-wide peptides as a proof-of-concept, then applied the technology to protein and whole proteomes for arginylation discovery. As a result, a large catalog of unbiased arginylation sites has been established from various cell and tissue samples. To facilitate data processing, a new computational tool, “ArginylomePlot”, was developed and made publicly available for isotopic pair analysis. To be beneficial to the arginylation field and broader research community, we have established a publicly available database indexing all arginylation sites discovered from this study, allowing others interested in protein arginylation to have easy access to this growing type of data. This work could serve as the technological foundation for studying the functions of this essential PTM, and it will have a long-lasting impact on the arginylation field by opening new biochemical and biological frontiers.

## Results

### Isotopic arginylation of substrate peptide by ATE1 *in vitro*

To test the proof-of-concept of the designed arginylation profiling strategy based on ATE1 enzymatic activity (**Fig. 1b**), we started by isotopically labeling a standard peptide (modified from human β-actin sequence) with an unmodified Asp (D) residue at the *N*-terminus (**Fig. 1c, Fig. S1**). The reconstituted ATE1 assay was slightly modified from the well-established conditions in our previous report^29^, where arginine-tRNA ligase (RARS) and ATE1 enzymes were kept at μM range. The arginylation activity of the assay was tested to be dependent on essential components such as arginine, ATP, tRNA, RARS, ATE1, and substrate (**Fig. S2**). Replacement of *N*-terminal D residue to acetylated D or Val produced no detectable arginylation products (**Fig. S3-4**). The arginylation of the standard peptide was time-dependent (0-60 min), and most of the peptides were arginylated with an incubation time of 30 min or longer (**Fig. S5**). Higher starting concentrations (20-300 μM) of substrate produced more arginylation product but correlated to lower overall product yields (**Fig. S5**). The incubation time and substrate concentration were set at 30 min and 100 μM, respectively, while arginine (2 mM) and ATP (2 mM) remained excessive.

After the ATE1 assay using isotopic Arg^10^ and Arg^0^ respectively, the reactions on standard peptide were mixed (1:1) and desalted for LC-MS analysis (**Fig. 1c**). The extracted ion chromatograms of the differentially labeled products were manually examined for hypothetical isotopic ratio (1.0) based on co-elution peak intensities (**Fig. 1c**), demonstrating that the isotopic labeling strategy was capable of introducing unnatural Arg^10^ to arginylation sites for unbiased identification. The data was also searched by Byonic to confirm the product identities with *N*-terminal arginylation (**Fig. S1**). The MS1 spectra of isotopic products were extracted to validate the Arg^10^/Arg^0^ pair features in doublets (“1+1” = 2 isotopic peaks, error ≤ 10 ppm) and summarized into a boxplot for relative intensity overviews (**Fig. 1d**). The co-eluting isotopic peaks indicated unbiased identification of arginylation. The average value of H/L intensity ratios calculated from individual MS1 spectra was considered the H/L ratio in an LC-MS run. The isotopic labeling of the standard peptide was replicated 4 times and produced consistent ratio profiles (**Fig. 1e**). An increasingly stringent MS1 pair detection using co-eluting quartet (“2+2” = 4 isotopic peaks, error ≤ 10 ppm) and sextet (“3+3” = 6 isotopic peaks, error ≤ 10 ppm) yielded H/L ratios closer to hypothetical 1.0 than doublet (“1+1” = 2 isotopic peaks) but gave decreased numbers of MS1 pairs in all replicates. To ensure all MS1 pairs are included, we decided to use MS1 doublet as the threshold filter and include duodecet (“6+6” peaks) for pair overview as shown in **Fig. 1d**. In addition, when the isotopically labeled peptides were mixed at other ratios (R^10^/R^0^-peptide = 1:2 and 7:10) than 1:1 used previously, experimental H/L ratios around 0.5 and 0.7 were observed expectedly (**Fig. S6-7**), another approach to confirm the labeled peptides. For the convenience of experiments and data analysis, an isotopic 1:1 mix strategy with a hypothetical H/L ratio of 1.0 is used in the rest of this study.

### Development of in-house software for dataset analysis

Encouraged by the proof-of-concept data, we then aimed to develop a computational tool “ArginylomePlot” (**Fig. 2a**) with several capabilities to facilitate handling datasets from future complex samples. The software should be able to 1) filter out MS2 pairs with the same sequence modified by Arg^10^ and Arg^0^ (Δmass = 10.008269 Da) disregarding other modifications (*e*.*g*., Met oxidation and Cys carbamidomethylation), 2) extract their MS1 scans from raw data, 3) summarize heavy/light ratios of each arginylation peptide, and 4) export paired MS1 and MS2 scans for data visualization and indexing. The software is publicly available to download on GitHub (https://github.com/BeckyHan/Garcia-Lab/tree/main/ArginylomePlot). Briefly, mzXML data converted from raw data was used as the source input data, and search results from proteomics software (*e*.*g*., Byonic) were used as peptide input. Once a co-elution MS2 pair was identified from the peptide list, their MS1 information (*e*.*g*., *m/z* and charge) was used to extract pairs from mzXML for ratio calculation. To present high-confidence proteomics data, peptide score (H or L ≥ 300) was applied as a filter before MS2 spectrum export. In the end, the software also exports spectra as Excel files for establishing a website database. An exemplary dataset and detailed instructions to execute the software are provided (**Supplementary Dataset 1**).

**Figure 2.**
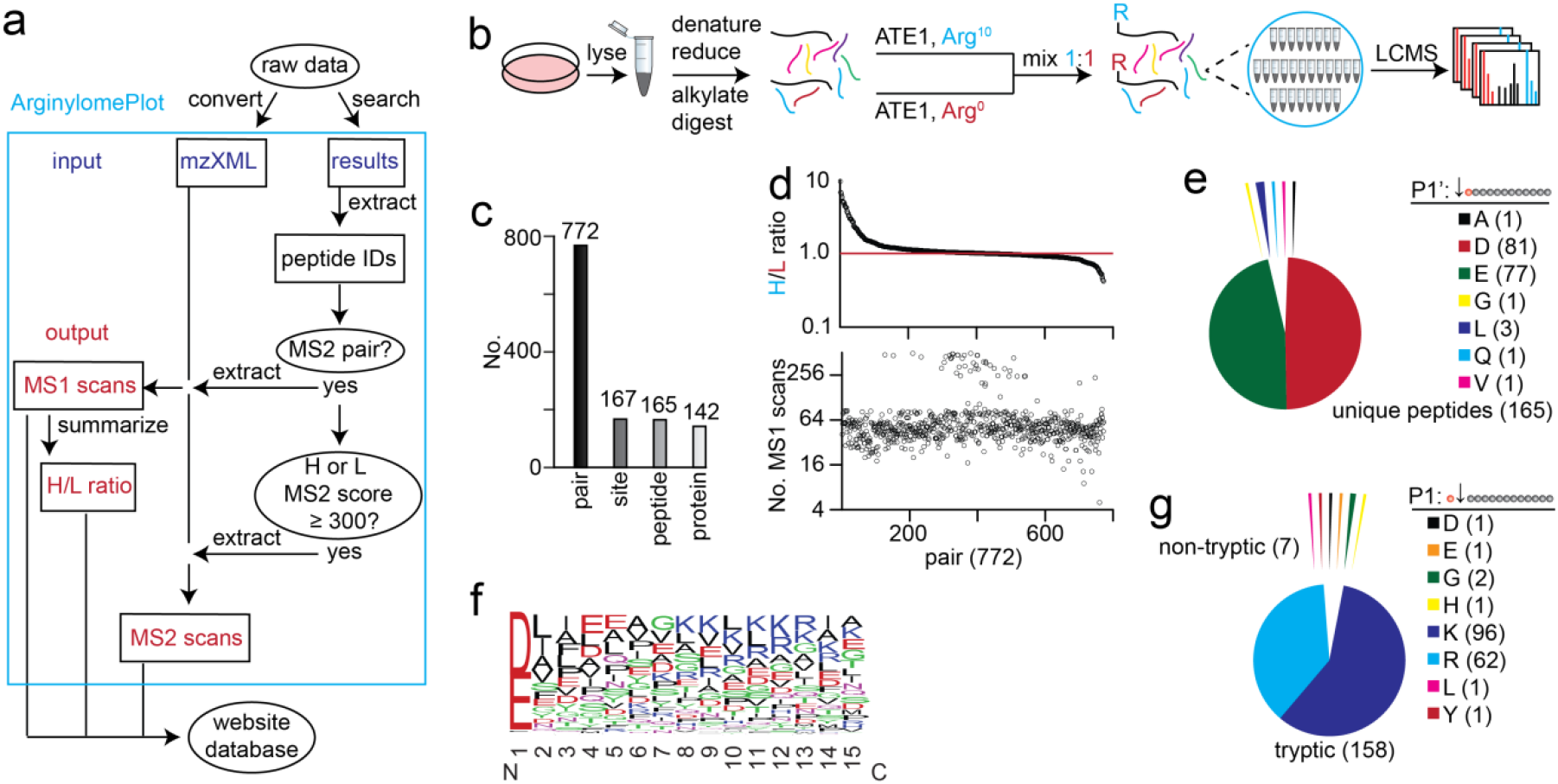
Arginylation analysis of whole-proteome peptides using in-house software. **a**, software flowchart of customized “ArginylomePlot”. Input data are mzXML and peptide identification files. Output data are MS1, MS2, and the summary of unbiased arginylation sites. **b**, experimental workflow for arginylation of tryptic peptides from HEK293T (ATE1 KO) cells. **c**, numbers of identified MS1 pairs, arginylation sites, unique peptides, and unique proteins. **d**, H/L ratio distribution of all MS1 pairs and their corresponding numbers of MS1 scans (doublets, error ≤ 10 ppm). The retention times of peptide IDs belonging to a pair were averaged. The software will extract matching MS1 scans within 1.25 min (± 1.25 min) of the averaged retention times. **e**, analysis of the *N*-terminal residues of all unique arginylated peptides. Q is arginylated after deamidation (Q→E). **f**, sequence logo calculated from unmodified forms of all unique arginylated peptides. The frequency plot is generated by WebLogo. The peptide sequences were aligned and extended to include 14 positions downstream of the arginylation sites as the P1’ position. **g**, comparison of tryptic and non-tryptic *N*-term of all arginylated peptides. Non-tryptic *N*-term may indicate endogenously exposed *N*-termini of proteins.

### Isotopic arginylation of HEK293T peptides by ATE1 *in vitro*

Assisted by the software, we next tested the arginylation of a whole-proteome peptide mixture prepared from the whole proteome of HEK293T cells with ATE1 knocked out to avoid potential interference from endogenous arginylation (**Fig. 2b**). ATE1 KO has been confirmed by genomic sequencing and Western blotting (section Methods, **Supplementary Information**). After data analysis, a total of 772 arginylation pairs belonging to 167 unique arginylation sites were detected at the MS1 level by the software (**Fig. 2c, Fig. S8, Table S1, Supplementary Dataset 2**). The data showed exclusive *N*-terminal arginylation (**Supplementary Dataset 2**), even though we searched for both *N*-terminal and sidechain arginylation at the same time. While charges 2 (83.7%) and 3 (13.8%) account for 97.5% of all identified peptides, most of the arginylated peptide pairs (98.3%) are detected at charge states 2 (47.3%) and 3 (51.0%) with a small portion in charge 4, indicating a charge shift for arginylated peptides (**Fig. S9**). The distribution profile of H/L ratios (threshold: 0.1 ≤ ratio ≤ 10) from all MS1 pairs is centered at 1.0 (**Fig. 2d**) which is the hypothetical ratio for each labeled peptide per experimental design, demonstrating the success of the workflow for complex peptide sample. The doublet ratios were also compared with the quartet and sextet profiles (**Fig. S10**). When a peptide pair was observed with the same charge state and a Δmass of 10.008269 Da, the averaged retention time of relevant peptide IDs containing both Arg^10^ and Arg^0^ was used to create a ± 1.25 min window for MS1 paired scans extraction. MS1 scans containing doublets (10-ppm error) were exported. The average value of H/L intensity ratios calculated from individual MS1 scans was considered the H/L ratio of a peptide pair. The numbers of MS1 scans with the doublet feature used for generating respective H/L ratios are provided (**Fig. 2d, Fig. S11**). When looking at the arginylated *N*-term residues of all unique peptides, a majority of arginylation happened on D and E (including Q→E) residues (159 out of 165, 96.36%) (**Fig. 2e**) while the data also suggested possible arginylation on other *N*-term residues (*e*.*g*., A, G, L and V), consistent with reported specificities and activities of ATE1^30,31^. We then generated the sequence logo of unique peptides using WebLogo 3.0^32^ (**Fig. 2f**) and the arginylation motif using pLogo^33^ (**Fig. S12**). The result suggested that arginylation favors *N*-term D and E residues significantly, while subsequent nonpolar residues (I, L, F, V and A) and acidic residues (D and E) may facilitate the peptide *N*-term arginylation (**Fig. 2f, Fig. S12**). While most of the unique peptides (158 out of 165, 95.76%) are tryptic after K or R cleavage, a few non-tryptic peptides were arginylated (**Fig. 2g**). For example, the arginylation of SSBP E17 was on a non-tryptic peptide (**Fig. S13**), this may have resulted from endogenous protein *N*-term processing and cleavage^34^. This observation demonstrates the potential application of our peptide workflow for the discovery of the cleavage and arginylation of endogenous protein *N*-term.

### Isotopic arginylation of protein substrates by ATE1 *in vitro*

We then moved forward from peptide to protein arginylation using CALR since it is known for *N*-term arginylation at the E18 site after endogenous cleavage of a signaling peptide (AA 1-17)^18^. Purified 18E-CALR after overexpression in HEK293T ATE1 KO cells was isotopically labeled and yielded the H/L ratio of 1.0 in the arginylated E18 peptide (R^10^/R^0^-E^18^PAVYFK) (**Fig. 3a** and **Fig. S14**). The *in vitro* arginylation efficiency of CALR is preliminarily estimated to be 64.7% for the E18 site based on MS2 spectrum counts of modified (R^10^/R^0^-EPAVYFK, counts = 33) and unmodified (EPAVYFK, counts = 18) peptides. When counting the unmodified peptide (EPAVYFK) in samples with and without ATE1, the arginylation efficiency was estimated to be 80.2% (*n* = 3, **Fig. S15**). Replication of CALR arginylation (*n* = 5) showed consistent ratios near 1.0 (**Fig. 3a** and **Fig. S16**). In addition, arginylation results (*n* = 5) of a commercial 18E-CALR (Abcam, ab276554) showed consistent E18 labeling (**Fig. S17**). In comparison, the 18R-CALR protein was purified and tested, and the results (*n* = 5) showed nearly no R^10^ arginylation due to the absence of an open *N*-term E18 residue (**Fig. S18**). The results indicated that the isotopic arginylation strategy could be used to discover modification sites in pure proteins.

**Figure 3.**
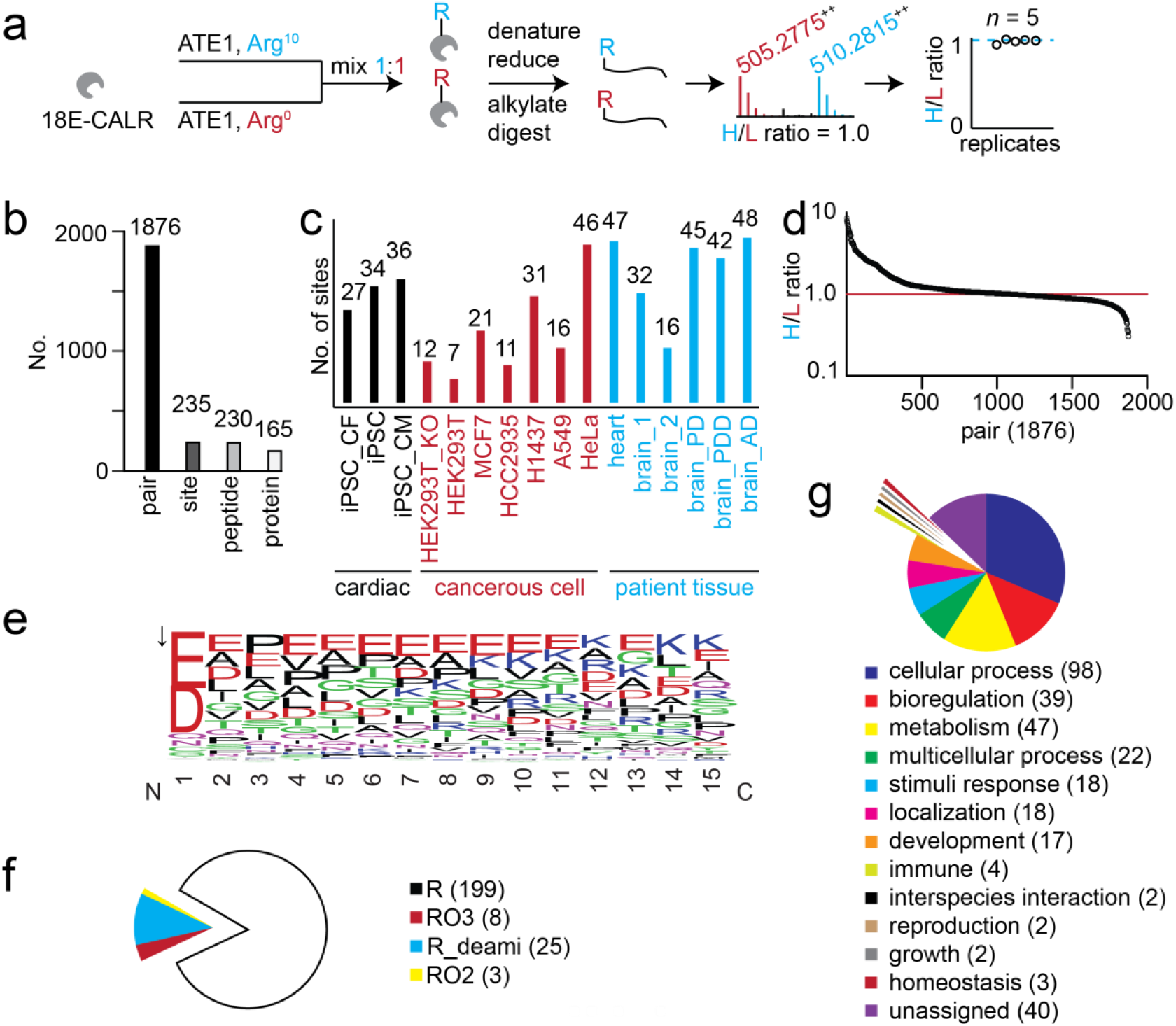
Summary of arginylation sites and ATE1 substrates in human proteins and proteomes. **a**, experimental workflow for CALR arginylation. **b**, numbers of identified MS1 pairs, arginylation sites, unique peptides and unique proteins in human cells and patient tissues. **c**, overview of arginylation sites in human cells and patient tissues. **d**, ratio plot of all MS1 pairs detected in all sample fractions. **e**, sequence logo calculated from unmodified forms of all unique arginylated peptides. Frequency plots are generated by WebLogo. The arrow indicates the cleavage site before arginylation. **f**, arginylation type comparison of all unique sites. **g**, biological function analysis of ATE1 protein substrates using PANTHER. iPSC, induced pluripotent stem cell. CF, cardiac fibroblasts. CM, cardiomyocytes. R, arginylation. RO3, Cys tri-oxidation and arginylation. RO2, Cys di-oxidation and arginylation. R_deami, N/Q arginylation after deamidation. PD, Parkinson’s disease. PDD, PD with dementia. AD, Alzheimer’s disease.

### Arginylation profiling in human proteomes

Using this established platform, we profiled 14 human samples after cell/tissue lysis and ATE1 assay *ex vivo*, a key to installing isotopic Arg post-translationally under ribosome-free conditions to bypass translational Arg incorporation and bias. A total of 1876 isotopically labeled pairs belonging to 235 unique arginylation sites on 165 proteins were identified (**Fig. 3b, Table 1, Supplementary Dataset 3**, and **Fig. S19-20**). As a negative control, the HEK293T sample without the addition of ATE1 didn’t show any arginylation site (RAW data is available). Only 7 arginylation sites (ACTC C259 RO2, BI1 N9 R_deami, HBA N79 R_deami, HMSD Q43 R_deami, PSD12 E348 R, S10AE C74 RO3, and SCN1A C1588 RO2) were assigned as sidechain arginylation accounting for 3% of all unique sites, while the rest 97% of sites were identified as *N*-term arginylation. Similar to results from HEK293T peptides, most of the peptide pairs are detected at charge states 2 (45.0%) and 3 (49.9%), with small portions in charges 4 (4.9%) and 5 (0.2%) (**Fig. S21**). As cross-validation, CALR E18 and PDIA D18 are among the top high-frequency sites with 185 and 135 detections (MS1 pairs) respectively, consistent with the literature on their confirmed E18^10^ and D18^24^ arginylation. The number of unique arginylation sites from each sample is listed in **Fig. 3c** which shows an average of 29 sites/sample could be identified. The biological samples cover induced pluripotent stem cells (iPSC), cardiac cells derived from the iPSC, commonly cultured cell lines, and clinical patient tissues, demonstrating the potential applicability of this method to a wide range of other human proteomes. The H/L ratios (threshold used: 0.1 ≤ ratio ≤ 10) of MS1 pairs are distributed symmetrically and perfectly centered at the hypothetical ratio of **1.0** (**Fig. 3d**). Endogenous Arg^0^ at µM level may interfere with the Arg^10^ labeling, however, the final concentration of Arg^10^ is at 2 mM thus Arg^0^ interference on final H/L ratios might be minimal. Since Arg^0^ and Arg^10^ labeling by ATE1 was carried out separately on complex whole proteomes, deviations of H/L ratios (range: 0.3-8.5) from the hypothetical ratio of 1.0 were expected which might be introduced from the multi-step preparation procedures including ATE1 arginylation on substrates in the whole proteome, H/L mixing, digestion, and peptide fractionation (**Fig. 1b**). The doublet ratios were also compared with the quartet and sextet profiles (**Fig. S22**). The numbers of MS1 scans with the doublet, quartet, and sextet features for generating respective H/L ratios are provided, quartet and sextet pairs showed lower numbers of qualified MS1 scans than doublets based on their trendlines (**Fig. S23**). The sequence logo (**Fig. 3e, Fig. S24**) and arginylation motif (**Fig. S25**) from all unique peptides suggested that arginylation favors *N*-term D and E residues significantly, while subsequent nonpolar residues (A, L, V, and P) and acidic residues (D and E) may facilitate the peptide *N*-term arginylation. Interestingly, all four types of arginylation based on different modification masses (**Fig. S26**) have been detected including arginylation (R, 84.7%), Cys tri-oxidation arginylation (RO3, 3.4%), N/Q deamidation arginylation (R_deami, 10.6%), and Cys di-oxidation arginylation (RO2, 1.3%), (**Fig. 3f**). Comparison of arginylation sites between iPSC, cancerous cells, and patient samples showed only 32 (13+11+8) shared sites, while the remaining majority of sites are unique to their respective cell/sample groups (**Fig. S27**). Comparisons between individual samples are listed in **Fig. S28** (cardiac), **Fig. S29** (brains), and **Fig. S30** (cancerous), indicating unique and shared arginylation sites from individual samples. Replication (*n* = 3) analysis of 4 samples showed that most arginylation sites in each sample were detected in replicates (**Fig. S31**), indicating the repeatability of the profiling method. Among 165 proteins identified, 162 protein targets are involved in many key biological processes (*e*.*g*., cellular process, biological regulation, and cellular metabolism) according to the PANTHER classification (**Fig. 3g**).

**Table 1.**
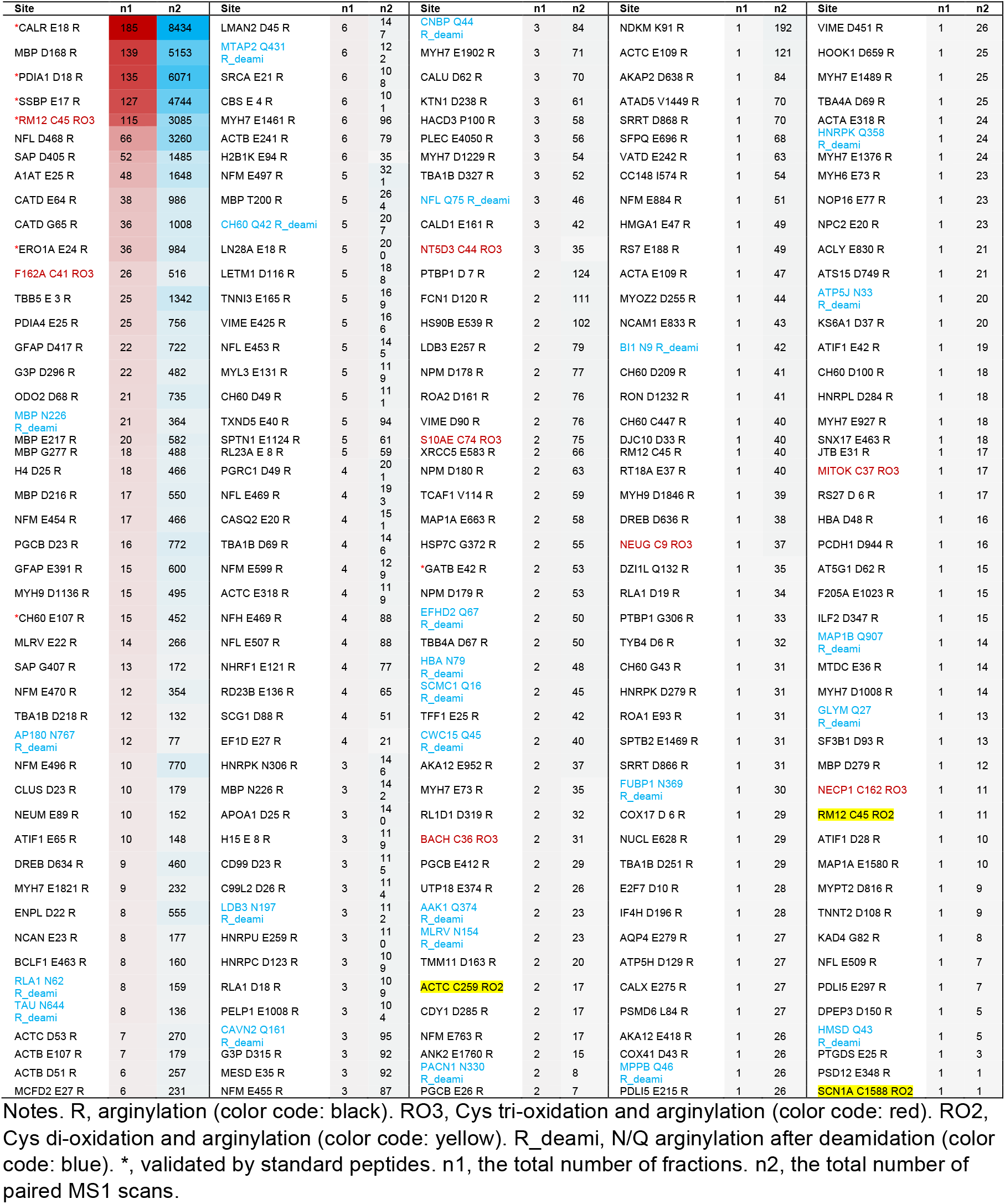
Number of detections in isotopic pairs and MS1 scans for each arginylation site from human proteomes.

### Validation of representative arginylation sites using standard peptides

We aimed to validate representative sites using synthetic peptides (*e*.*g*., REPAVYFK for CALR E18 peptide) whose MS behaviors (MS1 spectra in **Fig. 4a** and MS2 spectra in **Fig. S32**) matched those in our MS data. The chosen sites are a mixture of known (CALR and PDIA) and unknown, and they represent high-frequency (CALR, PDIA, SSBP, and RM12), mid-frequency (ERO1A and CH60), and low-frequency (GATB) sites in our data from biological samples (**Table 1**). Such a validation added extra confidence to the Arg10 arginylation which is an unnatural process and thus can be considered an internal validation of our workflow. For example, H/L ratio = 1.0 for RM12 C45 arginylation (**Fig. 4b**) eliminated the possibility of tryptic missed cleavage between Arg44 and tri-oxidized Cys45, standard peptide confirmed this minor type of arginylation with co-presence of Cys trioxidation^1^ (**Fig. S32**). It is noteworthy that the low frequency of a site doesn’t necessarily mean low confidence since most of the reported sites are based on many isotopic co-eluting MS1 scans (**Fig. S23** and **Supplementary Dataset 3**) and paired MS2 spectra (**Supplementary Dataset 4**). To further confirm that these representative sites possess arginylation at the *N*-termini but not on the side chain of *N*-terminal residues, we purchased peptides containing arginylation on the side chain [*e*.*g*., E(R)PAVYFK for CALR E18 peptide]. Analyzed by LCMS, side chain arginylation peptides may behave similarly (*e*.*g*., CALR E18 peptide) or differently (*e*.*g*., SSBP E17 peptide) with *N*-terminal arginylation peptides (**Fig. S33**). Interestingly, arginine residue on the side chain produced a signature ion at *m/z* 175.1190 when the peptide didn’t end with a *C*-terminal R residue [*e*.*g*., E(R)PAVYFK]. When *C*-terminal R residue was present on side chain arginylation peptides [*e*.*g*., E(R)EQPPETAAQR], both R residues could produce the same ion at *m/z* 175.1190 with a higher intensity than that from *N*-terminal arginylation peptides (*e*.*g*., REEQPPETAAQR) (**Fig. S33**). By adding the side chain arginylation peptides after ATE1 assay, the *N*-terminal arginylation was further confirmed for standard peptide (DIAALVHSSGNleS-NH_2_) (**Fig. S34**), CALR pure protein (**Fig. S35**) and CALR in HEK293T cells (**Fig. S36**).

**Figure 4.**
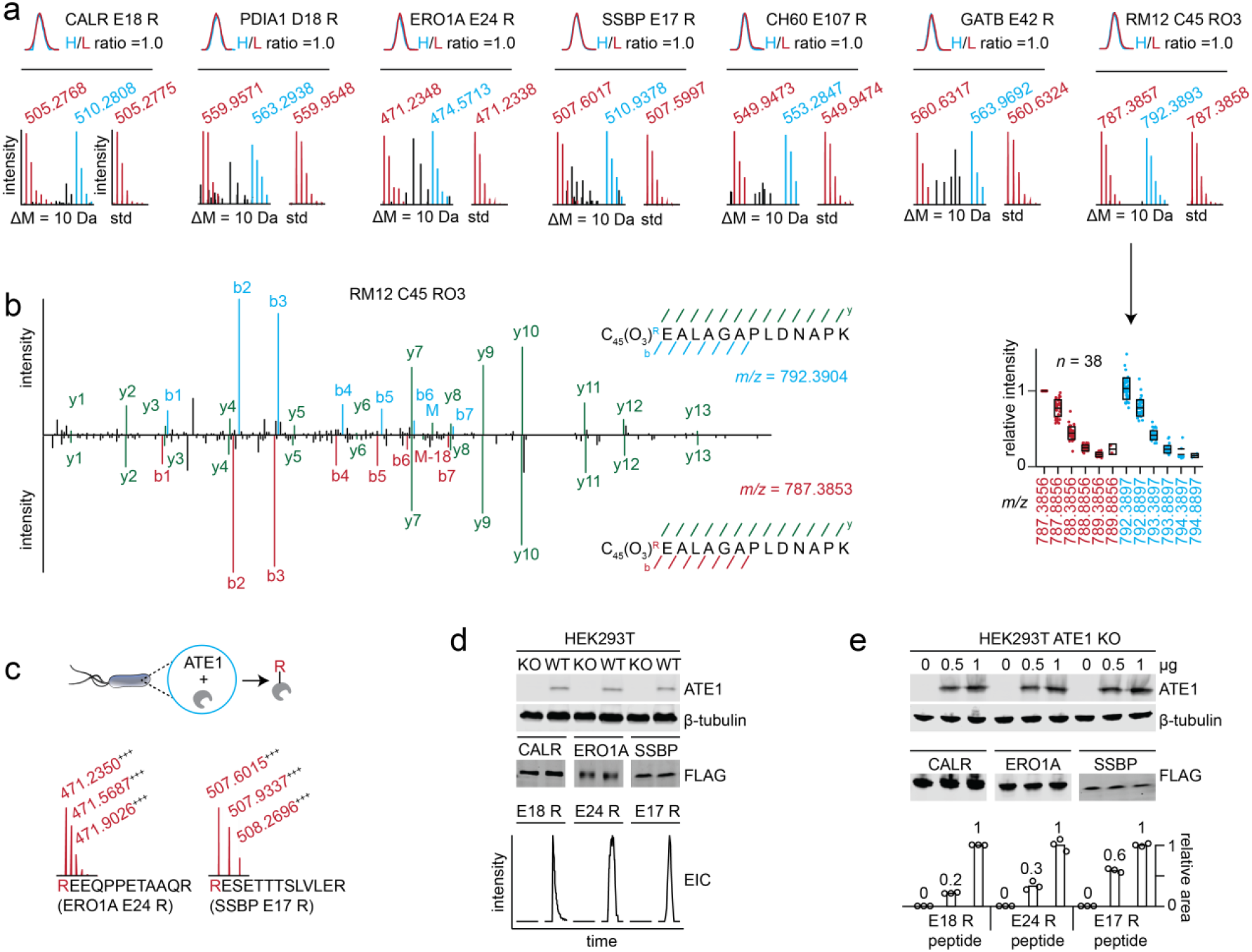
Validation of representative arginylation sites. **a**, synthetic peptide validation of 7 arginylation sites. **b**, representative MS2 spectra indicating RM12 arginylation on Cys45 with tri-oxidation (C45 RO3). **c**, open E24 in ERO1A and E17 in SSBP are arginylated in an in-bacteria arginylation system. Protein and ATE1 are co-expressed in E. coli for in-bacteria arginylation. The protein was purified and digested for proteomics analysis. **d**, arginylation dependency of CALR, ERO1A, and SSBP sites on endogenous ATE1. Anti-ATE1 was used to detect the expression of endogenous ATE1. β-tubulin was used as a loading control. Anti-FLAG antibody was used to detect expressions of CALR, ERO1A, and SSBP proteins. Extracted ion chromatograms (EICs) of arginylated peptide in each protein in WT and KO cells after pulldown and proteomics are provided. **e**, relative arginylation levels of CALR, ERO1A, and SSBP sites after co-overexpression of ATE1. Protein was purified by antibody pulldown experiment followed by proteomics analysis. Peak areas of arginylated peptides were normalized to the sample with the highest signal and relative ratios are displayed. Different amounts (0, 0.5, and 1 µg) of ATE1 plasmids were used for transfection. β-tubulin was used as a loading control. Anti-FLAG antibody was used to detect expressions of CALR, ERO1A, and SSBP proteins. std, standard peptide. KO, ATE1 KO. R, arginylation.

### Validation of representative arginylation sites using in-bacteria arginylation assay

We then selected two novel sites for validating their arginylation in an ATE1-protein co-expression system in *E. coli* reported in a recent study^35^. The two proteins are 1) a chaperone ERO1A in the endoplasmic reticulum (ER), being similar to CALR and PDIA; and 2) SSBP being distinct from ER proteins in molecular size (17 kDa), localization (mitochondria) and function (single-stranded DNA-binding). Briefly, a protein or peptide containing an arginylation site is fused to ubiquitin and co-expressed with human ATE1 and Ulp1 protease in *E. coli*. The expressed protein is cleaved by protease and exposes the *N*-terminal arginylation site for ATE1 arginylation. This system allows for in-bacteria validation of targeted arginylation. The results showed that both ERO1A E24 and SSBP 17E were arginylated (**Fig. 4c** and **Fig. S37-38**). As a bonus, we also detected a further leucylation on top of arginylation (**Fig. S37-38**), a unique process to modify *N*-term Arg due to the presence of the *E. coli* leucyl/phenylalanyl-tRNA-protein transferase (LFTR)^36^. Similarly, we also tested the in-bacteria arginylation of ERO1A and SSBP peptides, whose respective *N*-term E24 and 17E were also arginylated in the co-expression system (**Fig. S39**). The arginylation and leucylation were also observed by top-down proteomics (**Fig. S40-41**). The results concluded that ERO1A E24 and SSBP 17E are arginylation target sites of ATE1.

### Validation of representative arginylation sites in cells

The unbiased nature of the profiling method is owing to the co-elution strategy of isotopic Arg10/Arg0 peptides, however, the revealed sites are based on an *ex vivo* ATE1 assay and thus do not necessarily conclude their occurrences *in vivo* (in cells) where ATE1 is active. To confirm endogenous arginylation activity on the identified targets in cells, we overexpressed the wild-type ERO1A and SSBP in HEK293T cells using CALR as a positive control. Due to the low abundance of endogenous arginylation, arginylated peptides were not detectable in peptide mixtures obtained from whole proteomes if overexpressed proteins were usually not purified by immunoprecipitation. For example, the MS2 spectrum of arginylated peptide for the SSBP E17 site was only detected once from a pulldown sample (**Fig. S42**). Nevertheless, the arginylated peptides corresponding to all three sites (CALR E18, ERO1A E24 and SSBP E17) were identified by bottom-up proteomics after immunoprecipitation of overexpressed proteins (**Fig. 4d** and **Fig. S42**). As a comparison, we also performed the same experiment in HEK293T (ATE1 KO) cells (**Fig. 4d**) or on E-to-V mutants of ERO1A E24 and SSBP E17 (**Fig. S43**), which did not show any arginylation on those sites indicating arginylation dependency on ATE1 and arginylation selectivity on E residue. To further test their dose-dependency on ATE1, we then co-overexpressed the ATE1 and protein substrates in HEK293T (ATE1 KO) cells. The data show the arginylation levels of the tested sites are dose-dependent on the ATE1 expression (**Fig. 4e**). Such dose-dependency was also observed when co-overexpressing CALR and ATE1 in WT HEK293T cells (preliminarily estimated arginylation efficiencies between 0.025%-0.111%) (**Fig. S44**). Both ERO1A^37^ and SSBP^38^ contain cleavage peptides before the arginylation sites (E24 and 17E respectively), similar to that of CALR. Taking advantage of this feature, we further prepared 24R-ERO1A and 17R-SSBP plasmids by inserting an R residue before E into the full-length plasmids as positive transfection controls, the spectra of their arginylated peptides (**Fig. S45**) resulted from overexpression, affinity purification and proteomics further confirmed the arginylation of their wild-type counterparts.

We have additionally validated the endogenous arginylation in HEK293T cells of two more sites (proteins): A1AT E25 arginylation (**Fig. S46**) and Tau N644 arginylation after deamidation (**Fig. S47**). Similar to CALR, ERO1A and SSBP, A1AT has a signal peptide^39^ before E25 thus it is not surprising to observe its E25 arginylation by endogenous ATE1. Tau N644 has been arginylated after deamidation in our data, which has never been observed previously. We also confirmed the arginylation of this site using overexpression of wild-type Tau plasmid in HEK293T cells (**Fig. S47**). We have additionally confirmed the arginylation of the Tau E3 site (**Fig. S47**), which was proposed previously as an arginylation site of ATE1, but not validated for its arginylation^40^. A previous study showed that purified Tau did not show detectable arginylation by ATE1 *in vitro*^16^, possibly due to the lack of exposed arginylation sites (*e*.*g*., E3 and N644 deamidation). Different from Tau E3 as a calpains cleavage product^40^, the mechanism of generating an open N644 followed by deamidation for arginylation remains to be elucidated.

### Biological functions associated with ERO1A arginylation

To investigate whether arginylated ERO1A resides in ER, we did imaging on overexpressed ERO1A and 24R-ERO1A in HEK293T ATE1 KO cells. Both proteins co-localized with ER marker and PDI (**Fig. 5a** and **Fig. S48**), suggesting that 24R-ERO1A was translocated into ER after signaling peptide AA1-23 cleavage. As a positive control, we showed that CALR-halo and its arginylation form co-localized with a GFP-tagged ER marker (**Fig. S49**). Similar to CALR and PDIA, ERO1A is translocated into ER after signal peptide cleavage. A previous study reported the formation of the ERO1A-PDI complex in regulating cellular function^41^, our pulldown data showed that 24R-ERO1A interacted with PDI to a comparable extent with and without PDI co-overexpression (**Fig. S50-51**). We also showed that arginylation of ERO1A maintains its enzymatic activity on oxidative protein folding compared with WT ERO1A as indicated by the disappearance of the reduced JcM band (**Fig. S52**).

**Figure 5.**
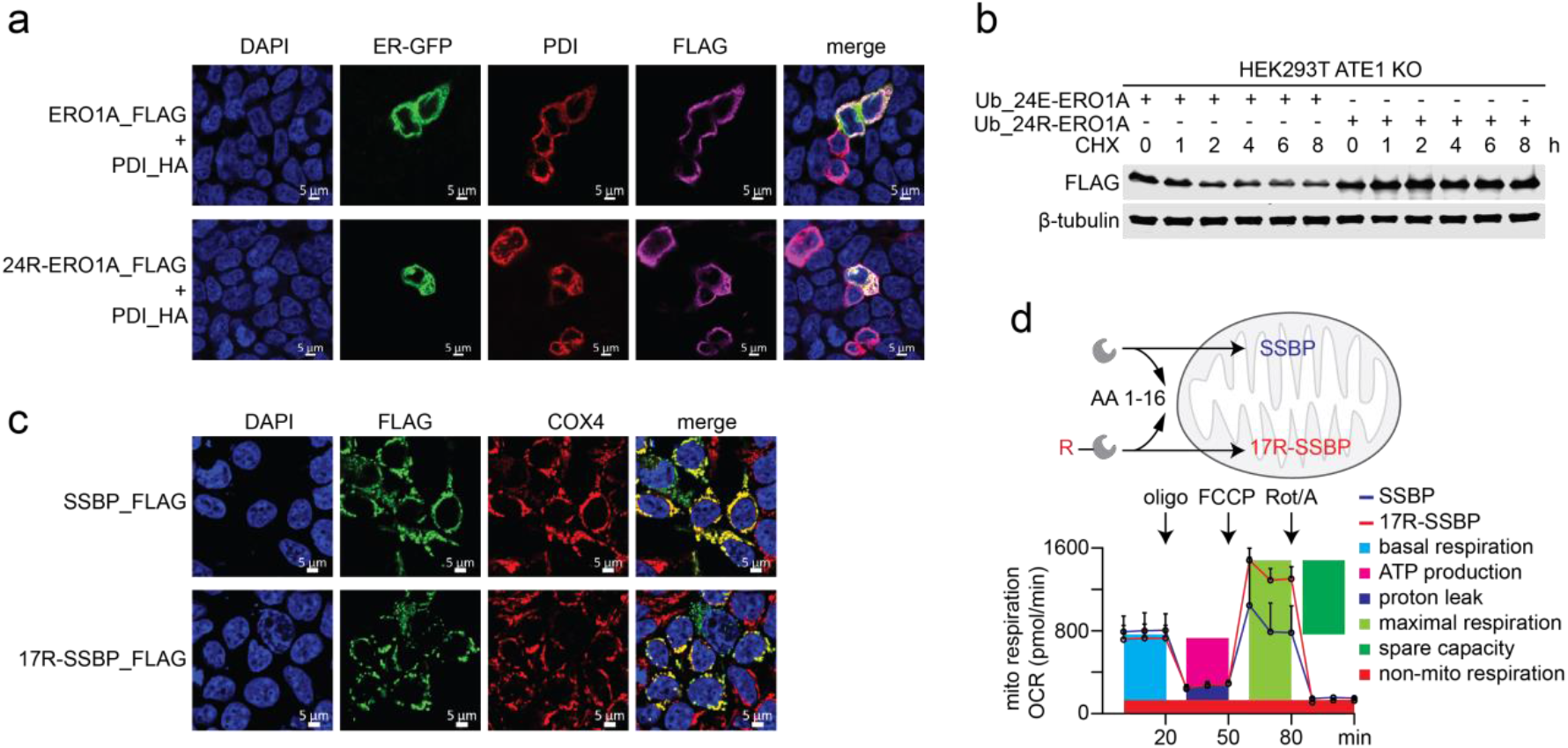
Biological functions of representative arginylation sites. **a**, imaging of ERO1A and 24R-ERO1A compared with ER marker and co-overexpressed PDI. **b**, expression levels of cytosolic ERO1A species in CHX chase experiment. **c**, imaging of SSBP and 17R-SSBP compared with mitochondria protein COX4. **d**, bioenergetic profiles of HEK293T cells after SSBP and 17R-SSBP transfection measured by Seahorse XF24. Mitochondria respiration was analyzed with basal respiration, ATP production, proton leak, maximal respiration, spare capacity, and non-mitochondria respiration. The average values of 3 Seahorse measurements were used for comparison.

To investigate the arginylation effect on ERO1A stability and degradation using HEK293T ATE1 KO cells, we compared the degradation patterns of ubiquitin-ERO1A fusion proteins with and without arginylation (**Fig. S52**) according to similar experiments performed on CALR^42^. Briefly, ubiquitin is cleaved by de-ubiquitinase after protein synthesis to expose E24 and R24 residues in ERO1A (no arginylation on E24) and 24R-ERO1A (∼100% arginylation on E24) respectively. Proteasome inhibitor MG132 treatments after transient transfection increased the levels of ERO1A and 24R-ERO1A, suggesting that both are involved in proteasome-mediated degradation. Similar dose dependence on MG132 was observed in co-treatments when the ribosomal translation of both species was inhibited by cycloheximide (CHX). In addition, 24R-ERO1A showed slightly higher levels after MG132-CHX co-treatment, indicating better stability than ERO1A against proteasomal degradation (**Fig. S52**). To take a closer look at their stabilities, we did CHX chase experiments^42^ where cells were treated with CHX after a 2-day transient transfection. The results showed that 24R-ERO1A was less prone to degradation than ERO1A with clear degradation after CHX treatment (**Fig. 5b, Fig. S53**). Such behavioral differences have been previously observed in cytosolic CALR^42^, suggesting that arginylation of ERO1A may also have a stabilization effect against proteasomal degradation.

### Biological functions associated with SSBP arginylation

SSBP matures by translocation into mitochondria after cleavage of its transit peptide (AA 1-16)^38^. We did imaging on SSBP and 17R-SSBP to investigate whether arginylated SSBP resides in the mitochondria. Both species co-localized with mitochondria protein COX4 (**Fig. 5c** and **Fig. S54**), suggesting that 17R-SSBP was translocated into mitochondria after cleavage of its transit peptide AA1-16. To assess the cytosolic degradation of SSBP with and without arginylation in the *N*-degron pathway, we evaluated its expression levels using a ubiquitin cleavage plasmid system for transfection. As a result, we did not observe an obvious response toward MG132 treatment (proteasome inhibitor) for both SSBP species (**Fig. S55**). CHX (protein synthesis inhibitor) treatment reduced the expression levels of both SSBP and 17R-SSBP compared with cells without CHX treatment, indicating that ribosomal synthesis was contributing to the cellular expression levels of both species. 17R-SSBP expression levels were higher than WT SSBP with and without CHX treatment, indicating a slower turnover of 17R-SSBP than SSBP. The results suggest that the major role of SSBP arginylation might not be degradation (**Fig. S55**).

We then turned our attention to mitochondrial SSBP. SSBP KO in mouse models is embryonic lethal whereas conditional KO in the heart results in cardiomyopathy and reduced life span (∼18 weeks) accompanied by heart respiratory chain deficiency^43^. It is noteworthy that ATE1 KO is also embryonic lethal with heart defects^1^, being similar to SSBP KO. Thus arginylation of SSBP may regulate cells through mitochondria functions. Considering the importance of mitochondria in energy generation and heart functions, we then investigated the functional differences between SSBP and 17R-SSBP on cellular bioenergetics (**Fig. 5d**) based on a previous study on SSBP mutants^44^. Toward this, the Seahorse XF24 Cell Mito Stress Test was performed to measure the arginylation effects of SSBP on mitochondrial respiration before and after sequential addition of oligomycin (oligo), carbonyl cyanide-4- (trifluoromethoxy)phenylhydrazone (FCCP) and rotenone/antimycin A (Rot/A). Analysis of cellular oxygen consumption rate (OCR) showed that HEK293T cells overexpressing 17R-SSBP displayed comparable basal respiration, ATP production, and non-mitochondria respiration than WT SSBP (**Fig. 5d**). Interestingly, 17R-SSBP resulted in significantly higher maximal respiration (*p* = 0.0104) and spare capacity (*p* = 0.0076) than cells overexpressing WT SSBP (**Fig. 5d, Fig. S56, Supplementary Dataset 5**). The relative ratios of maximal respiration and spare capacity in 17R-SSBP are 1.6 and 8.5 compared with WT SSBP (**Fig. S56**), indicating improved mitochondria respiration potential after SSBP arginylation. 17V-SSBP did not display these effects on mitochondria when compared with WT SSBP (**Supplementary Dataset 6**).

### Arginylation profiling in mouse tissues

Our profiling platform has been working in human cells and tissues, to test its general applicability we next applied it to mouse tissues. We acknowledge that it is not ideal to use human ATE1 to arginylate mouse proteomes, although such a cross-species approach has been previously used^16^. Results from three tissues (lung, heart, and brain) showed that the platform is applicable to arginylation discovery from mouse proteome (**Supplementary Dataset 7**). A total of 14 sites from 69 MS1 pairs were identified (**Fig. S57**). Interestingly, CALR D18 and A1AT1 E25 are the two shared arginylation sites across three tissues. We noticed that the mouse A1AT1 E25 site was identified by a different peptide sequence (EDVQETDTSQK) (**Fig. S58**) from the human A1AT E25 site (EDPQGDAAQK) (**Fig. S46**). In addition, human CALR E18 and mouse CALR D18 sites were also identified from species-dependent sequences (EPAVYFK and DPAIYFK, respectively) (**Fig. S59**), indicating that mouse and human sites can be used to cross-validate each other when overlapped.

### Establishing a public database for arginylation sites and ATE1 substrates

We envision that the arginylation sites discovered from the various samples will greatly advance the arginylation and *N*-degron fields. The fact that our high-frequency hits (CALR, PDIA, and TAU *et al*)^25,40^ are among the most intensively studied arginylation proteins instilled confidence to follow the biology of new sites. This represents a major advantage of our innovative method compared to existing profiling methods^21,28^. Meanwhile, we understand that we alone cannot follow up on all the targets/sites for biological functions, a publicly accessible database/website (www.arginylation.com, **Fig. 6**) was thus formulated to allow everyone to examine our data and follow the biology of their arginylation targets of interest. The isotopic MS1 and MS2 scans discussed in this work from standard peptides, whole-proteome peptide mixtures, pure proteins, human proteomes, and mouse proteomes are included on the website. The source code for data visualization is publicly available to download at https://github.com/ChenfengZhao/Arginylation. The website was designed to allow the following two capabilities, 1) users can search protein names, accession numbers, sample types, species, and peptide sequences of their interest; 2) users can manually select species, sample types, proteins, peptides, fractions of a sample, and charge states of the peptides from our indexed database (**Fig. 6**). As a result, all isotopic MS1 scans, H/L ratio summaries, and H/L MS2 scans with default settings are displayed on the website. To allow users to change our default parameters, we enabled changes in *m/z* errors and MS2 ion types. The most important feature of our website is that users can download our indexed data (MS1 and MS2 spectra in Excel format) based on species (*e*.*g*., human and mouse) or samples (*e*.*g*., iPSC or patient brain).

**Figure 6.**
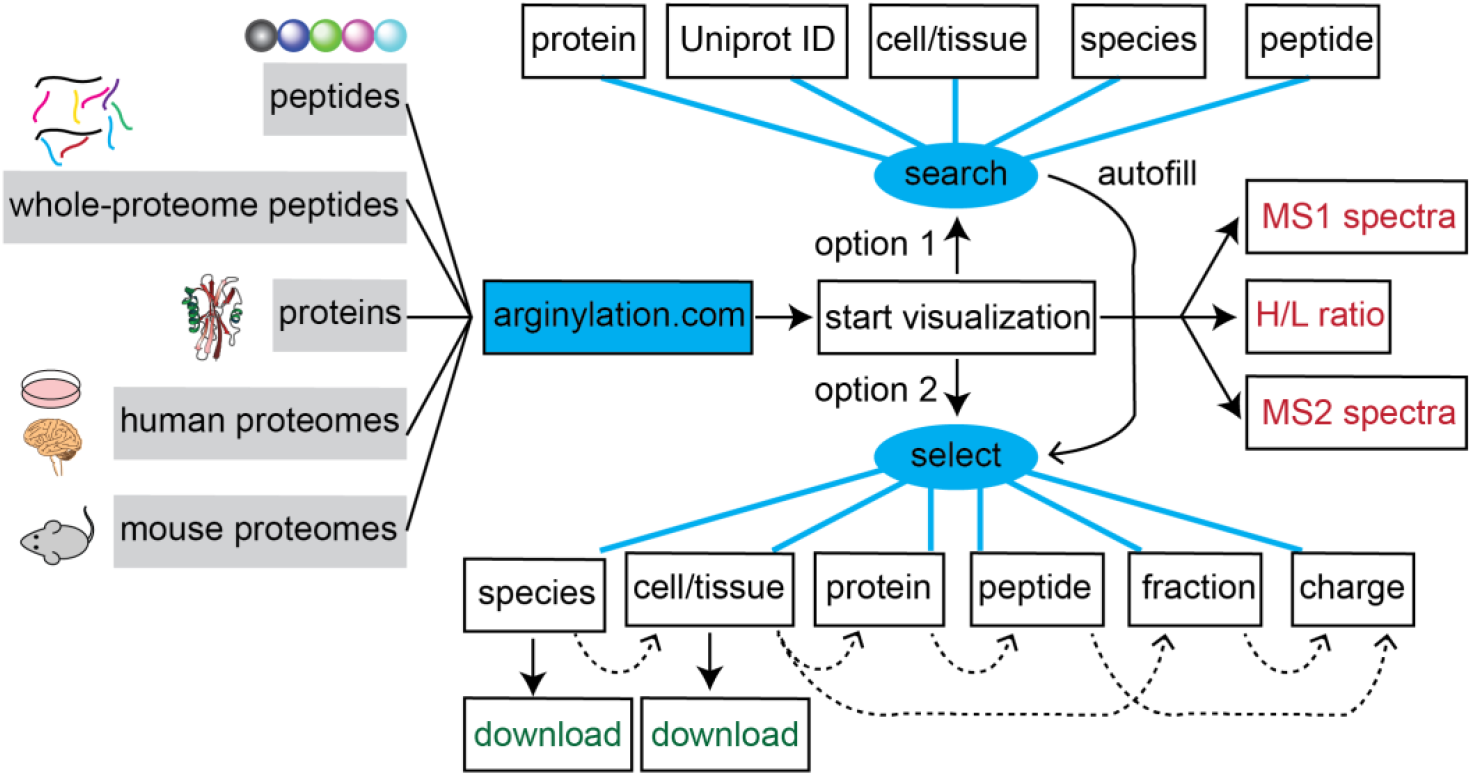
Design and flowchart of the arginylation database and its website. Arginylation data are from peptides, whole-proteome peptides, proteins, human proteomes, and mouse proteomes. Visualization includes isotopic ratios and annotated isotopic MS1 and MS2 spectra. All indexed MS1 and MS2 scans are accessible to the public to download. A dashed arrow means a one-to-multiple inclusion relationship.

## Discussion

ATE1 is the only enzyme known to catalyze arginylation in mammalian systems (*e*.*g*., human and mouse), it has two isoforms in humans (ATE1-1 and ATE1-2) and four isoforms in mice (ATE1-1, ATE1-2, ATE1-3, ATE1-4). (**Fig. S60**). The expression levels and enzymatic functions vary among different isoforms^24,45^. While our work mostly focuses on establishing a general proteomic platform using human ATE1-1, using other isoforms (ATE1-2 for human proteomes, and mouse isoforms for mouse proteomes) for arginylation discovery is a necessary next step to cover the ATE1 substrate scope comprehensively. We preliminarily compared human ATE1-1 and ATE1-2 for arginylation profiling in two samples (patient brain and HeLa cells). ATE1-1 and ATE1-2 sites showed unique and shared sites, more arginylation sites were revealed by ATE1-1 than ATE1-2 for both samples (**Fig. S61-62**), indicating that ATE1-1 may have a slightly broader substrate scope for such an *ex vivo* arginylation assay. In addition, previous studies showed that the expression of ATE1 isoforms is tissue-specific and cell-dependent^24,45^, thus the expression levels might be a significant consideration when choosing ATE1 isoforms for arginylation profiling experiments.

It was initially believed that arginylation occurred only on exposed *N*-terminal Asp, Glu and Cys residues resulting in protein degradation by the ubiquitin-dependent *N-*degron pathway^46^. Later studies identified proteins (*e*.*g*., α-synuclein^16^) potentially modified by arginylation on their sidechain Asp and Glu residues^27^, resulting from a different but minor catalysis mechanism of ATE1. Our arginylation sites were identified using a search algorithm considering both *N*-term and sidechain arginylation situations (**Fig. S26**). The resulting sites from the whole-proteome peptide mixture (**Fig. 2**) and proteomes (**Fig. 3**) are in agreement with the current understanding that ATE1 predominantly arginylate protein/peptide *N*-term C (oxidized), D, and E residues, while other residues and sidechains can be arginylated less preferably. Since the ATE1 assay was performed on proteomes, we believe most sites were arginylated as protein *N*-term (**Fig. 3** and **Supplementary Dataset 3**). A preliminary analysis of the P1 residues (protease cleavage site) in arginylated peptides showed potential protease cleavage after each of the 20 amino acid residues (**Fig. S63** and **Supplementary Dataset 3**). Cleavages at K and R sites accounting for 18.6% and 18.2% respectively might be produced from endopeptidases such as trypsin-like proteases, in contrast, how cleavages at A (13.3%) happened remains unanswered. While some sites (*e*.*g*., CALR E18, ERO1A E24, and SSBP E18) were known to be protein *N*-term based on the current knowledge on cleavages of signaling peptides and transit peptides, many sites (*e*.*g*., TAU N644) are left unexplained as to how the parent proteins were cleaved to expose the *N*-term sites for ATE1 arginylation. Previous studies have offered some insights into protease cleavage (*e*.*g*., calpains for TAU E3^40^ and caspase 3 for CDC6 D101^25^) for *N*-term arginylation, however, the cleavage products of more than 600 proteases remain elusive. Great efforts are needed to understand the interplay between proteolytic cleavage and arginylation.

Proteomic discovery of endogenously low-level arginylation from cells or tissues is challenging^21,28^. This is largely due to 1) the mass difference (+156 Da) introduced by arginylation being the same as the normal arginyl residue from a protein sequence, making label-free proteomics biased to predefined arginylation sites by nature; and 2) efficient enrichment (purification) methods are lacking. The fact that arginyl-tRNA is the same source of arginylation and ribosomal translation makes unbiased endogenous arginylation discovery almost impossible since one can’t exclude ribosomal incorporation when cells are alive^13^. Our initial effort using Arg10 and Arg0 for pulse labeling of arginylation in HEK293T cells yielded a putative list of sites (**Fig. S63**). Additional efforts using cycloheximide to stop protein synthesis, MG132 to accumulate arginylation, and peptide fractionation to boost peptide IDs did not seem to increase our confidence. Arginylation identification from a standard proteome-wide peptide mixture is not ideal. Our efforts on *in vivo* strategy also suggested that endogenous arginylation should have minimal contribution to the light (Arg^0^) labeling in our *ex vivo* isotopic approach. Previous studies tried to use anti-arginylation antibodies to reduce sample complexity and enrich endogenous arginylation^21,28^, however, these efforts by design were biased toward non-degradative ATE1 targets and high-abundance proteins. Sites from those studies showed nearly no overlap with those previously characterized by the Varshavsky group as degradation-inducing arginylation substrates^13^. We thus turned our attention to an *ex vivo* ATE1 enzyme approach using cell lysates, which might be less physiologically relevant than those targeting endogenous arginylation. In addition, arginylated proteins by *ex vivo* assay using lysates were not localized correctly in cell compartments, thus may be false positive hits (not arginylated in cells). However, due to its unbiased nature, this method represents a superior strategy for arginylation discovery.

This profiling method relies on deep peptide fractionation followed by LC-MS/MS analysis to discover the low-abundance arginylated peptides, and intact peptides that are not of interest (accounting for >99.99%) are obvious interfering signals in proteomics data. The reduced signal complexity of each peptide fraction after fractionation is key to identifying arginylated pairs from proteome-wide samples. The additional charge in ionization (**Fig. S9**) and polarity in chromatography of peptides (**Fig. 1c**) resulting from arginylation may help validate peptide identity using LC-centric prediction tools such as DeepLC (**Supplementary Dataset 8**)^47^ and Chronologer^48^. Without enrichment, the sites revealed by our method could be dependent on the expression levels of their parent proteins. We envisioned enrichment approaches are essential to achieve improved sensitivity and efficiency in arginylation site detection. Since ATE1 mainly installs Arg to *N*-termini of substrates, some technologies targeting *N*-term peptides might be helpful for arginylation enrichment. Examples are TAILS (terminal amine isotopic labeling of substrates) technology^49^ and iNrich (integrated *N*-terminal peptide enrichment) method^50^ which are currently under our investigation as future directions. The anti-arginylation antibody enrichment is also under consideration, one potential advantage of such an antibody enrichment strategy is that antibodies can be used for enriching *in vivo* arginylated proteins. However, antibody enrichment may require scaling up the ATE1 reaction to milligram proteomes from the current 20-µg scale. Taken together, our *ex vivo* ATE1 approach provides a robust, unbiased way to map arginylation at the proteome-wide scale, setting the stage for more systematic and comprehensive analyses. Looking ahead, combining this strategy with targeted enrichment techniques and in vivo validation should significantly advance our understanding of arginylation’s functional roles in diverse biological contexts.

## Methods

The methods section is provided in the Supplementary Information with details available.

## Acknowledgments

Z.L. thanks an NIH R21 (CA292191) and R01 (HL177113), a WUSM ICTS JIT grant (JIT1181), and support from the Research Education Component through an NIA P30 grant (AG066444). B.A.G. thanks NIH (HL177113, NS111997 and HD106051) and NSF (CHE 2127882) grants. D.L. thanks an NCI R21 grant (CA286307) and the support of the Mays Cancer Center Early Career Pilot Award from CCSG (P30 CA054174). Y.Z. thanks an NIH R35 grant (GM150678). M.J.G. thanks partial support from NIH (R01 HL141086), Children’s Discovery Institute of Washington University and St. Louis Children’s Hospital (PM-LI-2019-829), and the American Heart Association (970198). A. K. thanks NIH R35 (GM122505) and R01 (NS102435) grants. The authors thank Professor Natalie Niemi (WashU) for guidance on the Seahorse assay, Professor Xia Liu and Professor Guangyong Peng (WashU) for providing mouse tissues, and Washington University Diabetes Research Center supported by NIH Grant 5P30 DK020579.

## Author contributions

Z.L. and B.A.G. conceived the project. Z.L., Y.X., and J.G. performed proof-of-concept experiments. Z.L., Y.X., J.G., X.Liu, E.Z., B.B.P., D.H.R., R.M.S., F.N.V., R.K., G.P.D., B.M., X.Lan, D.F., and L.G. performed experiments. Z.L., Y.X., J.G., X.Liu, E.Z., B.B.P., D.H.R., R.K., B.M., X.Lan, D.F., L.G., Y.Z., K.J.L., M.J.G., D.L., A.K. and B.A.G. analyzed data. Z.L. wrote the draft. C.Z. wrote Python codes and established the website, and X.H. wrote R codes and ArginylomePlot software. K.J.L., B.M., and A.K. provided patient tissues. All authors wrote, revised, and approved the manuscript.

## Competing interests

Z.L., D.L., and B.A.G. are co-founders of LasNova Therapeutics, LLC.

## Data availability

Arginylation site data are publicly available to view and download from www.arginylation.com (the website will be accessible during peer review and after publication, it will be inaccessible during revision).

RAW data are uploaded to MassIVE under dataset numbers MSV000097195, MSV000097197, and MSV000097196. The data index is listed in **Supplementary Dataset 9**.

Log in information for reviewer: MSV000097195_reviewer, MSV000097197_reviewer, and MSV000097196_reviewer, password: xxx.

Data supporting the findings in this study are available within the paper. Any associated data are available from the corresponding authors upon request.

## Code availability

Python codes for website construction are available to download from GitHub (link: https://github.com/ChenfengZhao/Arginylation)

R codes for ArginylomePlot software are available to download from GitHub (link: https://github.com/BeckyHan/Garcia-Lab/tree/main/ArginylomePlot).

The arginylation website is publicly accessible (link: www.arginylation.com) for data visualization and download.

## Notes

### Summary of Updates

This revision addressed reviewers' comments, added more data, and more supporting information.

