## Supplemental Information for "An Unbiased Proteomic Platform for ATE1-based Arginylation Profiling"

### Supplementary figures and tables

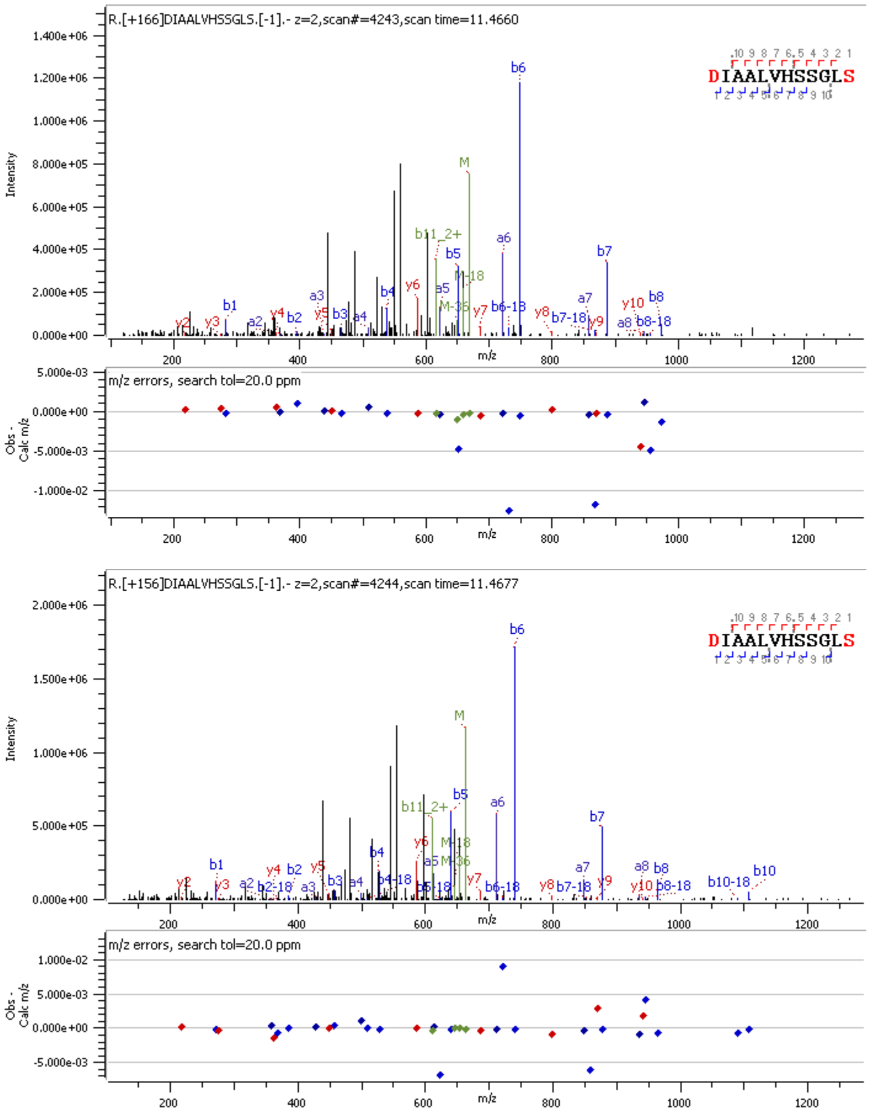


**Fig. S1**. MS2 identification of standard peptide arginylated by ATE1. DIAALVHSSGNleS-NH2 was arginylated by Arg10 (+166 Da) and Arg0 (+156 Da) with a mix ratio of 1:1. Leucine (L) was used as norleucine (Nle) replacement for search purposes. *m/z* values and intensities are displayed in the MS1 spectrum. Results are displayed in Byonic software.

**Related to Figure 1c.**

**a**, substrate peptide

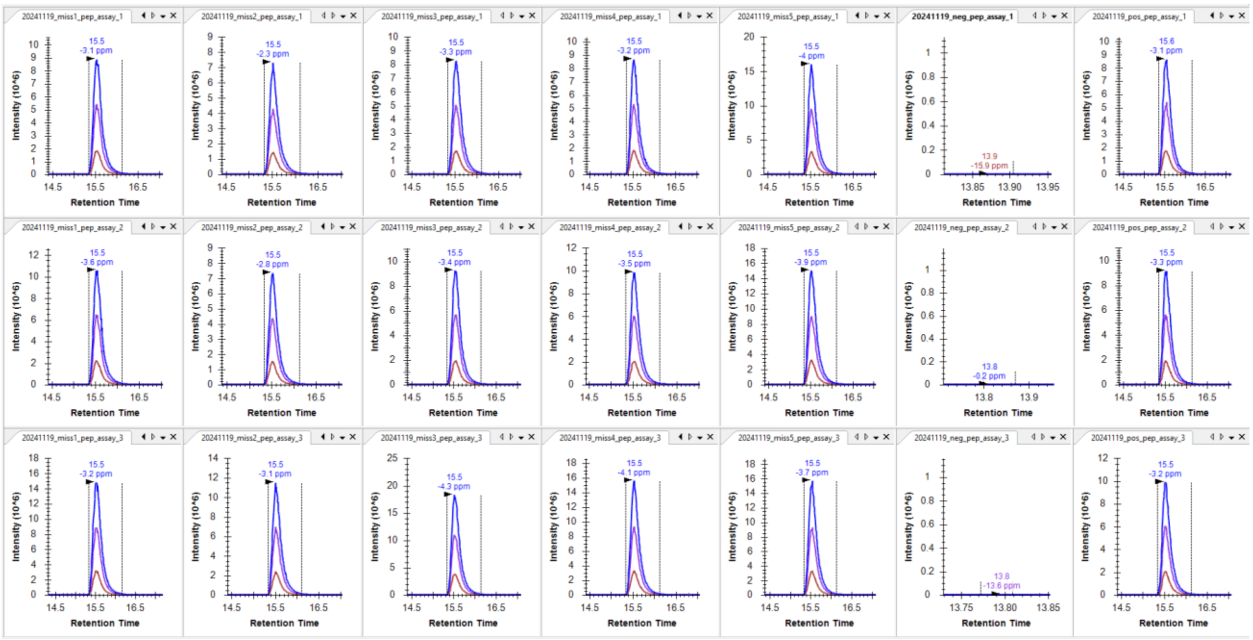


**b**, arginylated peptide

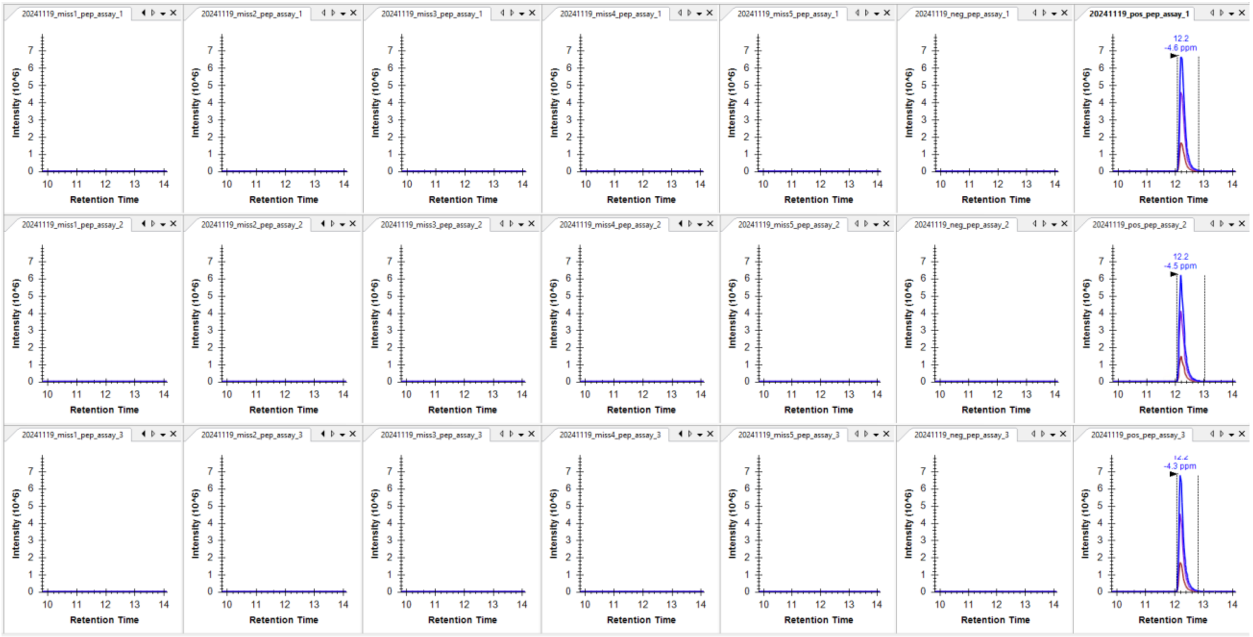


**Fig. S2**. Arginylated product from ATE1 assay consisted of different components. **a**, EIC of substrate peptide using Skyline software. **b**, EIC of arginylated peptide using Skyline software. ATE1 assays missing arginine, ATP, tRNA, RARS, ATE1, substrate, and nothing (from left to right). The assay was performed in triplicates (*n* = 3).

**Related to Figure 1c.**

**a**, substrate peptide

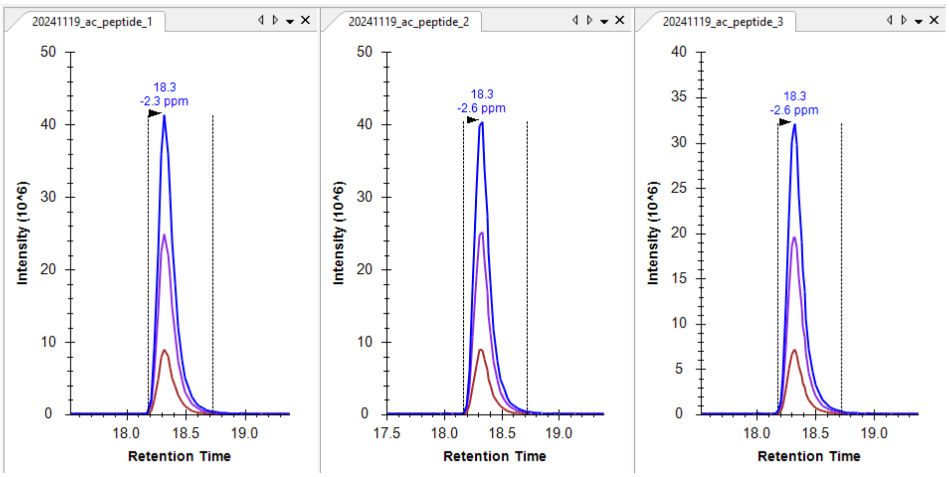


**b**, arginylated peptide

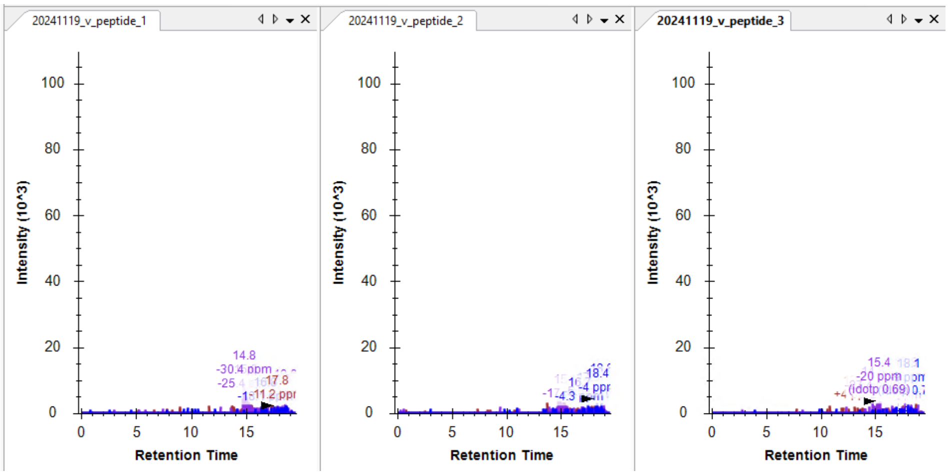


**Fig. S3**. ATE1 assay using N-terminal acetylated peptide. **a**, EIC of substrate peptide (acetyl-DIAALVHSSGNleS-NH2) using Skyline software. **b**, EIC of arginylated peptide using Skyline software. The assay was performed in triplicates (*n* = 3).

**Related to Figure 1c.**

**a**, substrate peptide

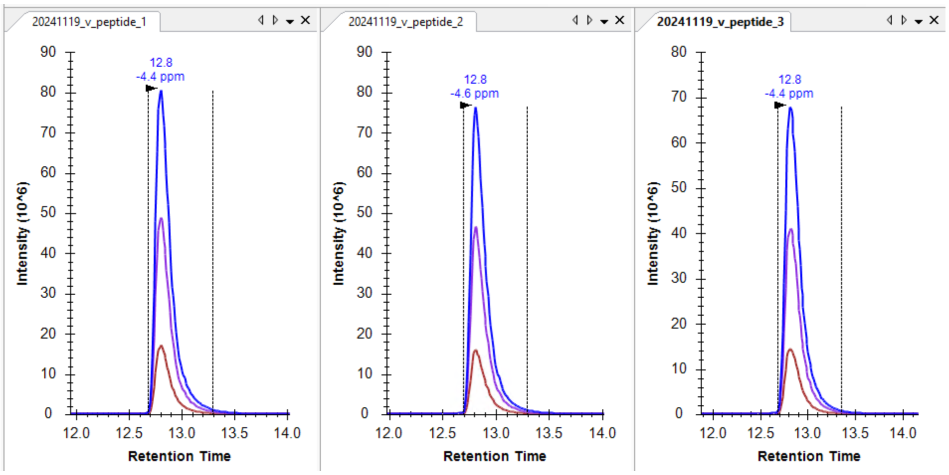


**b**, arginylated peptide

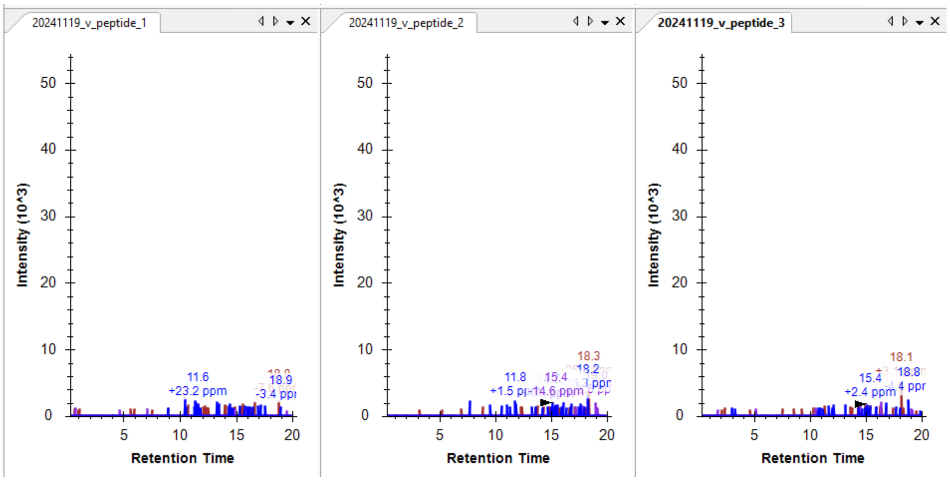


**Fig. S4**. ATE1 assay using a peptide with V as N-terminal residue. **a**, EIC of substrate peptide (VIAALVHSSGNleS-NH2) using Skyline software. **b**, EIC of arginylated peptide using Skyline software. The assay was performed in triplicates (*n* = 3).

**Related to Figure 1c.**

**a**, DIAALVHSSGNleS-NH2 → R-DIAALVHSSGNleS-NH2

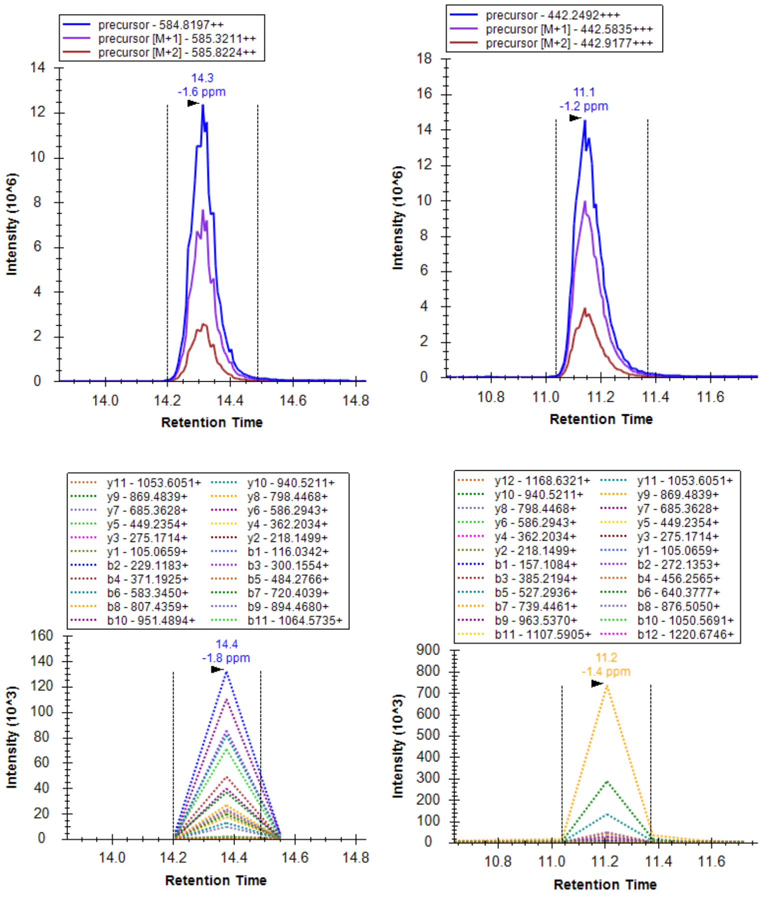


**b**,

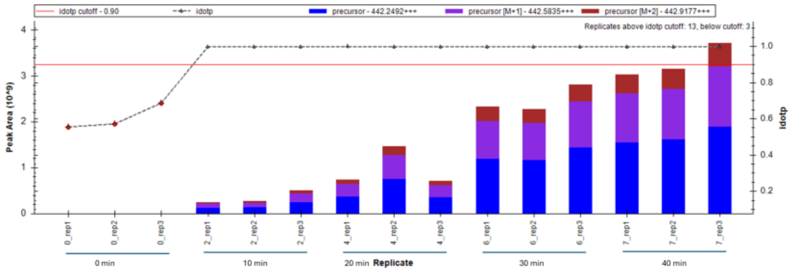


**c**,

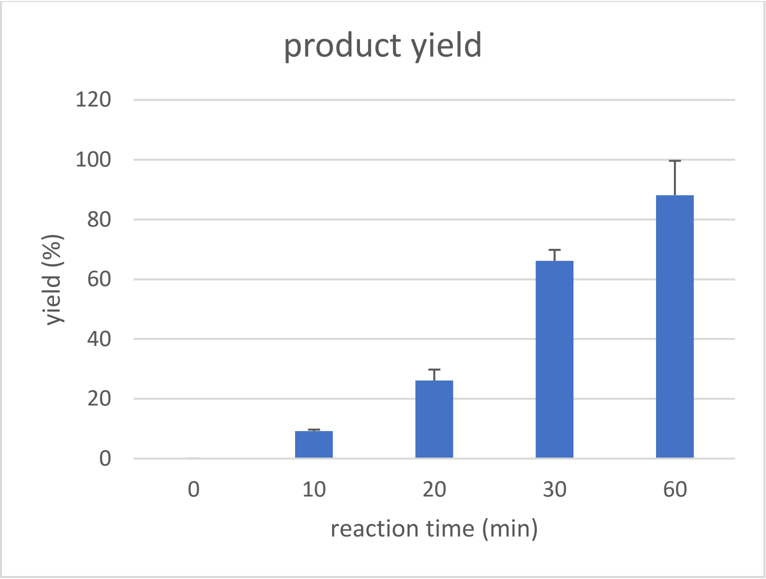


**d**,

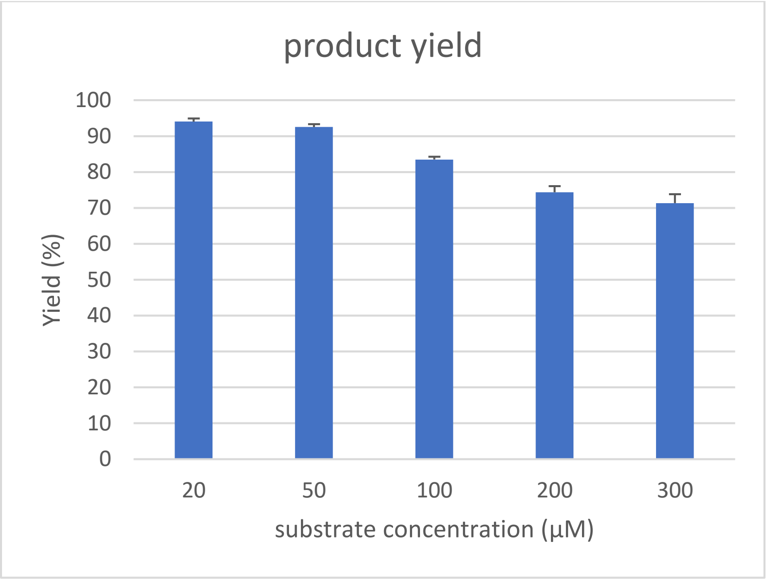


**Fig. S5**. ATE1 assay on standard peptide to monitor time-dependent and dose-dependent production of arginylated product. **a**, EIC and MS2 of standard peptide DIAALVHSSGNleS-NH2 and its arginylated product R-DIAALVHSSGNleS-NH2. **b**, quantification of product peptide using peak areas in Skyline software. Each time points are measured in triplicates (*n* = 3). **c**, time-dependent yield in the percentage of arginylated product. **d**, comparison of product yield using various concentrations of the peptide substrate.

**Related to Figure 1c.**

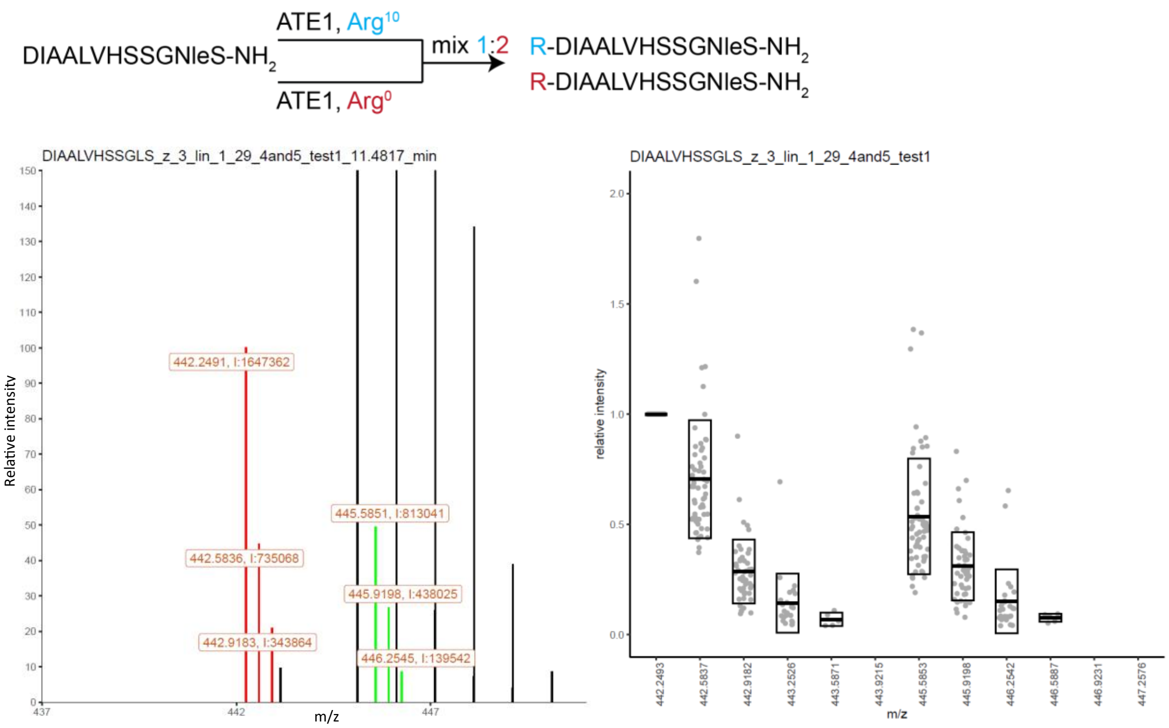


**Fig. S6**. MS1 ratio quantification of arginylated standard peptide. DIAALVHSSGNleS-NH2 was arginylated by Arg10 (+166 Da) and Arg0 (+156 Da) with a mix ratio of 1:2. Leucine (L) was used as norleucine (Nle) replacement for search purposes. *m/z* values and intensities (symbol: I) for highlighted peaks are displayed in the MS1 spectrum. A box plot is summarized from 61 MS1 scans.

**Related to Figure 1c.**

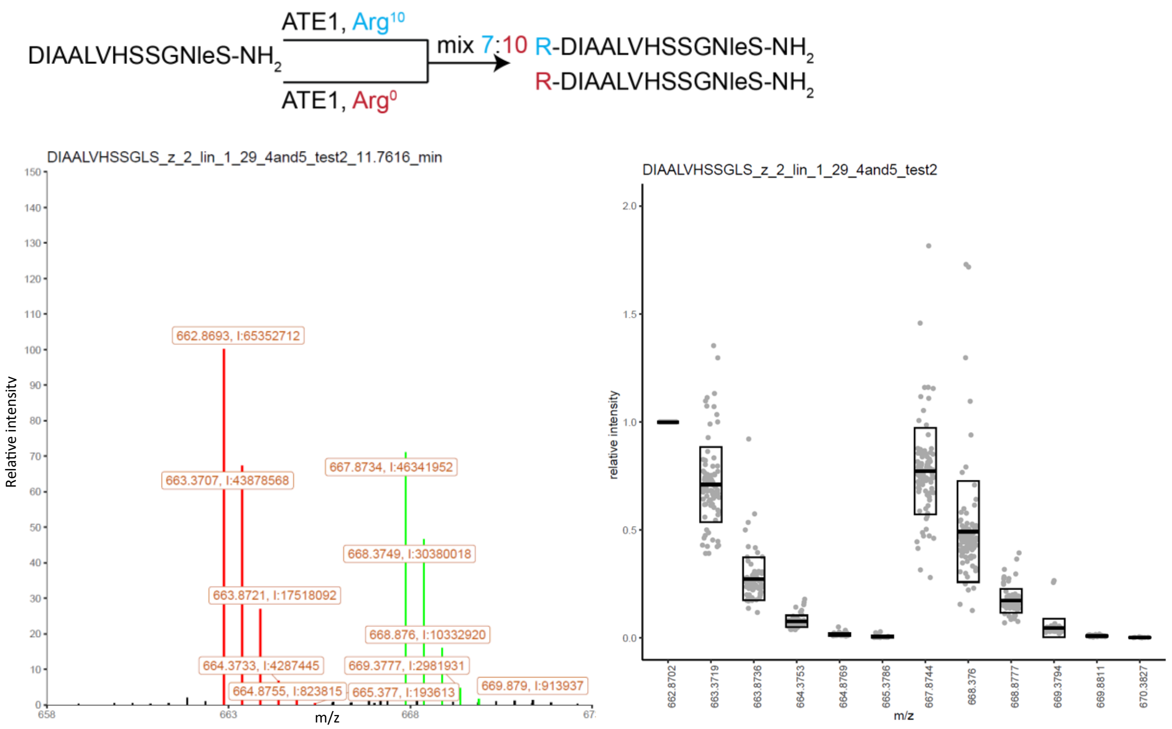


**Fig. S7**. MS1 ratio quantification of arginylated standard peptide. DIAALVHSSGNleS-NH2 was arginylated by Arg10 (+166 Da) and Arg0 (+156 Da) with a mix ratio of 7:10. Leucine (L) was used as norleucine (Nle) replacement for search purposes. *m/z* values and intensities (symbol: I) for highlighted peaks are displayed in the MS1 spectrum. A box plot is summarized from 102 MS1 scans.

**Related to Figure 1c.**

**Table S1**. Number of detections/pairs for each arginylation site from HEK293T tryptic peptide library.

| | unique sites (167) | | | | | | | |
|---|---|---|---|---|---|---|---|---|
| number of detections/pairs | 64 | ACTB D51 R | 5 | LDHA D82 R | 3 | RL18A D99 R | 2 | STMN1 E30 R |
| | 34 | SYK Q510 R_deami | 5 | MDHM E270 R | 3 | RL23A E 8 R | 2 | TCPQ D156 R |
| | 28 | H4 D25 R | 5 | PROF1 D27 R | 3 | RL36 E46 R | 1 | AT2A2 E656 R |
| | 24 | ACTB E316 R | 5 | RD23B D37 R | 3 | ROA1 D123 R | 1 | ATPA E208 R |
| | 20 | TBCD9 L1061 R | 5 | ROA2 D130 R | 3 | RRP5 D1816 R | 1 | CALM1 E15 R |
| | 16 | H31 D124 R | 5 | ROAA E233 R | 3 | RS14 E87 R | 1 | CALX E275 R |
| | 14 | H2B1K E94 R | 5 | TBA1B D327 R | 3 | RS3 D215 R | 1 | CDC37 E70 R |
| | 13 | SRSF1 D66 R | 4 | ANM5 E53 R | 3 | RS6 D120 R | 1 | CH10 D87 R |
| | 12 | H2B1K E36 R | 4 | G3BP1 E321 R | 3 | SRSF1 E143 R | 1 | CHD2 D1714 R |
| | 12 | RS12 D122 R | 4 | HNRPD D115 R | 3 | TCPG E382 R | 1 | COX5B E50 R |
| | 11 | ATP5I E60 R | 4 | HNRPU D627 R | 3 | TIF1B D128 R | 1 | DDX17 E255 R |
| | 11 | SRSF9 D64 R | 4 | HS90B E492 R | 3 | TKT D610 R | 1 | EF2 E440 R |
| | 10 | CH60 D353 R | 4 | ILF3 E298 R | 2 | AATM D154 R | 1 | EIF3G E181 R |
| | 10 | HSP7C D160 R | 4 | NIPA4 E152 R | 2 | AL7A1 D529 R | 1 | ETFB E165 R |
| | 10 | NUCL E411 R | 4 | NOLC1 E630 R | 2 | BIP E625 R | 1 | HERC2 V3548 R |
| | 10 | PTMA E22 R | 4 | PABP1 E313 R | 2 | CAH2 D19 R | 1 | HS90A E47 R |
| | 9 | RL26L D52 R | 4 | PLXA3 D1780 R | 2 | CDV3 E92 R | 1 | IFNA2 A168 R |
| | 9 | SF3A2 E92 R | 4 | RL38 D10 R | 2 | DNMT1 L899 R | 1 | METK2 D383 R |
| | 8 | CH10 D93 R | 4 | RL7 E10 R | 2 | EBP2 E263 R | 1 | NOLC1 D24 R |
| | 8 | ENPL E548 R | 4 | RS27 D 6 R | 2 | ELAV1 D105 R | 1 | NOP16 D114 R |
| | 8 | H31 E74 R | 4 | RS28 E52 R | 2 | FUBP2 D72 R | 1 | OFCC1 L127 R |
| | 8 | HDGF D62 R | 4 | SRSF9 D29 R | 2 | GUAA E425 R | 1 | PAIRB E274 R |
| | 8 | LRC59 D117 R | 4 | SSBP D96 R | 2 | H2B1J E36 R | 1 | PRDX3 E249 R |
| | 8 | RL10A D 8 R | 4 | TBA1A D327 R | 2 | HS90B E539 R | 1 | PUR6 D37 R |
| | 8 | RS19 D 8 R | 3 | 1433E E 5 R | 2 | IRS4 D675 R | 1 | PUR6 E23 R |
| | 7 | ENOA E10 R | 3 | CALR E25 R | 2 | LARP1 D115 R | 1 | RL19 D47 R |
| | 7 | GRP75 D207 R | 3 | CSPP1 E43 R | 2 | LKHA4 D574 R | 1 | RPN1 D580 R |
| | 7 | HSP7C E129 R | 3 | DLDH E496 R | 2 | MBB1A E1156 R | 1 | RS16 D110 R |
| | 7 | RS29 D49 R | 3 | ECHB D349 R | 2 | MCTS1 E75 R | 1 | RS25 E61 R |
| | 6 | DDX21 E185 R | 3 | EF2 E846 R | 2 | NOP56 E565 R | 1 | RS7 E42 R |
| | 6 | G6PI D117 R | 3 | GARS E109 R | 2 | PAIRB D229 R | 1 | RU17 D17 R |
| | 6 | H4 D69 R | 3 | H2B1B E36 R | 2 | PAIRB E93 R | 1 | SAFB1 D720 R |
| | 6 | HS90A D500 R | 3 | HNRH3 E68 R | 2 | PGK1 D98 R | 1 | SP100 D832 R |
| | 6 | HS90B E42 R | 3 | HNRPC D131 R | 2 | PRDX6 D9 R | 1 | SP16H E480 R |
| | 6 | KCRB D87 R | 3 | HNRPC E199 R | 2 | PRDX6 D42 R | 1 | SPB1 D683 R |
| | 6 | ODPB D220 R | 3 | HNRPK D397 R | 2 | PUR6 E20 R | 1 | SRSF1 D155 R |
| | 6 | RL12 E131 R | 3 | HS71A D160 R | 2 | RL26 D52 R | 1 | SRSF3 D4 R |
| | 6 | RL36 E88 R | 3 | MATR3 D121 R | 2 | ROA0 E177 R | 1 | THIC E201 R |
| | 6 | STMN1 D44 R | 3 | MMP3 G304 R | 2 | RS11 E49 R | 1 | TPIS E105 R |

| | | | | | | | | |
|---|---|---|---|---|---|---|---|---|
| | 5 | ALDOA E15 R | 3 | PGK1 D92 R | 2 | SRRT D866 R | 1 | TSYL2 D284 R |
| | 5 | DOPD E88 R | 3 | PININ D529 R | 2 | SSBP E17 R | 1 | VIME D176 R |
| | 5 | ENPL D672 R | 3 | RB6I2 E356 R | 2 | STIP1 E154 R | | |

R, arginylation (color code: black). RO3, Cys tri-oxidation and arginylation. RO2, Cys di-oxidation and arginylation. R_deami, N/Q arginylation after deamidation (color code: blue).

**Related to Figure 2c.**

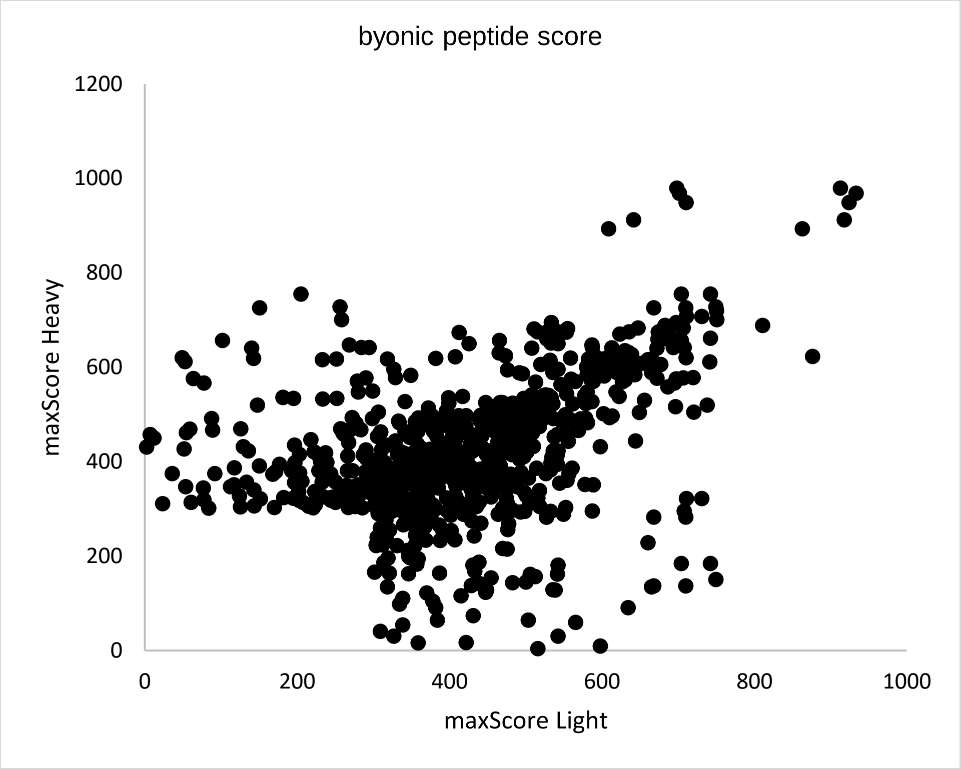


**Fig. S8**. Peptide score distribution for all detections/pairs from HEK293T tryptic peptide library. Peptides with both H and L scores < 300 are excluded.

**Related to Figure 2c.**

**a**,

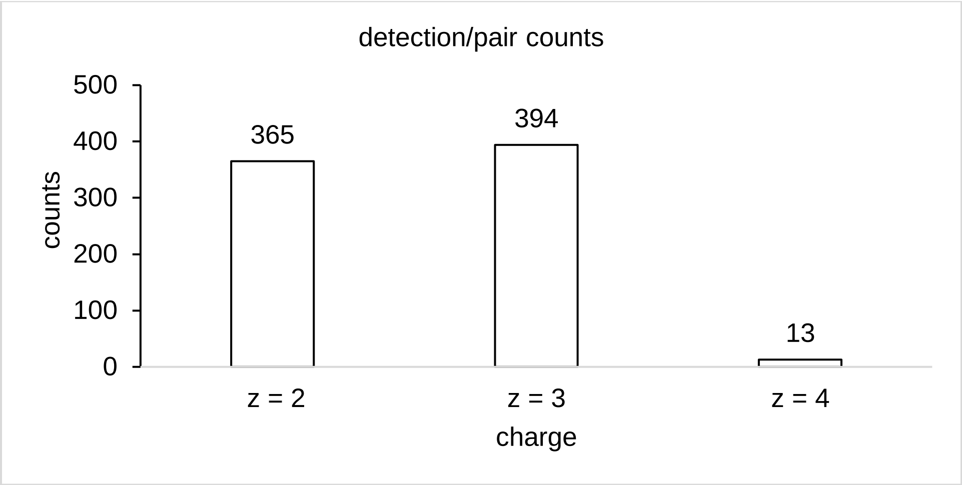
detection/pair counts
counts
500
400
300
200
100
0
365
394
13
z = 2
z = 3
z = 4
charge

**b**,

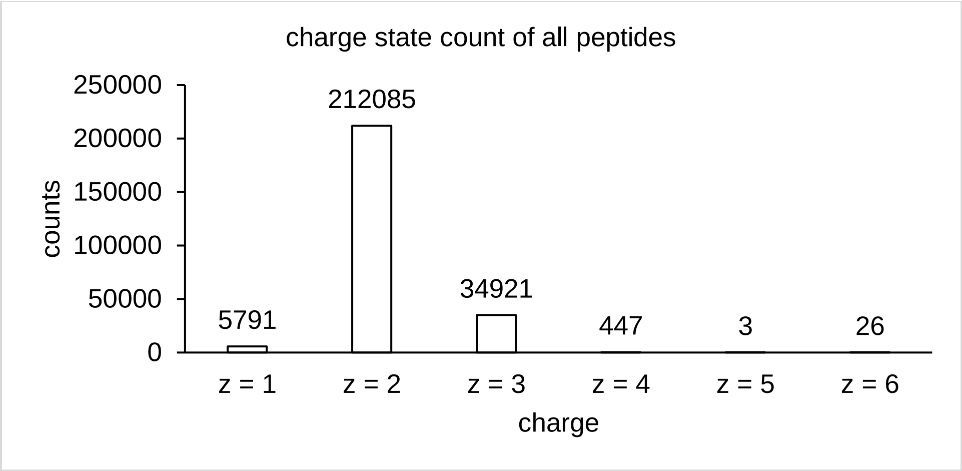
charge state count of all peptides
counts
250000
200000
150000
100000
50000
0
5791
212085
34921
447
3
26
z = 1
z = 2
z = 3
z = 4
z = 5
z = 6
charge

**c**,

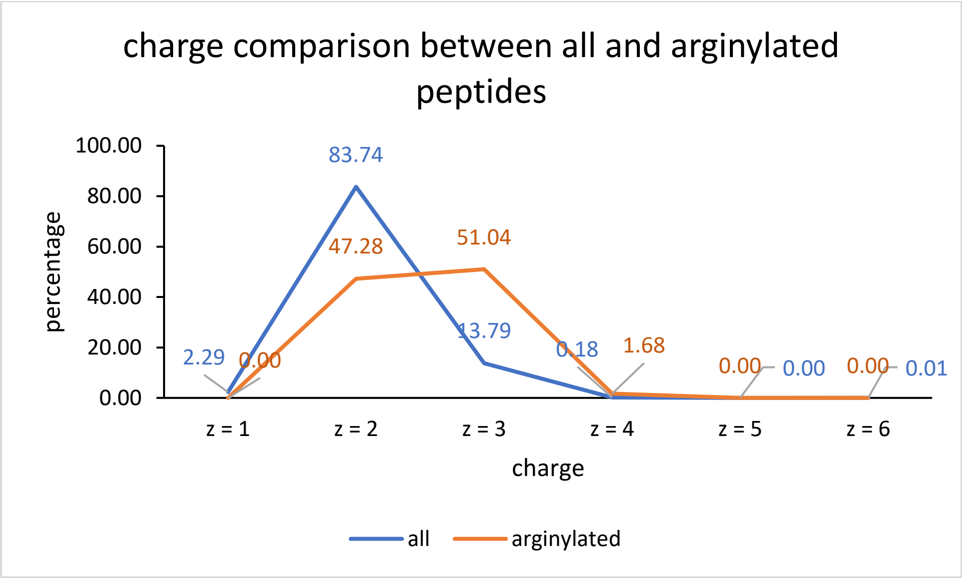
charge comparison between all and arginylated peptides
percentage
100.00
80.00
60.00
40.00
20.00
0.00
2.29
0.00
83.74
47.28
51.04
13.79
0.18
1.68
0.00
0.00
0.00
0.01
z = 1
z = 2
z = 3
z = 4
z = 5
z = 6
charge
all
arginylated

**Fig. S9**. Charge distribution comparison between all peptides and arginylated peptides from HEK293T tryptic peptide library. **a**, charge distribution pattern of arginylated peptides. **b**, the charge distribution pattern of all peptides. **c**, percentage comparison of all and arginylated peptides. Charges 2-3 in arginylated peptides shifted towards higher charges due to arginylation (charge 2: 47%, charge 3: 51%) compared with all peptides (charge 2: 84%, charge 3: 14%).

**Related to Figure 2c.**

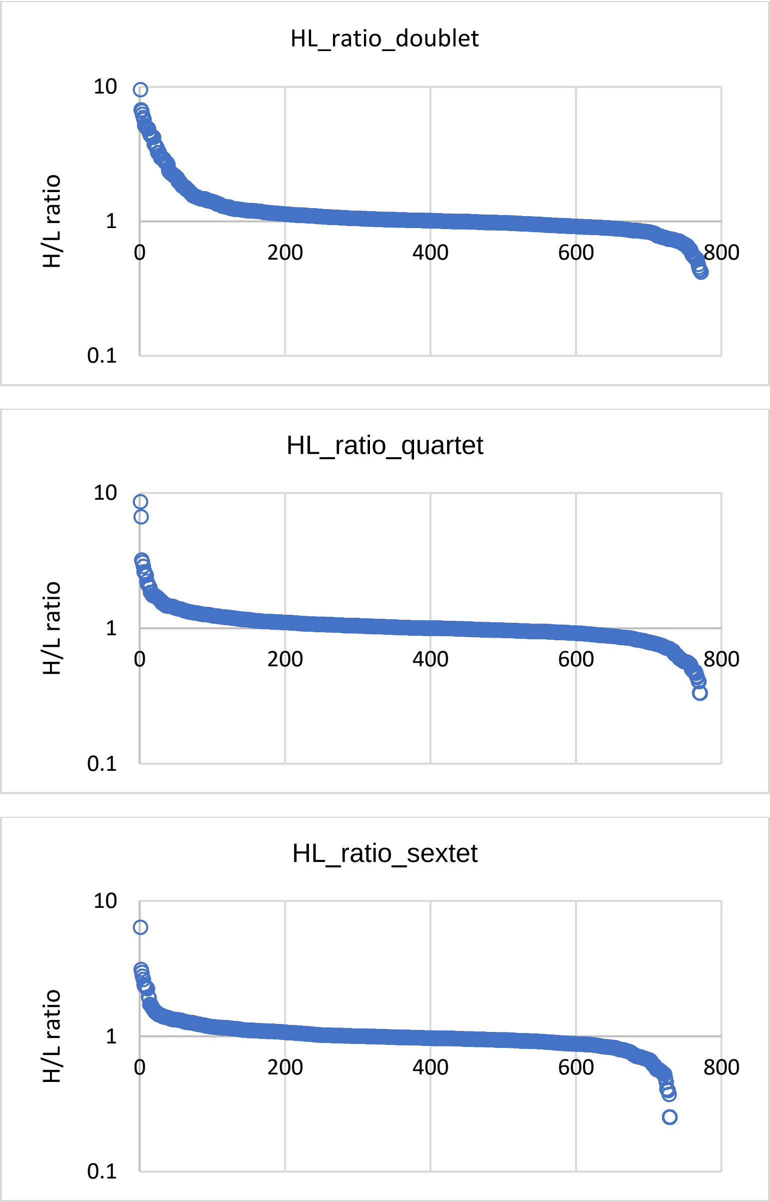


**Fig. S10**. H/L ratio distributions of all MS1 pairs in doublet, quartet, and sextet from HEK293T tryptic peptide library.

**Related to Figure 2d.**

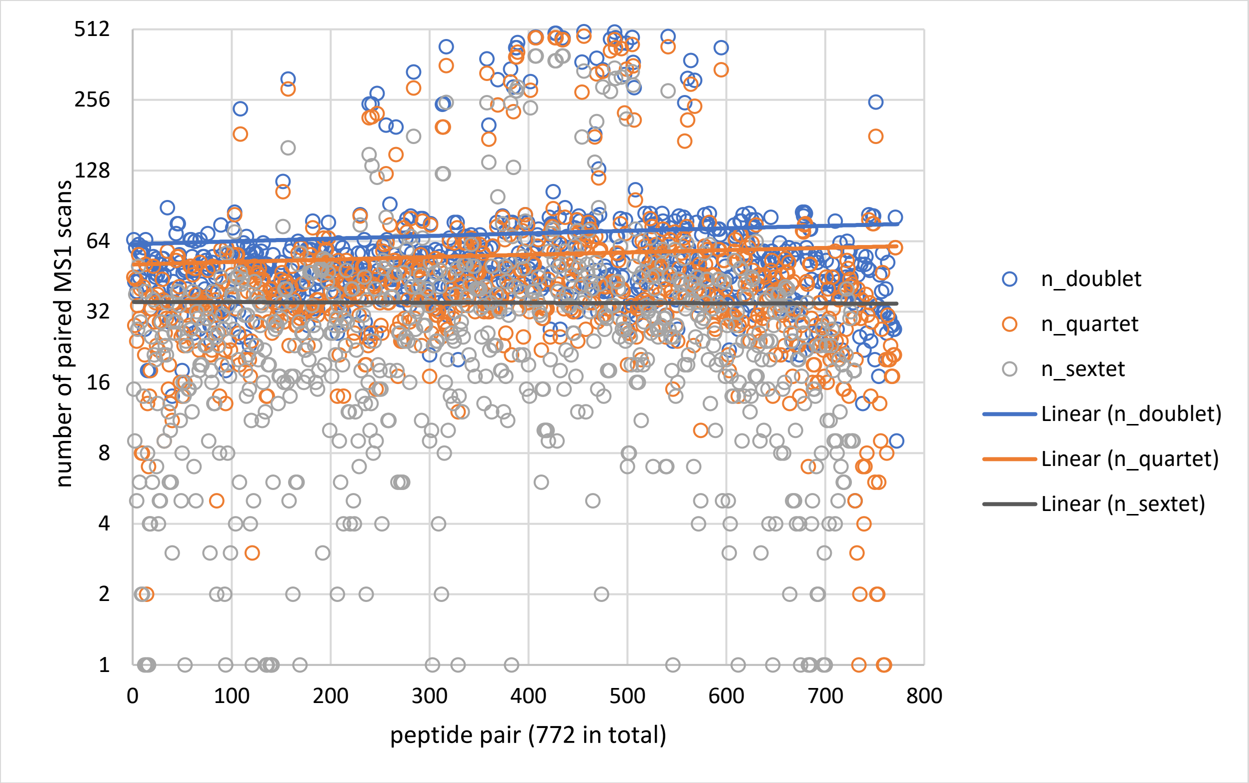


**Fig. S11**. Numbers of MS1 scans of all 772 peptide pairs in doublet, quartet, and sextet from HEK293T tryptic peptide library. Trendlines for all doublets, quartets and sextets are generated respectively.

**Related to Figure 2d.**

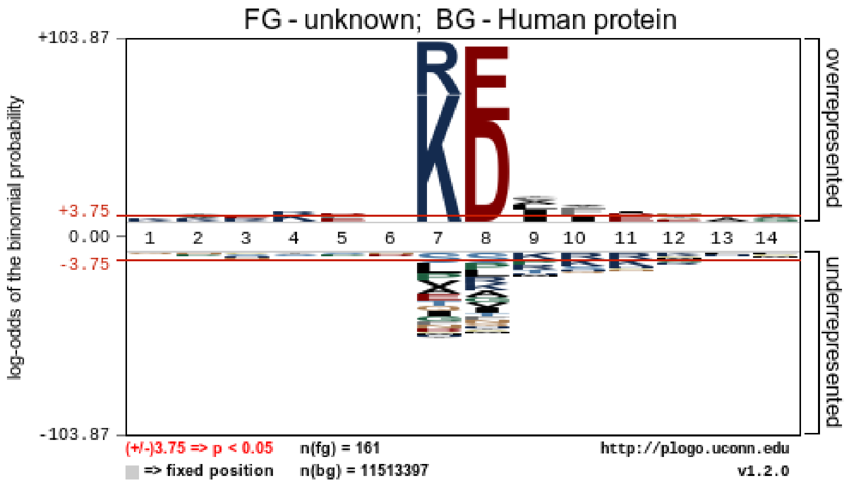


**Fig. S12.** Arginylation motif calculated from unmodified forms of all unique arginylated peptides from HEK293T tryptic peptide library. The significant motif is generated by pLogo using the human proteome as the background.

**Related to Figure 2f.**

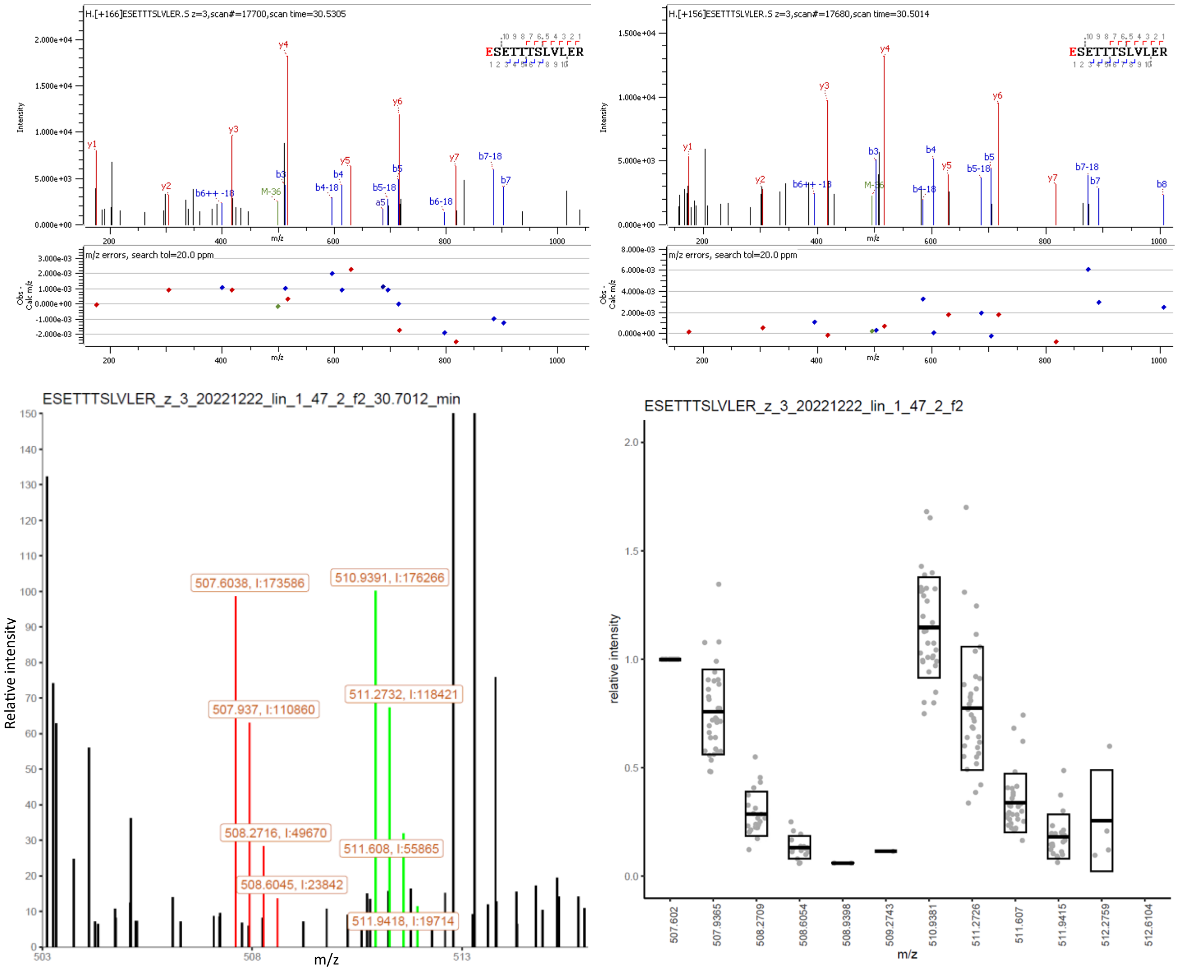


**Fig. S13**. MS2 identification and MS1 ratio quantification of arginylated SSBP E17 from HEK293T tryptic peptide library. *m/z* values and intensities for highlighted peaks are displayed in the MS1 spectrum. A box plot is summarized from 33 MS1 scans. Arg10 modification, +166 Da. Arg0 modification, +156 Da.

**Related to Figure 2g.**

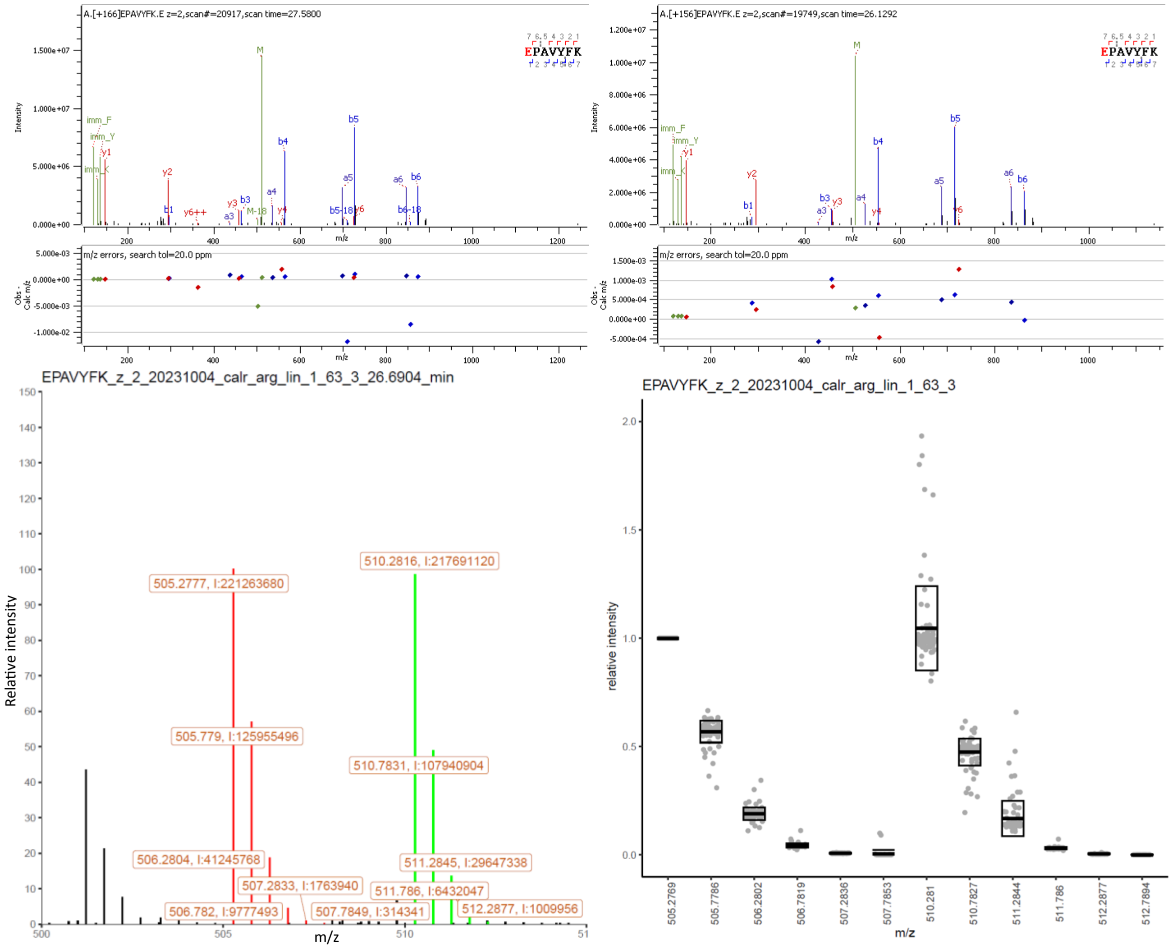


**Fig. S14**. MS2 identification and MS1 ratio quantification of arginylated CALR E18 after overexpression and purification. *m/z* values and intensities for highlighted peaks are displayed in the MS1 spectrum. A box plot is summarized from 33 MS1 scans. Arg10 modification, +166 Da. Arg0 modification, +156 Da.

**Related to Figure 3a.**

**a**, EPAVYFK

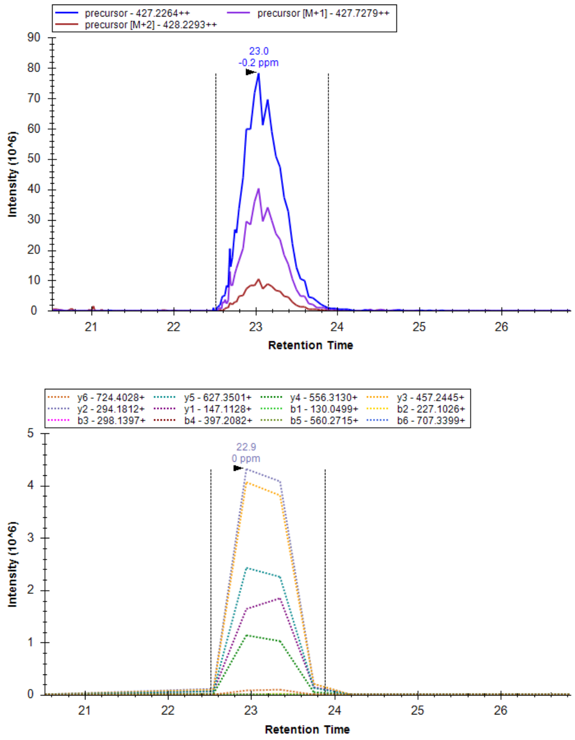


**b**,

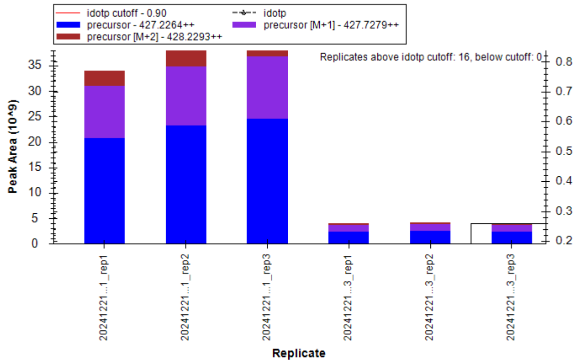


**c**, REPAVYFK

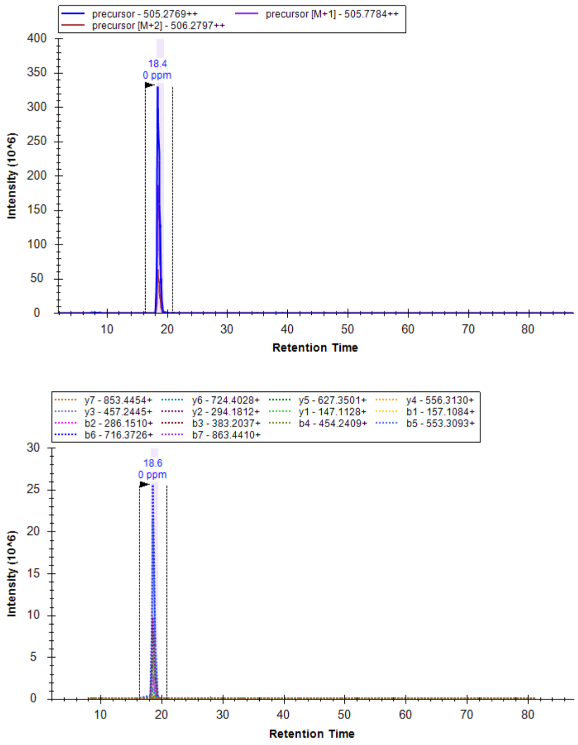


**d**,

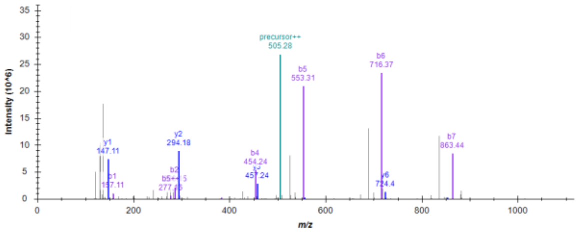


**Fig. S15**. ATE1 assay on pure CALR protein to calculate arginylation efficiency. **a**, EIC of MS1 precursor and MS2 fragments of EPAVYFK in CALR. **b**, quantification of EPAVYFK peptide using peak areas in Skyline software (89.1 ± 0.9%). Experiments are measured in triplicates (*n* = 3). Left (*n* = 3): negative control without ATE1, right (*n* = 3): experiments with ATE1. Arginylation efficiency is calculated as 61.7 ± 3.1% by spectra counts in arginylation samples. Arginylation efficiency is calculated as 80.2 ± 6.6% by spectra counts of EPAVYFK in negative and arginylation samples. **c**, EIC of MS1 precursor and MS2 fragments of REPAVYFK in CALR. **d**, MS2 spectrum annotation of REPAVYFK peptide in Skyline software.

**Related to Figure 3a.**

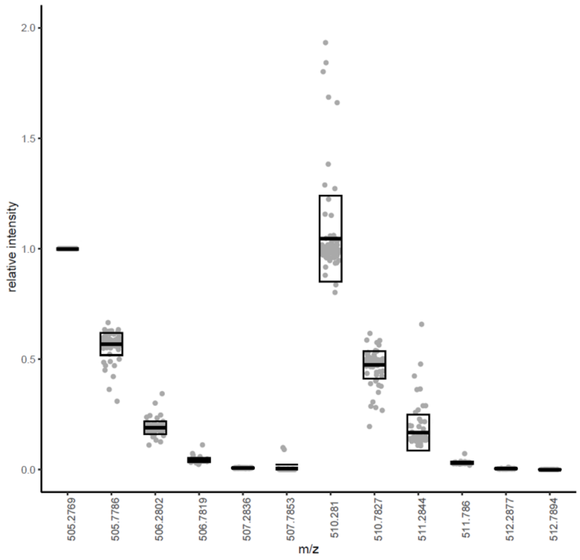
2.0
1.5
1.0
0.5
0.0
relative intensity
505.2769
505.7786
506.2802
506.7819
507.2836
507.7853
510.281
510.7827
511.2844
511.786
512.2877
512.7894
m/z

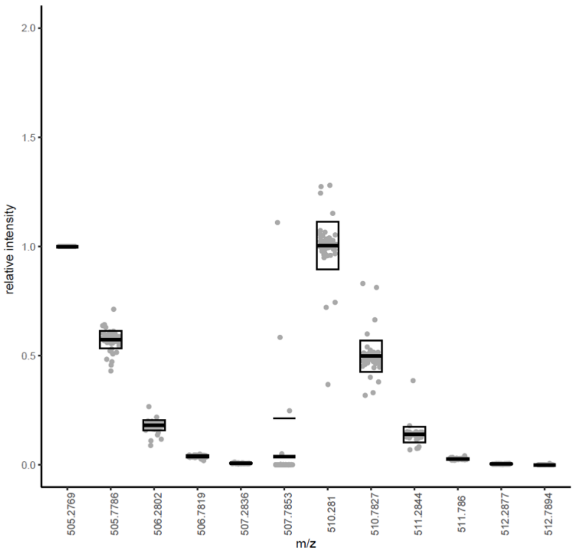
2.0
1.5
1.0
0.5
0.0
relative intensity
505.2769
505.7786
506.2802
506.7819
507.2836
507.7853
510.281
510.7827
511.2844
511.786
512.2877
512.7894
m/z

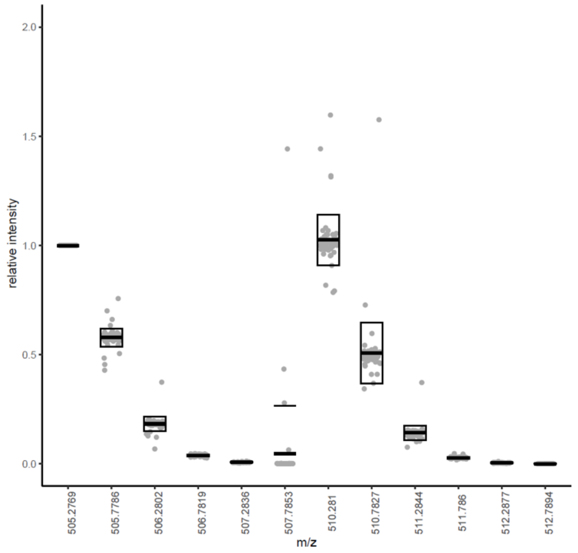
2.0
1.5
1.0
0.5
0.0
relative intensity
505.2769
505.7786
506.2802
506.7819
507.2836
507.7853
510.281
510.7827
511.2844
511.786
512.2877
512.7894
m/z

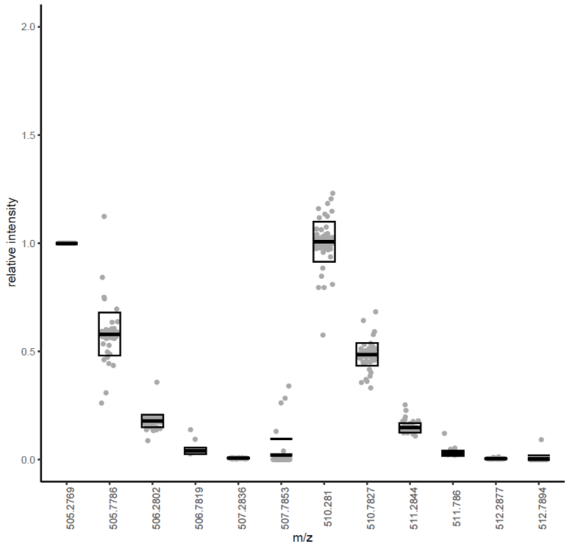
2.0
1.5
1.0
0.5
0.0
relative intensity
505.2769
505.7786
506.2802
506.7819
507.2836
507.7853
510.281
510.7827
511.2844
511.786
512.2877
512.7894
m/z

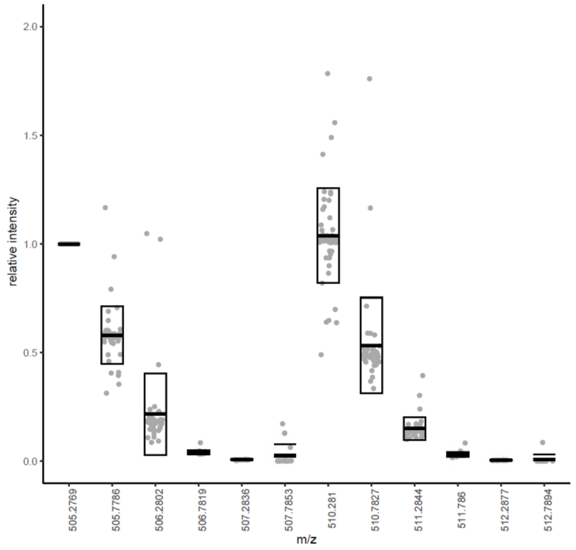
2.0
1.5
1.0
0.5
0.0
relative intensity
505.2769
505.7786
506.2802
506.7819
507.2836
507.7853
510.281
510.7827
511.2844
511.786
512.2877
512.7894
m/z

**Fig. S16**. MS1 ratio quantification of arginylated CALR E18 after overexpression and purification. H/L ratios in box plots are 0.95, 1.01, 0.99, 1.00, 1.06, and 0.99 summarized from 97, 71, 67, 72, and 53 MS1 scans in five replicates (*n* = 5), respectively.

**Related to Figure 3a.**

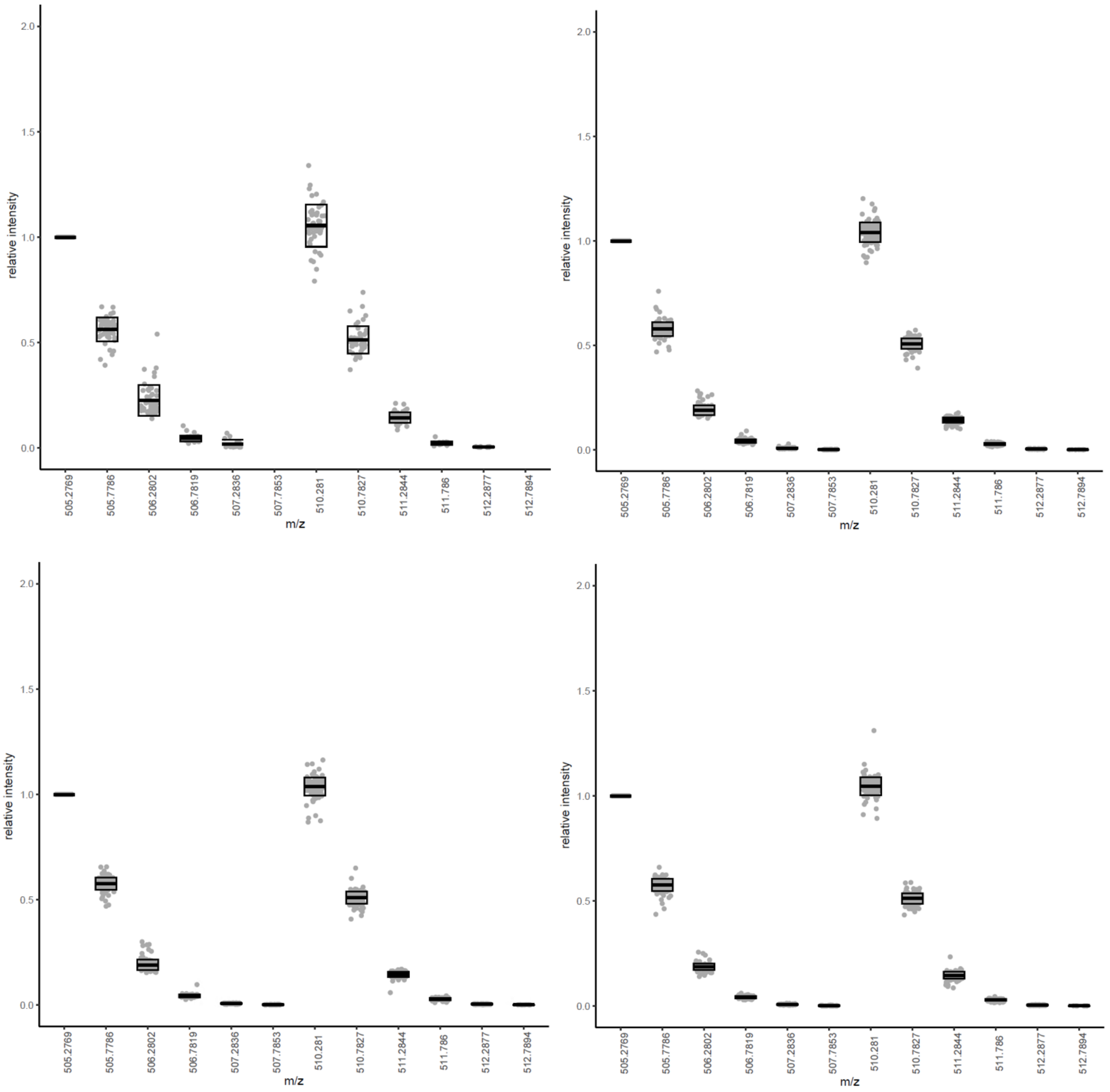
relative intensity
m/z
2.0
1.5
1.0
0.5
0.0
505.2769
505.7786
506.2802
506.7819
507.2836
507.7853
510.281
510.7827
511.2844
511.786
512.2877
512.7894

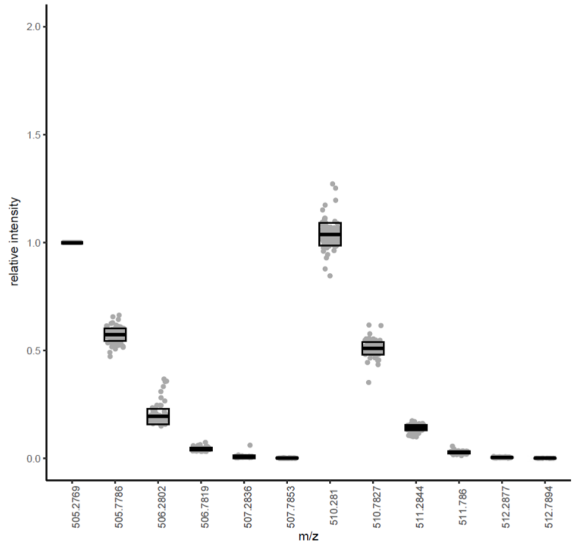


**Fig. S17**. MS1 ratio quantification of arginylated CALR E18 from commercial source. Protein was purchased from Abcam (cat no.: ab276554). H/L ratios in box plots are 0.96, 0.96, 0.97, 0.96, and 0.97 summarized from 50, 122, 127, 131, and 130 MS1 scans in five replicates ($n$ = 5), respectively.

**Related to Figure 3a.**

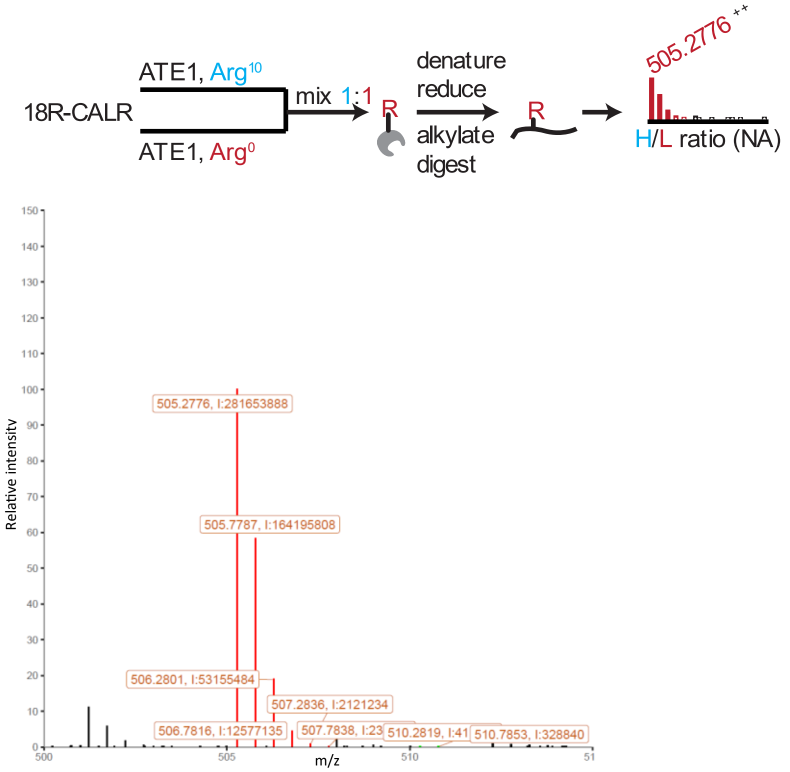


**Fig. S18**. Representative MS1 scan of overexpressed and purified 18R-CALR arginylation by ATE1. *m/z* values and intensities for highlighted peaks are displayed in the MS1 spectrum. H/L ratio is not available due to the lack of the E18 site for isotopic (Arg10) arginylation. The experiment was performed 5 times (*n* = 5).

**Related to Figure 3a.**

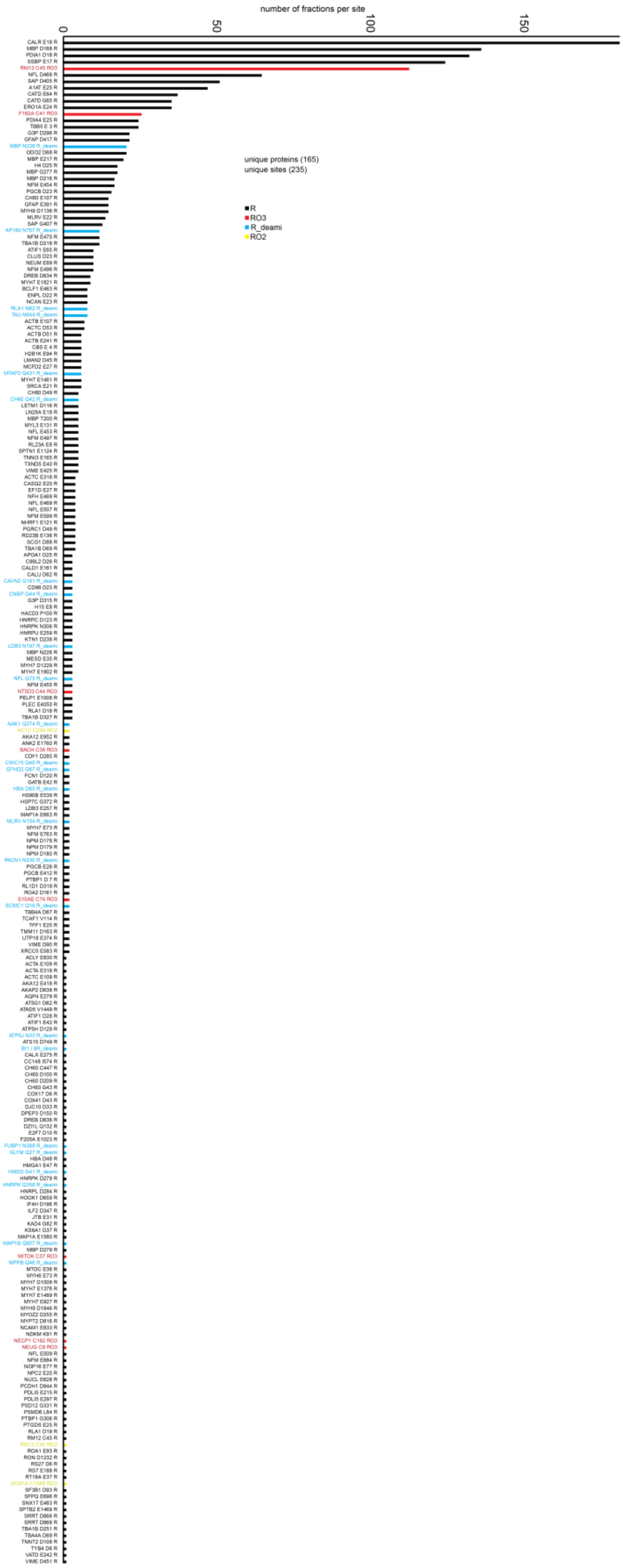
number of fractions per site
0
50
100
150
unique proteins (165)
unique sites (235)
R
RO3
R_deami
RO2

**Fig. S19.** Number of detections/pairs for each arginylation site from human proteomes. R, arginylation. RO3, Cys tri-oxidation and arginylation. RO2, Cys di-oxidation and arginylation. R_deami, N/Q arginylation after deamidation. *, validated by standard peptides.

**Related to Figure 3b.**

**Table S2. Number of detections in isotopic pairs and MS1 scans for each arginylation site from human proteomes.**

| Site | n1 | n2 | Site | n1 | n2 | Site | n1 | n2 |
|---|---|---|---|---|---|---|---|---|
| *CALR E18 R | 185 | 8434 | LMAN2 D45 R | 6 | 147 | CNBP Q44 R_deami | 3 | 84 |
| MBP D168 R | 139 | 5153 | MTAP2 Q431 R_deami | 6 | 122 | MYH7 E1902 R | 3 | 71 |
| *PDIA1 D18 R | 135 | 6071 | SRCA E21 R | 6 | 108 | CALU D62 R | 3 | 70 |
| *SSBP E17 R | 127 | 4744 | CBS E 4 R | 6 | 101 | KTN1 D238 R | 3 | 61 |
| *RM12 C45 RO3 | 115 | 3085 | MYH7 E1461 R | 6 | 96 | HACD3 P100 R | 3 | 58 |
| NFL D468 R | 66 | 3260 | ACTB E241 R | 6 | 79 | PLEC E4050 R | 3 | 56 |
| SAP D405 R | 52 | 1485 | H2B1K E94 R | 6 | 35 | MYH7 D1229 R | 3 | 54 |
| A1AT E25 R | 48 | 1648 | NFM E497 R | 5 | 321 | TBA1B D327 R | 3 | 52 |
| CATD E64 R | 38 | 986 | MBP T200 R | 5 | 264 | NFL Q75 R_deami | 3 | 46 |
| CATD G65 R | 36 | 1008 | CH60 Q42 R_deami | 5 | 207 | CALD1 E161 R | 3 | 42 |
| *ERO1A E24 R | 36 | 984 | LN28A E18 R | 5 | 200 | NT5D3 C44 RO3 | 3 | 35 |
| F162A C41 RO3 | 26 | 516 | LETM1 D116 R | 5 | 188 | PTBP1 D 7 R | 2 | 124 |
| TBB5 E 3 R | 25 | 1342 | TNNI3 E165 R | 5 | 169 | FCN1 D120 R | 2 | 111 |
| PDIA4 E25 R | 25 | 756 | VIME E425 R | 5 | 166 | HS90B E539 R | 2 | 102 |
| GFAP D417 R | 22 | 722 | NFL E453 R | 5 | 145 | LDB3 E257 R | 2 | 79 |
| G3P D296 R | 22 | 482 | MYL3 E131 R | 5 | 119 | NPM D178 R | 2 | 77 |
| ODO2 D68 R | 21 | 735 | CH60 D49 R | 5 | 111 | ROA2 D161 R | 2 | 76 |
| MBP N226 R_deami | 21 | 364 | TXND5 E40 R | 5 | 94 | VIME D90 R | 2 | 76 |
| MBP E217 R | 20 | 582 | SPTN1 E1124 R | 5 | 61 | S10AE C74 RO3 | 2 | 75 |
| MBP G277 R | 18 | 488 | RL23A E 8 R | 5 | 59 | XRCC5 E583 R | 2 | 66 |
| H4 D25 R | 18 | 466 | PGRC1 D49 R | 4 | 201 | NPM D180 R | 2 | 63 |
| MBP D216 R | 17 | 550 | NFL E469 R | 4 | 193 | TCAF1 V114 R | 2 | 59 |
| NFM E454 R | 17 | 466 | CASQ2 E20 R | 4 | 151 | MAP1A E663 R | 2 | 58 |
| PGCB D23 R | 16 | 772 | TBA1B D69 R | 4 | 146 | HSP7C G372 R | 2 | 55 |
| GFAP E391 R | 15 | 600 | NFM E599 R | 4 | 129 | *GATB E42 R | 2 | 53 |
| MYH9 D1136 R | 15 | 495 | ACTC E318 R | 4 | 119 | NPM D179 R | 2 | 53 |
| *CH60 E107 R | 15 | 452 | NFH E469 R | 4 | 88 | EFHD2 Q67 R_deami | 2 | 50 |
| MLRV E22 R | 14 | 266 | NFL E507 R | 4 | 88 | TBB4A D67 R | 2 | 50 |
| SAP G407 R | 13 | 172 | NHRF1 E121 R | 4 | 77 | HBA N79 R_deami | 2 | 48 |
| NFM E470 R | 12 | 354 | RD23B E136 R | 4 | 65 | SCMC1 Q16 R_deami | 2 | 45 |
| TBA1B D218 R | 12 | 132 | SCG1 D88 R | 4 | 51 | TFF1 E25 R | 2 | 42 |
| AP180 N767 R_deami | 12 | 77 | EF1D E27 R | 4 | 21 | CWC15 Q45 R_deami | 2 | 40 |
| NFM E496 R | 10 | 770 | HNRPK N306 R | 3 | 146 | AKA12 E952 R | 2 | 37 |
| CLUS D23 R | 10 | 179 | MBP N226 R | 3 | 142 | MYH7 E73 R | 2 | 35 |
| NEUM E89 R | 10 | 152 | APOA1 D25 R | 3 | 140 | RL1D1 D319 R | 2 | 32 |
| ATIF1 E65 R | 10 | 148 | H15 E 8 R | 3 | 119 | BACH C36 RO3 | 2 | 31 |
| DREB D634 R | 9 | 460 | CD99 D23 R | 3 | 115 | PGCB E412 R | 2 | 29 |
| MYH7 E1821 R | 9 | 232 | C99L2 D26 R | 3 | 114 | UTP18 E374 R | 2 | 26 |
| ENPL D22 R | 8 | 555 | LDB3 N197 R_deami | 3 | 112 | AAK1 Q374 R_deami | 2 | 23 |
| NCAN E23 R | 8 | 177 | HNRPU E259 R | 3 | 110 | MLRV N154 R_deami | 2 | 23 |
| BCLF1 E463 R | 8 | 160 | HNRPC D123 R | 3 | 109 | TMM11 D163 R | 2 | 20 |
| RLA1 N62 R_deami | 8 | 159 | RLA1 D18 R | 3 | 109 | ACTC C259 RO2 | 2 | 17 |
| TAU N644 R_deami | 8 | 136 | PELP1 E1008 R | 3 | 104 | CDY1 D285 R | 2 | 17 |
| ACTC D53 R | 7 | 270 | CAVN2 Q161 R_deami | 3 | 95 | NFM E763 R | 2 | 17 |
| ACTB E107 R | 7 | 179 | G3P D315 R | 3 | 92 | ANK2 E1760 R | 2 | 15 |
| ACTB D51 R | 6 | 257 | MESD E35 R | 3 | 92 | PACN1 N330 R_deami | 2 | 8 |
| MCFD2 E27 R | 6 | 231 | NFM E455 R | 3 | 87 | PGCB E26 R | 2 | 7 |

Notes. R, arginylation (color code: black). RO3, Cys tri-oxidation and arginylation (color code: red). RO2, Cys di-oxidation and arginylation (color code: yellow). R_deami, N/Q arginylation after deamidation (color code: blue). *, validated by standard peptides. n1, number of total fractions. n2, number of total paired MS1 scans.

**Related to Table 1 and Figure 3c.**

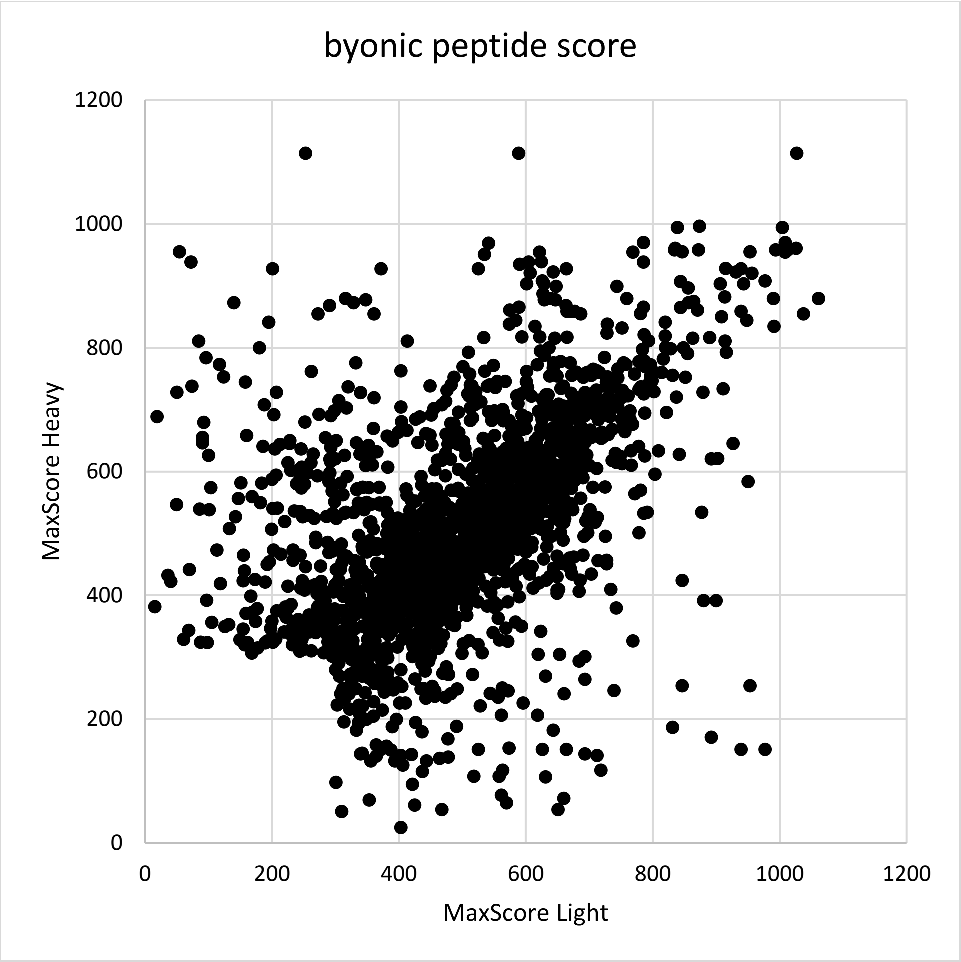


**Fig. S20**. Peptide score distribution for all detections/pairs from human proteomes. Peptides with both H and L scores < 300 are excluded.

**Related to Figure 3b.**

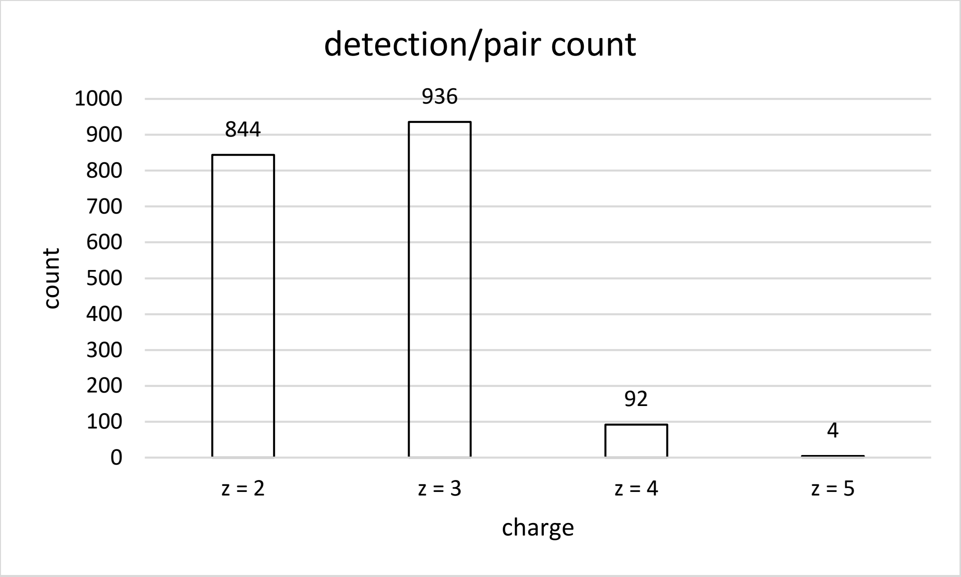


**Fig. S21**. Peptide charge distribution for all detections/pairs from human proteomes.

**Related to Figure 3d.**

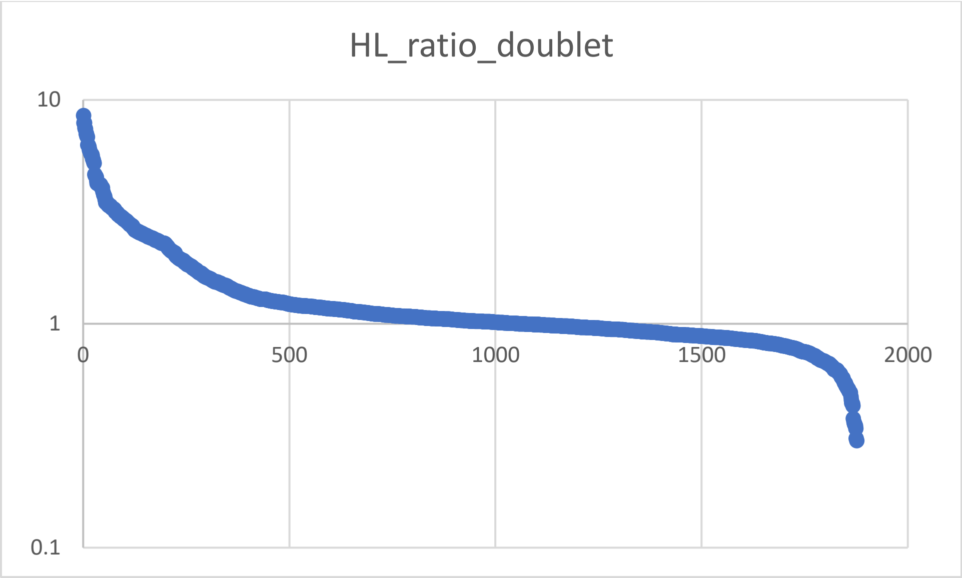
HL_ratio_doublet
10
1
0.1
0
500
1000
1500
2000

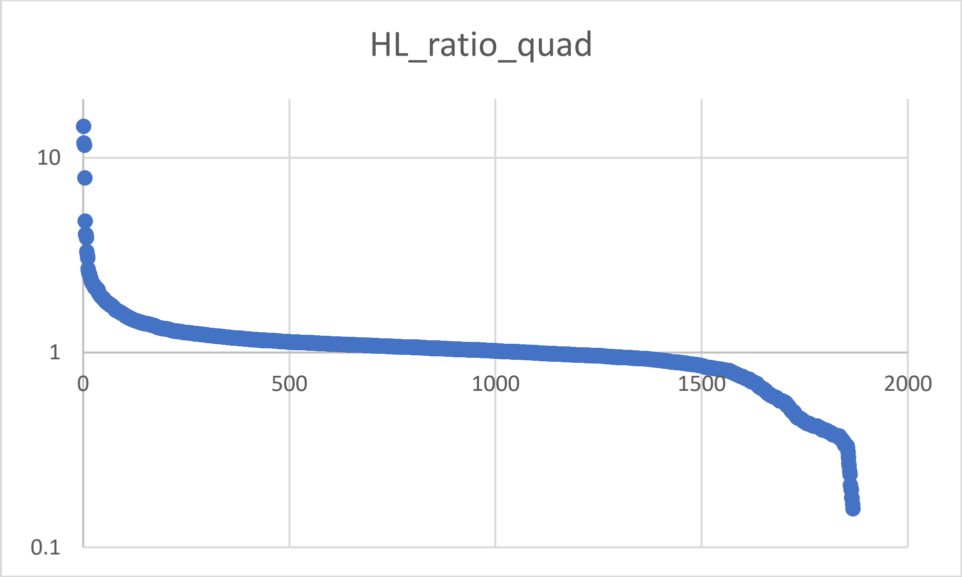
HL_ratio_quad
10
1
0.1
0
500
1000
1500
2000

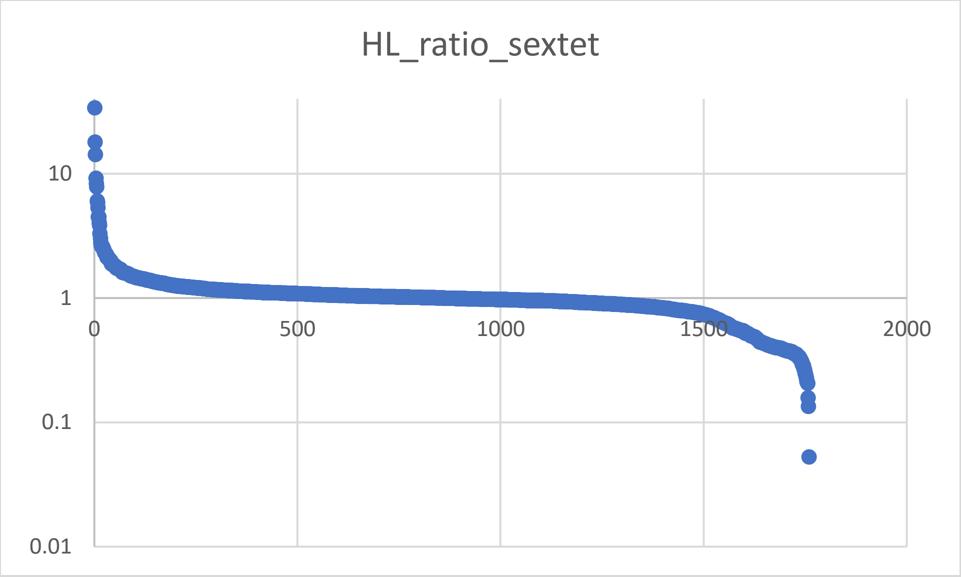


**Fig. S22**. H/L ratio distribution of all MS1 pairs in doublet, quartet, and sextet from human proteomes.

**Related to Figure 3d.**

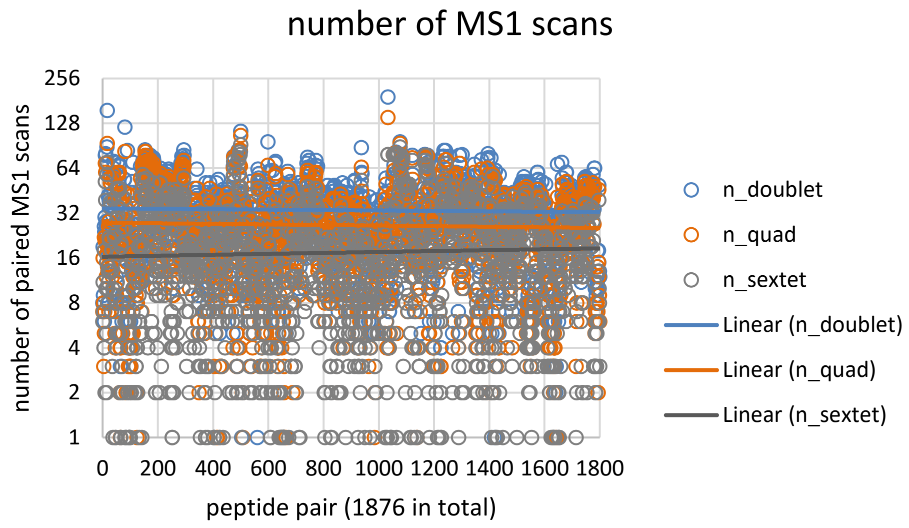


**Fig. S23**. Numbers of MS1 scans of all 1876 peptide pairs in doublet, quartet, and sextet from human proteomes. Trendlines for all doublets, quartets and sextets are generated respectively.

**Related to Figure 3d.**

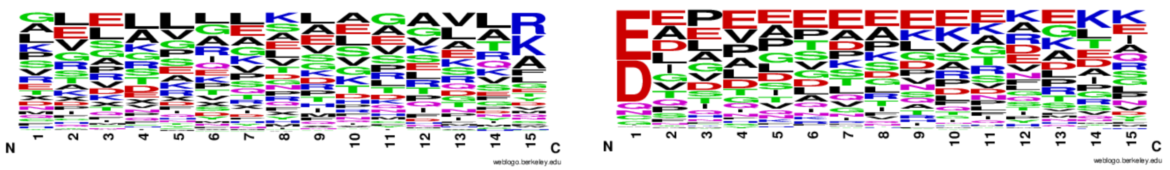

**Fig. S24**. Sequence logo calculated from unmodified forms of all unique arginylated peptides from human proteomes. Frequency plots are generated by WebLogo.

**Related to Figure 3e.**

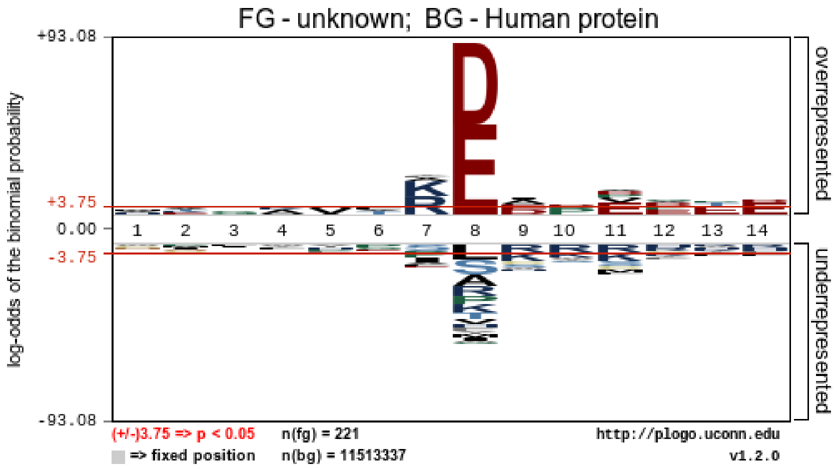


**Fig. S25**. Arginylation motif calculated from unmodified forms of all unique arginylated peptides from human cells and patient tissues. The significant motif is generated by pLogo using the human proteome as the background.

**Related to Figure 3e.**

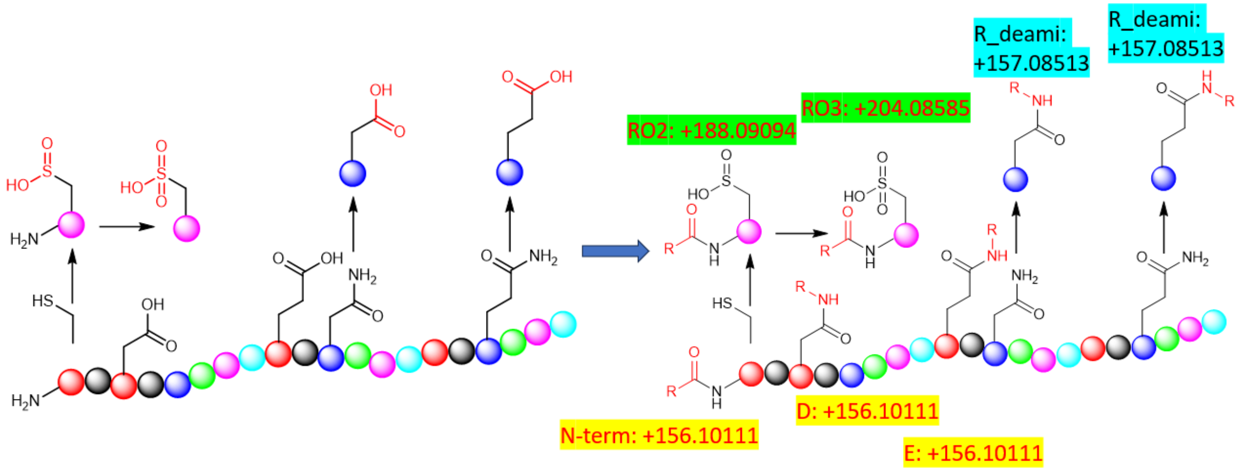


**Fig. S26**. ATE1 mediated arginylation types. Possible sites for arginylation and their products. Several types of arginylation are proposed including N-terminal arginylation (R: +156 Da), midchain arginylation (R: +156 Da) on D and E, Cys tri-oxidation and arginylation (RO3: +204 Da), Cys di-oxidation and arginylation (RO2: +188 Da), N/Q arginylation after deamidation (R_deami: +157 Da).

**Related to Figure 3f.**

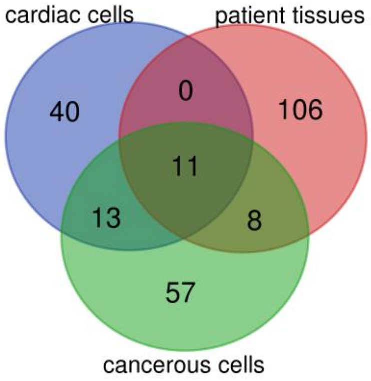


**Fig. S27**. Venn diagram of arginylation sites between sample types.

**Related to Figure 3c.**

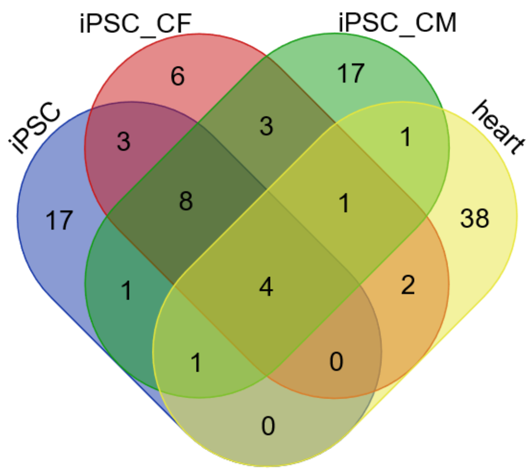


Summary table of arginylation sites.

| sample | number of elements | number of unique elements |
|---|---|---|
| heart | 47 | 47 |
| iPSC | 34 | 34 |
| iPSC_CF | 27 | 27 |
| iPSC_CM | 36 | 36 |
| Overall number of unique elements | | 102 |

Detailed table of sample overlaps.

| sample | total | elements |
|---|---|---|
| heart<br>iPSC<br>iPSC_CF<br>iPSC_CM | 4 | ODO2 D68 R, CATD E64 R, PDIA1 D18 R, CALR E18 R |
| iPSC<br>iPSC_CF<br>iPSC_CM | 8 | SSBP E17 R, H4 D25 R, SAP D405 R, ENPL D22 R, DREB D634 R, ERO1A E24 R, RLA1 N62 R_deami, TBB5 E 3 R, |
| heart<br>iPSC<br>iPSC_CM | 1 | RM12 C45 RO3 |
| heart<br>iPSC_CF<br>iPSC_CM | 1 | MCFD2 E27 R |
| iPSC<br>iPSC_CF | 3 | ACTB D51 R, RLA1 D18 R, BCLF1 E463 R |
| iPSC<br>iPSC_CM | 1 | CH60 E107 R |
| iPSC_CF<br>iPSC_CM | 3 | CH60 Q42 R_deami, MESD E35 R, VIME E425 R |
| heart<br>iPSC_CF | 2 | CATD G65 R, PDIA4 E25 R |
| heart<br>iPSC_CM | 1 | F162A C41 RO3 |
| iPSC | 17 | SPTB2 E1469 R, H15 E 8 R, PTBP1 D 7 R, RT18A E37 R, SFPQ E696 R, PSMD6 L84 R, XRCC5 E583 R, LN28A E18 R, HACD3 P100 R, CH60 C447 R, PGRC1 D49 R, PELP1 E1008 R, COX17 D 6 R, HMGA1 E47 R, HNRPU E259 R, CBS E 4 R, DREB D636 R |

| | | |
|---|---|---|
| iPSC_CF | 6 | MYH9 D1846 R, DJC10 D33 R, ACTB E107 R, AKAP2 D638 R, VIME D90 R, ATAD5 V1449 R |
| iPSC_CM | 17 | TXND5 E40 R, LDB3 N197 R_deami, HS90B E539 R, MPPB Q46 R_deami, CH60 G43 R, MYOZ2 D255 R, CWC15 Q45 R_deami, RM12 C45 R, ACTC E318 R, CH60 D100 R, ACTC D53 R, ACTC E109 R, LDB3 E257 R, ACTA E318 R, ACTA E109 R, MYH6 E73 R, PDLI5 E215 R |
| heart | 38 | PDLI5 E297 R, MLRV E22 R, ATP5J N33 R_deami, APOA1 D25 R, TMM11 D163 R, HBA D48 R, MYH7 D1229 R, CLUS D23 R, MYH7 D1008 R, MYH7 E1376 R, LMAN2 D45 R, JTB E31 R, TNNI3 E165 R, MYPT2 D816 R, MYH7 E1902 R, CASQ2 E20 R, SRCA E21 R, MYH7 E1461 R, MTDC E36 R, TNNT2 D108 R, ATIF1 E42 R, MYH7 E73 R, MYH7 E1489 R, A1AT E25 R, ATIF1 E65 R, CAVN2 Q161 R_deami, MYL3 E131 R, MYH7 E927 R, MLRV N154 R_deami, AT5G1 D62 R, MYH7 E1821 R, MITOK C37 RO3, SCMC1 Q16 R_deami, ACTC C259 RO2, KTN1 D238 R, ATP5H D129 R, LETM1 D116 R, CALD1 E161 R |

**Fig. S28**. Venn diagram of arginylation sites in cardiac samples. Summary and detailed overlapping sites are listed. iPSC, induced pluripotent stem cell. CF, cardiac fibroblasts. CM, cardiomyocytes.

**Related to Figure 3c.**

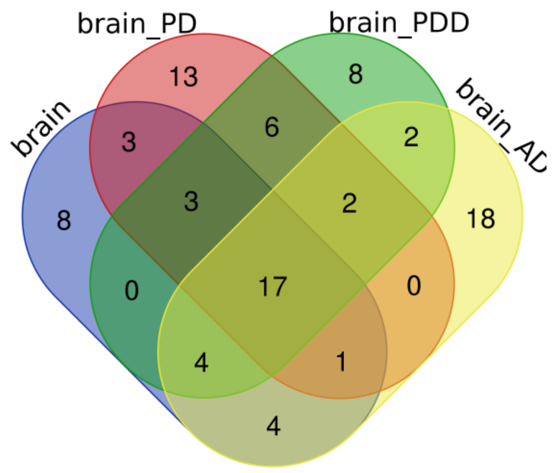


Summary table of arginylation sites.

| sample | number of elements | number of unique elements |
|---|---|---|
| brain (brain_117849 and 117504) | 40 | 40 |
| brain_AD (brain_110007) | 48 | 48 |
| brain_PD (brain_112568 ) | 45 | 45 |
| brain_PDD (brain_108180) | 42 | 42 |
| Overall number of unique elements | | 89 |

Detailed table of sample overlaps.

| sample | total | elements |
|---|---|---|
| brain brain_AD brain_PD brain_PDD | 17 | GFAP E391 R CATD G65 R F162A C41 RO3 MBP D168 R NCAN E23 R AP180 N767 R_deami TAU N644 R_deami MBP E217 R SPTN1 E1124 R A1AT E25 R NFM E496 R NFL D468 R GFAP D417 R CATD E64 R NFM E497 R NEUM E89 R CALR E18 R |
| brain brain_PD brain_PDD | 3 | RD23B E136 R CLUS D23 R PGCB D23 R |
| brain brain_AD brain_PD | 1 | NHRF1 E121 R |
| brain brain_AD brain_PDD | 4 | MBP N226 R_deami MBP D216 R NFH E469 R NFM E455 R |
| brain_AD brain_PD brain_PDD | 2 | MTAP2 Q431 R_deami PDIA1 D18 R |
| brain brain_PD | 3 | NFM E763 R C99L2 D26 R CDY1 D285 R |
| brain brain_AD | 4 | NFL E469 R SAP G407 R SAP D405 R NFM E454 R |
| brain_PD brain_PDD | 6 | NT5D3 C44 RO3 TBB4A D67 R BACH C36 RO3 ODO2 D68 R ERO1A E24 R ANK2 E1760 R |
| brain_AD brain_PDD | 2 | NFM E599 R PLEC E4050 R |
| brain | 8 | MAP1A E1580 R MBP D279 R NFM E470 R MAP1B Q907 R_deami AQP4 E279 R HOOK1 D659 R SCN1A V1579 RO2 NFL E509 R |

| | | |
|---|---|---|
| brain_PD | 13 | TBA4A D69 R NEUG C 9RO3 TBA1B D251 R PTGDS E25 R PCDH1 D944 R PGCB E412 R ATS15 D749 R HBA D65 R_deami TBA1B D69 R ROA2 D161 R SCG1 D88 R NECP1 C162 RO3 LETM1 D116 R |
| brain_PDD | 8 | PACN1 N330 R_deami KS6A1 D37 R PGCB E26 R ATIF1 D28 R TYB4 D 6 R PDIA4 E25 R UTP18 E374 R RM12 C45 RO3 |

**Fig. S29**. Venn diagram of arginylation sites in brain samples. Summary and detailed overlapping sites are listed. PD, Parkinson's disease. PDD, PD with dementia. AD, Alzheimer's disease. brain, healthy brains including brain_117849 and brain_117504. brain_AD, brain_110007. brain_PD, brain_112568. brain_PDD, brain_108180.

**Related to Figure 3c.**

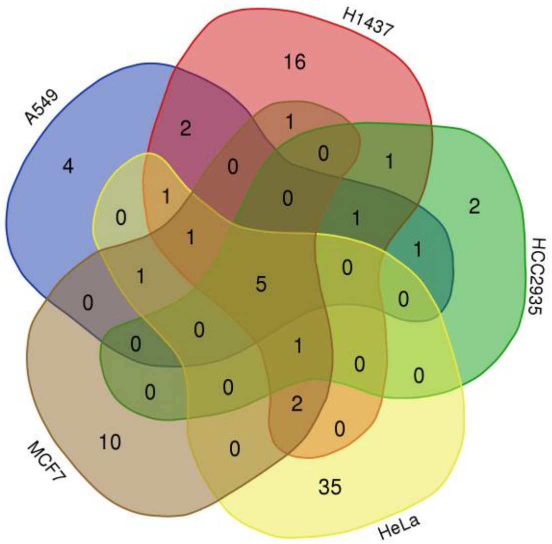


Summary table of arginylation sites.

| sample | number of elements | number of unique elements |
|---|---|---|
| A549 | 16 | 16 |
| H1437 | 31 | 31 |
| HCC2935 | 11 | 11 |
| HeLa | 46 | 46 |
| MCF7 | 21 | 21 |
| Overall number of unique elements | | 84 |

Detailed table of sample overlaps.

| sample | total | elements |
|---|---|---|
| A549 H1437 HCC2935 HeLa MCF7 | 5 | CATD E64 R ERO1A E24 R PDIA1 D18 R CALR E18 R RM12 C45 RO3 |
| A549 H1437 HeLa MCF7 | 1 | F162A C41 RO3 |
| H1437 HCC2935 HeLa MCF7 | 1 | SSBP E17 R |
| A549 H1437 HCC2935 | 1 | CH60 E107 R |
| A549 H1437 HeLa | 1 | SAP D405 R |
| A549 HeLa MCF7 | 1 | MCFD2 E27 R |
| H1437 HeLa MCF7 | 2 | LMAN2 D45 R PDIA4 E25 R |
| A549 H1437 | 2 | CH60 D49 R BCLF1 E463 R |
| A549 HCC2935 | 1 | CD99 D23 R |
| H1437 HCC2935 | 1 | S10AE I68 RO3 |
| H1437 MCF7 | 1 | ODO2 D68 R |
| A549 | 4 | NPM D178 R CWC15 Q45 R_deami NPM D179 R NPM D180 R |

| | | |
|---|---|---|
| H1437 | 16 | H4 D25 R SRRT D866 R RL1D1 D319 R RS27 D 6 R MESD E35 R CH60 D209 R ATIF1 E65 R RLA1 D18 R PGRC1 D49 R ACTB E107 R SRRT D868 R RM12 C45 RO2 SF3B1 D93 R TFF1 E25 R NOP16 E77 R HNRPU E259 R |
| HCC2935 | 2 | NPC2 E20 R PSD12 G331 R |
| HeLa | 35 | TXND5 E40 R NUCL E628 R SNX17 E463 R HNRPK D279 R BI1 I 8R_deami ACTB D51 R HSP7C G372 R HNRPK N306 R RON D1232 R RS7 E188 R CALX E275 R MYH9 D1136 R ACTB E241 R G3P D296 R VATD E242 R ROA1 E93 R TBA1B D218 R VIME D451 R EF1D E27 R HNRPL D284 R COX41 D43 R HNRPK Q358 R_deami ILF2 D347 R RL23A E 8 R CNBP Q44 R_deami FCN1 D120 R G3P D315 R VIME E425 R FUBP1 N369 R_deami HNRPC D123 R H2B1K E94 R ACLY E830 R IF4H D196 R RLA1 D19 R CALU D62 R |
| MCF7 | 10 | CATD G65 R GLYM Q27 R_deami F205A E1023 R JTB E31 R MTDC E36 R MITOK C37 RO3 GATB E42 R SCMC1 Q16 R_deami KTN1 D238 R LETM1 D116 R |
| A549 H1437 HCC2935 HeLa MCF7 | 5 | CATD E64 R ERO1A E24 R PDIA1 D18 R CALR E18 R RM12 C45 RO3 |

**Fig. S30**. Venn diagram of arginylation sites in cancer cells. Summary and detailed overlapping sites are listed.

**Related to Figure 3c.**

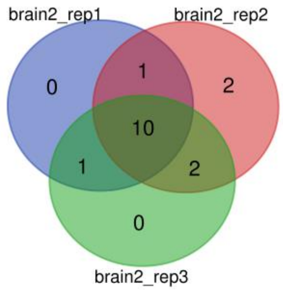


| Names | total | elements |
|---|---|---|
| brain2_rep1<br>brain2_rep2<br>brain2_rep3 | 10 | NFL D468 R CATD G65 R GFAP D417 R MBP N226 R_deami SAP D405 R MBP D168 R MBP E217 R NFM E454 R CALR E18 R AP180 N767 R_deami |
| brain2_rep1<br>brain2_rep2 | 1 | GFAP E391 R |
| brain2_rep1<br>brain2_rep3 | 1 | SAP G407 R |
| brain2_rep2<br>brain2_rep3 | 2 | NFL E469 R A1AT E25 R |
| brain2_rep2 | 2 | CDY1 D285 R CATD E64 R |

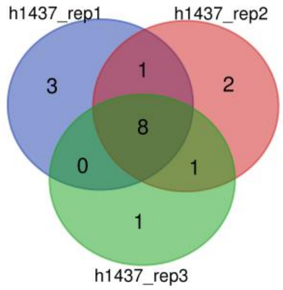


| sample | total | elements |
|---|---|---|
| h1437_rep1<br>h1437_rep2<br>h1437_rep3 | 8 | ACTB E107 R SSBP E17 R PDIA4 E25 R PDIA1 D18 R ERO1A E24 R ATIF1 E65 R CALR E18 R RM12 C45 RO3 |
| h1437_rep1<br>h1437_rep2 | 1 | H4 D25 R |
| h1437_rep2<br>h1437_rep3 | 1 | CATD E64 R |
| h1437_rep1 | 3 | MESD E35 R CH60 D49 R RM12 C45 RO2 |
| h1437_rep2 | 2 | RL1D1 D319 R SF3B1 D93 R |
| h1437_rep3 | 1 | BCLF1 E463 R |

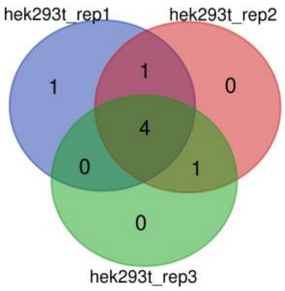


| sample | total | elements |
|---|---|---|
| hek293t_rep1 hek293t_rep2 hek293t_rep3 | 4 | SSBP E17 R LETM1 D116 R RM12 C45 RO3 CALR E18 R |
| hek293t_rep1 hek293t_rep2 | 1 | PDIA1 D18 R |
| hek293t_rep2 hek293t_rep3 | 1 | PDIA4 E25 R |
| hek293t_rep1 | 1 | F162A C41 RO3 |

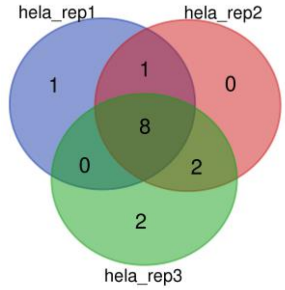


| sample | total | elements |
|---|---|---|
| hela_rep1 hela_rep2 hela_rep3 | 8 | ACTB E241 R G3P D296 R SSBP E17 R PDIA1 D18 R TBA1B D218 R H2B1K E94 R CALR E18 R RM12 C45 RO3 |
| hela_rep1 hela_rep2 | 1 | ERO1A E24 R |
| hela_rep2 hela_rep3 | 2 | RL23A E 8 R EF1D E27 R |
| hela_rep1 | 1 | SAP D405 R |
| hela_rep3 | 2 | CNBP Q44 R_deami F162A C41 RO3 |

**Fig. S31**. Venn diagram of arginylation sites in replicates of 4 biological samples. H1437, HEK293T, and HeLa cells were used to compare sites from biological replicates (*n* = 3). Brain_c2 (117504) was used to compare sites from technical replicates (*n* = 3). Summary and detailed overlapping sites are listed.

**Related to Figure 3c.**

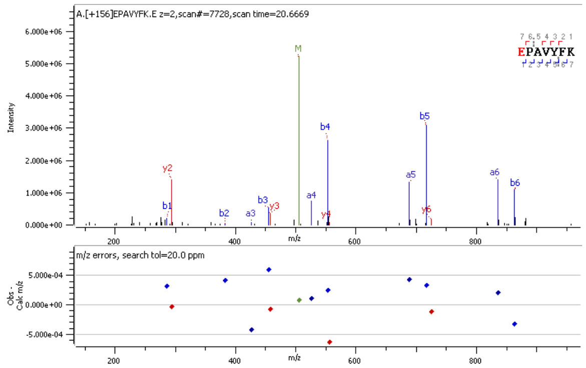


Standard peptide MS2 of CALR E18 R

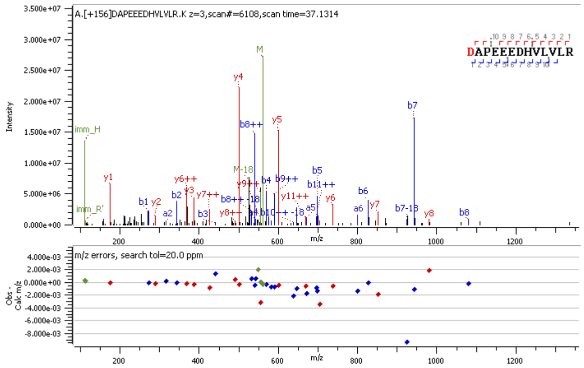


Standard peptide MS2 of PDIA1 D18 R

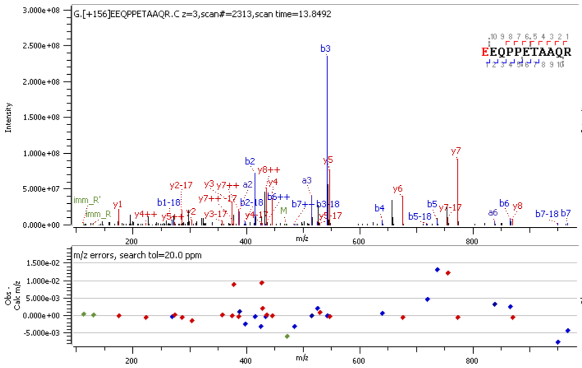


Standard peptide MS2 of ERO1A E24 R

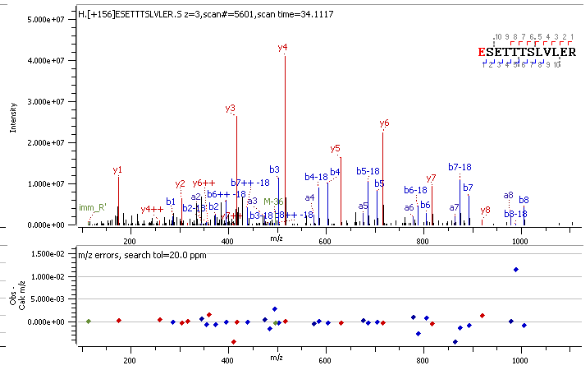


Standard peptide MS2 of SSBP E17 R

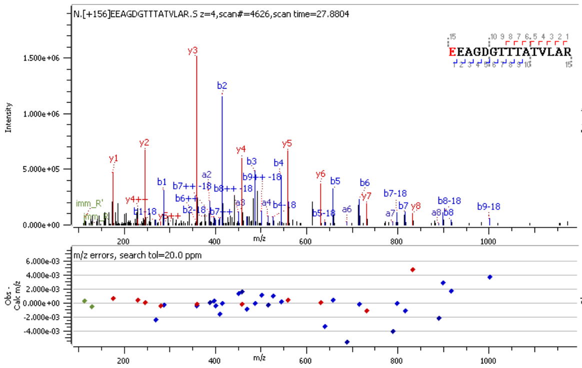


Standard peptide MS2 of CH60 E107 R

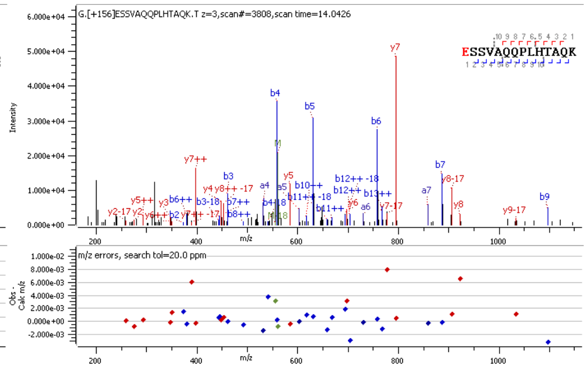


Standard peptide MS2 of GATB E42 R

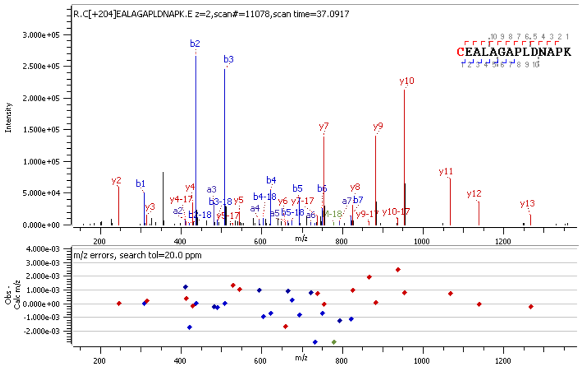


Standard peptide MS2 of RM12 C45 RO3

**Fig. S32**. MS2 spectra of 7 synthetic arginylated peptides for site validation. Arg0 modification, +156 Da. Arg0 modification and Cys tri-oxidation, +204 Da.

**Related to Figure 4a.**

**a**,

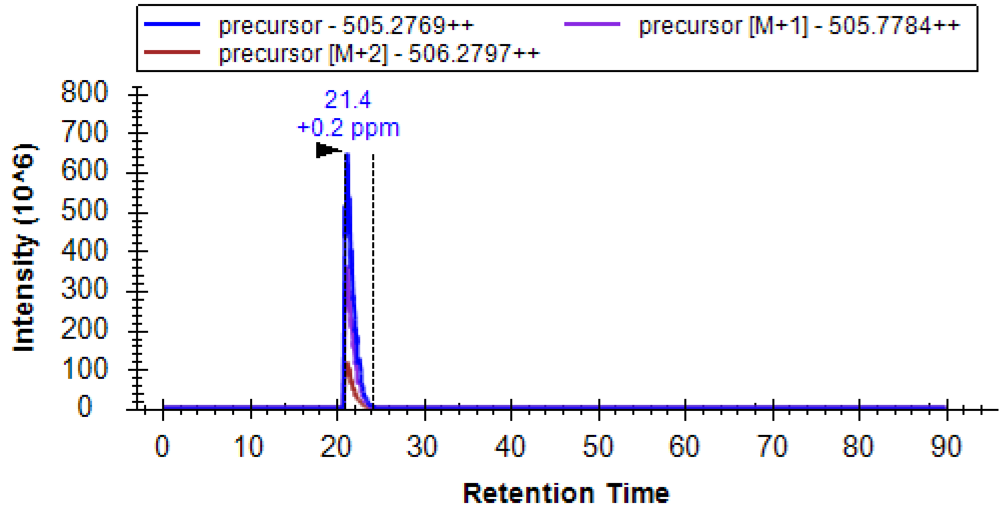


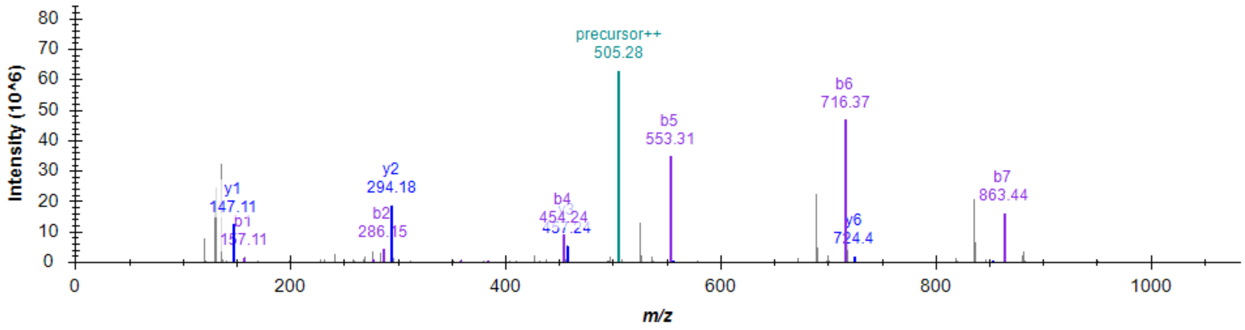


Retention time (21.4 min), EIC, MS2 of REPAVYFK.

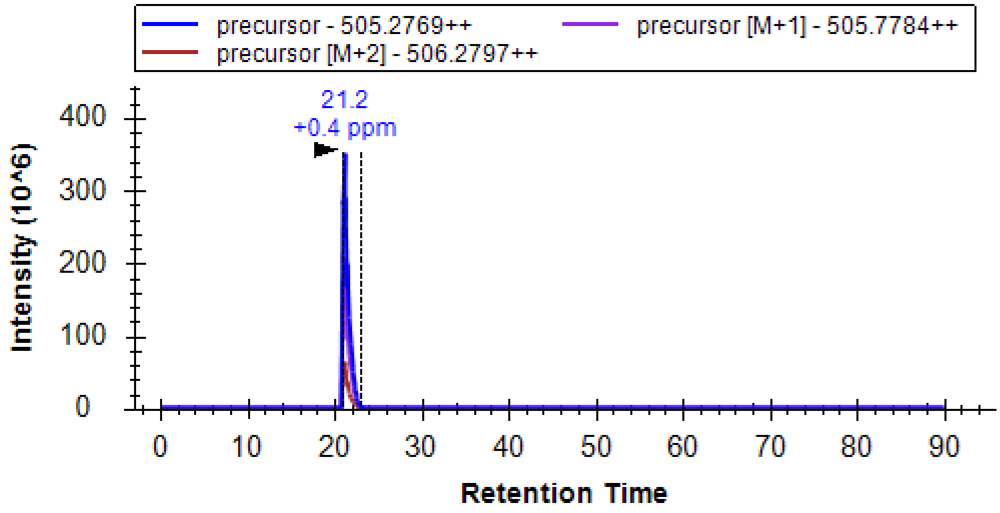

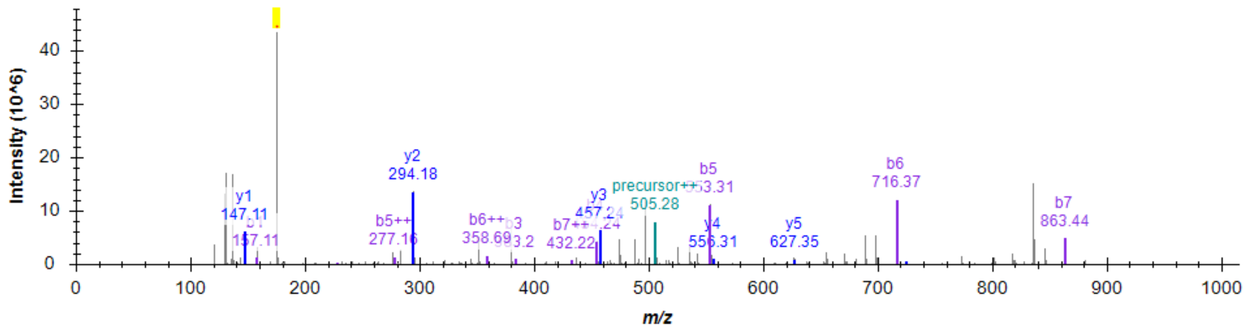


Retention time (21.2 min), EIC, MS2 of E(R)PAVYFK. The signature ion at m/z 175.1190 from Arg residue on the side chain is highlighted.

These two peptides show similar retention times and thus can not be distinguished clearly by LC behaviors.

**b**,

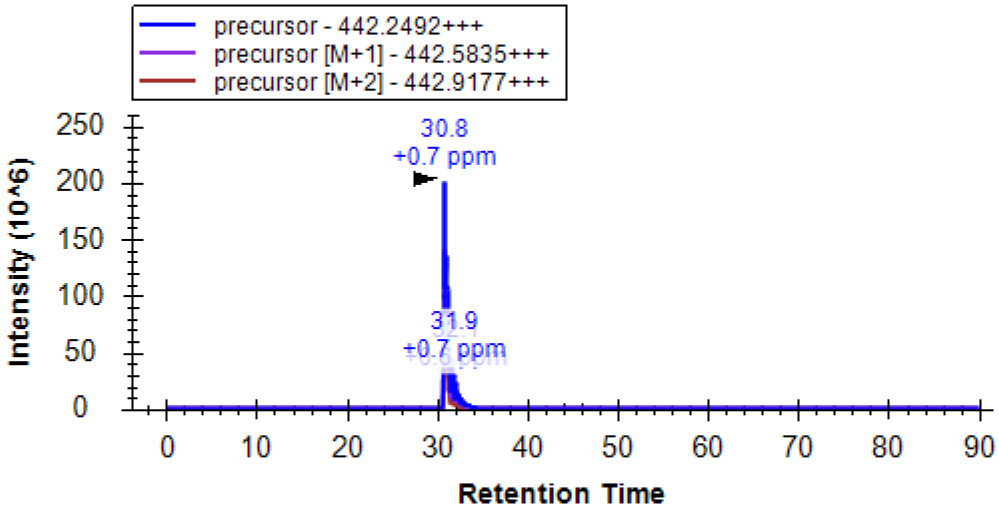


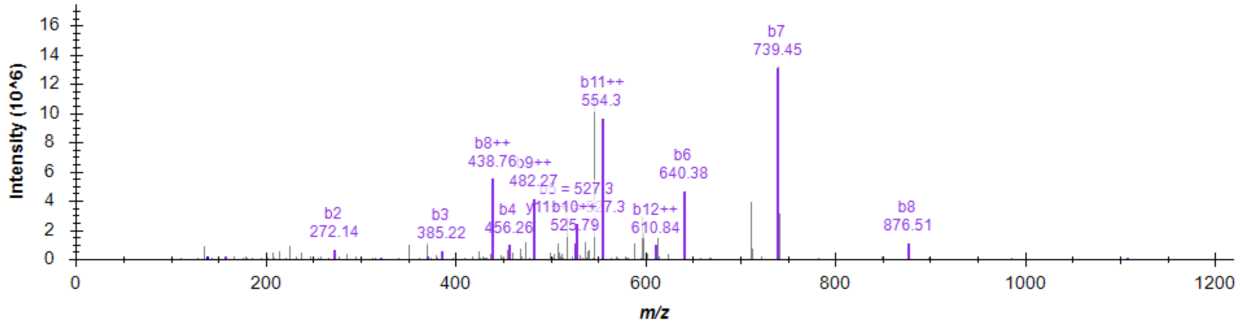


Retention time (30.8 min), EIC, MS2 of RDIAALVHSSGNleS-NH2.

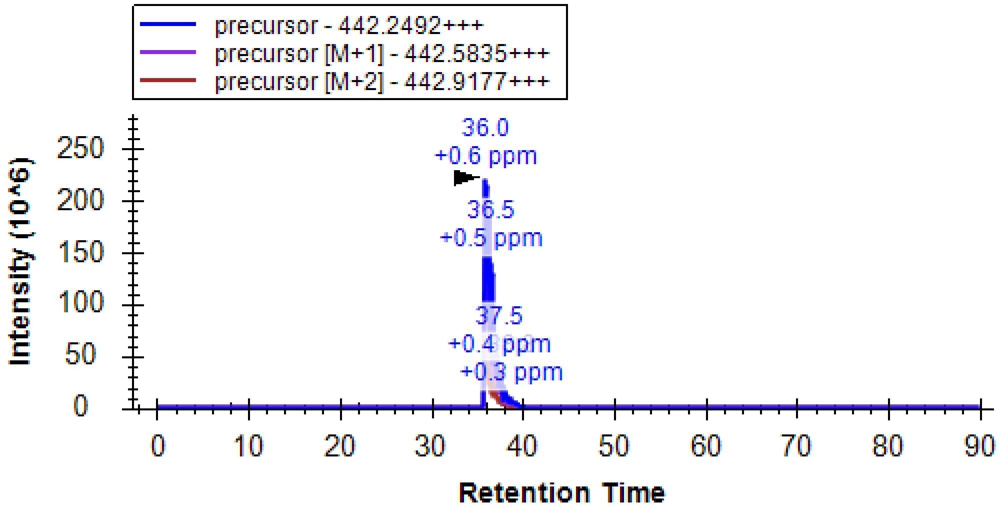


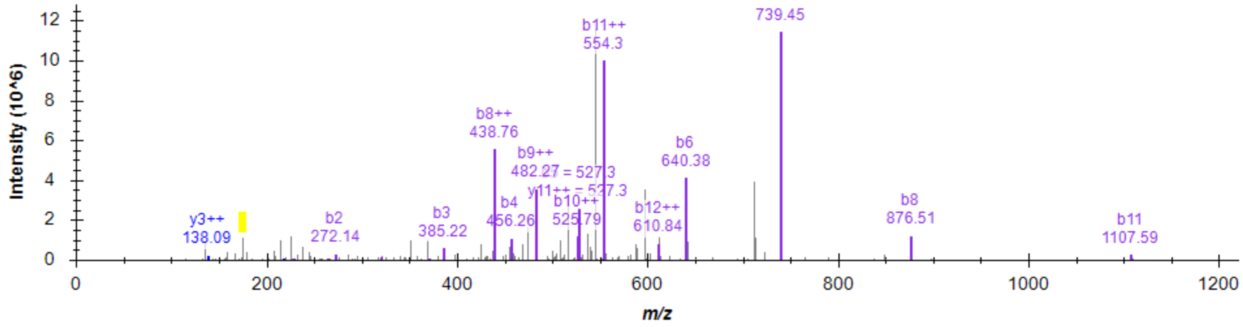


Retention time (36.0 min), EIC, MS2 of D(R)IAALVHSSGNleS-NH2. The signature ion at m/z 175.1190 from Arg residue on the side chain is highlighted.

These two peptides show different retention times and thus can be distinguished by LC behaviors.

**c**,

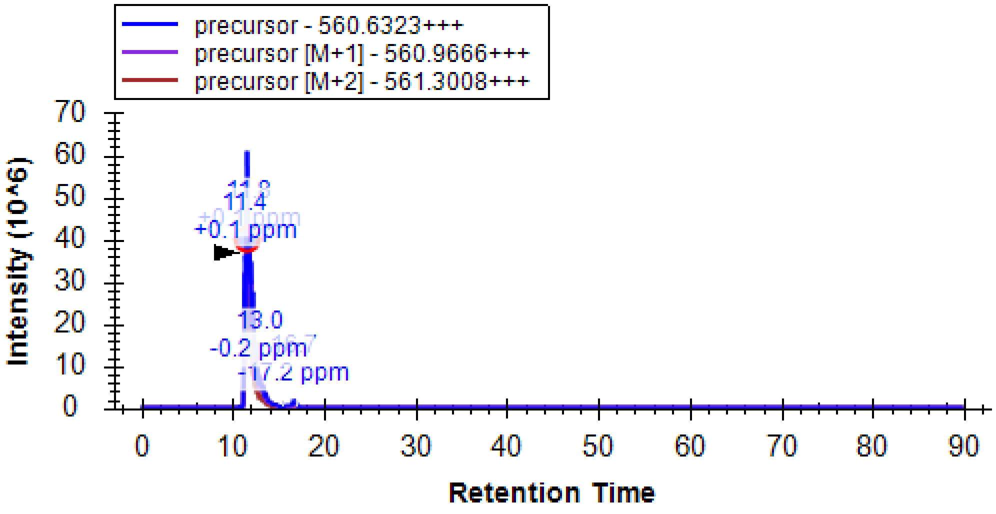

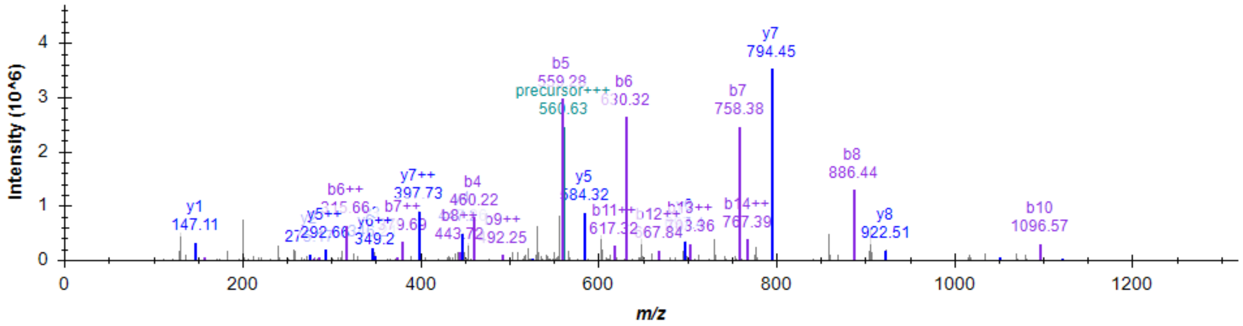


Retention time (11.6 min), EIC, MS2 of RESSVAQQPLHTAQK.

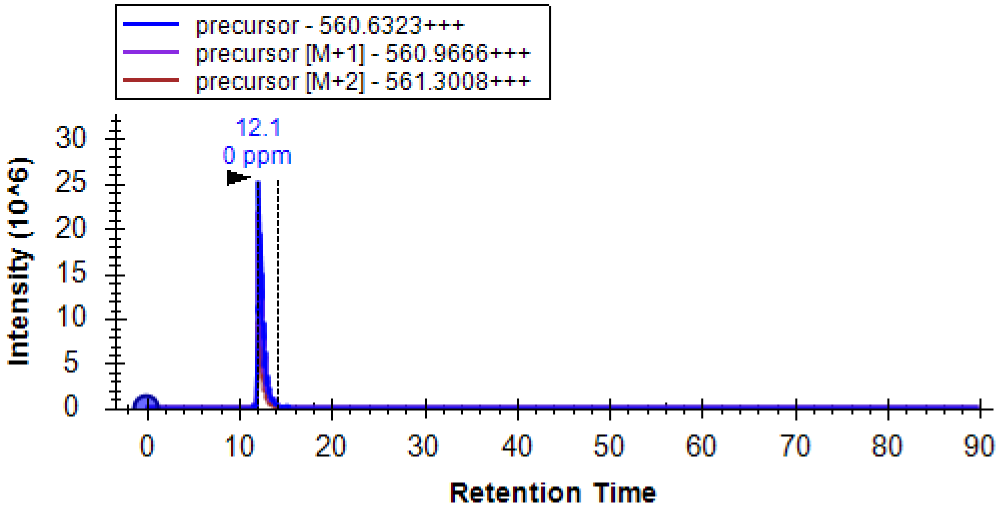


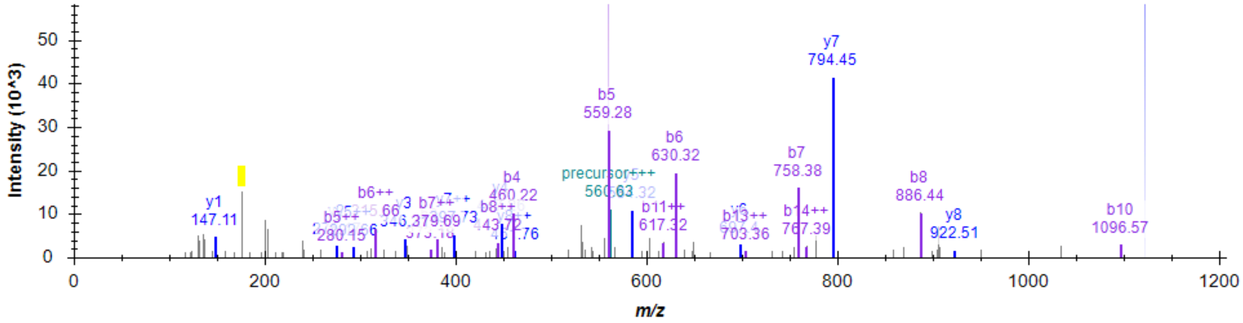


Retention time (12.1 min), EIC, MS2 of E(R)SSVAQQPLHTAQK. The signature ion at m/z 175.1190 from Arg residue on the side chain is highlighted.

These two peptides show close but different retention times and thus can be distinguished by LC behaviors.

**d**,

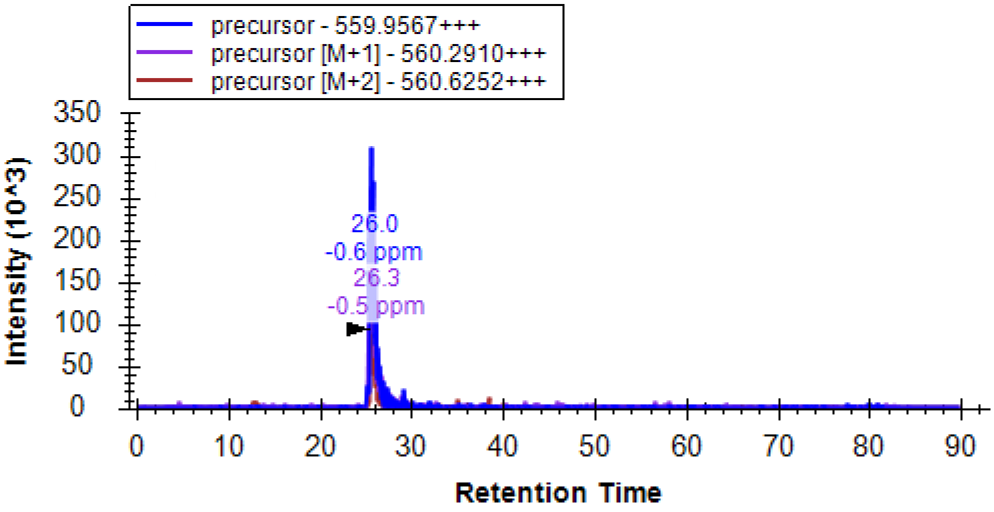


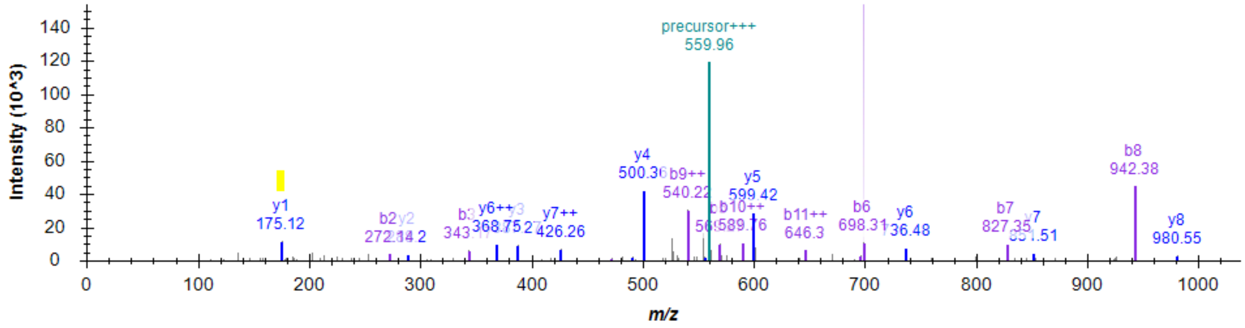


Retention time (25.6 min), EIC, MS2 of RDAPEEEDHVLVLR. The signature ion at m/z 175.1190 from C-term Arg residue is highlighted.

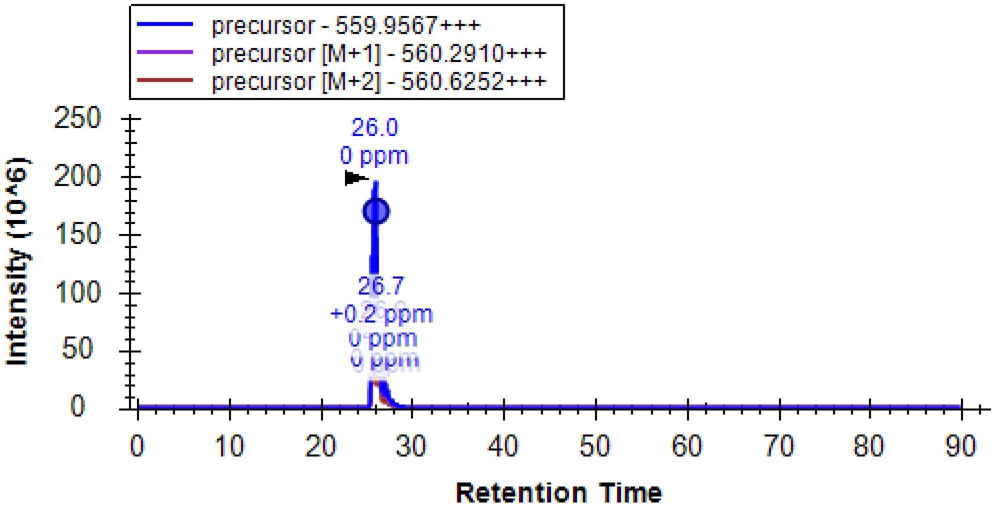

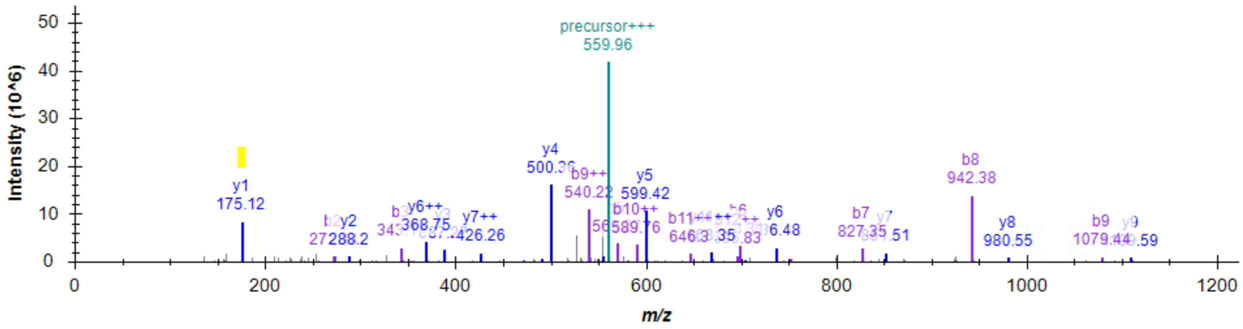


Retention time (26.0 min), EIC, MS2 of D(R)APEEEDHVLVLR. The signature ion at m/z 175.1190 from C-term Arg residue and Arg residue on the side chain is highlighted.

These two peptides show close but different retention times and thus can be distinguished by LC behaviors. Due to the presence of C-term Arg residue, m/z 175.1190 is not unique ion to side chain arginylation. Still, it shows a higher intensity in side chain arginylation than in N-term arginylation.

**e**,

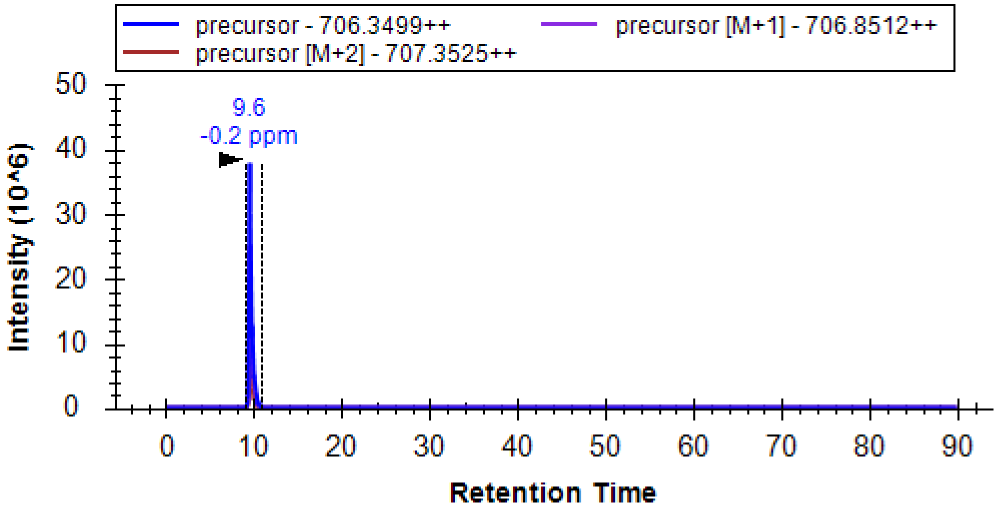


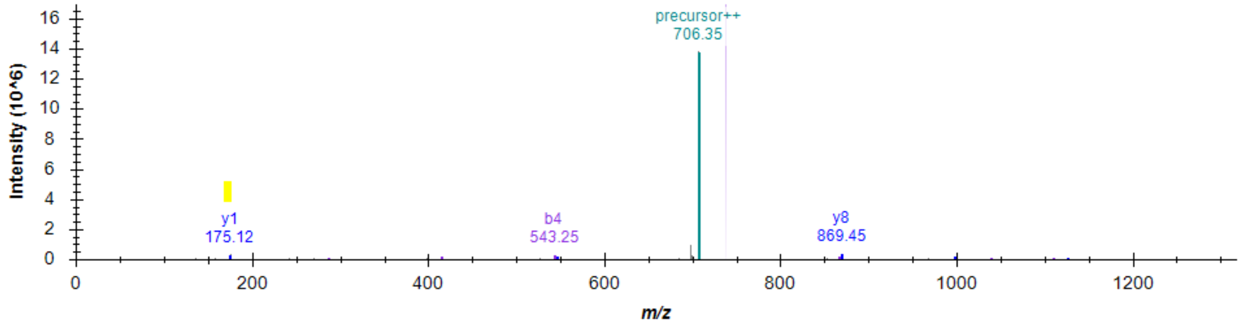


Retention time (9.6 min), EIC, MS2 of REEQPPETAAQR. The signature ion at m/z 175.1190 from C-term Arg residue is highlighted.

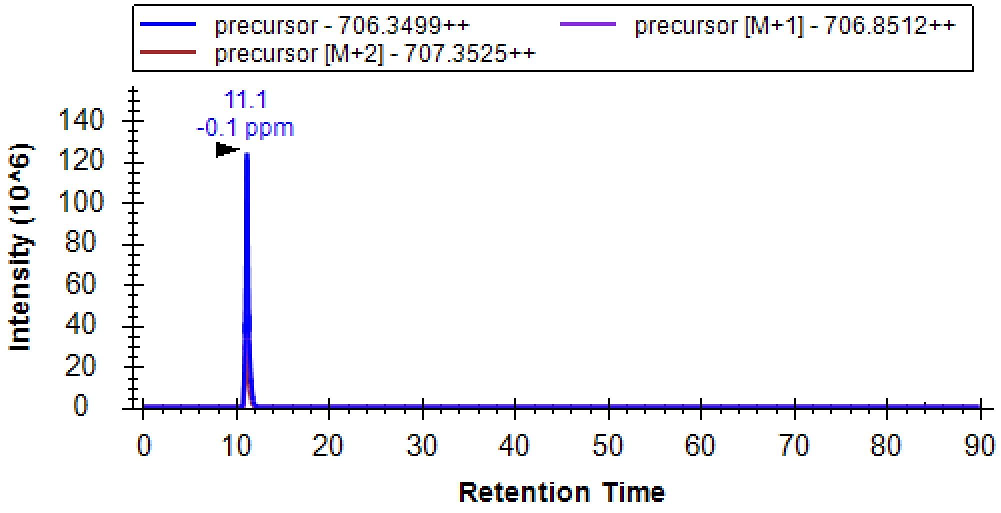


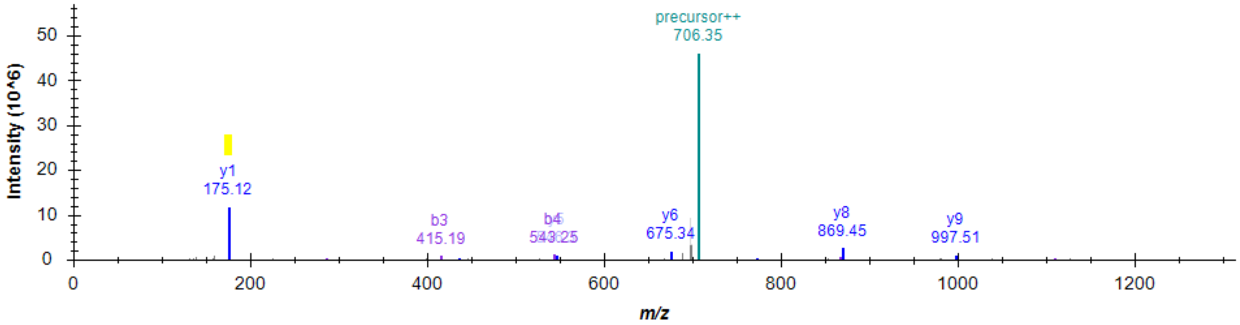


Retention time (11.1 min), EIC, MS2 of E(R)EQPPETAAQR. The signature ion at m/z 175.1190 from C-term Arg residue and Arg residue on the side chain is highlighted.

These two peptides show different retention times and thus can be distinguished by LC behaviors. Due to the presence of C-term Arg residue, m/z 175.1190 is not a unique ion to side chain arginylation. Still, it shows a higher intensity in side chain arginylation than in N-term arginylation.

**f**,

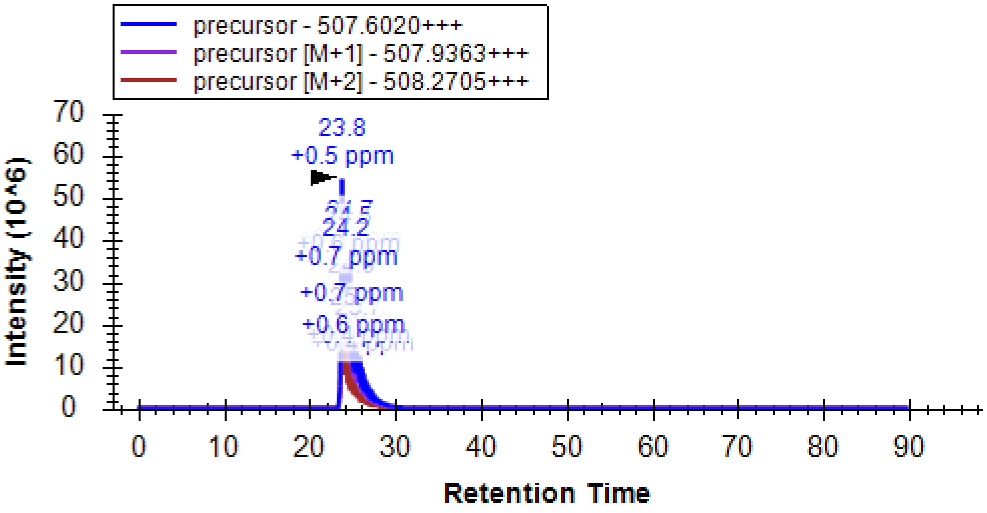

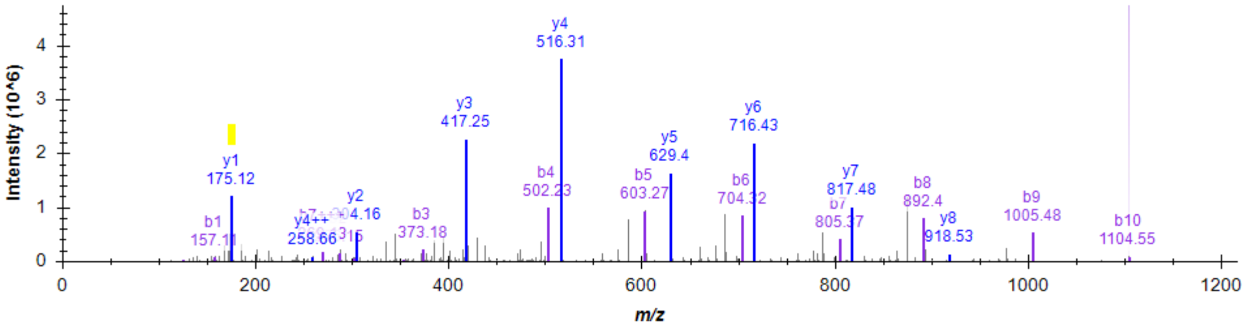


Retention time (23.8 min), EIC, MS2 of RESETTTSLVLER. The signature ion at m/z 175.1190 from C-term Arg residue is highlighted.

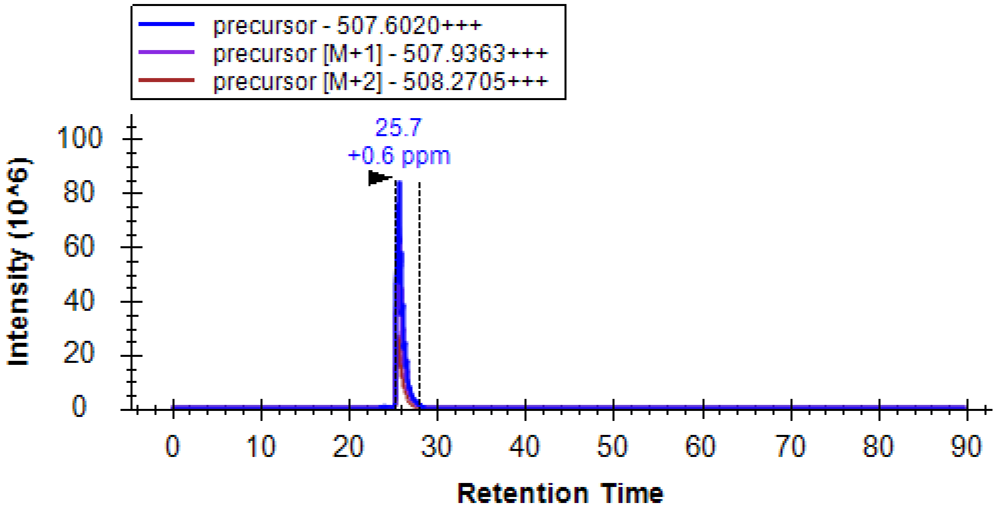


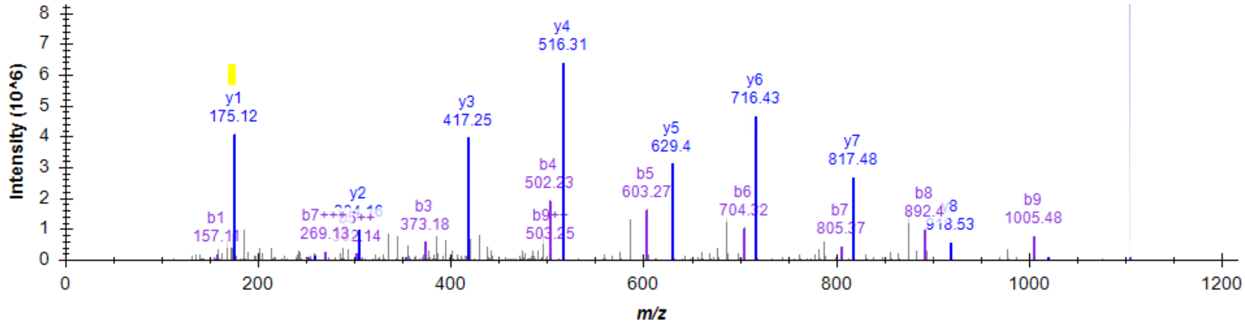


Retention time (25.7 min), EIC, MS2 of E(R)SETTTSLVLER. The signature ion at m/z 175.1190 from C-term Arg residue and Arg residue on the side chain is highlighted.

These two peptides show different retention times and thus can be distinguished by LC behaviors. Due to the presence of C-term Arg residue, m/z 175.1190 is not unique ion to side chain arginylation. Still, it shows a higher intensity in side chain arginylation than in N-term arginylation.

**g**,

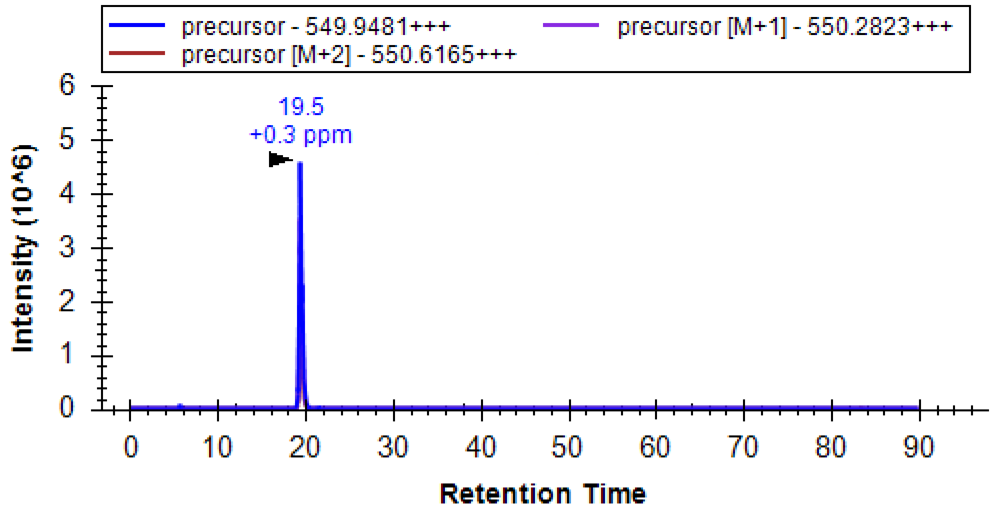


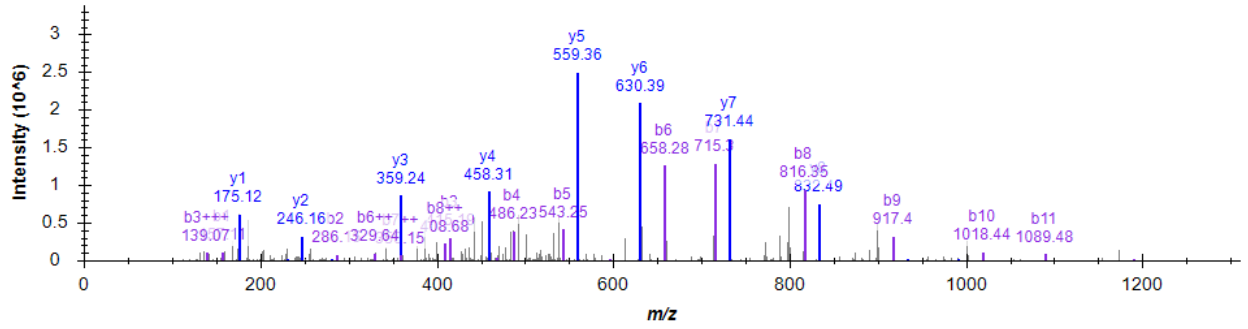


Retention time (19.5 min), EIC, MS2 of REEAGDGTTTATVLAR. The signature ion at m/z 175.1190 from C-term Arg residue is highlighted.

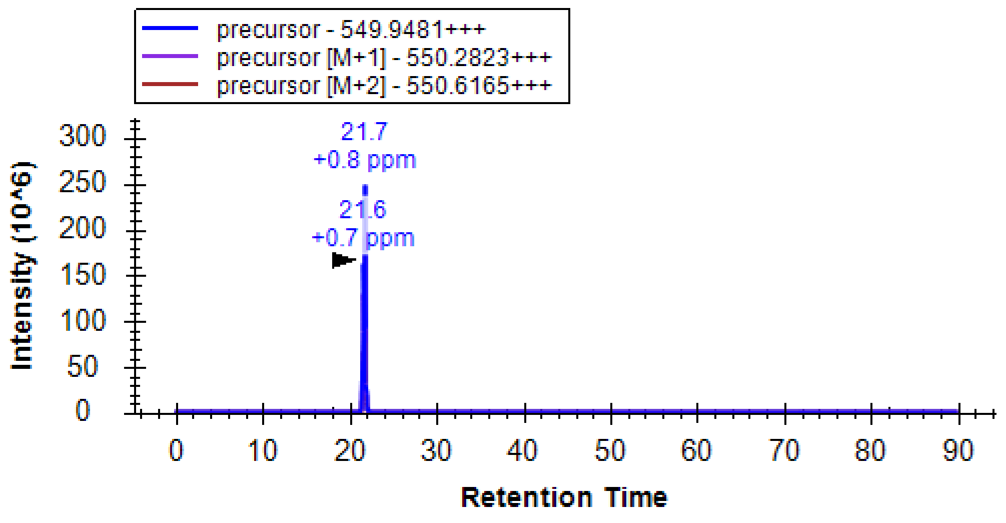

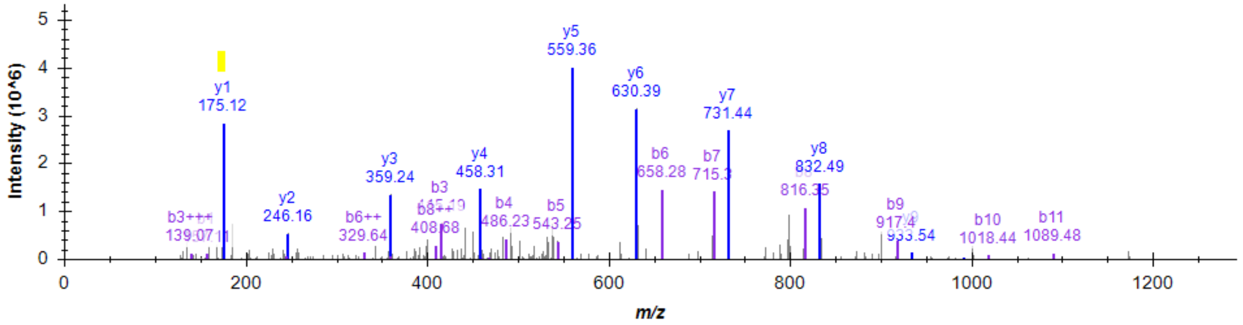


Retention time (21.7 min), EIC, MS2 of E(R)EAGDGTTTATVLAR. The signature ion at m/z 175.1190 from C-term Arg residue and Arg residue on the side chain is highlighted.

These two peptides show different retention times and thus can be distinguished by LC behaviors. Due to the presence of C-term Arg residue, m/z 175.1190 is not unique ion to side chain arginylation. Still, it shows a higher intensity in side chain arginylation than in N-term arginylation.

**Fig. S33**. Chromatographic and MS differentiation of N-terminal and side chain arginylation on N-terminal D/E residues in 7 peptides. **a**, chromatographic and MS behaviors of REPAVYFK and E(R)PAVYFK. **b**, chromatographic and MS behaviors of RDIAALVHSSGNleS-NH2 and D(R)IAALVHSSGNleS-NH2. **c**, chromatographic and MS behaviors of RESSVAQQPLHTAQK and E(R)SSVAQQPLHTAQK. **d**, chromatographic and MS behaviors of RDAPEEEDHVLVLR and D(R)APEEEDHVLVLR. **e**, chromatographic and MS behaviors of REEQPPETAAQR and E(R)EQPPETAAQR. **f**, chromatographic and MS behaviors of RESETTTSLVLER and E(R)SETTTSLVLER. **g**, chromatographic and MS behaviors of REEAGDGTTTATVLAR and E(R)EAGDGTTTATVLAR.

**Related to Figure 4a.**

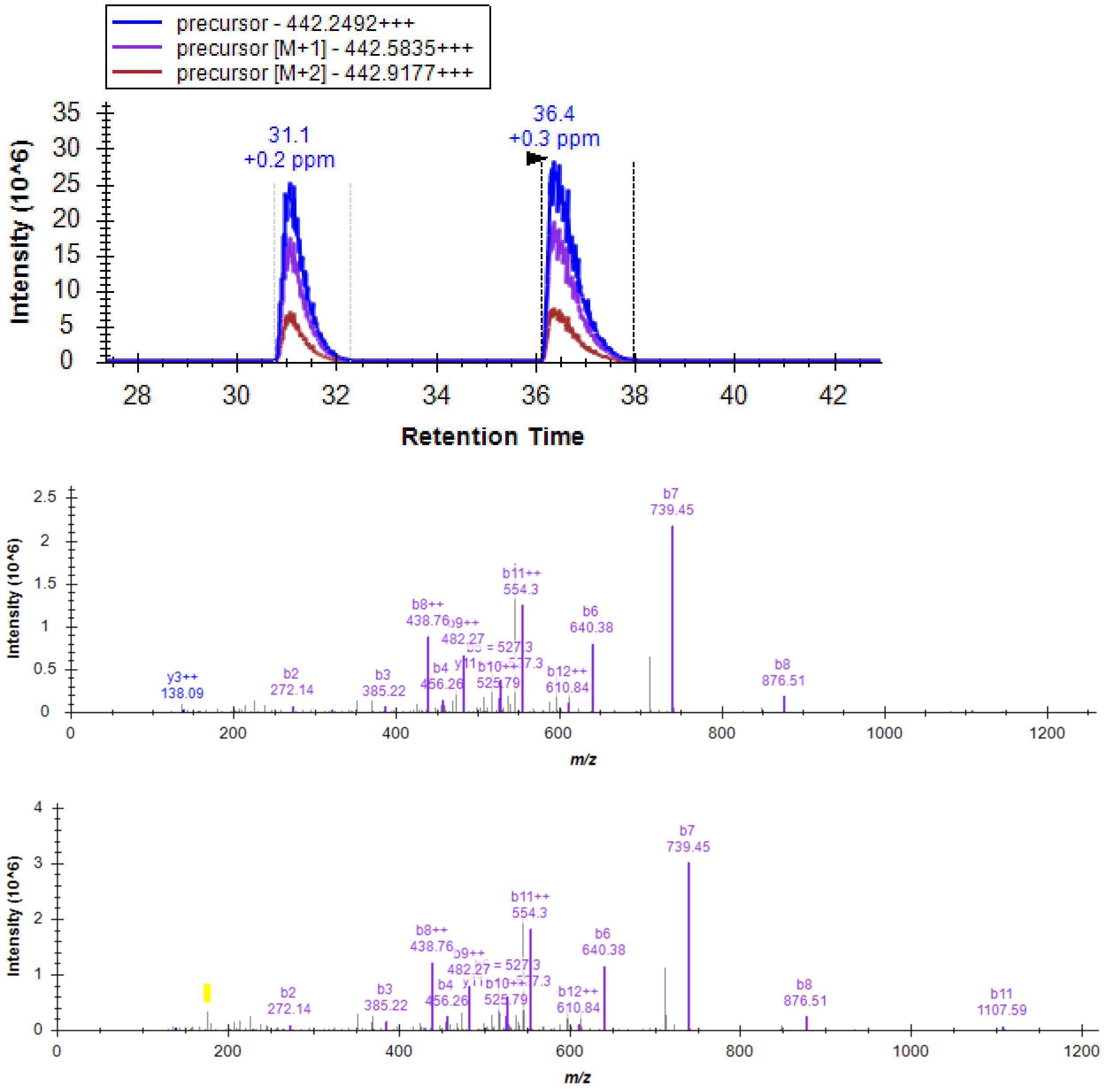


**Fig. S34**. Confirmation of arginylation product RDIAALVHSSGNleS-NH2 (retention time = 31.1 min). Retention time (36.4 min), EIC, MS2 of D(R)IAALVHSSGNleS-NH2 which was added to the arginylation assay using substrate DIAALVHSSGNleS-NH2. The signature ion at m/z 175.1190 from Arg residue on the side chain is highlighted. These two peptides show different retention times and thus can be distinguished by LC behaviors.

**Related to Figure 4a.**

**a**,

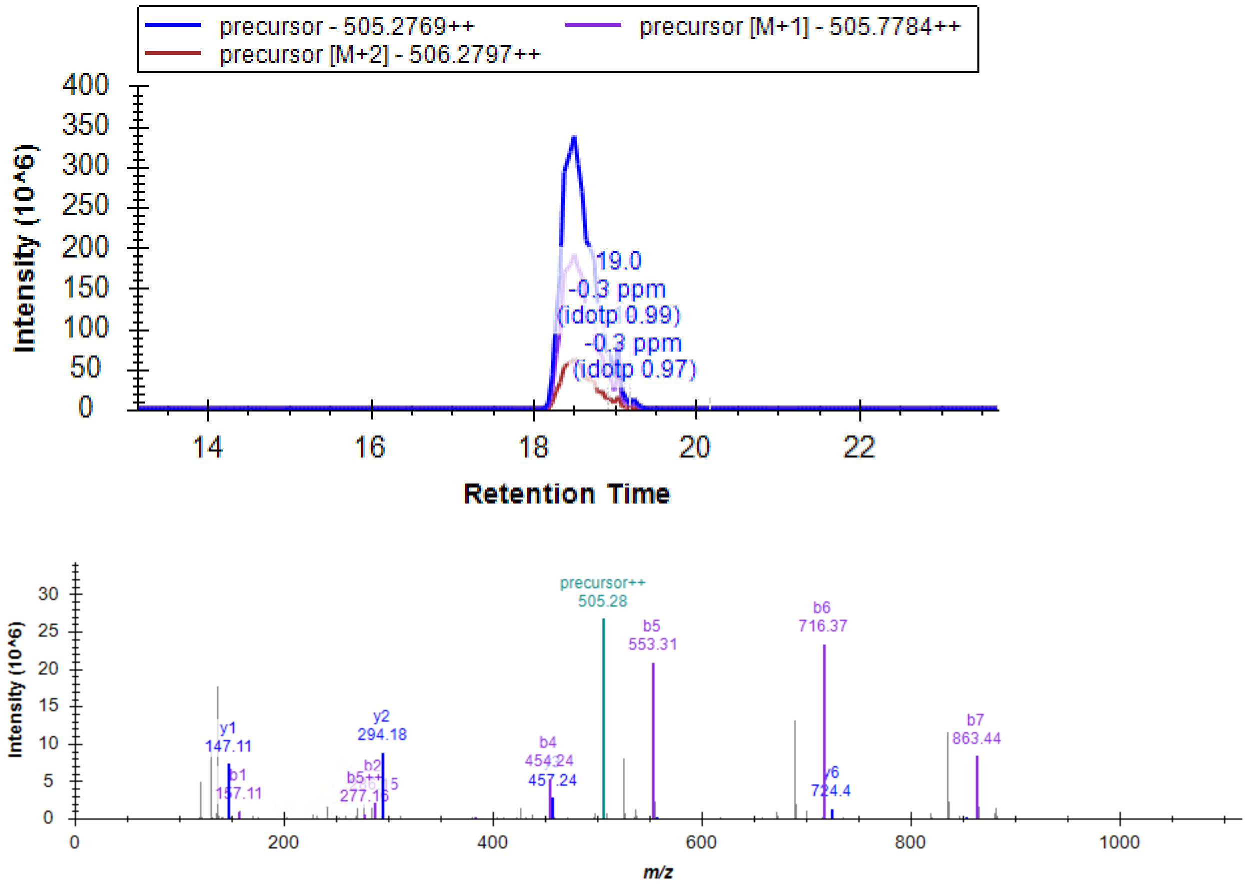
precursor - 505.2769++
precursor [M+1] - 505.7784++
precursor [M+2] - 506.2797++
Intensity (10^6)
Retention Time
19.0
-0.3 ppm
(idotp 0.99)
-0.3 ppm
(idotp 0.97)
precursor++
505.28
b5
553.31
b6
716.37
b7
863.44
y1
147.11
b1
157.11
y2
294.18
b4
454.24
457.24
y6
724.4
m/z

**b**,

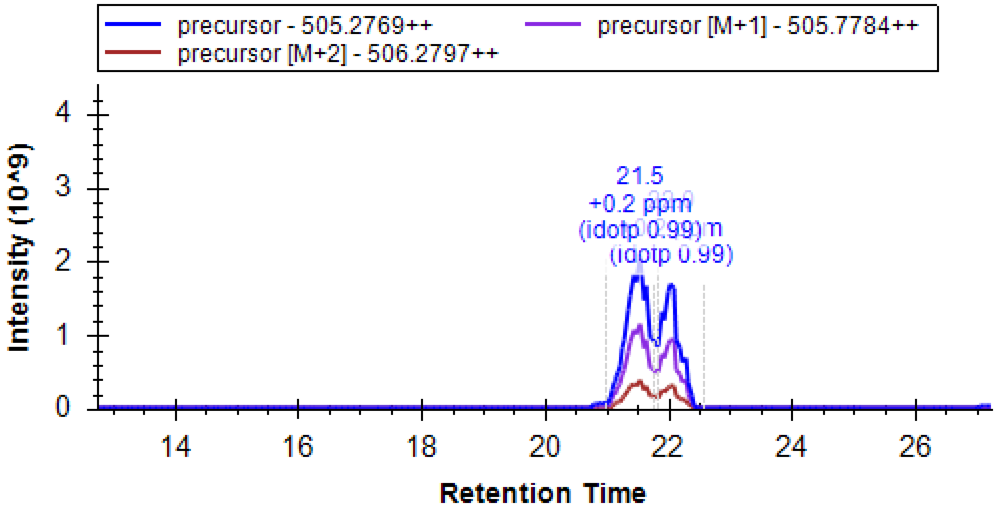
precursor - 505.2769++
precursor [M+1] - 505.7784++
precursor [M+2] - 506.2797++
Intensity (10^9)
Retention Time
21.5
+0.2 ppm
(idotp 0.99)
(idotp 0.99)

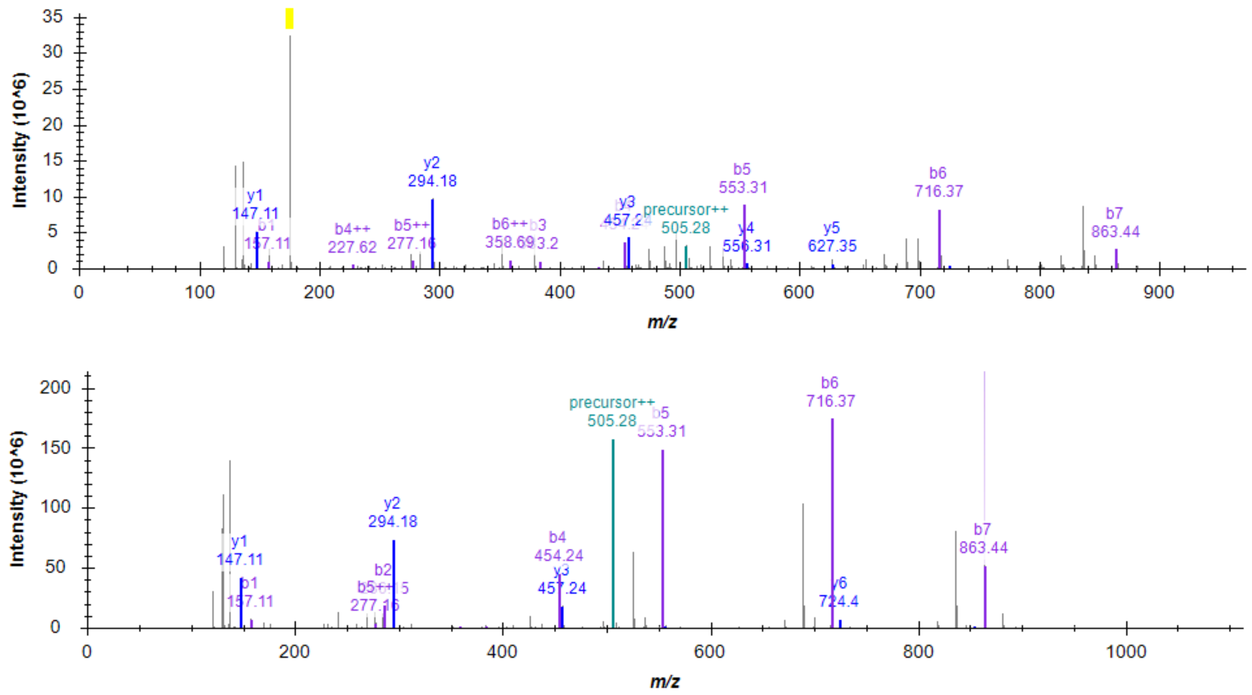


**Fig. S35**. Confirmation of arginylated peptide REPAVYFK from CALR protein arginylation by ATE1 assay. **a**, EIC, MS2 of REPAVYFK from CALR arginylation by ATE1 assay. MS2 spectra of REPAVYFK did not show the signature ion at m/z 175.1190. **b**, E(R)PAVYFK peptide was added to the CALR sample. The MS2 spectra show the signature ion at m/z 175.1190 (highlighted) from Arg residue on the side chain.

**Related to Figure 4a.**

**a**, EIC of REPAVYFK and added E(R)PAVYFK.

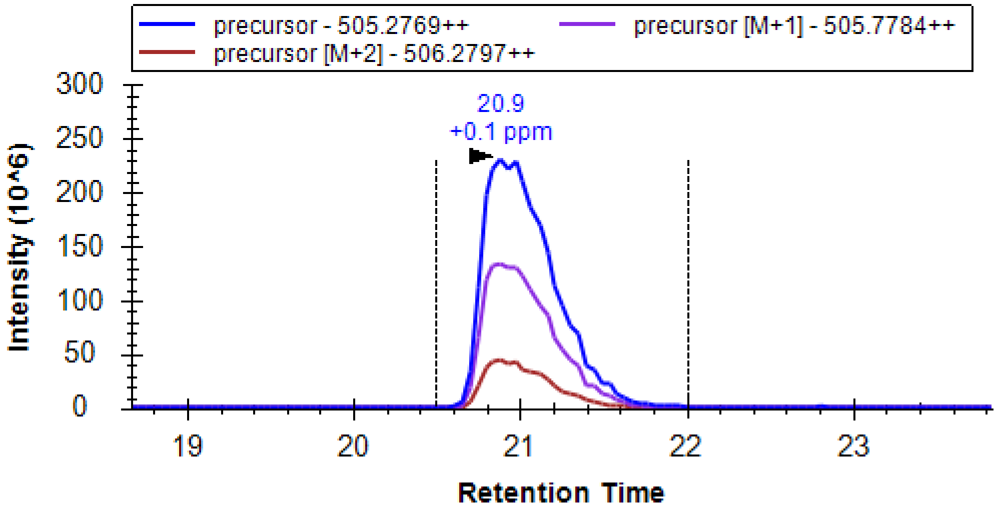


**b**, EIC of R10EPAVYFK.

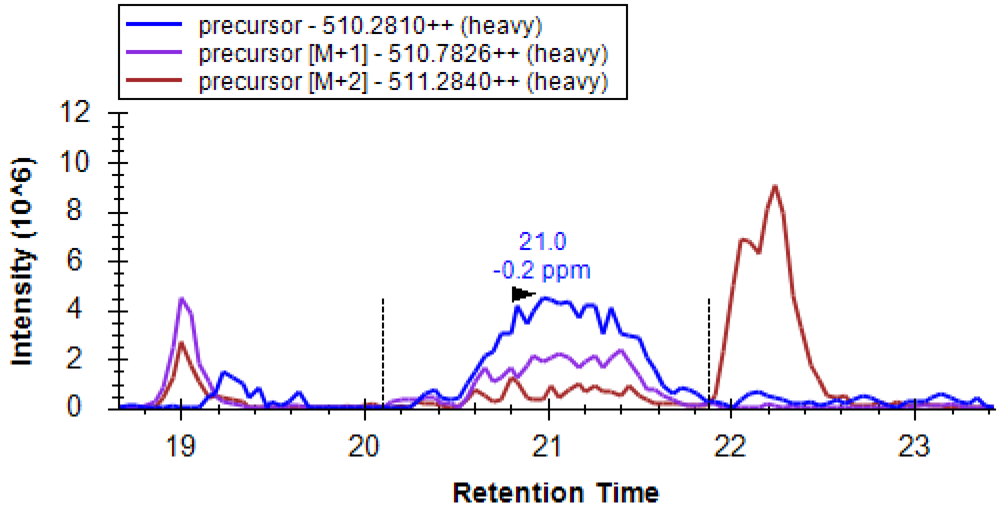


**c**, MS2 of REPAVYFK and added E(R)PAVYFK.

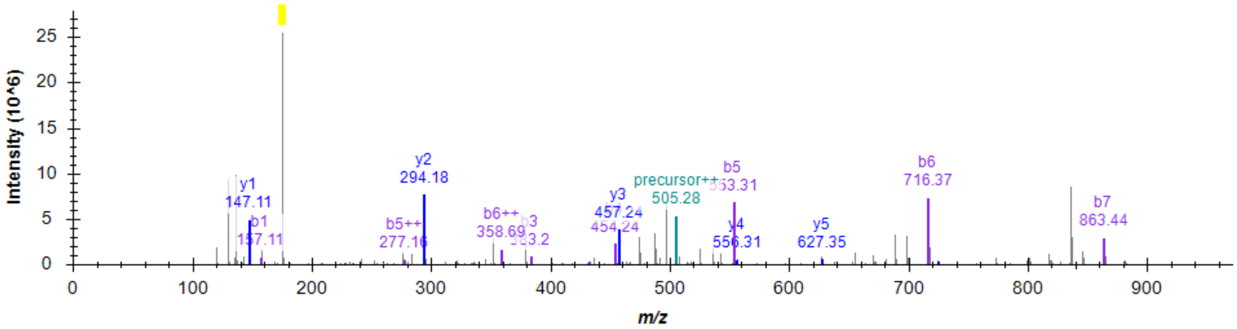

**d**, MS2 of R10EPAVYFK.

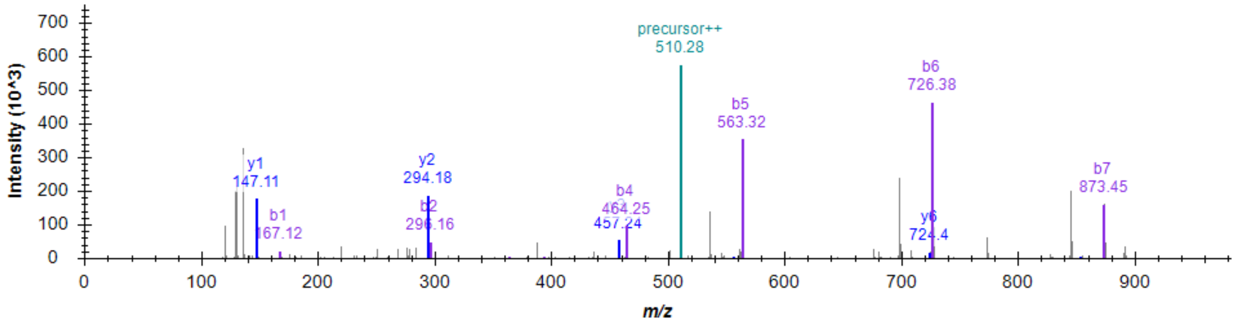


**Fig. S36**. Confirmation of CALR's arginylated peptide R[10]/R-EPAVYFK from arginylation profiling of HEK293T cells. E(R)PAVYFK was added to a peptide fraction (containing R[10]/R-EPAVYFK, H/L ratio = 0.9) of HEK293T cells. **a**. EIC of REPAVYFK and added E(R)PAVYFK. **b**. EIC of R[10]EPAVYFK. **c**. MS2 of REPAVYFK and added E(R)PAVYFK leading to the presence of highlighted signature ion at m/z 175.1190. **d**. MS2 of R10EPAVYFK.

**Related to Figure 4a.**

MS2 spectrum of peptide REEQPPETAAQR

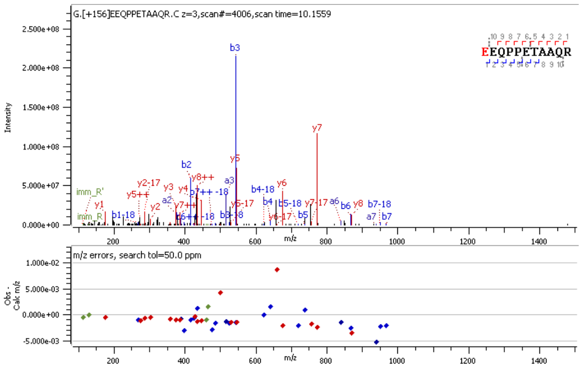


EICs of MS1 precursors and MS2 fragments of peptide LREEQPPETAAQR

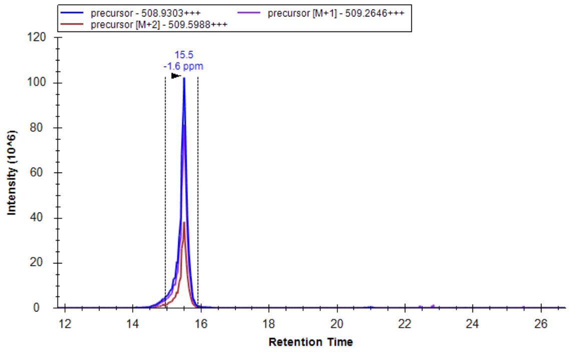


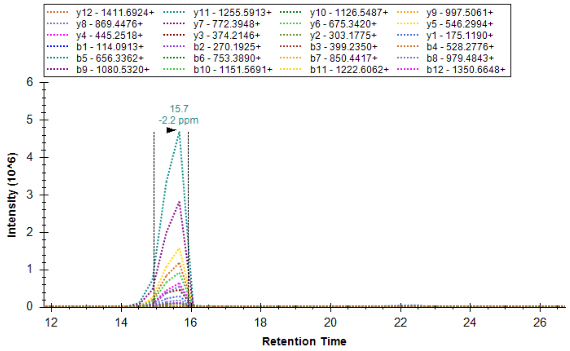


MS2 spectrum of peptide LREEQPPETAAQR

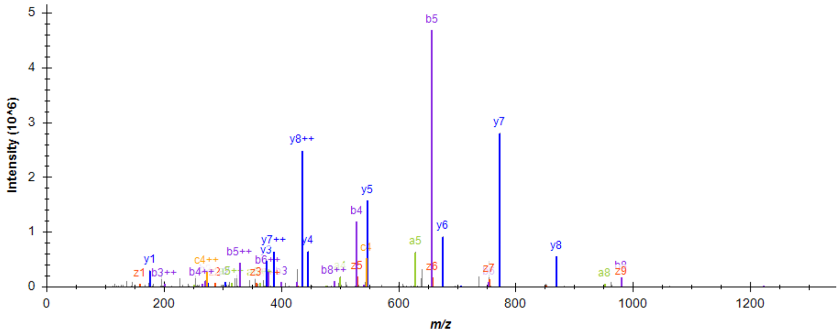


ERO1A sequence:
E24EQPPETAAQRCFCQVSGYLDDCTCDVETIDRFNNYRLFPRLQKLLESDYFRYYKVNLKRPCPFWNDIS
QCGRRDCAVKPCQSDEVPDGIKSASYKYSEEANNLIEECEQAERLGAVDESLSEETQKAVLQWTKHDDS

SDNFCEADDIQSPEAEYVDLLLNPERYTGYKGPDAWKIWNVIYEENCFKPQTIKRPLNPLASGQGTSEEN
TFYSWLEGLCVEKRAFYRLISGLHASINVHLSARYLLQETWLEKKWGHNITEFQQRFDGILTEGEGPRRLK
NLYFLYLIELRALSKVLPFFERPDFQLFTGNKIQDEENKMLLLEILHEIKSFPLHFDENSFFAGDKKEAHKLK
EDFRLHFRNISRIMDCVGCFKCRLWGKLQTQGLGTALKILFSEKLIANMPESGPSYEFHLTRQEIVSLFNAF
GRISTSVKELENFRNLLQNIHHHHHHHHH

**Fig. S37**. MS2 spectra of ERO1A E24 arginylation and further leucylation after ATE1-protein co-expression in *E. coli*. Protein N-term is an open E24 residue for arginylation. Arg0 modification, +156 Da. Leucylation after arginylation is visualized by Skyline.

**Related to Figure 4c.**

MS2 spectrum of peptide RESETTTSLVLER

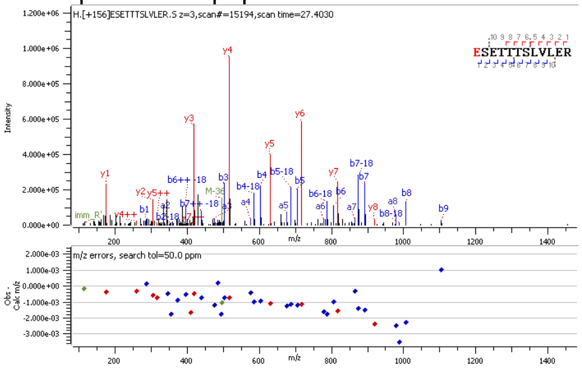


EICs of MS1 precursors and MS2 fragments of peptide LRESETTTSLVLER

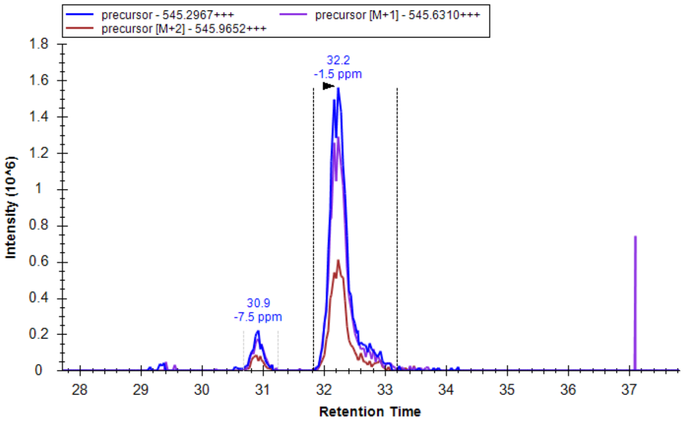


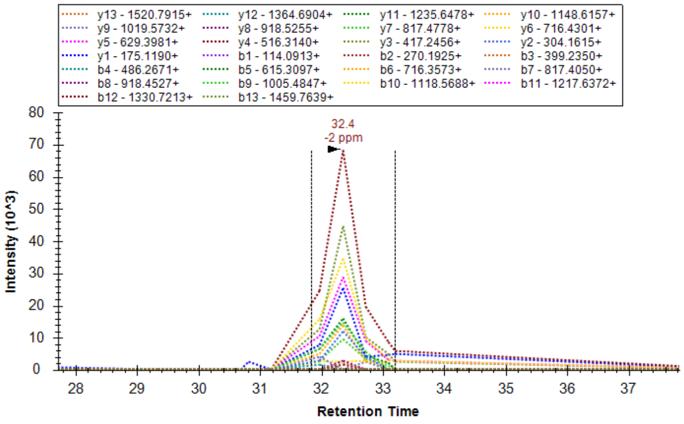


MS2 spectrum of peptide LRESETTTSLVLER

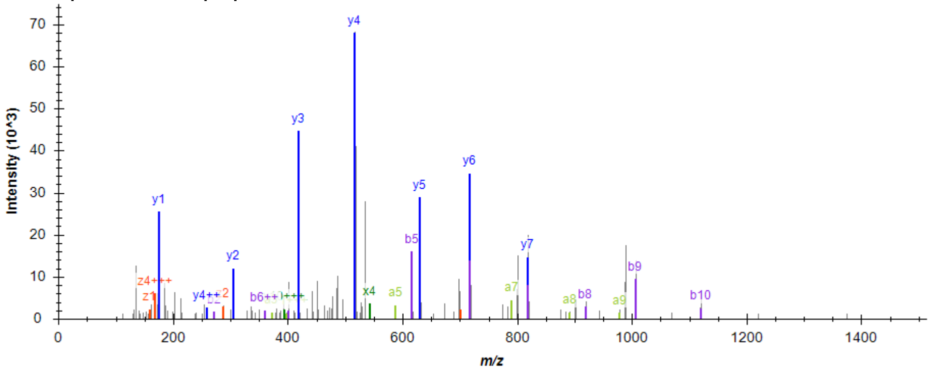

SSBP sequence:
E17SETTTSLVLERSLNRVHLLGRVGQDPVLRQVEGKNPVTIFSLATNEMWRSGDSEVYQLGDVSQKTTW
HRISVFRPGLRDVAYQYVKKGSRIYLEGKIDYGEYMDKNNVRRQATTIIADNIIFLSDQTKEKEHHHHHHHH

**Fig. S38**. MS2 spectra of SSBP E17 arginylation and further leucylation after ATE1-protein co-expression in *E. coli*. Protein N-term is an open E17 residue for arginylation. Arg0 modification, +156 Da. Leucylation after arginylation is visualized by Skyline.

**Related to Figure 4c.**

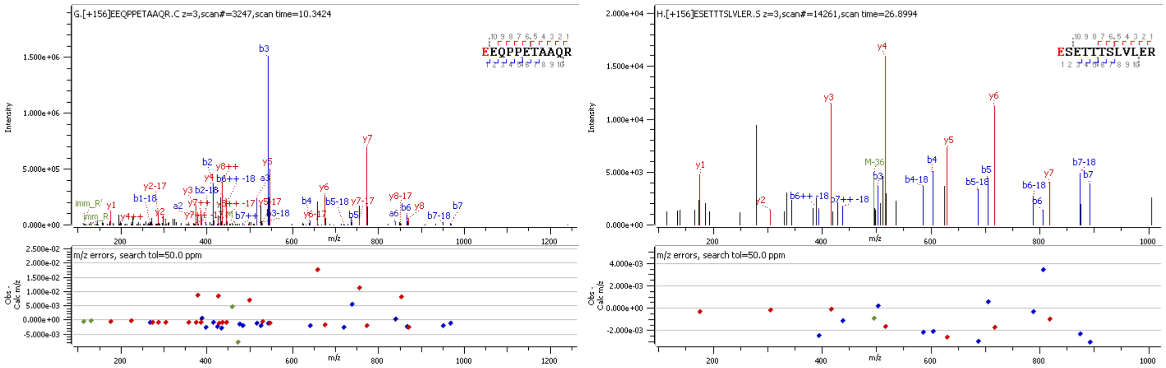


ERO1A peptide_ub sequence:
E$_{24}$EQPPETAAQRGSMQIFVKTLTGKTITLEVEPSDTIENVKAKIQDKEGIPPDQQRLIFAGKQLEDGRTLSDYNIQKESTLHLVLRLRGGHHHHHHHH

SSBP peptide_ub sequence:
E$_{17}$SETTTSLVLERSLNRGSMQIFVKTLTGKTITLEVEPSDTIENVKAKIQDKEGIPPDQQRLIFAGKQLEDGRTLSDYNIQKESTLHLVLRLRGGHHHHHHHH

**Fig. S39**. MS2 spectra of ERO1A E24 (left) and SSBP E17 (right) arginylation after ATE1-peptide co-expression in *E. coli*. Protein N-term is an open E residue for arginylation. Arg0 modification, +156 Da.

**Related to Figure 4c.**

### ERO1A Peptide + Ubiquitin + 8His

**a**

L-R- EEQPPETAAQRGSMQIFVKTLTGKTITLEVEPSDTIENVK
AKIQDKEGIPPDQQRLIFAGKQLEDGRTLSDYNIQKESTL
HLVLRLRGGHHHHHHHH

**b**

| | | Predicted | | Observed | |
|---|---|---|---|---|---|
| Protein | Modification | Monoisotopic Mass | Average Mass | Monoisotopic Mass | Average Mass |
| ERO1A pep Ubi 8his | WT | 11036.71 | 11043.25 | 11036.75 | 11043.61 |
| | R | 11192.82 | 11199.44 | 11192.84 | 11199.81 |
| | LR | 11305.90 | 11312.59 | 11305.93 | 11312.94 |

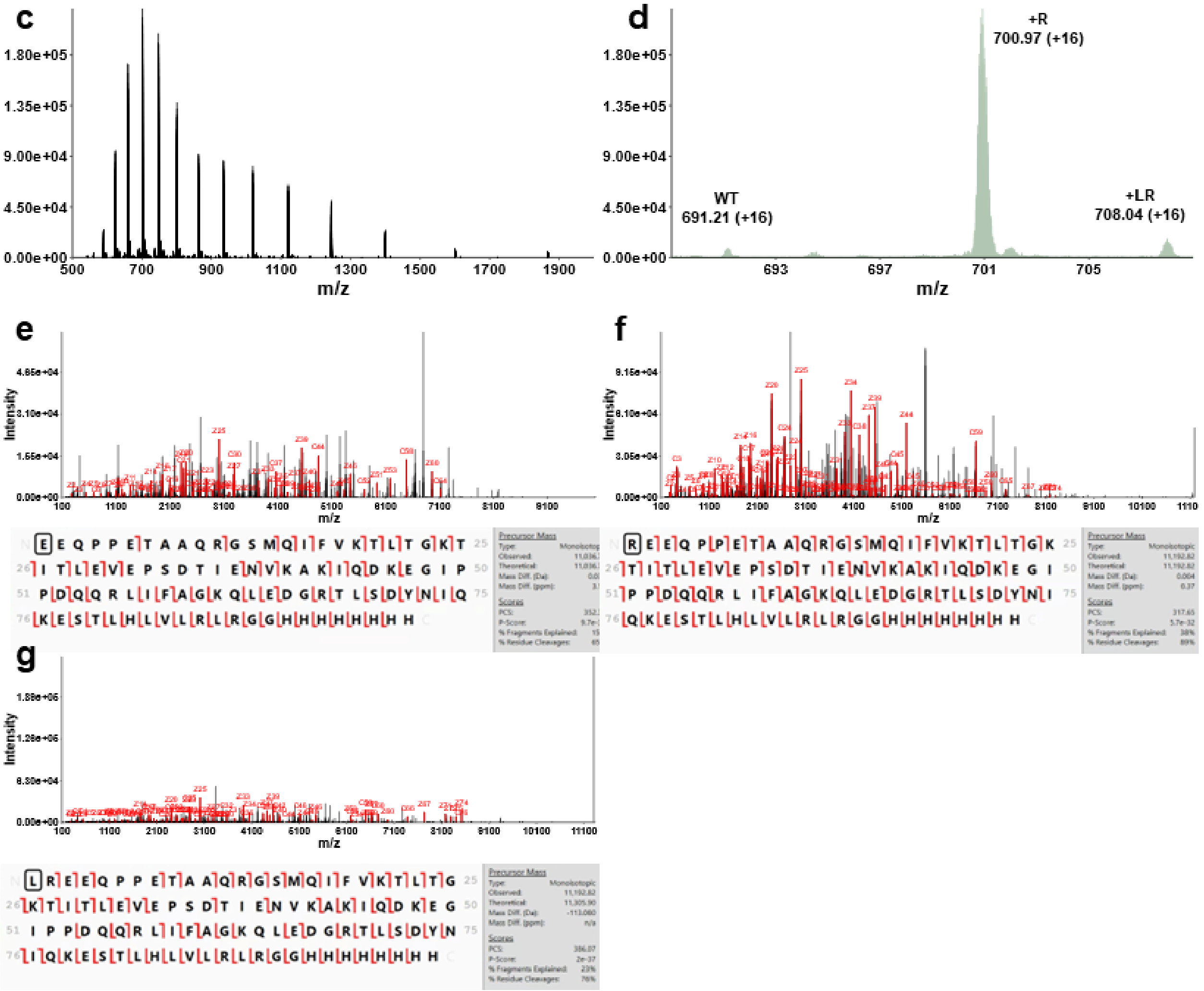


**Fig. S40**. Top-down characterization of ERO1A E24 arginylation using peptide_Ub fusion protein. **a**, Sequence of ERO1A peptide with C-terminal ubiquitin and 8his tag, and possible N-terminal arginine modification or leucine/arginine modification (Red). **b**, Predicted and observed monoisotopic and average masses of ERO1A species. **c**, Average MS1 spectra of coeluting ERO1A species. **d**, Zoomed spectra of ERO1A species showing +16 charge states. **e**, MS2 spectra and sequence coverage of WT ERO1A. **f**, MS2 spectra and sequence coverage of N-terminal arginylated ERO1A. **g**, MS2 spectra and sequence coverage of N-terminal leucylated and arginylated ERO1A.

**Related to Figure 4c.**

### SSBP1 Peptide + Ubiquitin + 8His

**a**

L-R-ESETTTSLVLERSLNRGSMQIFVKTLTGKTITLEVEPSDT
IENVKAKIQDKEGIPPDQQRLIFAGKQLEDGRTLSDYNIQ
KESTLHLVLRLRGGHHHHHHHH

**b**

| | | Predicted | | Observed | |
|---|---|---|---|---|---|
| Protein | Modification | Monoisotopic Mass | Average Mass | Monoisotopic Mass | Average Mass |
| SSBP1 pep Ubi 8his | WT | 11616.07 | 11622.94 | 11616.11 | 11623.31 |
| | R | 11772.17 | 11779.12 | 11772.22 | 11779.53 |
| | LR | 11885.26 | 11892.28 | 11885.30 | 11892.67 |

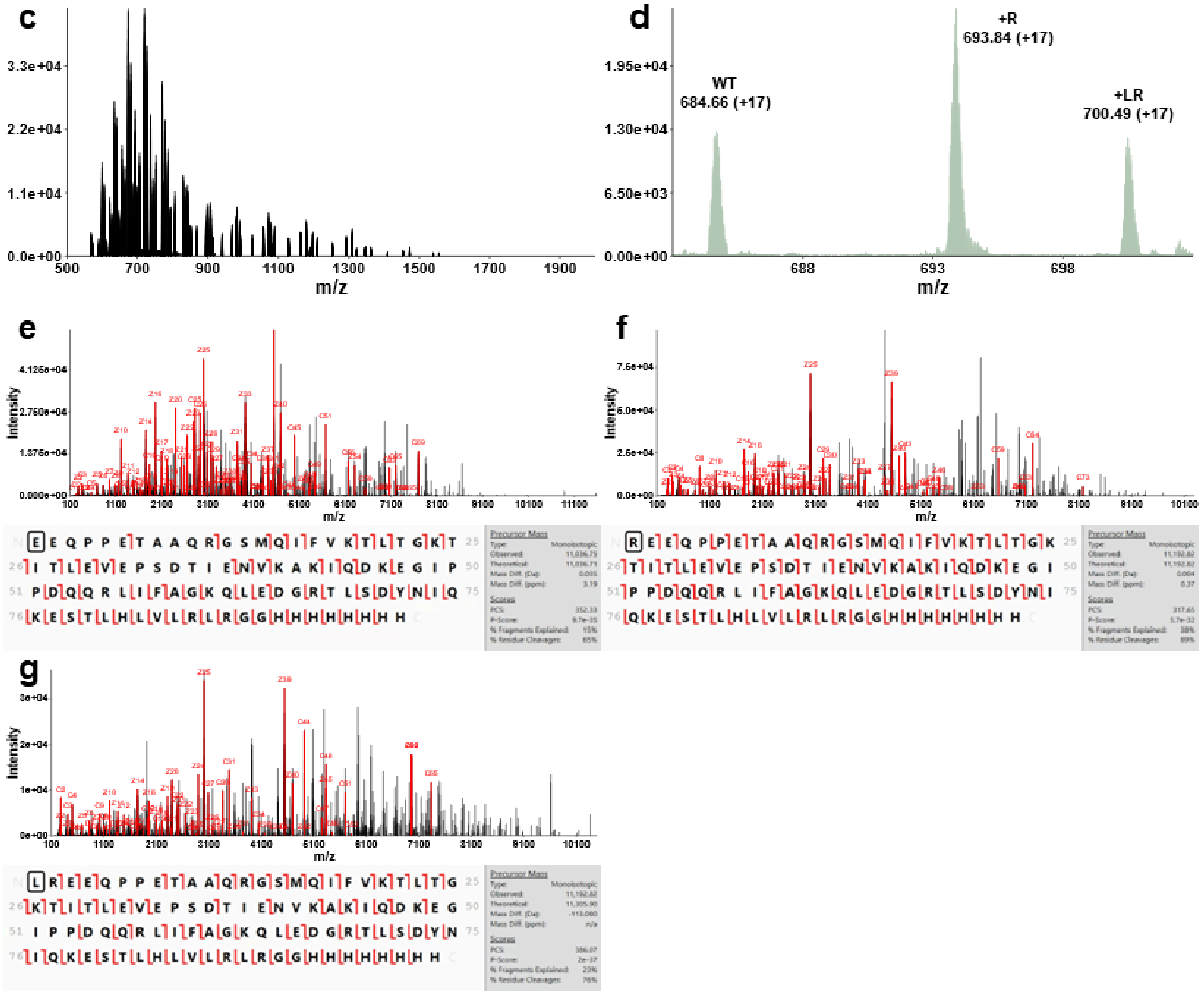


**Fig. S41**. Top-down characterization of SSBP E17 arginylation using peptide_Ub fusion protein. **a**, Sequence of SSBP1 peptide with C-terminal ubiquitin and 8his tag, and possible N-terminal arginine modification or leucine/arginine modification (Red). **b**, Predicted and observed monoisotopic and average masses of SSBP1 species. **c**, Average MS1 spectra of coeluting SSBP1 species. **d**, Zoomed spectra of SSBP1 species showing +17 charge states. **e**, MS2 spectra and sequence coverage of WT SSBP1. **f**, MS2 spectra and sequence coverage of N-terminal arginylated SSBP1. **g**, MS2 spectra and sequence coverage of N-terminal leucylated and arginylated SSBP1.

**Related to Figure 4c.**

**a**, CALR E18 R

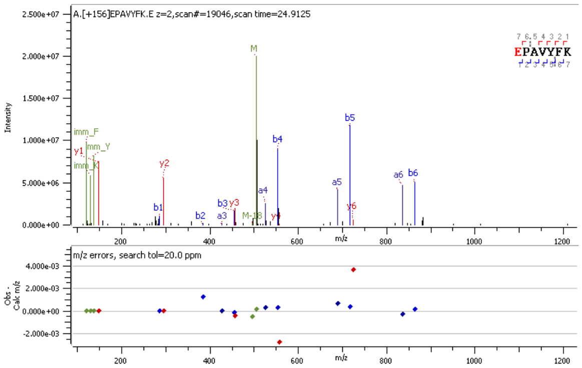


**b**, ERO1A E24 R

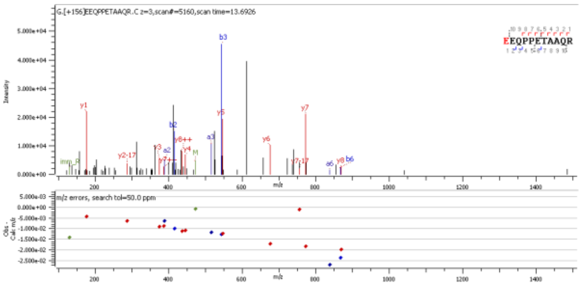


**c**, SSBP E17 R

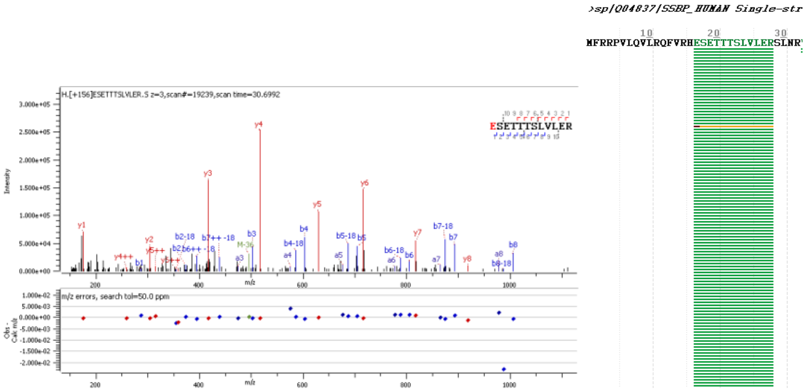


**Fig. S42**. MS2 spectra of arginylated peptides in CALR, ERO1A, and SSBP proteins purified from HEK293T cells after transfection. **a**, MS2 spectra of arginylated CALR E18 peptide. **b**, MS2 spectra of arginylated ERO1A E24 peptide. **c**, MS2 spectra of arginylated SSBP E17 peptide which was only detected once among unmodified. R, arginylation.

**Related to Figure 4d.**

**a**,

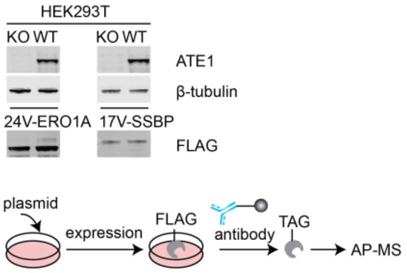


**b**, VEQPPETAAQR

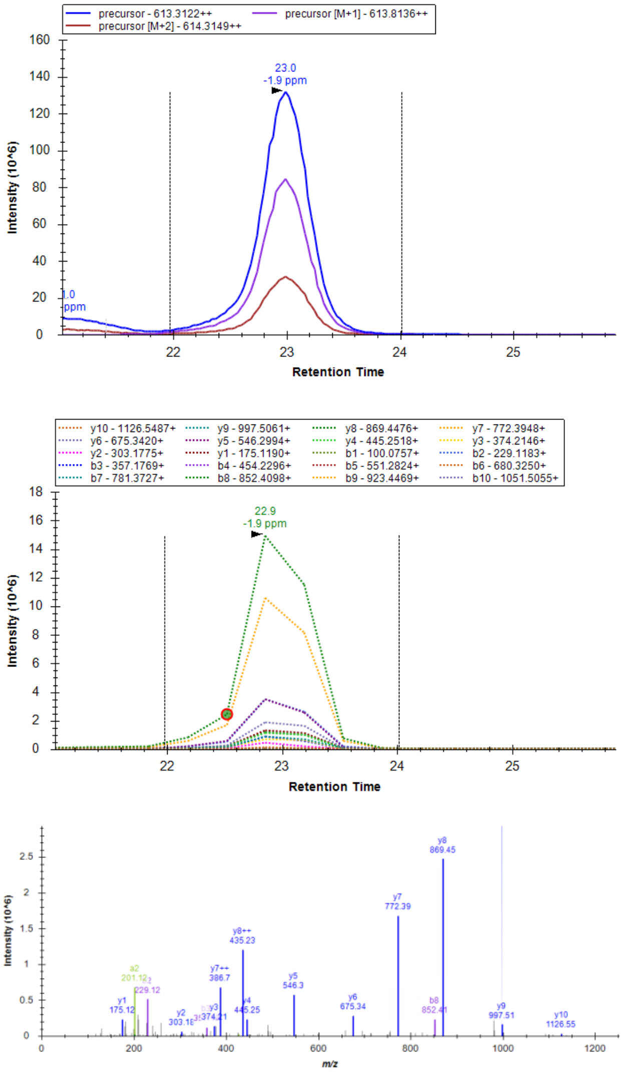


**c**, RVEQPPETAAQR (not detected in WT or ATE1 KO cells)

**d**, VSETTTSLVLER

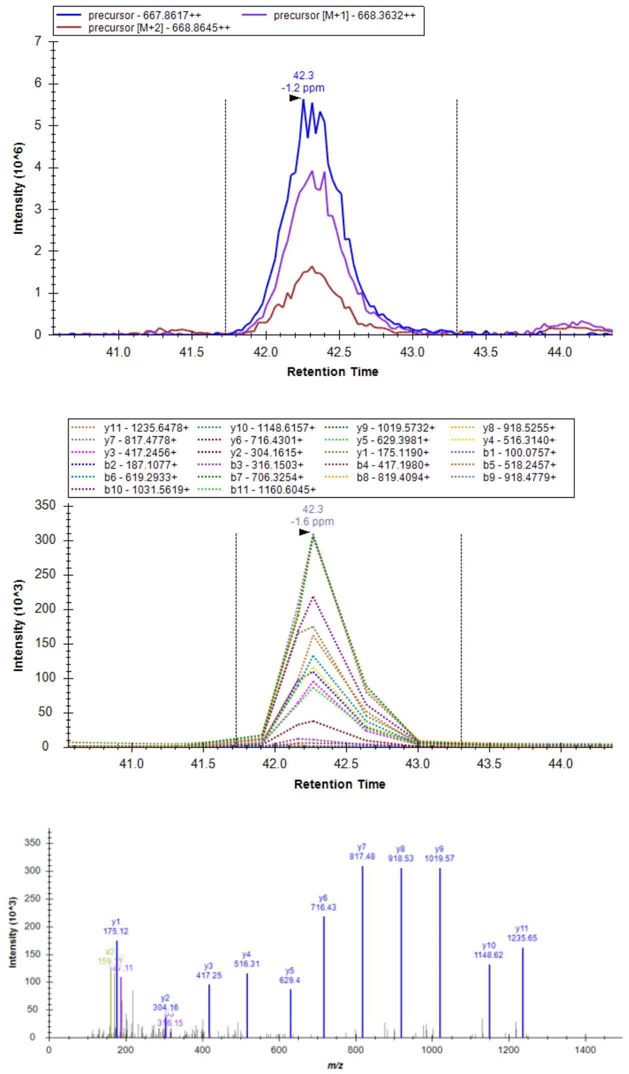


**e**, RVSETTTSLVLER (not detected in WT or ATE1 KO cells)

**Fig. S43**. Overexpression and IP pulldown of 24V-ERO1A and 17V-SSBP proteins from HEK293T cells for arginylation detection. **a**, overexpression of 24V-ERO1A and 17V-SSBP proteins. **b**, EIC of MS1 precursors and MS2 fragments of VEQPPETAAQR peptide. **c**, absence of RVEQPPETAAQR in WT or ATE1 KO cells. **d**, EIC of MS1 precursors and MS2 fragments of VSETTTSLVLER peptide. **c**, absence of RVSETTTSLVLER in WT or ATE1 KO cells.

**Related to Figure 4d.**

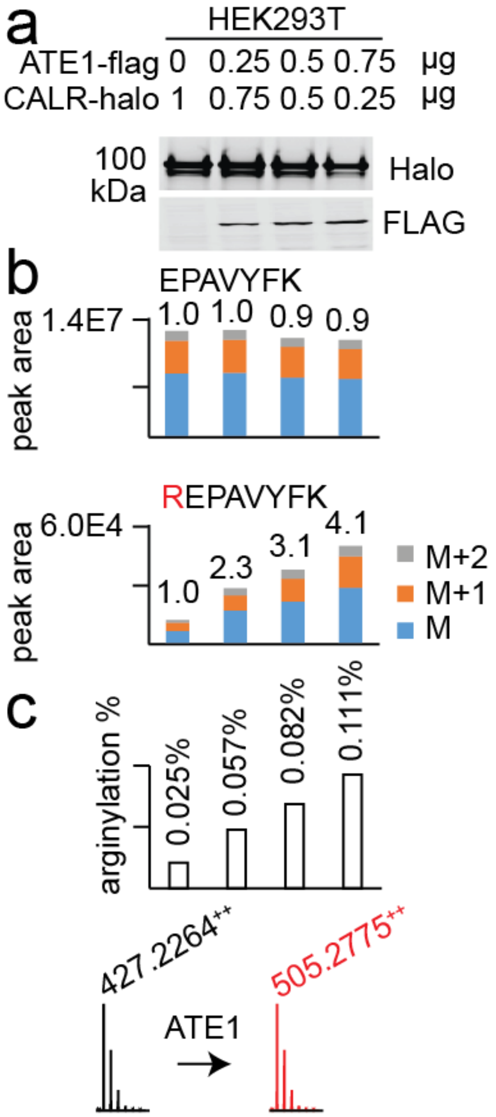


**Fig. S44**. Arginylation of CALR in WT HEK293T cells (*in vivo*) with ATE1 overexpression. **a**, overexpression of CALR-halo and ATE1-flag. **b**, quantification of unmodified and arginylated E18 peptide in overexpressed CALR using peak areas. Relative ratios are provided. **c**, quantification of CALR arginylation levels after normalization by CALR-halo expression levels. The ionization efficiency ratio of EPAVYFK and REPAVYFK at charge 2 was roughly determined to be 1:30.

**Related to Figure 4e.**

**a**, AA1-17_R18E_CALR

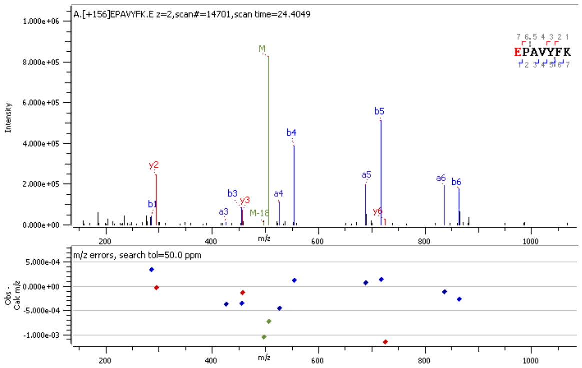


**b**, AA1-23_R24E_ERO1A

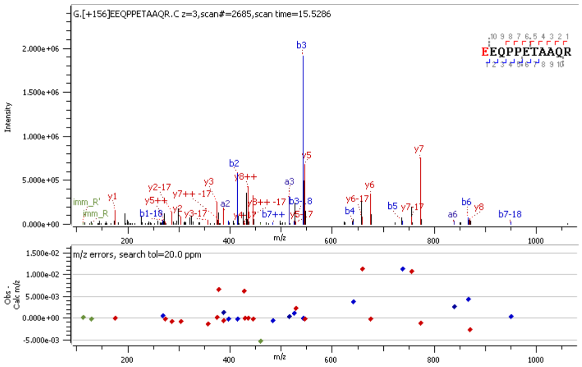


**c**, AA1-16_R17E_SSBP

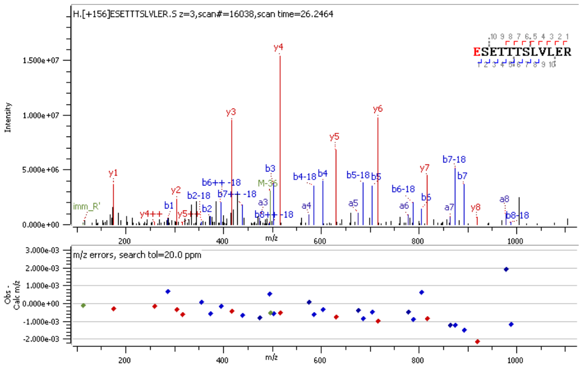


**Fig. S45**. MS2 spectra of arginylation sites after 18R-CALR, 24R-ERO1A, and 17R-SSBP overexpression and purification. Arginine residues were inserted to WT plasmids by mutagenesis. Arginylation peptides were used as *in vivo* or transfection positive controls for arginylation confirmation of respective proteins.

**Related to Figure 4e.**

**a**,

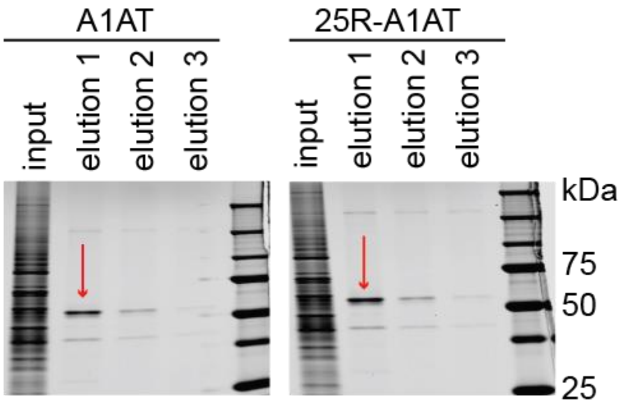


**b**,

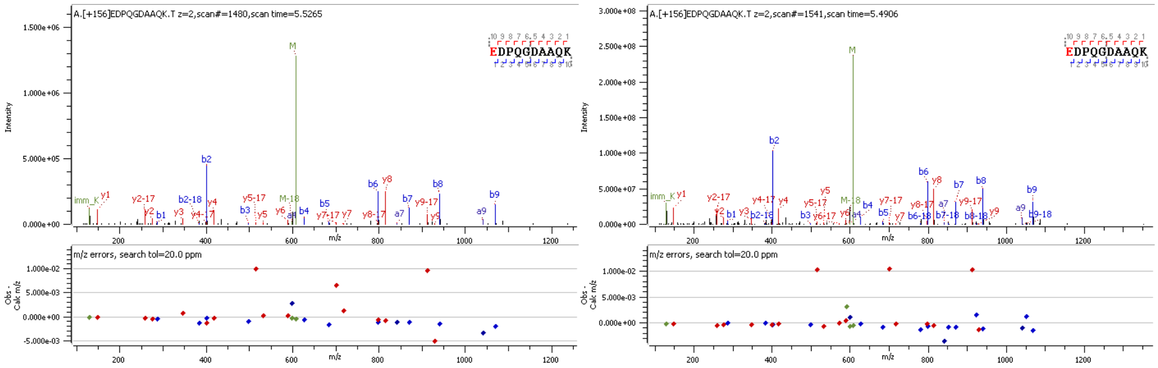


A1AT 25R-A1AT

**Fig. S46**. MS2 spectra of arginylation sites after A1AT and 25R-A1AT overexpression and purification. **a**, A1AT and 25R-A1AT overexpression and purification. **b**, detection of E24 arginylation (left) in A1AT and E24 arginylation peptide (right) in 25R-A1AT as a positive control.

**Related to Figure 4.**

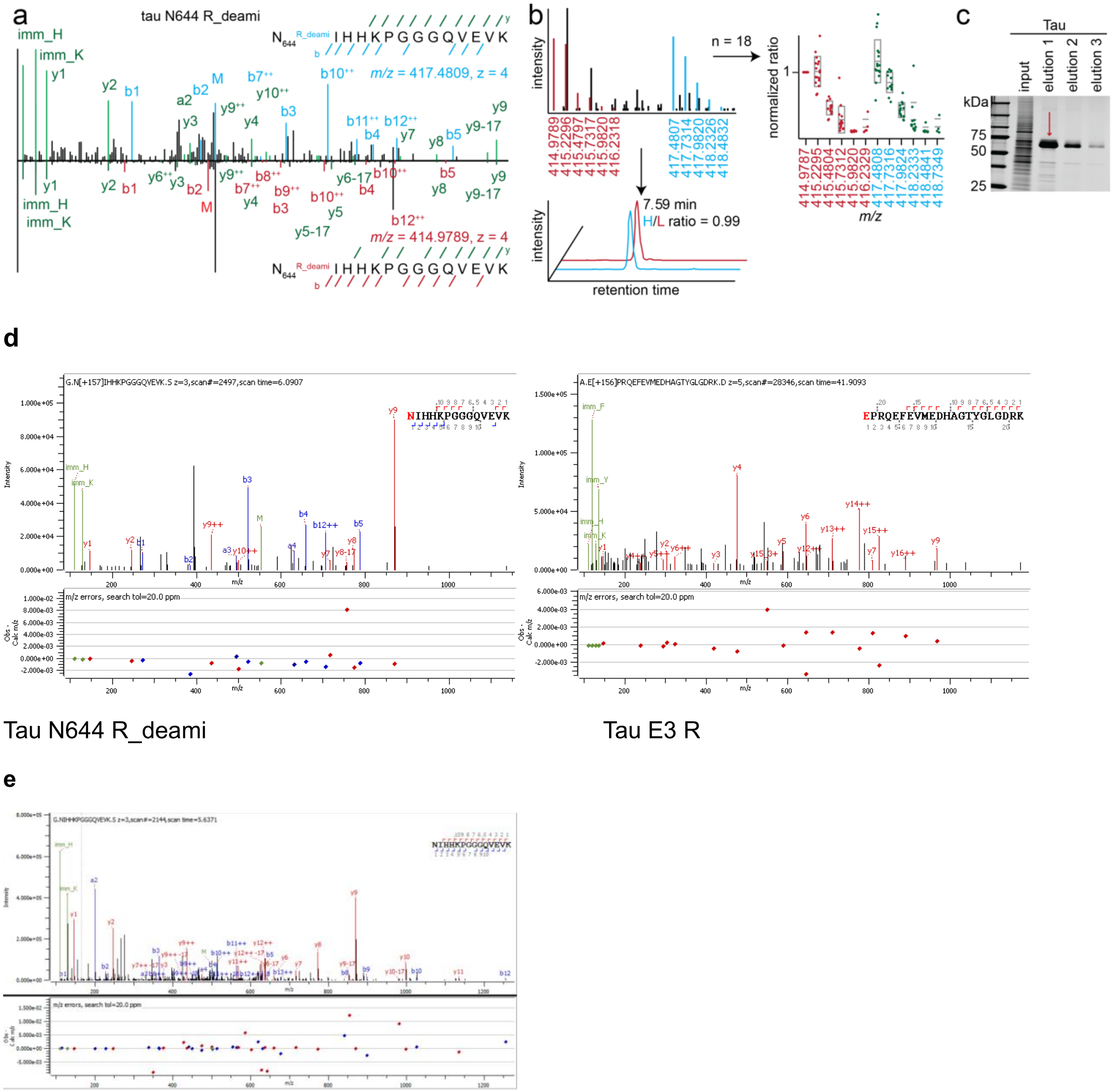


**Fig. S47**. Tau arginylation and validation. **a**, representative MS2 spectra indicating tau arginylation *ex vivo* on N644 arginylation after deamidation (tau N644 R_deami). **b,** statistics of MS1 pairs of tau N644 R_deami in one fraction of an AD sample. **c**, tau overexpression and purification. **d**, detection of N644 (left) and E3 (right) arginylation sites. **e**, detection of unmodified N644 peptide. R, arginylation. R_deami, arginylation after deamidation.

**Related to Figure 4.**

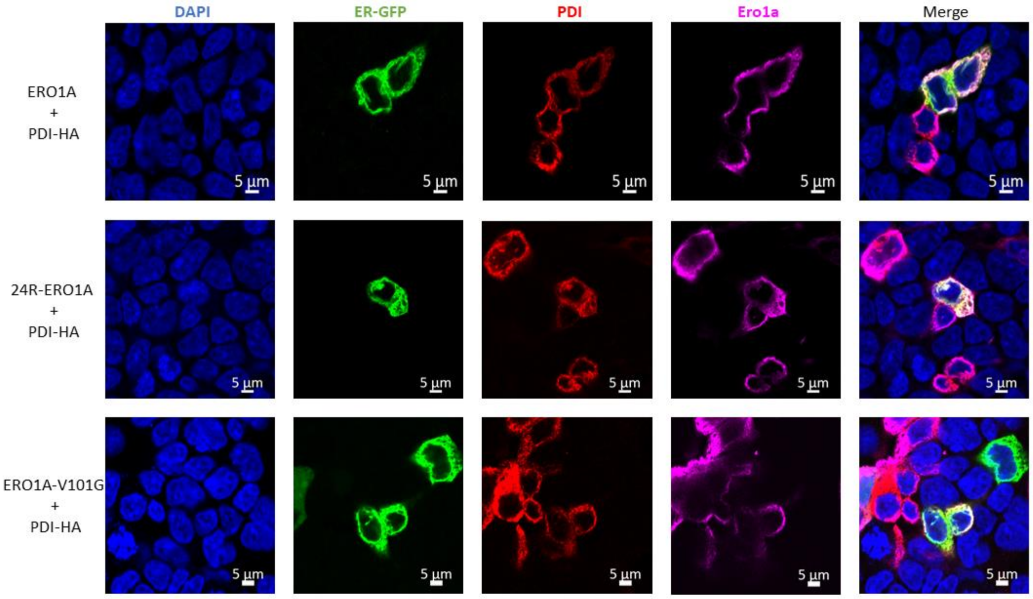


**Fig. S48.** Cell imaging of ERO1A and 24R-ERO1A compared with ER marker and PDI. ERO1A and arginylated ERO1A superimpose well with ER maker and PDI.

**Related to Figure 5a.**

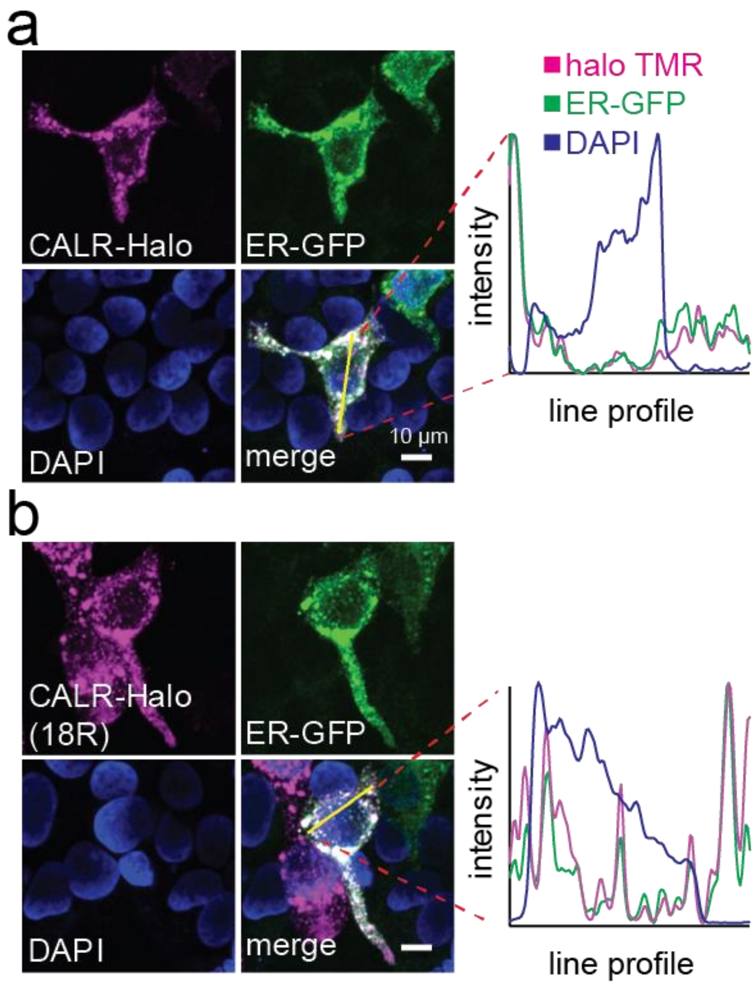


**Fig. S49.** Cell imaging of CALR-halo and 18R-CALR-halo compared with a GFP-labeled ER marker. CALR (**a**) and arginylated CALR (**b**) superimpose well with ER maker.

**Related to Figure 5a.**

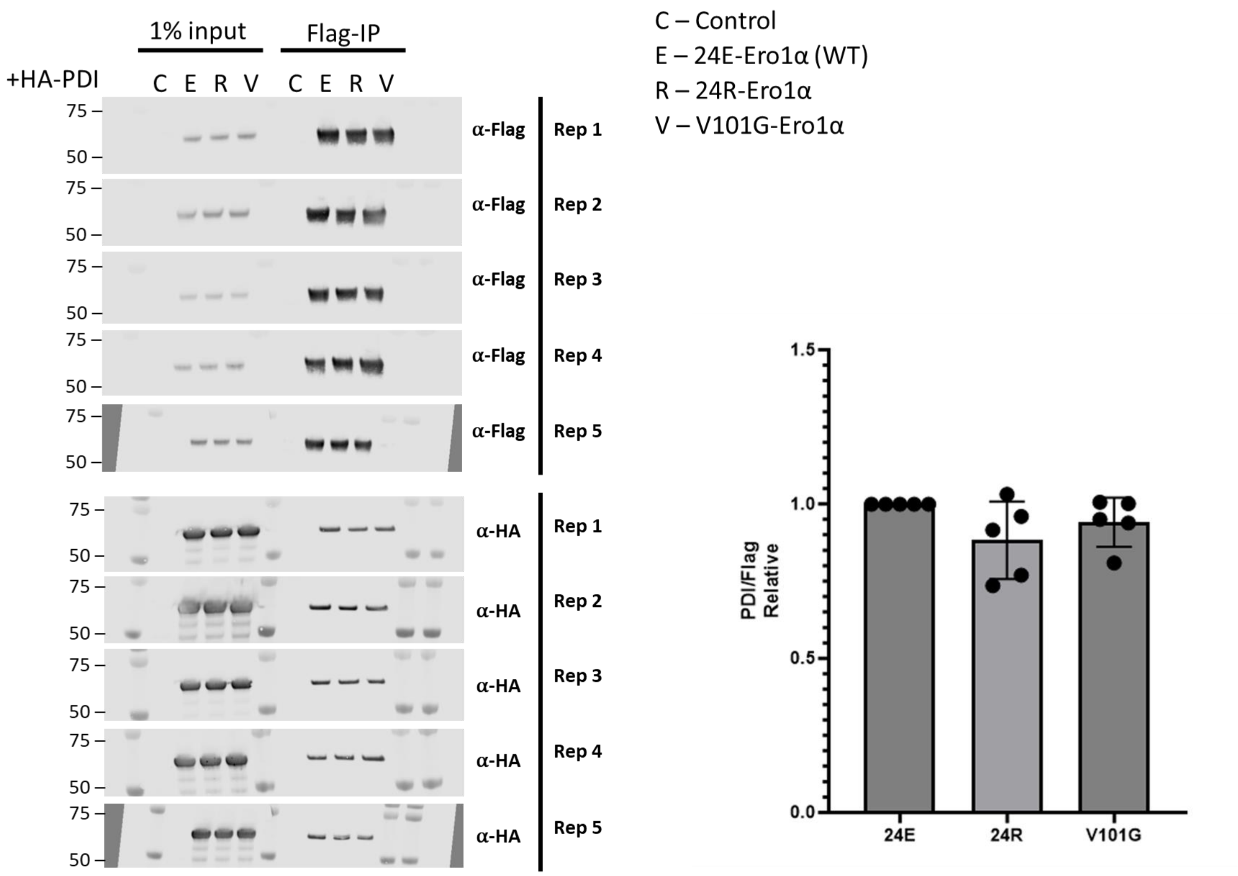


**Fig. S50**. Quantification of PDI pulled down by co-expressed ERO1A (bait) mutants. Five replicates were used (*n* = 5).

**Related to Figure 5b.**

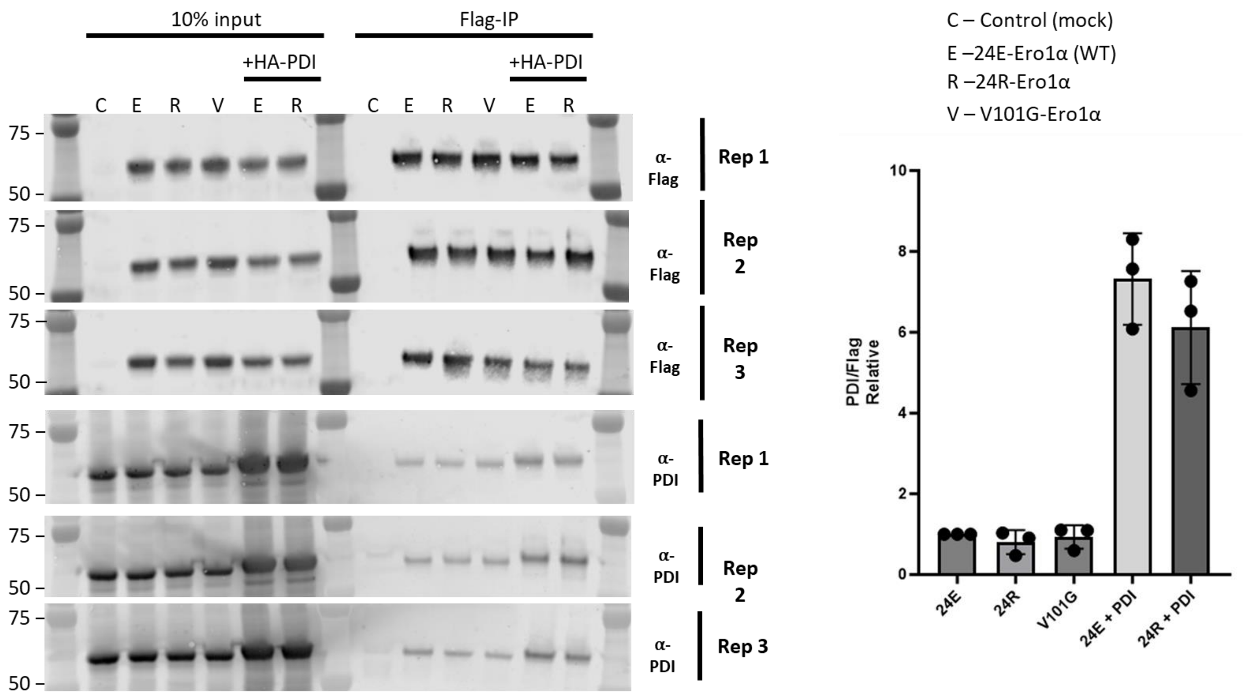


**Fig. S51**. Quantification of endogenous and overexpressed PDI pulled down by overexpressed ERO1A (bait) mutants. Three replicates were used (*n* = 3).

**Related to Figure 5b.**

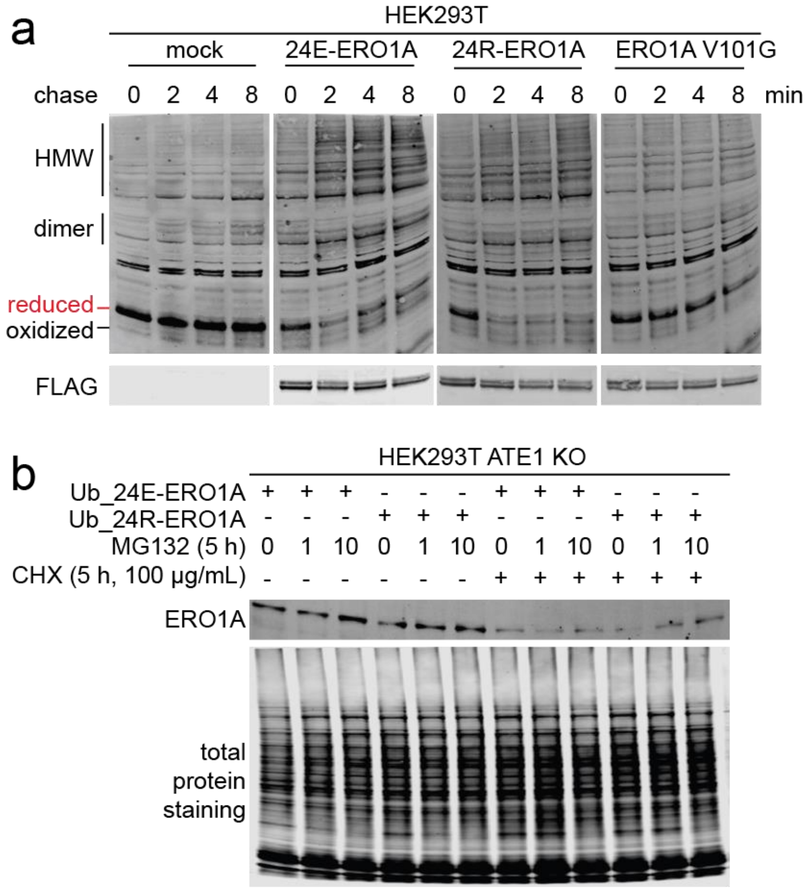


**Fig. S52.** Enzymatic and degradation activities of ERO1A. **a**, oxidative folding of mouse JcM. HEK293T cells were co-transfected with JcM and ERO1A mutants with mock as a control. Cells were pulsed by DTT and chased at various times after DTT removal. The disappearance of fully reduced JcM represents the ERO1A oxidative folding activity. **b**, expression levels of cytosolic ERO1A species after MG132 and CHX treatments. For MG132 treatments, DMSO was used as control. MG132 and CHX were added to cells at the same time.

**Related to Figure 5b.**

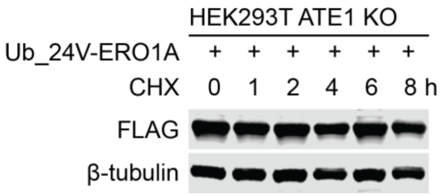


**Fig. S53.** Expression levels of cytosolic 24V-ERO1A species after CHX treatments. CHX was added to cells at different times to monitor protein stability. Cells were harvested at the same time to compare protein levels.

**Related to Figure 5b.**

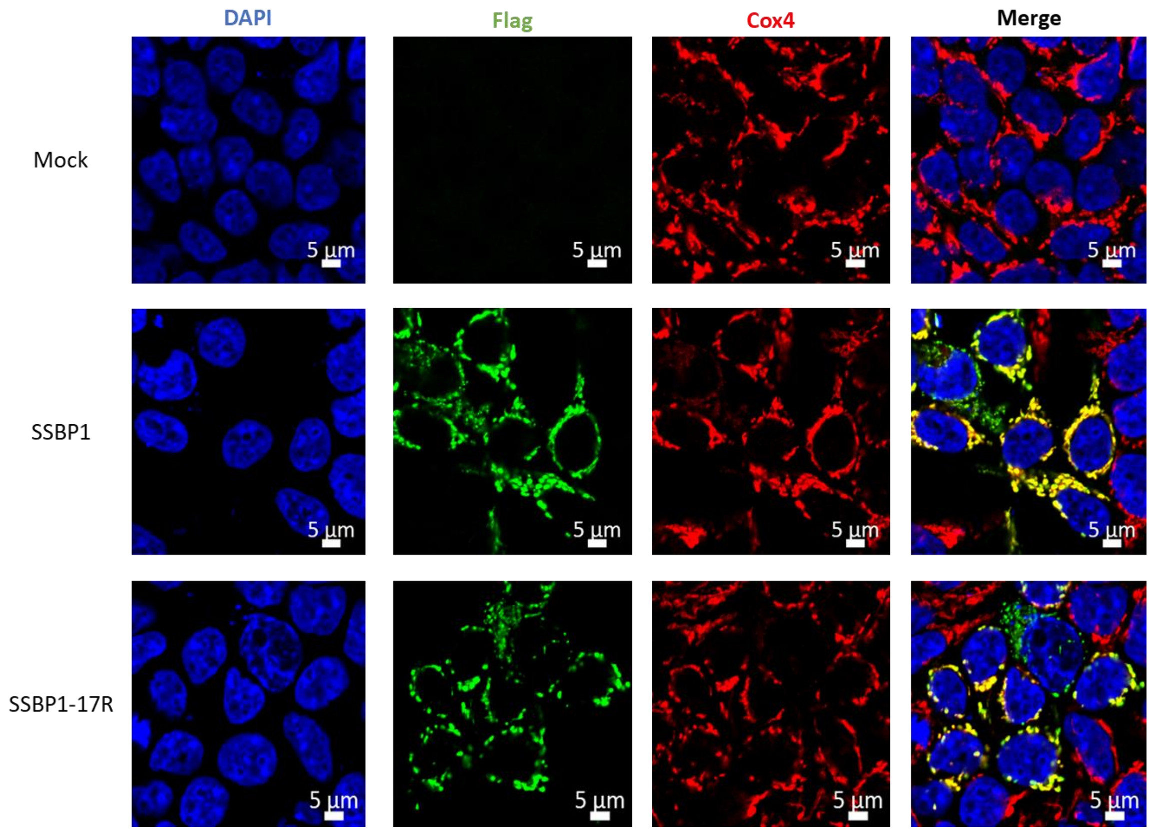


**Fig. S54.** Cell imaging of SSBP and 17R-SSBP compared with mitochondria protein COX4. SSBP and arginylated SSBP superimpose with COX4 as shown in merged figures.

**Related to Figure 5c.**

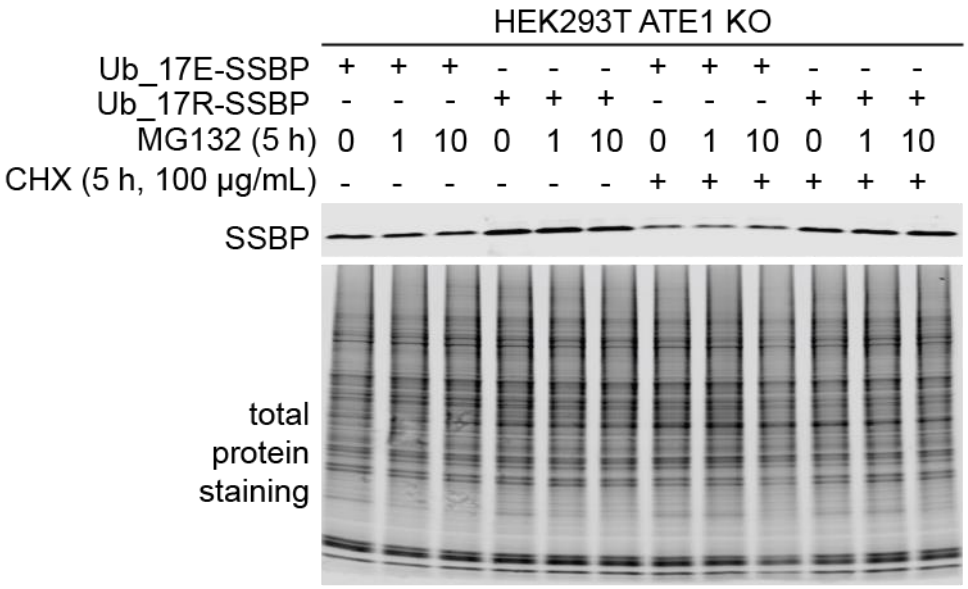


**Fig. S55**. Expression levels of cytosolic SSBP species after MG132 and CHX treatments. For MG132 treatments, DMSO was used as control. MG132 and CHX were added to cells at the same time.

**Related to Figure 5c.**

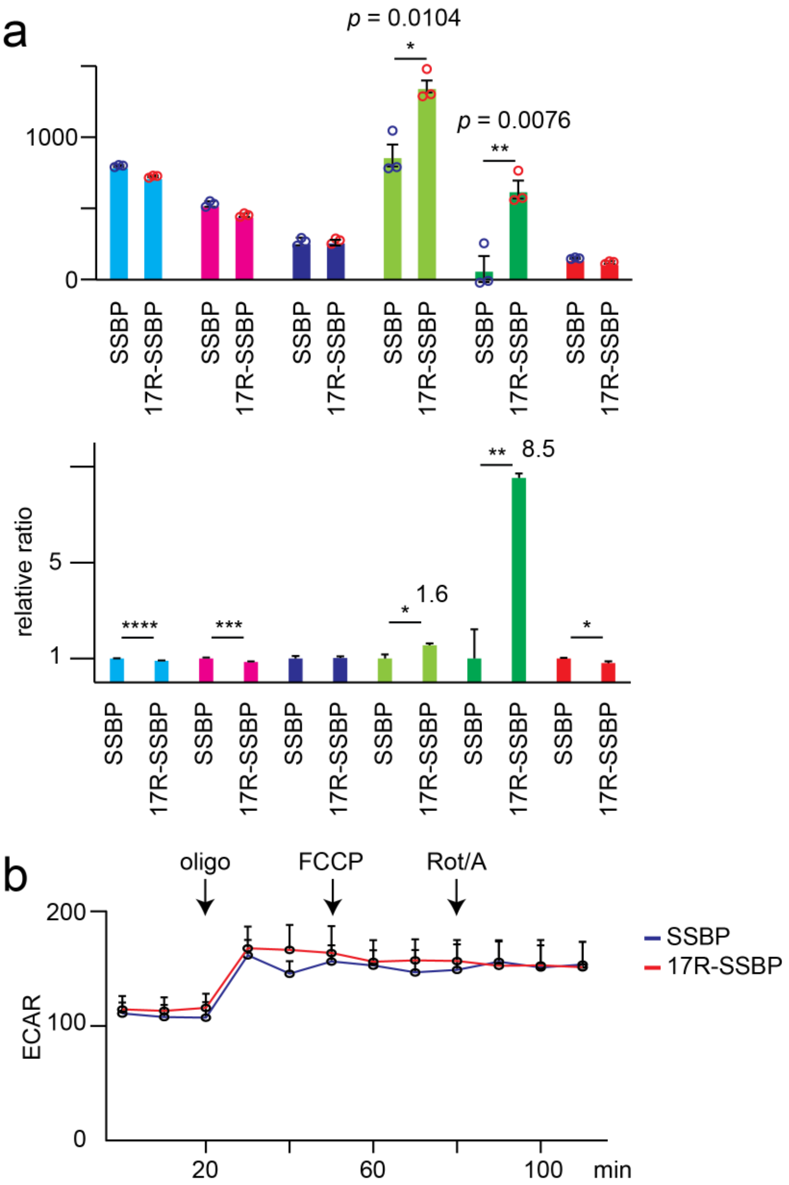


**Fig. S56.** Data summary of Seahorse XF24 Cell Mito Test assay. **a**, Relative ratios of OCR parameters are calculated for cells overexpressing SSBP and 17R-SSBP. The average values of 3 Seahorse measurements were used for comparison. A two-tail Student's *t*-test was used. $*p < 0.05$, $**p < 0.01$, $***p < 0.005$, $***p < 0.001$. **b**, ECAR profiles of SSBP and 17R-SSBP. 5 biological replicates ($n$ = 5 wells) were used for each measurement.

**Related to Figure 5d.**

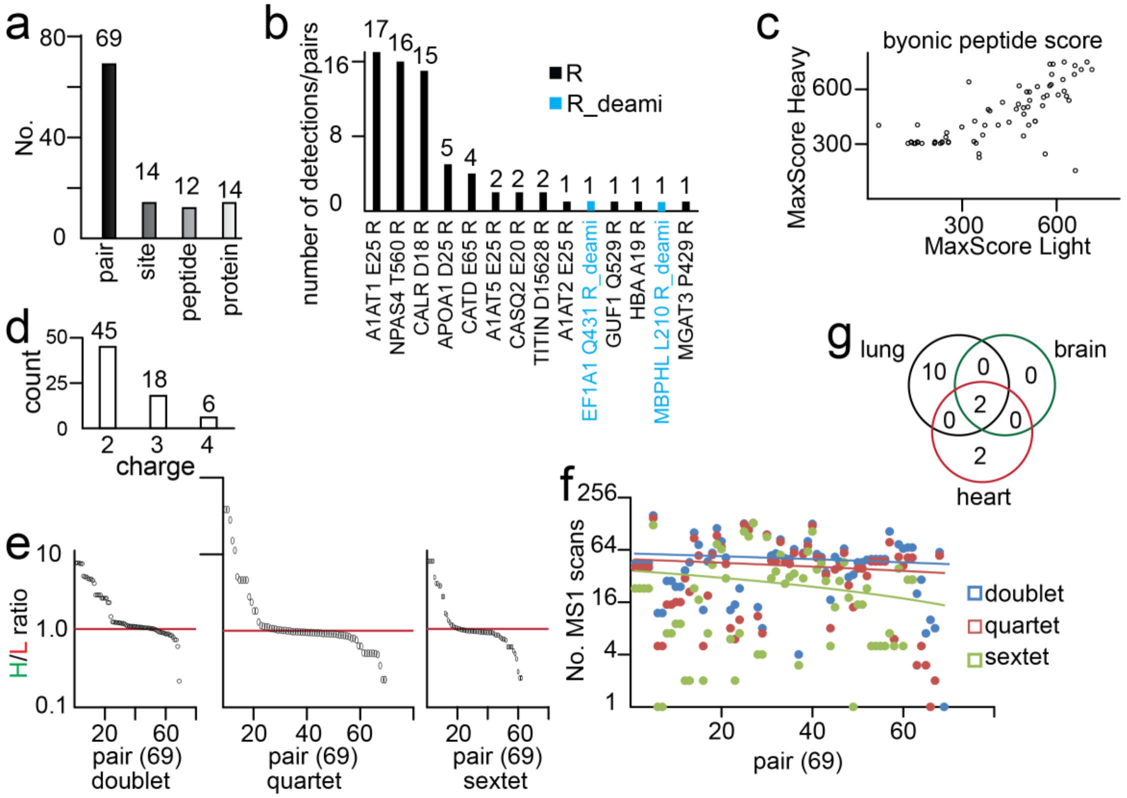


**Figure S57.** Summary of arginylation sites in mouse tissues. **a,** numbers of identified MS1 pairs, arginylation sites, unique peptides and unique proteins in mouse tissues. b, number of detections/pairs for each arginylation site from mouse tissues. **c**, peptide score distribution for all detections/pairs from mouse tissues. Peptides with both H and L scores < 300 are excluded. **d**, peptide charge distribution for all detections/pairs from mouse tissues. **e**, H/L ratio distribution of all MS1 pairs in doublet, quartet, and sextet from mouse tissues. **f**, numbers of MS1 scans of all MS1 pairs in doublet, quartet, and sextet from mouse tissues. Trendlines for all doublets, quartets, and sextets are generated respectively. **g**, Venn diagram on numbers of shared and unique arginylation sites in mouse tissues.

**Related to Supplementary Dataset 7.**

a,

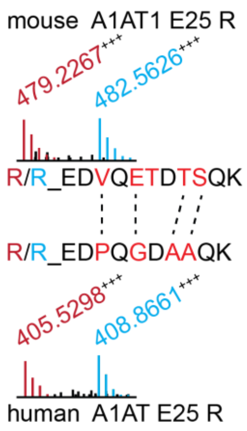


b,

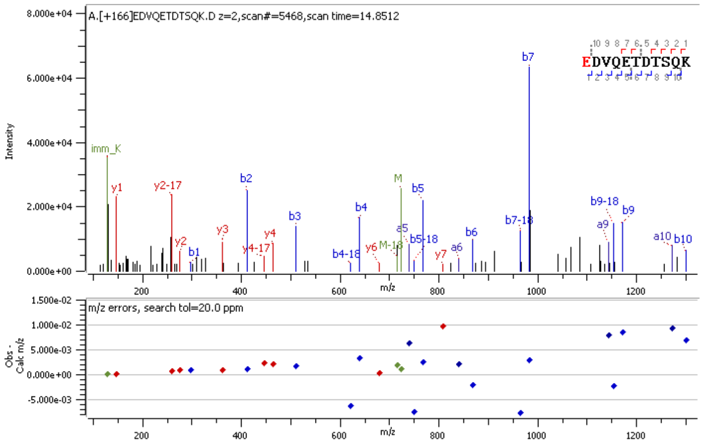


c,

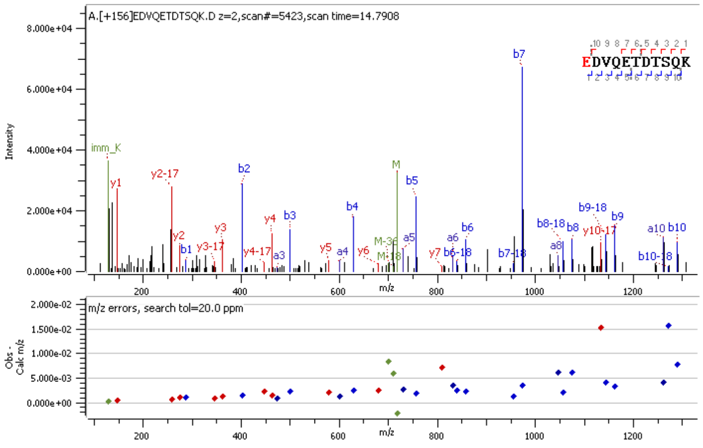


**Fig. S58.** Mouse A1AT1 E25 arginylation. **a**, sequence comparison of mouse and human A1AT arginylation from proteomics profiling. **b**, MS2 spectrum of A1AT arginylation peptide with Arg10 modification. **c**, MS2 spectrum of A1AT arginylation peptide with Arg0 modification.

**Related to Supplementary Dataset 7.**

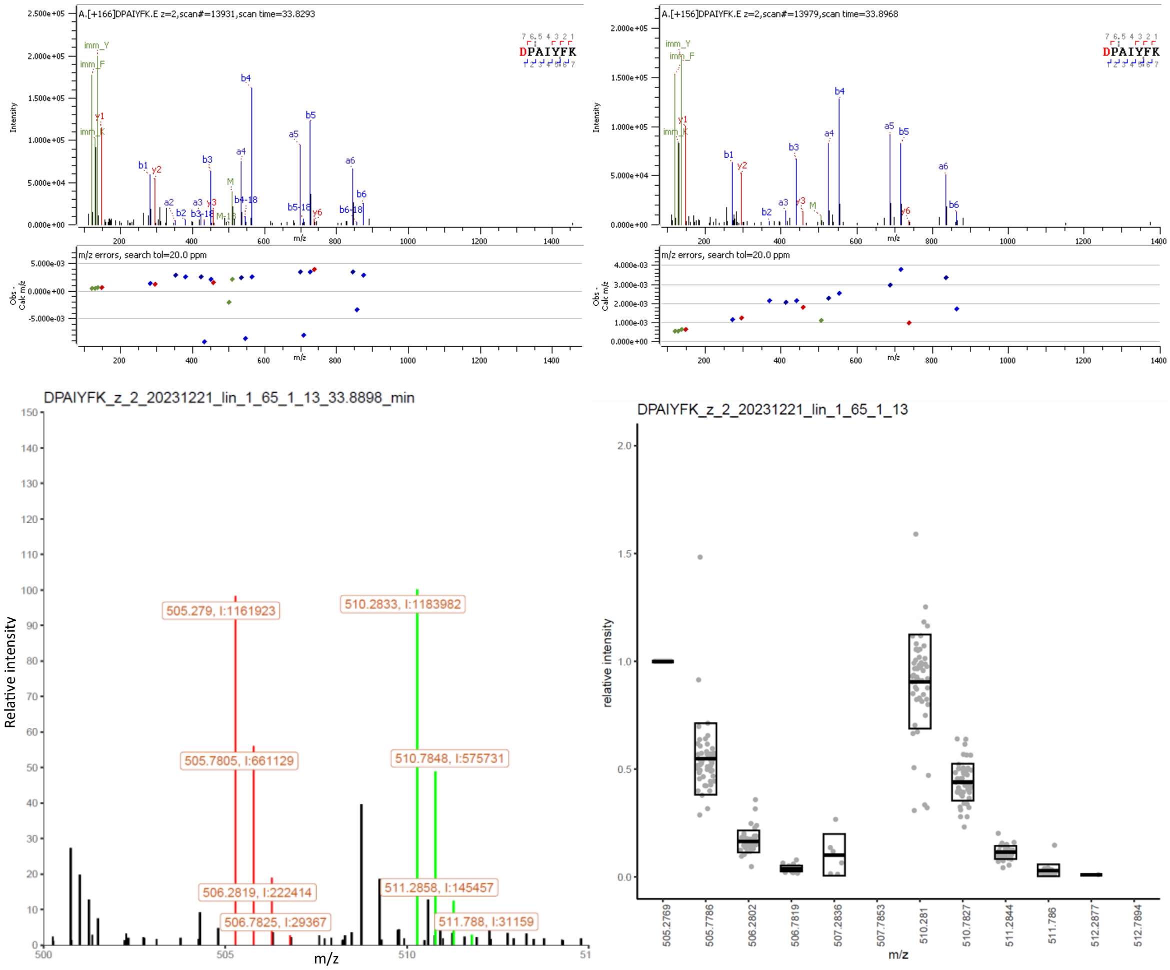


**Fig. S59**. MS2 identification and MS1 ratio quantification of mouse CALR D18 arginylation. DPAIYFK was arginylated by Arg10 (+166 Da) and Arg0 (+156 Da) with a mix ratio of 1:1. *m/z* values and intensities are displayed in the MS1 spectrum. A box plot is summarized from 57 MS1 scans.

**Related to Supplementary Dataset 7.**

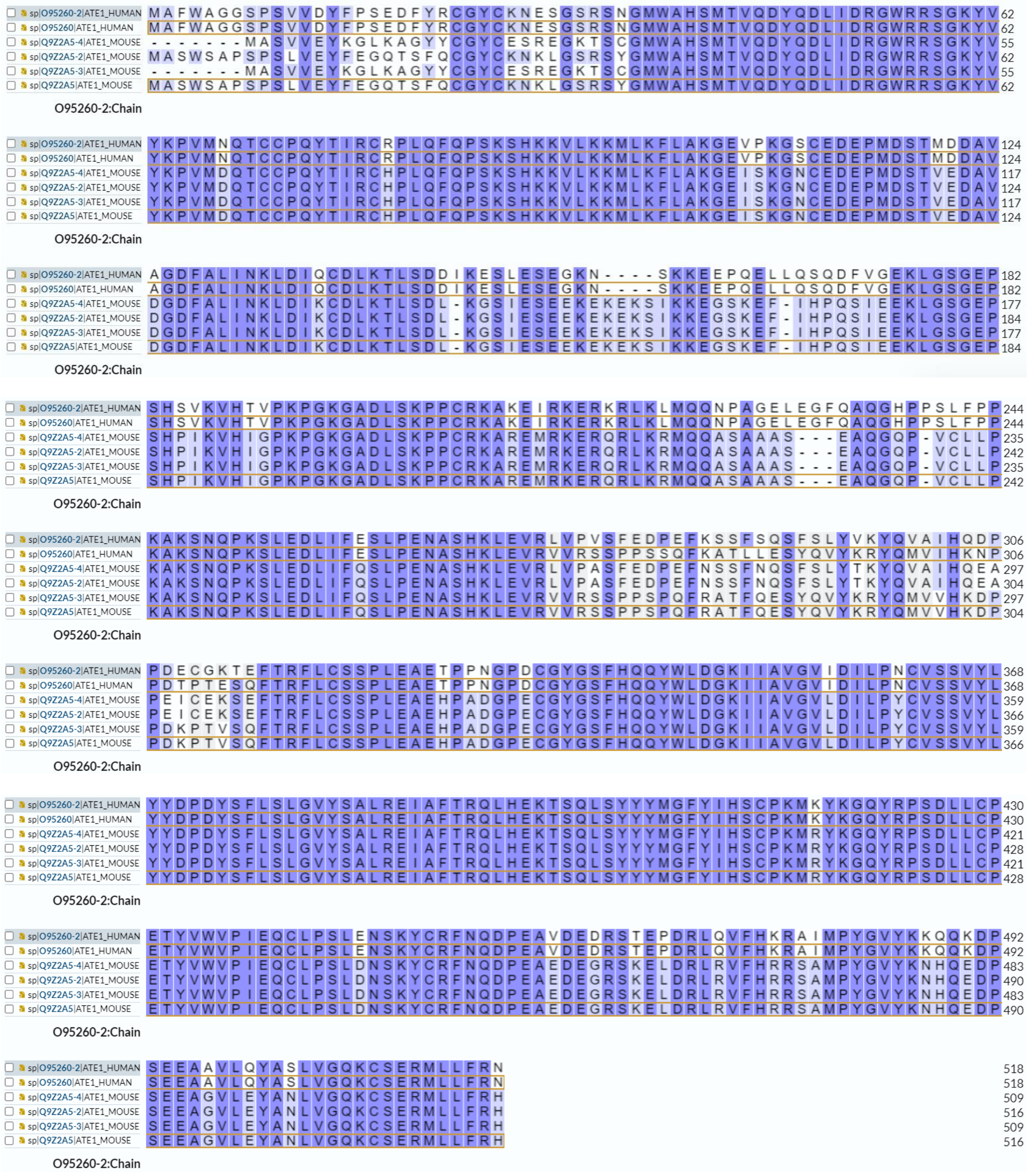


**Fig. S60**. Sequence alignment of human and mouse ATE1 isoforms. Sequences of 2 human isoforms and 4 mouse isoforms are aligned by multiple sequence alignment using CLUSTAL O(1.2.4).

**Related to Discussion.**

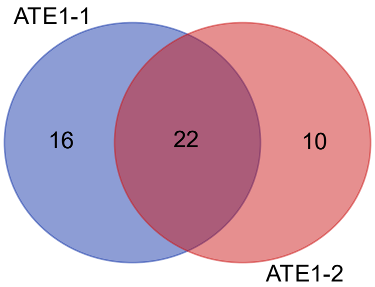


Summary table of arginylation sites in brain_AD sample.

| sample | number of elements | number of unique elements |
|---|---|---|
| ATE1-1 | 38 | 38 |
| ATE1-2 | 32 | 32 |
| Overall number of unique elements | | 48 |

Detailed table of sample overlaps.

| sample | total | elements |
|---|---|---|
| ATE1-1 ATE1-2 | 22 | NFL D468 R EFHD2 Q67 R_deami NFL Q75 R_deami GFAP E391 R CATD G65 R GFAP D417 R MBP D168 R NFM E599 R NFL E469 R MBP G277 R TCAF1 V114 R NFL E453 R NEUM E89 R AKA12 E952 R MBP N226 R_deami PLEC E4050 R MBP E217 R MTAP2 Q431 R_deami MBP D216 R A1AT E25 R NFM E454 R CALR E18 R |
| ATE1-1 | 16 | NFM E497 R NFL E507 R MAP1A E663 R NCAN E23 R NHRF1 E121 R TBA1B D327 R SAP G407 R AAK1 Q374 R_deami NFM E884 R TAU N644 R_deami SAP D405 R SPTN1 E1124 R NFH E469 R PDIA1 D18 R NFM E455 R NFM E496 R |
| ATE1-2 | 10 | MBP T200 R NCAM1 E833 R CATD E64 R F162A C41 RO3 AKA12 E418 R AP180 N767 R_deami E2F7 D10 R DZI1L Q132 R CC148 I574 R MBP N226 R |

**Fig. S61**. Venn diagram of arginylation sites in brain_AD sample using ATE1-1 and ATE1-2. Summary and detailed overlapping sites are listed.

**Related to Discussion.**

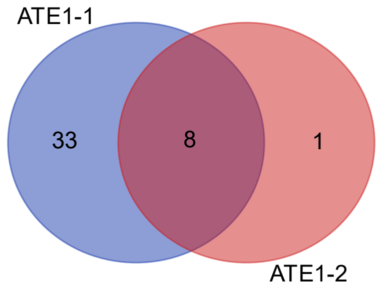


Summary table of arginylation sites in HeLa sample.

| **sample** | **number of elements** | **number of unique elements** |
|---|---|---|
| ATE1-1 | 41 | 41 |
| ATE1-2 | 9 | 9 |
| Overall number of unique elements | | 42 |

Detailed table of sample overlaps.

| **sample** | **total** | **elements** |
|---|---|---|
| ATE1-1 ATE1-2 | 8 | G3P D315 R HSP7C G372 R HNRPK N306 R ACTB E241 R G3P D296 R PDIA1 D18 R CALR E18 R RM12 C45 RO3 |
| ATE1-1 | 33 | TXND5 E40 R COX41 D43 R SSBP E17 R NUCL E628 R HNRPK Q358 R_deami SNX17 E463 R ILF2 D347 R RL23A E 8 R HNRPK D279 R BI1 I 8R_deami CNBP Q44 R_deami ACTB D51 R CATD E64 R ERO1A E24 R MCFD2 E27 R LMAN2 D45 R VIME E425 R FUBP1 N369 R_deami RON D1232 R RS7 E188 R HNRPC D123 R PDIA4 E25 R CALX E275 R MYH9 D1136 R SAP D405 R VATD E242 R ACLY E830 R IF4H D196 R ROA1 E93 R RLA1 D19 R VIME D451 R CALU D62 R HNRPL D284 R |
| ATE1-2 | 1 | FCN1 D120 R |

**Fig. S62**. Venn diagram of arginylation sites in HeLa sample using ATE1-1 and ATE1-2. Summary and detailed overlapping sites are listed.

**Related to Discussion.**

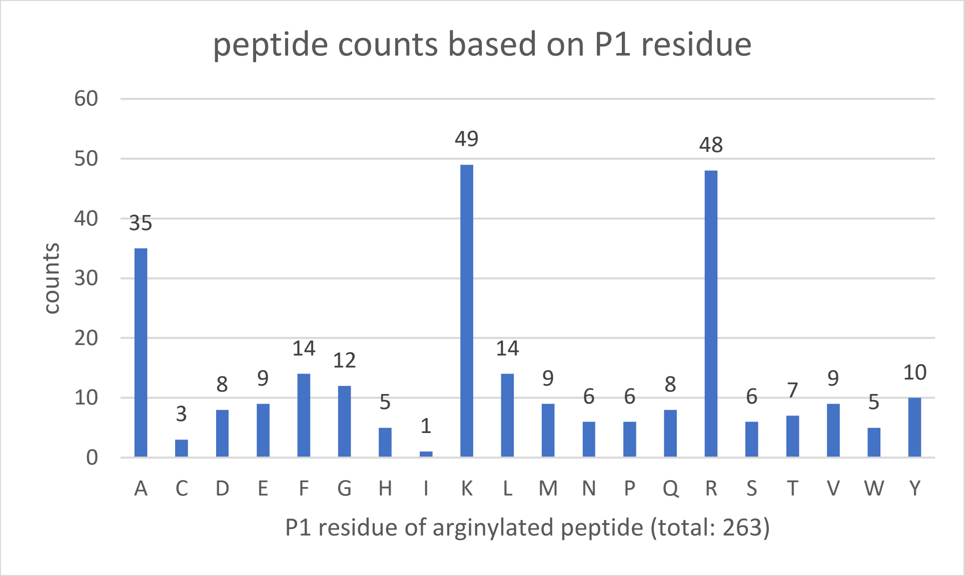
peptide counts based on P1 residue
counts
P1 residue of arginylated peptide (total: 263)
A C D E F G H I K L M N P Q R S T V W Y
35 3 8 9 14 12 5 1 49 14 9 6 6 8 48 6 7 9 5 10
0 10 20 30 40 50 60

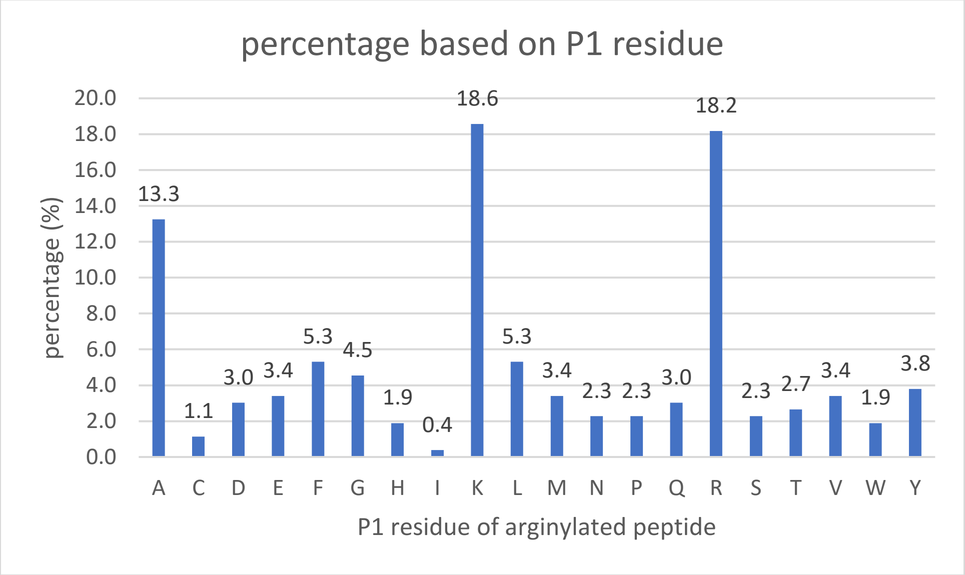
percentage based on P1 residue
percentage (%)
P1 residue of arginylated peptide
A C D E F G H I K L M N P Q R S T V W Y
13.3 1.1 3.0 3.4 5.3 4.5 1.9 0.4 18.6 5.3 3.4 2.3 2.3 3.0 18.2 2.3 2.7 3.4 1.9 3.8
0.0 2.0 4.0 6.0 8.0 10.0 12.0 14.0 16.0 18.0 20.0

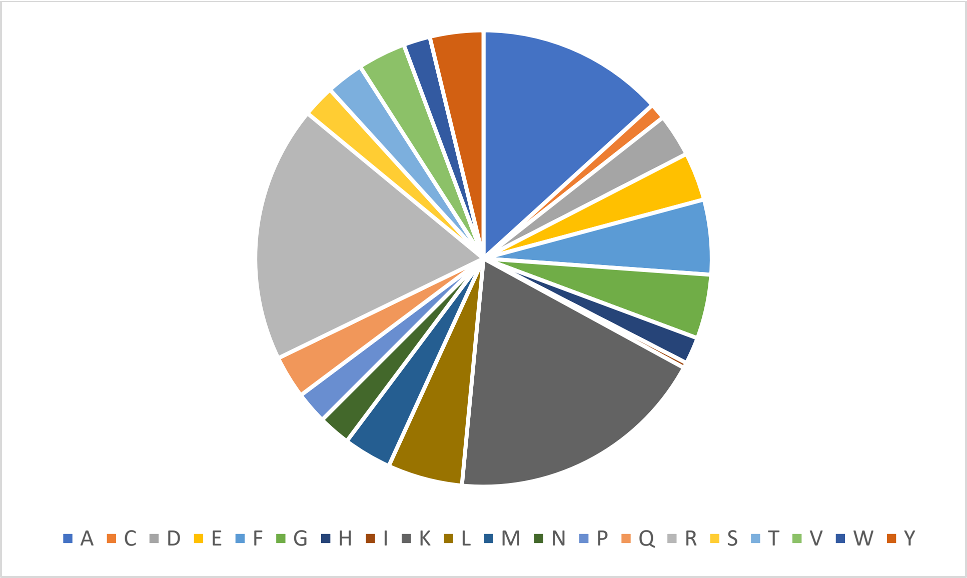


**Fig. S63.** Insights into proteolytic cleavage sites of arginylated peptides from human proteomes.

**Related to Discussion.**

**a**

| No. | protein | UniProt ID | peptide |
|---|---|---|---|
| 1. | LRC36 | Q1X8D7 | ALLPPE$^{R/R}_{711}$K |
| 2. | RL36 | Q9Y3U8 | AME$^{R/R}_{59}$LLK |
| 3. | LBHD2 | A0A0U1RRK4 | AVAGAWE$^{R/R}_{30}$K |
| 4. | KIF24 | Q5T7B8 | AVSIPE$^{R/R}_{75}$R |
| 5. | PI5L1 | Q5T9C9 | D$^{R/R}_{244}$LNFQGK |
| 6. | NOP2 | P46087 | GVNLD$^{R/R}_{336}$PLGK |
| 7. | CH60 | P10809 | IGIE$^{R/R}_{446}$IIK |
| 8. | FERM1 | Q9BQL6 | INQLYE$^{R/R}_{294}$QAR |
| 9. | RS17 | P08708 | LGND$^{R/R}_{27}$FHTNK |
| 10. | DDR1 | Q08345 | NC*LVGE$^{R/R}_{776}$NFTIK |
| 11. | TSH3 | Q63HK5 | NLTE$^{R/R}_{856}$SHTSK |
| 12. | HS71A | P0DMV8 | NQVALNPQNTVFD$^{R/R}_{69}$AK |
| 13. | HSP7C | P11142 | NQVAMNPTNTVFD$^{R/R}_{69}$AK |
| 14. | F124B | Q9H5Z6 | QVHC*GSE$^{R/R}_{136}$ILR |
| 15. | HGF | P14210 | SGLTC*SMWD$^{R/R}_{416}$K |
| 16. | HOOK3 | Q86VS8 | SLQD$^{R/R}_{533}$QGSK |
| 17. | ZFHX3 | Q15911 | TALE$^{R/R}_{2727}$AHIR |
| 18. | H4 | P62805 | TVTAMD$^{R/R}_{86}$VVYALK |
| 19. | TM131 | Q92545 | VASISFD$^{R/R}_{394}$ASK |
| 20. | RS19 | P39019 | VLQALE$^{R/R}_{108}$GLK |
| 21. | MBNL1 | Q9NR56 | WLTLE$^{R/R}_{17}$VC*R |
| 22. | NP1L1 | P55209 | YAVLYQPLFD$^{R/R}_{115}$K |

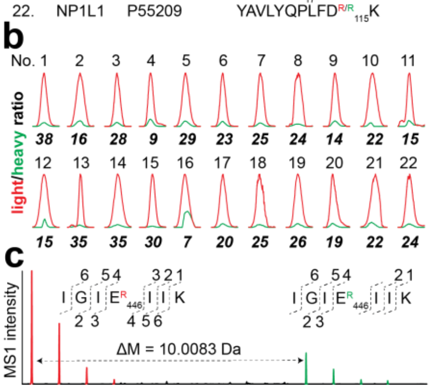


**Fig. S64.** Preliminary data on putative arginylation sites identified from HEK293T cells after pulse isotopic arginine treatment. **a,** list of identified peptides containing incorporated isotopic arginines (Arg$^{10}$ and Arg$^{0}$). *carbamidomethyl. **b**, extracted MS1 chromatograms and corresponding light/heavy ratios of co-eluting peptide pairs. **c**, MS1 characterization of a co-eluting peptide pair in protein CH60 as an example. MS1 isotopic envelopes for light and heavy Arg modified peptides.

**Related to Discussion.**

### Supplementary methods

#### Chemicals

The following key chemicals are used in this study.

| chemical | abbreviation | source | catalog No. |
|---|---|---|---|
| *L*-arginine hydrochloride | Arg0 or R0 | Silantes GmbH | 201003902 |
| *L*-[$^{13}C_6{}^{15}N_4$]arginine hydrochloride | Arg10 or R10 | Silantes GmbH | 201604102 |
| cycloheximide | CHX | Sigma Aldrich | C7698 |
| MG132 | | MedChem Express | HY-13259 |
| DIAALVHSSGNleS-$NH_2$ | Standard peptide | GenScript | N.A. |
| REPAVYFK | CALR E18 R | GenScript | N.A. |
| RDAPEEEDHVLVLR | PDIA1 D18 R | GenScript | N.A. |
| REEQPPETAAQR | ERO1A E24 R | GenScript | N.A. |
| RESETTTSLVLER | SSBP E17 R | GenScript | N.A. |
| REEAGDGTTTATVLAR | CH60 E107 R | GenScript | N.A. |
| RESSVAQQPLHTAQK | GATB E42 R | GenScript | N.A. |
| RC(O3)EALAGAPLDNAPK | RM12 C45 RO3 | GenScript | N.A. |
| Acetyl-DIAALVHSSGNleS-NH2 | Standard peptide | GenScript | N.A. |
| VIAALVHSSGNleS-NH2 | Standard peptide | GenScript | N.A. |
| D(R)IAALVHSSGNleS-NH2 | Standard peptide | GenScript | N.A. |
| E(R)PAVYFK | Standard peptide | GenScript | N.A. |
| E(R)SSVAQQPLHTAQK | Standard peptide | GenScript | N.A. |
| D(R)APEEEDHVLVLR | Standard peptide | GenScript | N.A. |
| E(R)EQPPETAAQR | Standard peptide | GenScript | N.A. |
| E(R)SETTTSLVLER | Standard peptide | GenScript | N.A. |
| E(R)EAGDGTTTATVLAR | Standard peptide | GenScript | N.A. |

#### Plasmids

The following plasmids are used in this study.

| construct | species | source | identifier | note |
|---|---|---|---|---|
| ATE1-1-his_pET30a(+) | human | GenScript | N.A. | codon-optimized |
| RARS1-his_pET30a(+) | human | GenScript | N.A. | codon-optimized |
| ATE1_pcDNA3.1+/C-(k)DYK | human | GenScript | OHu12263D | |
| ERO1A_pcDNA3.1+/C-(k)DYK | human | GenScript | OHu06553D | |
| 24R-ERO1A_pcDNA3.1+/C-(k)DYK | human | GenScript | OHu06553D | mutagenesis |
| ERO1A-V101G_pcDNA3.1+/C-(k)DYK | human | GenScript | OHu06553D | mutagenesis |
| ERO1A-E24V_pcDNA3.1+/C-(k)DYK | human | GenScript | OHu06553D | mutagenesis |
| SSBP_pcDNA3.1+/C-(k)DYK | human | GenScript | OHu19765C | |
| SSBP-E17V_pcDNA3.1+/C-(k)DYK | human | GenScript | OHu19765C | mutagenesis |
| 17R-SSBP_pcDNA3.1+/C-(k)DYK | human | GenScript | OHu19765C | mutagenesis |
| CALR_pcDNA3.1+/C-(k)DYK | human | GenScript | OHu23892 | |
| 18R-CALR_pcDNA3.1+/C-(k)DYK | human | GenScript | OHu23892 | Mutagenesis |
| Ub-K48R_ERO1A-flag_pRK5 | human | GenScript | OHu06553D | |
| Ub-K48R_24R-ERO1A-flag_pRK5 | human | GenScript | OHu06553D | mutagenesis |
| Ub-K48R_SSBP-flag_pRK5 | human | GenScript | OHu19765C | |
| Ub-K48R_17R-SSBP-flag_pRK5 | human | GenScript | OHu19765C | mutagenesis |
| CALR-HA-TEV-halo_pcDNA5/FRT | human | GenScript | OHu23892 | |
| 18R-CALR-HA-TEV-halo_pcDNA5/FRT | human | GenScript | OHu23892 | Mutagenesis |
| A1AT_pcDNA3.1+/C-(k)DYK | human | GenScript | OHu22141D | |
| 25R-A1AT_pcDNA3.1+/C-(k)DYK | human | GenScript | OHu22141D | Mutagenesis |
| MAPT_pcDNA3.1+/C-(k)DYK | human | GenScript | OHu28029C | |
| P4HB_pcDNA3.1(+)-C-HA | human | GenScript | OHu25382C | |
| JcM-Cys_pcDNA3.1(+)-myc-His A | mouse | GenScript | OMu02621C | |
| ATE1-2_pGEX6p | human | Addgene 79687 | NP_008972.2 | reference[1] |

#### Antibodies

The following antibodies are used in this study.

| antibody | source | catalog No. |
|---|---|---|

| | | |
|---|---|---|
| anti-FLAG M2 | Sigma-Aldrich | F3165 |
| β-tubulin antibody | Cell Signaling | 2146 |
| IRDye 680RD goat anti-mouse | LI-COR | 925-68070 |
| IRDye 800CW goat anti-rabbit | LI-COR | 925-32211 |
| Anti-ATE1 antibody, clone 6F11 | Sigma-Aldrich | MABS436 |
| Anti-P4HB antibody | Abcam | ab2792 |
| anti-Myc antibody | Sigma-Aldrich | 05-419 |
| anti-HA antibody | Cell Signaling | 3724S |
| anti-mouse IR 800CW | LI-COR | 925-32210 |
| anti-rabbit IR 680RD | LI-COR | 926-68071 |
| anti-mouse alexa fluor 647 | Thermo Scientific | A-21235 |
| anti-rabbit alexa fluor 568 | Thermo Scientific | A-11011 |

### Materials

HEK293T, MCF7, HCC2935, H1437, and A549 cells were purchased from ATCC and cultured following the manufacturer's protocols. HEK293T ATE1 KO cells were generated by CRISPR technology from HEK293T cells. The ATE1 KO was confirmed by sequencing which showed two events (A insertion and CA deletion) at exon 5 on two alleles. iPSC_CF, iPSC and iPSC_CM cells were prepared by Michael J. Greenberg lab with details described in a later section. HeLa cells were purchased from ATCC and prepared by Yi Zhang lab. Frozen brain samples were obtained from Anna Kashina lab. Detailed information on the Parkinson's disease patient samples and matching controls is listed in **Additional File 1** from a previous study[2], as follows: brain_c1(117849), healthy; brain_c2 (117504), healthy; brain_112568, Parkinson's; brain_108180, Parkinson's with dementia. brain_110007 sample from an Alzheimer's disease patient (age 88, female) was obtained from the same source in a previous report[3]. Frozen human heart tissue was obtained from the Translational Cardiovascular Biobank and Repository (Kory J. Lavine, IRB# 201104172). Frozen mouse lung, heart and brain tissues in-whole (C57/BL, male, 40 days old) were gifts from Professors Xia Liu and Guangyong Peng (Department of Otolaryngology-Head and Neck Surgery, Washington University School of Medicine). CALR with open 18E as N-terminal was purchased from Abcam (catalog No.: ab276554).

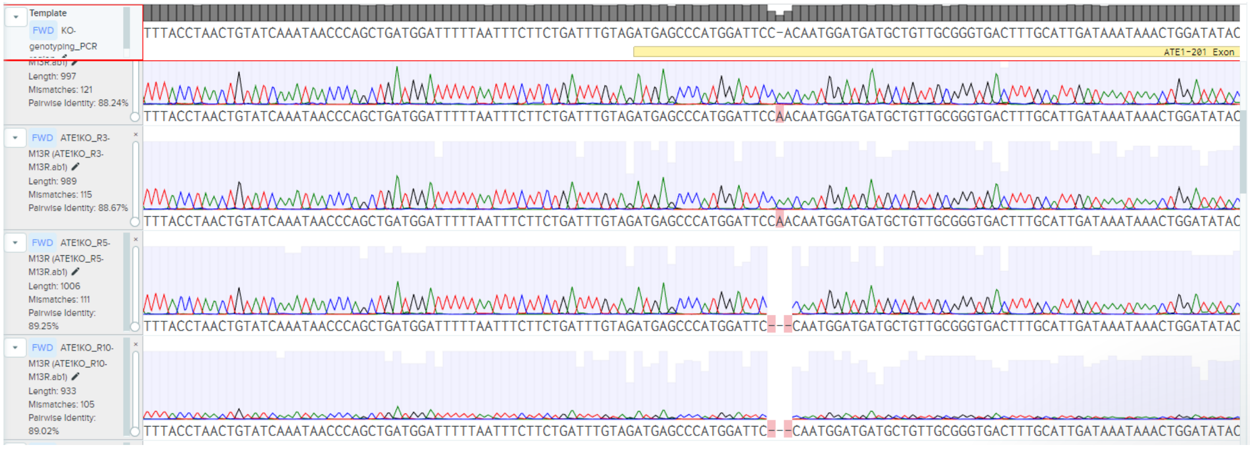

Note: confirmation of ATE1 KO in HEK293T cells.

### Human induced pluripotent stem cell culture and differentiation to cardiomyocytes and cardiac fibroblasts

Human induced pluripotent stem cells (iPSCs), stem cell derived cardiomyocytes, and stem cell derived cardiac fibroblasts were all derived from the same parent line (BJ fibroblast line CRL-2522, ATCC). The parent line was reprogrammed by the Washington University Genome Engineering and Stem Cell Core using the CytoTune127 iPS 2.0 Sendai reprogramming kit (A16517, ThermoFisher). We have previously shown that this line is pluripotent, and we have used this line previously to derive cardiac lineages [4-8]. Stem cells were grown as colonies and maintained in feeder-free conditions on Matrigel (Corning) coated plates in StemFlex media (ThermoFisher) as previously described [4-7]. Stem cells cells were passaged using 0.02% EDTA at least 4 times before starting the differentiation process.

Stem cell derived cardiomyocytes were derived from iPSCs in monolayer culture by temporally modulating WNT signaling [4,9]. Spontaneous beating was observed by days 9-12. On days 12-17, the population of cardiomyocytes was enriched using metabolic selection (No Glucose RPMI-1640 with B27 supplement plus lactate) as previously described [4,10]. Cells were aged at least 30 days, and experiments used cells derived from multiple differentiations. We routinely obtain >90% cardiomyocytes as assessed by immunofluorescence staining for the sarcomeric-specific marker, troponin T [4].

Stem cell derived cardiac fibroblasts were derived from iPSCs in monolayer culture using the method of Zhang et al. [11], as we have previously done [7,8]. Temporal modulation of WNT signaling differentiated iPSCs towards a mesoderm lineage, and then the cells were differentiated to cardiac fibroblasts by modulation of FGF, WNT, and TGFβ signaling. Using this procedure, we obtain cardiac fibroblasts as validated by RT-PCR measurements of the expression levels of cardiac fibroblast-specific genes, GATA4, and TCF21 and the general fibroblast genes COL1A1 and DDR2 [7].

**Expression and purification of RARS1 and ATE1 proteins**

BL21 (DE3*) (Invitrogen) and BL21-CodonPlus (Agilent Technologies, catalog No.: 230245) were used for expressing RARS1 and ATE1 respectively according to a previous study[12]. Bacteria expressing protein were grown in large cultures (1 L/flask) at 37 °C in rich LB broth to an optical density of 0.7-0.9 at 600 nm. The flasks were cooled rapidly on ice before induction with isopropyl β-d-1-thiogalactopyranoside (IPTG) at a final concentration of 1 mM. Cultures were grown at 16 °C for an additional 16-20 h and cells were harvested by centrifugation (5,000 g, 30 min, 4 °C), flash frozen in liquid nitrogen, and stored at -80 °C. Cells were lysed in lysis buffer (50 mM Tris, 500 mM NaCl, 5 mM β-ME, 0.5 mM PMSF, 10% glycerol, pH 7.8) at 4 °C by probe sonication. The His-tagged proteins were purified on the HisTrap HP column (Cytiva, catalog No.: 17-5248-01) in the lysis buffer containing 300 mM imidazole by FPLC (Cytiva). Fractions containing the protein were pooled and concentrated by centrifugation to a final volume of ~1 mL. Protein was loaded to the size exclusion column, eluted by elution buffer (50 mM Tris, 100 mM NaCl, 2 mM TCEP, 10% glycerol, pH 7.8), and concentrated in Millipore 30 kDa MWCO filters. Protein concentrations were determined and 10-μL aliquots were stored at -80 °C.

***In vitro* transcription and production of tRNA**

Double strand DNA template (TAATACGACTCACTATA-GGGCCAGTGGCGCAATGGATAACGCGTCTGACTACGGATCAGAAGATTCTAGGTTCGACTCCTGGCT GGCTCGCCA) containing a T7 RNA polymerase recognition sequence (TAATACGACTCACTATA) and a 76-base $tRNA^{arg}$ sequence[13] was ordered from IDT. The *in vitro* transcription was carried out following the manufacturer's protocol on HiScribe T7 High Yield RNA Synthesis Kit (NEB, catalog No.: E2040S) using 250 ng of DNA template for a 20-μL reaction incubated at 37 °C for 4 h. DNAase and its buffer were added (final volume 30 μL) to remove DNA template by incubation for 30 min at 37 °C. 170 μL ddH2O was added followed by 200 μL of phenol:chloroform:isoamylalcohol pH 6.7 (Sigma, catalog No.: 516726). The supernatant was added $1/10^{th}$ volume of 3 M NaOAc and 2.5 volumes of cold 200-proof ethanol, incubated at -20 °C for 20 min, and centrifuged at 4 °C. The pellet was air-dried until moist and resuspended in 20 μL ddH2O. tRNA concentration was determined by NanoDrop, and 2-μL aliquots were stored at -80 °C.

**Preparation of whole proteome peptide mixture**

HEK293T (ATE1 KO) cell pellet harvested from a 6-cm dish was added 200 μL ice-cold PBS, lysed by probe sonication (Fisherbrand™ Model 120 Sonic Dismembrator, setting: 1 sec on, 2 sec off, 20% energy, 60 cycles) to yield a homogeneous solution. Lysates were ultracentrifuged at 100,000 g for 30 min at 4 °C to yield soluble and insoluble proteomes. To the insoluble fraction, 200 μL PBS was added, and resuspended by probe sonication. Protein concentration was determined by DC assay (Bio-Rad). 100 μg of soluble and insoluble proteomes were diluted in 40 μL of 8 M urea in PBS, reduced by 5 mM of TCEP with 30 min incubation at 37 °C, alkylated by 15 mM of iodoacetamide (IAA) with 30 min incubation at room temperature in the dark. The solution was diluted to 2 M urea by 50 mM ammonium bicarbonate in H2O, digested by trypsin (sequence grade, Promega) at 1:50 trypsin/protein ratio (*w*/*w*) with overnight (~12 h) incubation at 37 °C. The resulting peptide solutions were acidified by formic acid at a final concentration of 5%, desalted, and resuspended in $H_2O$ for ATE1 assay.

#### Preparation of cell and tissue proteomes

Ice-cold PBS (200 and 400 µL for pellet from a 6-cm and 10-cm dish, respectively) was added to cell pellets which were then lysed by probe sonication (Fisherbrand™ Model 120 Sonic Dismembrator, setting: 1 sec on, 2 sec off, 20% energy, 60 cycles) to yield a homogeneous solution. A few milligrams (5-10) of tissue pieces/samples (human or mouse) were transferred to tubes (Thermo Scientific, catalog No.: 3468) and ice-cold PBS was added (200-400 µL). A scoop of beads (GB05 beads for the brain and ZRB05 beads for other tissues, Next Advance, Inc.) was added to each tissue. Samples were homogenized in the cold room using FastPrep-24 Classic bead beating grinder (FastPrep-24, MP Biomedicals) at default settings (4.0 m/s, 20 sec, 2 cycles). Lysates were ultracentrifuged at 100,000 g for 30 min at 4 °C to yield soluble and insoluble proteomes. To the insoluble fraction, the same volume of PBS as the soluble fraction was added to resuspend by probe sonication until homogeneity (5-10 cycles). Protein concentration was determined by DC assay (Bio-Rad).

#### Affinity purification of Halo-tagged CALR

C-terminal Halo-tagged CALR was cloned into the pcDNA5/FRT vector (Invitrogen) for mammalian expression[14]. The CALR coding sequence was amplified from clone ID OHu23892 (GenScript). The C-terminal version of the Halo tag was amplified from pFC14A (Promega, Madison, WI, USA). 18R-CALR was generated by inserting an Arg codon after the signal peptide of CALR using mutagenesis. Wildtype or ATE1-/- HEK293T cells were transfected with plasmids expressing Halo-tagged wildtype or 18R CALR using FuGENE® HD (Promega, Madison, WI, USA). For experiments where ATE1 was co-expressed, various amounts of the plasmid expressing flag-tagged ATE1 were also transfected together with CALR. After 2 days of transfection, cells were harvested, and pellets were snap-frozen with liquid nitrogen. Halo purification was performed according to the manual of HaloTag® Mammalian Pull-Down Systems (Promega, Madison, WI, USA). Briefly, thawed cell pellets were lysed with Mammalian Lysis Buffer with the addition of protease inhibitors (Promega, Madison, WI, USA). Crude lysates were centrifuged at 20,000 × g at 4 °C for 10 minutes, and supernatants were collected to bind with Magne® HaloTag® Beads (Promega, Madison, WI, USA). After rotating at 4 °C overnight, beads were washed with cold Wash Buffer 5 times, then eluted with AcTEV Protease (Invitrogen, Carlsbad, CA, USA) at room temperature for 1 hour with shaking. Eluates were subjected to sample processing for bottom-up mass spectrometry analysis.

#### Isotopic labeling by ATE1 arginylation assay

The ATE1 assay was adapted from a previous study[12]. The assay was set up on ice by mixing a 20-µL reaction containing 1x reaction buffer, 2 mM Arg (Arg10 and Arg0, respectively), 2 mM ATP, 3 µM $tRNA^{Arg}$, 1 µM RARS1, 3 µM human ATE1 (isoform 1 or 2), and 1 µg/µL substrate. The 5x reaction buffer consists of 50 mM HEPES, 30 mM KCl, and 10 mM $MgCl_2$. The substrate can be peptide, peptide mixture, protein, or proteome. A total of 20 µg substrate was always used unless the protein substrate was purified from overexpression and pulldown experiments where protein concentration was too low to quantify. The amount of commercial CALR was 20 µg/reaction. The concentration of standard peptide was 100 µM unless otherwise noticed. Volumes of ddH2O were adjusted every time depending on the volumes of other reagents. The pair reactions were incubated at 37 °C for 30 min for peptides, 1 h for proteins, and 2 h for proteomes. The pair reactions were mixed 1:1 for sample preparation or -80 °C storage.

| | | | heavy (Arg10) | light (Arg0) | |
|---|---|---|---|---|---|
| **order** | **reagent** | **stock concentration** | **volume (µL)** | **volume (µL)** | **final concentration** |
| 1 | ddH2O | N.A. | 5 | 5 | |
| 2 | Reaction buffer | 5X | 4 | 4 | 1X |
| 3 | Arg10 or Arg0 | 40 mM | 1 | 1 | 2 mM |
| 4 | ATP | 10 mM | 4 | 4 | 2 mM |

| | | | | | |
|---|---|---|---|---|---|
| 5 | Substrate | N.A. | 2 | 2 | 1 µg/µL for proteome |
| 6 | tRNA | 60 µM | 1 | 1 | 3 µM |
| 7 | RARS1 | 20 µM | 1 | 1 | 1 µM |
| 8 | ATE1 | 30 µM | 2 | 2 | 3 µM |

**Sample preparation after ATE1 arginylation assay**

The peptide sample was acidified by formic acid to a final concentration of 5%, desalted, dried by SpeedVac, and reconstituted for LCMS analysis. Whole-proteome peptide sample was acidified by formic acid to a final concentration of 5%, desalted, dried by SpeedVac, and reconstituted for peptide fractionation. Protein or proteome sample was added accurately 20 mg urea (final concentration: 6 M) to denature proteins, TCEP (final concentration: 10 mM) for 30 min incubation at 37 °C to reduce disulfides, and iodoacetamide (final concentration: 30 mM, final volume: 54 µL) for 30 min at room temperature protected from light to alkylate reduced thiols. The solution was diluted to 2 M urea by 50 mM ammonium bicarbonate in $H_2O$, digested by trypsin (sequence grade, Promega) at a 1:50 trypsin/protein ratio (*w*/*w*) with overnight (~12 h) incubation at 37 °C. The resulting peptide solution was acidified by formic acid at a final concentration of 5%, desalted, dried by SpeedVac, and reconstituted for peptide fractionation.

**Peptide desalting**

Peptides were resuspended in 0.1% formic acid and desalted by in-house packed stage-tips. Stage tips were self-prepared by sealing five disks of C18 material (cat. No.: 2315, Empore, 3M Company) at the bottom of a P200 tip. C18 disks were cut by sample corers (cat. No.: 18035-01 for peptides < 20 µg, cat. No.: 18035-02 for 20-100 µg peptides, Fine Science Tools). Stage tips were equilibrated with 50 µL of methanol, 50 µL of 80% acetonitrile in $H_2O$ containing 0.1% formic acid (FA), and 50 µL of water containing 0.1% FA by centrifugation (1,000 g, ~1-2 min). The sample was loaded to the stage-tip and centrifuged to flow through, washed with 75 µL of water containing 0.1% FA. The stage-tip was transferred to a new collection tube, eluted by 2x 75 µL of 80% acetonitrile in $H_2O$ containing 0.1% FA. The sample was dried by SpeedVac (SAVANT SVC100H Refrigerated Condensation Trap) under vacuum for 30-60 min at room temperature, stored at –80 °C.

**Peptide fractionation**

Peptides (20 µg, half material from ATE1 assay) were resuspended in 50 µL ammonium formate (AF, pH 10). Stage tips were self-prepared by sealing five disks of C18 material (cat. No.: 2315, Empore, 3M Company) at the bottom of a P200 tip. C18 disks were cut by sample corers (cat. No.: 18035-02, Fine Science Tools). Stage tips were equilibrated with 50 µL of methanol, 50 µL of 80% acetonitrile in $H_2O$ containing 0.1% formic acid (FA), and 50 µL of AF pH 10 by centrifugation (1,000 g, ~1-2 min). The sample was loaded to the stage-tip and centrifuged to flow through (fraction 1), then eluted by 20 µL of the following buffers consisting of MeCN and ammonium formate (AF, pH 10) to yield an additional 29 fractions. The fractions were dried by SpeedVac (SAVANT SVC100H Refrigerated Condensation Trap) under vacuum for 30-60 min at room temperature, and stored at –80 °C. Each fraction was resuspended in 20 µL of 2% MeCN in 0.1% FA and 5 µL were injected (~167 ng per fraction in average) for analysis.

| buffer | MeCN% | MeCN (µL) | AF, pH 10 (µL) | total (µL) |
|---|---|---|---|---|
| 2 | 1 | 10 | 990 | 1000 |
| 3 | 2 | 20 | 980 | 1000 |
| 4 | 3 | 30 | 970 | 1000 |
| 5 | 4 | 40 | 960 | 1000 |
| 6 | 5 | 50 | 950 | 1000 |
| 7 | 6 | 60 | 940 | 1000 |
| 8 | 7 | 70 | 930 | 1000 |
| 9 | 8 | 80 | 920 | 1000 |
| 10 | 9 | 90 | 910 | 1000 |
| 11 | 10 | 100 | 900 | 1000 |
| 12 | 11 | 110 | 890 | 1000 |
| 13 | 12 | 120 | 880 | 1000 |

| | | | | |
|---|---|---|---|---|
| 14 | 13 | 130 | 870 | 1000 |
| 15 | 14 | 140 | 860 | 1000 |
| 16 | 15 | 150 | 850 | 1000 |
| 17 | 16 | 160 | 840 | 1000 |
| 18 | 17 | 170 | 830 | 1000 |
| 19 | 18 | 180 | 820 | 1000 |
| 20 | 19 | 190 | 810 | 1000 |
| 21 | 20 | 200 | 800 | 1000 |
| 22 | 21 | 210 | 790 | 1000 |
| 23 | 22 | 220 | 780 | 1000 |
| 24 | 23 | 230 | 770 | 1000 |
| 25 | 24 | 240 | 760 | 1000 |
| 26 | 25 | 250 | 750 | 1000 |
| 27 | 26 | 260 | 740 | 1000 |
| 28 | 27 | 270 | 730 | 1000 |
| 29 | 28 | 280 | 720 | 1000 |
| 30 | 80 | 800 | 200 | 1000 |

**LCMS for proteomics analysis**

A Vanquish Neo UHPLC was coupled to an Exploris 240 or Orbitrap Ascend (Thermo Scientific). Peptide samples were maintained at 7 °C on the sample tray in the LC system. Separation of peptides was carried out on an Easy-Spray™ PepMap™ Neo nano-column (2 µm, C18, 75 µm X 150 mm) at room temperature with a mobile phase consisting of a linear gradient of A (0.1%FA in $H_2O$) and B (acetonitrile containing 0.1% FA) under the following conditions: 0 → 80 → 83 → 90 min, 0% → 27% → 100% → 100% B. The flow rate was 300 nL/min. The injection volume is 5 µL. The voltage applied to the nano-LC electrospray ionization source was 1.9 kV. The temperature of the ion transfer tube (ITC) was set at 275 °C. Spectra were collected in a data-dependent acquisition mode with MS1 scan range of *m/z* 350–2000 in the orbitrap. For Exploris, the top 20 most intense peaks from a single high-resolution (120,000) full MS spectrum of parent ions were fragmented for MS2 spectra. For Ascend, the time of a scan cycle is 3 seconds. Parent ions assigned as peptides in charge states +2-5 with intensity higher than 1E4 were included for fragmentation. HCD-induced fragmentation (MS2) scans were recorded in orbitrap (scan from *m/z* 150–1500). Dynamic exclusion was set as a repeat count of 1 within an exclusion time of 20 seconds. All other parameters were left as default values.

A Waters M5 UHPLC was coupled to a ZenoTOF 7600 (Sciex) in positive data-dependent mode. Peptide samples were maintained at 7 °C on the sample tray in LC. Separation of peptides was carried out on an Waters nanoEase M/Z Symmetry C18 Analytical Column, (5 µm, 100 Å, 300 µm X 150 mm) at room temperature with a mobile phase consisting of a linear gradient of A (0.1%FA in $H_2O$) and B (acetonitrile containing 0.1% FA) under the following conditions: 0 → 1 → 46 → 47 → 50 → 50.5 → 55 min, 2 → 2 → 32 → 80 → 80 → 2 → 2% B. The flow rate was set at 5 µL/min. The ZenoTOF 7600 system was operated using the OptiFlow TurboV ion source with a vertical microflow probe (1-50 µL/min electrode). The ionization window was 2.5-48.5 min. Source and gas parameters are: 20 and 60 psi for ion source gas 1 and 2 respectively, 35 and 7 psi for curtain and CAD gas respectively, and 200 and 35 °C for temperature and column temperature respectively. Spray voltage was 5 kV. The MS1 mass range was 350-1500 Da with an accumulation time of 0.1 second, declustering potential of 80 V, and collision energy of 10 V. Top 45 monoisotopic ions assigned as peptide in charge states +2-6 with intensity higher than 100 cps were included for fragmentation. Dynamic exclusion was set as a repeat count of 1 within an exclusion time of 7 seconds. MS2 CID fragmentation with Zeno trapping was carried out over a mass range 120-1600 Da with an accumulation time of 0.02 second, declustering potential of 80 V, time bins to sum of 8, and a Zeno threshold of 1E5 cps. All other parameters were left as default values.

**Search parameters using Byonic software**

Each data file (in ".raw" format) was generated by the instrument (Xcalibur software), and searched using the byonic software v4.5.2 (Protein Metrics) against a reverse-concatenated, nonredundant database of the human proteome (20398 proteins in total). Peptides were required to have at least a tryptic *C*-terminus and up to one missed cleavage was allowed in the database search. The mass tolerances of precursor and product ions were set to 10 and 20 ppm, respectively. Carbamidomethylation (+57.021464 Da) on Cysteine residues was allowed for 2 per peptide as a “common” variable modification in the byonic parameter file. Oxidation (+15.99492 Da) on Methionine residues was allowed for 2 per peptide as a “common” variable modification. Total “common” modifications were limited to up to 4 per peptide. Arginylation (+156.10111 Da as $Arg^{0}$ modification or +166.10937 as $Arg^{10}$ modification) on peptide *N*-terminal, Asp, and Glu residues, arginylation after deamidation (+157.08513 Da as Arg0 modification or +167.09339 as Arg10 modification) on Asn and Gln residues, arginylation after di-oxidation (+188.09094 Da as Arg0 modification or +198.09920 as Arg10 modification) or tri-oxidation (+204.08585 Da as Arg0 modification or +214.09411 as Arg10 modification) on Cysteine residues was allowed for 1 per peptide as a “rare” modification. Total rare modifications were limited to 1 per peptide (see screenshot below). Peptides with 2 or more arginylation modifications were not included in the search. The false-positive rate was set at 1% or lower.

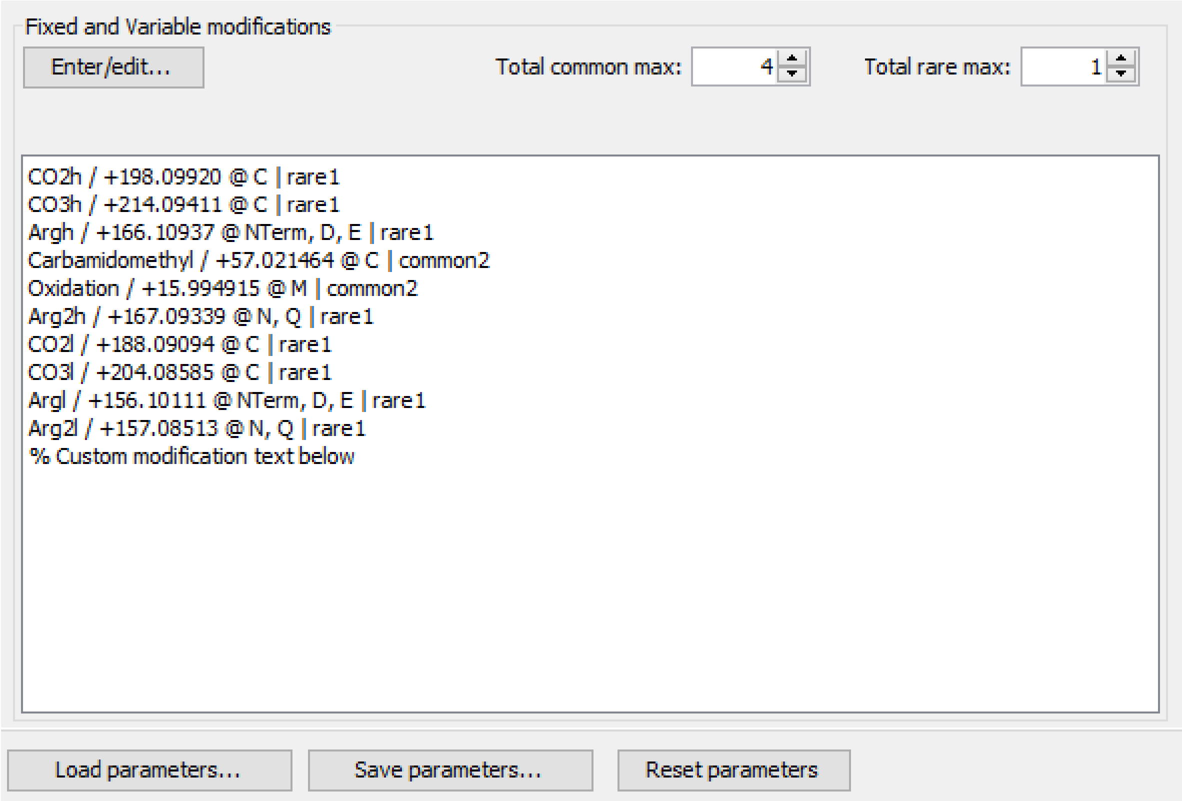


**ArginylomePlot software to determine arginylation sites after Byonic search**

Custom software based on *R* script, ArginylomePlot, was used to process all MS1 spectra of arginylated peptide and peptide IDs in Excel files generated from Byonic search. The software is publicly available to download from GitHub at https://github.com/BeckyHan/Garcia-Lab/tree/main/ArginylomePlot. A copy of the software together with a demo dataset is provided in the **Supplementary Dataset 1**. Briefly, .raw files from fractions of a sample were converted to .mxXML files using RawConverter (version 1.1.0.23) with monoisotopic selection (2015 released, publicly available at http://fields.scripps.edu/rawconv). The software first combines all peptide IDs into a single file, then extracts all co-eluting MS2 spectra with a mass difference of 10.008269 Da indicating co-presence of Arg10 and Arg0 modifications on the same peptide. Using the average retention time of all MS2 spectra from each arginylated peptide, the software then goes to a specific mzXML file and extracts their matching MS1 scans (doublet, quartet, or sextet matching with a threshold of 10 ppm for each m/z value) within a time window (±1.25 min of average retention time). The heavy/light ratio was calculated based on each MS1 scan, and the number of MS1 scans of each peptide was exported with respective ratios (doublet, quartet, or sextet). Failure to detect matching doublet MS1 will result in the exclusion of an MS2 pair. A ratio summary in a box plot (duodecet: “6+6” peaks) was generated for each arginylation site with a unique peptide sequence, charge, and raw. A summary table for each sample was generated with listed information on protein, unmodified peptide sequence, site, Arg10/Arg0 modified peptide ID, charge, raw, byonic scores of Arg10/Arg0 modified peptides, H/L ratios of doublet, quartet, or sextet MS1 scans, and their exact numbers of MS1 scans.

**Sequence logo preparation and motif analysis**

For sequence logo, all unique arginylated peptides with confidently identified arginylation sites were utilized as datasets to generate sequence logos. These peptides were initiated from the arginylated amino acid residues, aligned, and extended to include 14 positions downstream of the arginylation sites from HEK293T peptides, as well as 15 positions upstream and 14 positions downstream of the arginylation sites from all human proteomes. In cases involving C- and N-terminal peptides, the sequence was supplemented to 15 amino acids with the necessary number of "X"s, representing any amino acid. The sequence logos depicting these aligned peptide sequences were generated using the Weblogo program, employing a frequency plot.[15]

For motif analysis, potential motifs surrounding the identified arginylation sites were analyzed using pLogo[16]. Sequences spanning residues from -7 to +6 around the arginylation sites were extracted and compared to their corresponding background frequencies in the human proteome. Significant motifs ($p < 0.05$, Bonferroni corrected) were then visualized.

**Establishing a publicly available arginylation website**

A custom python script was written to enable data indexing of the arginylation database. The source code of the website is publicly available to download from GitHub at https://github.com/ChenfengZhao/Arginylation. The website is publicly available at www.arginylation.com (available during the review process, unavailable before publication, and available after publication). The website allows searching for sample (cell or tissue), proteins, and peptides. Alternatively, it also allows the manual selection of species, sample, protein, peptide, fraction of a sample, charge of a peptide for data visualization. Each MS1 or MS2 scan is available to be selected to view. Mass errors in both MS1 and MS2 can be changed manually with a default value of 20 ppm. MS2 ion types are available for selection with a default of b and y associated ions. This website allows for downloading detailed source MS1 and MS2 data in Excel format at species or sample levels, the downloadable data are organized based on unique peptide sequence, charge, and raw.

**In-bacteria arginylation assay**

In-bacteria arginylation assay was established with full details in our recent work[17]. Briefly, His-tagged protein or peptide is fused to ubiquitin and is co-expressed with human ATE1 and Ulp1 protease in *E. coli*. Following Ulp1 cleavage, protein N-terminal residue is exposed for arginylation by ATE1. Human ATE1 and Ulp1 protease were cloned into a pETDuet vector without tags. Substrate proteins ERO1A and SSBP1 were each cloned into a pCIOX vector with a C-terminal His-tag in two forms: starting from residues $E_{24}$ and $E_{17}$ respectively, or as peptides fused to ubiquitin ($E_{24}$EQPPETAAQ$R_{34}$-Ub for ERO1A and $E_{17}$SETTTSLVLERSLN$R_{32}$-Ub for SSBP1). These two plasmids were co-transformed and expressed in BL21(DE3)-RIL cells (Agilent). Protein production was induced with 0.2 mM IPTG overnight at 16 °C in Luria broth (Fisher). The His-tagged proteins were purified on Ni-NTA beads (GoldBio) in 50 mM Tris-HCl (pH 7.5) buffer, supplemented with 0.5 M NaCl, 5 mM *β*-ME, and 10 mM imidazole. Proteins were eluted in buffer containing 250 mM imidazole, further purified by size exclusion chromatography, and concentrated in Millipore concentrators. Arginylation states of each substrate protein were analyzed by Mass Spectroscopy.

**Sample preparation of protein or pulldown samples**

Protein (20 μg for pure protein or uncertain amount for pulldown protein) was denatured in 6 M urea, reduced by TCEP (final concentration: 10 mM) for 30 min incubation at 37 °C, and alkylated by iodoacetamide (final concentration: 30 mM) for 30 min at room temperature protected from light. The solution was diluted to 2 M urea by 50 mM ammonium bicarbonate in H2O, digested by trypsin (sequence grade, Promega) at a 1:50 trypsin/protein ratio (*w*/*w*) with overnight (~12 h) incubation at 37 °C. The resulting peptide solution was acidified by formic acid at a final concentration of 5%, desalted according to a previous study[18], dried by SpeedVac, and reconstituted for LCMS analysis. To the affinity-purified protein, urea powder was added directly to a final concentration of 6 M, and 1 μg of trypsin was used for each sample for digestion.

**Top-down analysis for protein arginylation**

Intact ERO1A and SSBP1 peptide-ubiquitin species were analyzed via Eksigent M5 LC coupled to a Sciex 7600 ZenoTOF mass spectrometer adapted from a recent study[19]. Intact protein was injected onto a Waters NanoEase BEH C4 column, 1.7 μm particle size, 300 Å pore size, 300 μm ID and 50 mm in length. The protein was separated over 7-minute gradient from 20%-90% B at 10 μL/min. The mass spectrometer was operated in the intact protein (> 5 kDa) protein mode. MS1 scans were collected over a scan range of 500-2000 m/z with 100 ms accumulation time. For MS2 scans, the instrument was operated in $MRM^{HR}$ mode and three charge states per species were targeted. Species were accumulated for 250 ms and fragmented with EAD fragmentation using 7 eV electron kinetic energy, 5000 nA electron beam current, and 5 ms reaction time. MS2 spectra were collected over a scan range of 100-2000 *m/z* with Zeno pulsing on. The resulting .WIFF files were converted to .MZML using MSConvert with the peak picking filter on all MS levels selected. The converted files were then deconvolved using FLASHDeconv to decharge and deisotope all MS1 and MS2 spectra using default parameters. Deconvolved monoisotopic MS2 data was visualized using ProSight Lite with matching fragment tolerance set to 10 ppm.

**Cell culture and transfection**

HEK293T cells were cultured in Dulbecco's Modified Eagle's Medium (DMEM) (with high glucose, *L*-glutamine and sodium pyruvate) supplemented with 10% (*v*/*v*) fetal bovine serum (FBS) (Corning), penicillin (100 U/mL)/streptomycin (100 μg/mL) at 37 °C in a humidified atmosphere containing 5% $CO_2$. Cells were grown to ~40% confluence under standard growth conditions before transfection. A standard transfection ratio of vector/transfection reagent (3:1, *w*/*w*) was used[18]. 4 μg plasmid in expression vector and 12 μg of neutralized (pH 7) polyethyleneimine (PEI, Polysciences Inc.) 'MAX' (MW 40,000, 1 mg/mL) were premixed in 200 μL FBS-free DMEM medium, the resulted mixture was incubated at room temperature for 30 min before added to a 6-cm dish. 10 μg plasmid and 30 μg PEI in 500 μL FBS-free DMEM was used for a 10-cm dish. 1 μg plasmid and 3 μg PEI in 50 μL FBS-free DMEM was used for a well in a 6-well plate. Empty vector was used as a control group ('mock') of transfection. Cells were incubated for ~48 h before harvest by scratching on ice.

**Western blotting**

Cell pellets were lysed by probe sonication in PBS. Protein concentration was determined by DC assay (Bio-Rad). Samples were diluted to 1 mg/mL followed by adding 4X sodium dodecyl sulfate (SDS) loading buffer. Samples were resolved by SDS-PAGE (acrylamide mini gel from Invitrogen). Proteins in gel were transferred to polyvinylidene difluoride (PVDF, MilliporeSigma) in Towbin buffer using a Trans-Blot® SD Semi-Dry Transfer Cell (Bio-Rad). The membrane was blocked for ~1 h at room temperature with 5% nonfat dry milk (*w*/*v*) in Tris-buffered saline with 0.05% Tween 20 (TBST) and incubated with primary antibody in the same solution overnight at 4 ºC or 1 h at room temperature. The membrane was washed (3 × 2 min, TBST), incubated with secondary antibody (1:10,000) in milk for 1 h at room temperature, washed with TBST (3 × 1 min) and TBS (3 × 1 min), and visualized on a Li-Cor scanner (LI-COR Biosciences, v1.0.0.55).

**Flag pulldown and purification**

Cell pellets were lysed by probe sonication in lysis buffer (50 mM HEPES/150 mM NaCl, pH 7.5) (400 µL for a pellet from 10-cm dish, and 200 µL for a pellet from 6-cm dish). Lysates were ultracentrifuged at 100,000 g for 30 min at 4 °C to yield soluble and insoluble proteomes. The supernatant was transferred to a new tube and 50 µL slurry of Anti-FLAG® M2 Magnetic Beads (catalog No.: M8823, Sigma-Aldrich) was added. NP40 was added to a final concentration of 1%. The mixture was rotated at 4 ºC for 3 h. Using a magnetic strip, the magnetic beads were washed 3x by HEPES/NaCl/NP40 buffer, 3x by PBS. 50 µL of 1% SDS (or 150 ng/µL 3x FLAG peptide) was added to the beads, and incubated at 37 °C for 10 min. Elution was transferred to a new tube for further analysis.

**Protein degradation assay**

The experiment was carried out according to a previous study[20]. HEK293T ATE1 KO cells in 6-well plates were transfected transiently with Ub_ERO1A or Ub_SSBP plasmids, cultured for two days, and treated with MG132 and CHX. Cells were harvested by scraping on ice for Western blotting analysis. For CHX chase experiment, the cells were transfected in 6-cm dishes for two days, treated with CHX at 100 µg/mL for 0-8 hours at different times, and harvested at the same time by scraping on ice.

**Cell mitochondria stress assay**

The experiment was carried out according to a previous study[21]. Oxygen concentrations were measured in XFe24 Extracellular Flux Analyzer (Seahorse Bioscience). 40k HEK293T cells were seeded in XFe24 cell plates preincubated with poly-D-lysine for 1 h (100 µL/well, 50 µg/mL). Cells were transfected with 200 ng plasmid (SSBP or 17R-SSBP) which was co-incubated with 600 ng PEI in 20 µL medium for 30 min. After two days, media was aspirated, added 1 mL of prewarmed Seahorse XF DMEM Medium pH 7.4 (Agilent, catalog No.: 103575-100) containing 1 mM pyruvate, 2 mM glutamine, and 10 mM glucose, incubated for 1 h at 37 °C in a non-CO2 incubator. XF DMEM medium was aspirated and 500 µL was added right before measurement. Oligomycin, FCCP, and Rot/AA were prepared to a final concentration of 1.5, 0.5, and 0.5 µM respectively according to the user guide of Agilent Technologies XF CELL MITO STRESS TEST KT (Agilent, catalog No.: 103015-100). Compounds were added to the ports on the XF24 sensor cartridge pre-incubated with calibrant solution (1 mL/well) overnight at 37 °C in a non-CO2 incubator. Wells containing no cells were used as a background group for the XFe24 analysis.

***In vivo* oxidative folding assay**

The *in vivo* myc-tagged Ig-J chains (JcM) folding assays were performed based on a previous study[22]. Hek293T cells in a 6-cm dish were co-transfected with JcM and Ero1α (WT, 24R, or V101G) for 48 h. A mock vector was used as a control group. Cells were harvested by scraping on ice and incubated in 200 µL DMEM (prewarmed at 37 °C) containing 5 mM DTT for 5 min at 37 °C. After two washes in ice-cold PBS, cells were resuspended in fresh 800 µL DMEM (prewarmed at 20 °C) at 20 °C and aliquoted to four tubes (200 µL/tube). Aliquots were quenched by 20 mM NEM at different time points (0, 2, 4, and 8 min), and centrifuged to remove the supernatant. The cell pellet was lysed in RIPA buffer containing a protease inhibitor cocktail and 20 mM NEM. Lysate was probe sonicated, and the supernatants were loaded and resolved by nonreducing 16% SDS-PAGE for Western blotting analysis.

**Flag pulldown for ERO1A interaction with PDI**

After 48 h of transfection, cells were collected and washed with 1X PBS once. Cells were lysed with RIPA lysis buffer (Sigma Aldrich, catalog No.: 20-188) containing 1× EDTA-free protease inhibitor cocktail (Bimake, B1400) and sonication 3 cycles (10 sec on, 5 sec off, 20% amplitude) (Fisher Brand, catalog No.: FB120). Protein concentrations were determined using a DC assay kit (BioRad, catalog No.: 5000111) on a multi-mode microplate reader (Biotek Synergy 2). Cell lysates were diluted with lysis buffer to 2.5 µg/µL and 500 µg of protein was incubated with 50 µL of a 50% slurry of anti-FLAG M2 magnetic beads (Sigma-Aldrich, catalog No.: M8823) overnight at 4 °C with rotation. Beads were washed with rotation 3 times with 1X PBS (1 mL) for 5 min. Enriched proteins were eluted with 50 µL of 200 nM 3X Flag peptide for 30 min at 37 °C. 17 µL of 4X sample buffer was added to the samples and 25 µL of sample was loaded for SDS-

PAGE and immunoblotting. For immunoblotting analysis, proteins were transferred to a nitrocellulose membrane using an iBlot 2 system (Thermo Scientific). Membranes were blocked with TBS containing 0.1% Tween-20 and 3% BSA (Sigma Aldrich, catalog No.: A9647) and incubated with primary antibodies (diluted 1:1,000) anti-Flag (M2) (Sigma Aldrich, catalog No.: F3165), anti-HA (Cell Signaling, catalog No.: 3724S), anti-PDI (Abcam, catalog No.: ab2792), and secondary antibodies (diluted 1:10,000) anti-mouse IR 800CW (Licor, catalog No.: 925-32210) and anti-Rabbit IR 680RD (Licor, catalog No.: 926-68071) sequentially. Immunoblot images were acquired using a Licor Scanner and analyzed with Licor Image Studio. All infrared fluorescence western blot images were converted to grayscale in image studio and unsaturated exposure images were used for quantification.

**Cell imaging**

Cells were seeded on 22 mm × 22 mm glass coverslips (No. 1.5) coated with collagen (Neuvitro Corporation, GG-22-15-Collagen) that had been placed in single wells of a 6-well plate for 24 h before transfection. For the ER-GFP marker, 100 µL of CellLight™ ER-GFP, BacMam 2.0 (Thermo Scientific, catalog No.: C10590) was added directly to the cells during transfection. Cells expressing Halo proteins (CALR-halo) were stained with HaloTag® TMRDirect™ Ligand (Promega, Madison, WI, USA) overnight in their regular culture medium. After 48 h of transfection, cells were washed with 1X PBS twice and fixed in freshly prepared 4% paraformaldehyde in 1X PBS for 20 min at 37 °C. After washing with PBS twice, cells were permeabilized and blocked with blocking buffer (1× PBS, 3% BSA, 0.3% Triton X-100) for 1 h at room temperature. The primary and secondary antibodies were diluted with blocking buffer (1× PBS, 3% BSA, 0.3% Triton X-100). Cells were incubated with diluted primary antibodies (1:1,000) anti-Flag (M2) (Sigma, catalog No.: F3165), anti-HA (Cell Signaling, catalog No.: 3724S) or anti-Cox4 (Cell Signaling, 3E11, 1:200) overnight at 4 °C. Cells were rinsed with 1X PBS three times followed by a 1-h incubation with secondary antibodies (diluted 1:1,000) anti-mouse Alexa Fluor 647 (Thermo Scientific, catalog No.: A-21235), anti-rabbit Alexa Fluor 568 (Thermo Scientific, catalog No.: A-11011) at room temperature in the dark. Cells were washed with 1X PBS three times and incubated with NucBlue Fixed Cell Stain ReadyProbes reagent (Invitrogen, catalog No.: R37606) for 5 min at room temperature. Coverslips were washed with 1X PBS, mounted in Antifade Diamond (Life Technologies, catalog No.: P36961) on glass slides, and sealed with gel polish. HaloTag® TMRDirect™ was excited by 543 nm laser and detected at 553-753 nm. Images were collected on a Zeiss LSM880 with an Airyscan laser-scanning confocal microscope and exported to ZEN Blue for final processing and assembly.
