## Supplemental Dataset 1 for "An Unbiased Proteomic Platform for ATE1-based Arginylation Profiling": ArginylomePlot_v12.3.Rmd

````
---
title: "auto_plot MS1_ratio&scan"
output: html_document
date: "2023-01-04"
---

```{r 1merge}
library(tidyverse)
library(readxl)
library(openxlsx)
library (htmltools)
library(Hmisc)
# Find all .csv files
files <- dir("./byonic", recursive=TRUE, full.names=TRUE, pattern="\\.xlsx$")

# Apply the function to all files.
result <- files %>% lapply(read_excel, sheet = 2) %>% bind_rows
#%>% rename(Peptide = `Peptide\n< ProteinMetrics Confidential >`), col_types = "text"
colnames(result) <-  c("Query", "Protein_Rank", "Peptide", "Glycans_NHFAGNa", "Modification", "Observed_m_z", "z", "Observed_MH", "Calcmass", "Offbyerror", "Masserror", "StartingPosition", "Cleavage", "Score", "Delta", "DeltaMod", "LogProb", "uniquePeptides", "ProteinName", "ProteinDBnum","Comment", "Scannum", "ScanTime")
result1 = result %>% filter(!(startsWith(ProteinName, ">Reverse >")))
write.xlsx(result1, "1combine.xlsx")
```


```{r 2heavy_extract}
heavy_list = list.files(pattern = "1combine.xlsx", full.names = TRUE)  
data_heavy = heavy_list %>% lapply(read_excel) %>% bind_rows #, col_names = TRUE, skip = 1, sheet = 1, , col_types = "text"
funb <- function(x) gsub("[^0-9A-Za-z///']", "", x)
data_mutate = data_heavy %>% mutate(modi = funb(.[[5]])) 
toMatch <- c('198','214','166','167')
match_heavy <- data_mutate %>% filter(., grepl(paste(toMatch, collapse="|"), modi)) %>% select(-modi) %>% arrange(Peptide, desc(Score))
#write.xlsx(match_heavy, "2.1heavy_match.xlsx")

# {r 2.2light_extract}
light_list = list.files(pattern = "1combine.xlsx", full.names = TRUE)  
data_light = light_list %>% lapply(read_excel) %>% bind_rows #, col_names = TRUE, skip = 1, sheet = 1, , col_types = "text"
funb <- function(x) gsub("[^0-9A-Za-z///']", "", x)
data_mutate = data_light %>% mutate(modi = funb(.[[5]])) 
toMatch <- c('188','204','156','157')
match_light <- data_mutate %>% filter(., grepl(paste(toMatch, collapse="|"), modi)) %>% select(-modi) %>% arrange(Peptide, desc(Score))
#write.xlsx(match_light, "2.2light_match.xlsx")

```

```{r 3pair_match}
library(tidyverse)
library(readxl)
library(openxlsx)
library(stringr)
funb <- function(x) gsub("[^a-zA-Z]", "", x)
match_heavy$Peptide <- gsub("\\.\\-", "U", match_heavy$Peptide)
heavy = match_heavy %>% mutate(peptide1 = funb(.[[3]])) %>% mutate(pep1 = str_sub(peptide1, 2, -2)) %>% separate(Comment, into=c("raw","other"), sep = "[.]", remove = FALSE, extra = "drop")
heavy$Peptide <- gsub("U", "\\.\\-", heavy$Peptide)
heavy$peptide1 <- gsub("U", "\\.\\-", heavy$peptide1)

match_light$Peptide <- gsub("\\.\\-", "U", match_light$Peptide)
light = match_light %>% mutate(peptide1 = funb(.[[3]])) %>% mutate(pep1 = str_sub(peptide1, 2, -2)) %>% separate(Comment, into=c("raw","other"), sep = "[.]", remove = FALSE, extra = "drop")
light$Peptide <- gsub("U", "\\.\\-", light$Peptide)
light$peptide1 <- gsub("U", "\\.\\-", light$peptide1)


merge1 = inner_join(heavy, light, by = c("ProteinName", "pep1", "raw")) %>% filter(abs(Calcmass.x - Calcmass.y) >= 9.998269 & abs(Calcmass.x - Calcmass.y) <= 10.018269) %>% filter(abs(ScanTime.x - ScanTime.y) <= 3) %>% unique(.)
data_enddash = merge1 %>% filter(str_detect(Peptide.x, ".-$")) %>% mutate(sub_pep = str_sub(peptide1.x, 2, -1)) %>% mutate(sub_pep_len = nchar(sub_pep)) %>% filter(sub_pep_len >= 5)
data_other = merge1 %>% filter(!(str_detect(Peptide.x, ".-$"))) %>% mutate(sub_pep = str_sub(peptide1.x, 2, -2)) %>% mutate(sub_pep_len = nchar(sub_pep)) %>% filter(sub_pep_len >= 5)
data_com = rbind(data_enddash, data_other) %>% arrange(pep1, desc(Score.x), desc(Score.y))
write.xlsx(data_com, "HLmatch.xlsx")

```


```{r 4outputfiles}
library(tidyverse)
library(readxl)
library(openxlsx)
library(stringr)
library(RaMS)

source = read_excel("HLmatch.xlsx", sheet = 1) %>% filter(!(startsWith(ProteinName, ">Reverse >")))

source_tmp1 = source %>% select(raw, sub_pep, Calcmass.y, z.x, z.y, ScanTime.x, ScanTime.y) %>% unique(.)
source_tmp2 = source_tmp1 %>% mutate(ScanTimeTemp = (ScanTime.x + ScanTime.y)/2) %>% group_by(raw, sub_pep, Calcmass.y) %>% summarise(ScanTimeMean = mean(ScanTimeTemp))
source_tmp3 = left_join(source_tmp2, source_tmp1[,-c(6,7)], by = c("sub_pep", "Calcmass.y")) %>% unique(.) %>% pivot_longer(., cols = starts_with("z."), names_to = "zn", values_to = "Z") %>% select(-zn) %>% unique(.) %>% pivot_longer(., cols = starts_with("raw."), names_to = "rawn", values_to = "raw") %>% select(-rawn) %>% unique(.)
source_final = left_join(source_tmp2, source_tmp3[,-c(3)], by = c("raw", "sub_pep", "Calcmass.y")) %>% unique(.) %>% arrange(sub_pep, raw)
source_final$sub_pep <- gsub("\\.\\-", "", source_final$sub_pep)

write.xlsx(source_final, "HLmatch_3min.xlsx")

source_temp4.1 = source %>% group_by(ProteinName, pep1, StartingPosition.x, Peptide.x, Peptide.y, Modification.x, Modification.y, z.x, raw) %>% summarise(maxScoreH = max(Score.x), maxScoreL = max(Score.y)) %>% ungroup(.)
source_temp4.2 = source_temp4.1 %>% arrange(pep1, raw, z.x) 
write.xlsx(source_temp4.2, "output_peptides.xlsx")

source_temp4.3 = source %>% group_by(ProteinName, pep1, StartingPosition.x) %>% summarise(maxScoreH = max(Score.x), maxScoreL = max(Score.y)) %>% ungroup(.)
source_temp4.4 = source_temp4.3 %>% arrange(pep1) 
write.xlsx(source_temp4.4, "output_proteins.xlsx")

```


```{r 5export}
total_row = nrow(source_final)
mainDir = './results/individual'

for (i in 1:total_row) # check rows.
{
subDir = paste(source_final$sub_pep[i], 'z',source_final$Z[i], source_final$raw[i],'row', i, sep = '_')
plotname = paste(source_final$sub_pep[i], 'z',source_final$Z[i], source_final$raw[i], sep = '_')
dir.create(file.path(mainDir, subDir))
#setwd(file.path(mainDir, subDir))

raw = paste(source_final$raw[i], '.mzXML', sep = '')
msdata <- grabMSdata(files = paste('./mzxml/',raw, sep = ''), grab_what = c("MS1"))
calculatedmass = source_final$Calcmass.y[i] # [M+H]+
z = source_final$Z[i]
m = (calculatedmass - 1.00728 + z*1.00728)/z # L m/z
diff = 10.008269
retentiontime =  source_final$ScanTimeMean[i]

df = msdata$MS1 %>% filter(abs(rt-retentiontime) < 1.25) %>% select(-filename) 

df1 = df %>% filter(abs((mz - m)/m*10^6) <= 10) 
df2 = df %>% filter(abs((mz - (m + 1*1.00335/z))/(m + 1*1.00335/z)*10^6) <= 10) 
df3 = df %>% filter(abs((mz - (m + 2*1.00335/z))/(m + 2*1.00335/z)*10^6) <= 10) 
df4 = df %>% filter(abs((mz - (m + 3*1.00335/z))/(m + 3*1.00335/z)*10^6) <= 10) 
df5 = df %>% filter(abs((mz - (m + 4*1.00335/z))/(m + 4*1.00335/z)*10^6) <= 10) 
df6 = df %>% filter(abs((mz - (m + 5*1.00335/z))/(m + 5*1.00335/z)*10^6) <= 10) 

df7 = df %>% filter(abs((mz - (m + diff/z))/(m + diff/z)*10^6) <= 10) 
df8 = df %>% filter(abs((mz - (m + diff/z + 1*1.00335/z))/(m + diff/z + 1*1.00335/z)*10^6) <= 10) 
df9 = df %>% filter(abs((mz - (m + diff/z + 2*1.00335/z))/(m + diff/z + 2*1.00335/z)*10^6) <= 10) 
df10 = df %>% filter(abs((mz - (m + diff/z + 3*1.00335/z))/(m + diff/z + 3*1.00335/z)*10^6) <= 10) 
df11 = df %>% filter(abs((mz - (m + diff/z + 4*1.00335/z))/(m + diff/z + 4*1.00335/z)*10^6) <= 10) 
df12 = df %>% filter(abs((mz - (m + diff/z + 5*1.00335/z))/(m + diff/z + 5*1.00335/z)*10^6) <= 10) 

#df_org = inner_join(df1, df7, by = c("rt")) %>% left_join(., df2, by = c("rt")) %>% left_join(., df3, by = c("rt")) %>% left_join(., df5, by = c("rt")) %>% inner_join(., df6, by = c("rt"))

#colnames(df_org) <- c("rt", "mz1", "int1", "mz2", "int2", "mz3", "int3", "mz4", "int4", "mz5", "int5", "mz6", "int6")

#df_org = df_org %>% mutate(mz1_avg = mean(mz1), mz4_avg = mean(mz4)) %>% mutate(error = (mz1_avg - m) / m * 10^6, error4 = (mz4_avg-(m + diff/z))/(m + diff/z) * 10^6) 


df_org1 = inner_join(df1, df7, by = c("rt")) %>% left_join(., df2, by = c("rt")) %>% left_join(., df3, by = c("rt")) %>% left_join(., df4, by = c("rt")) %>% left_join(., df5, by = c("rt")) %>% left_join(., df6, by = c("rt")) %>% left_join(., df8, by = c("rt")) %>% left_join(., df9, by = c("rt")) %>% left_join(., df10, by = c("rt")) %>% left_join(., df11, by = c("rt")) %>% left_join(., df12, by = c("rt"))
colnames(df_org1) <- c("rt", "mz1", "int1", "mz7", "int7", "mz2", "int2", "mz3", "int3", "mz4", "int4","mz5", "int5","mz6", "int6","mz8", "int8", "mz9", "int9", "mz10", "int10","mz11", "int11","mz12", "int12") 
df_org1 = df_org1 %>% select(rt, mz1, int1,  mz2, int2, mz3, int3, mz4, int4,mz5, int5,mz6, int6,mz7, int7,mz8, int8, mz9, int9, mz10, int10,mz11, int11,mz12, int12)

write.xlsx(df_org1, file = paste(mainDir, subDir,"df_filter.xlsx", sep = "/")) 

calculate_ratio_twonum <- function(a, b) {
  if (!is.na(a) && !is.na(b)) {
    ratio <- a / b
    return(ratio)
  } else {
    return(NA)
  }
}

calculate_ratio1 <- function(a, b, c, d) {
  if (!is.na(a) && !is.na(b) && !is.na(c) && !is.na(d) ) {
    sumA <- sum(a, b)
    sumB <- sum(c, d)
    ratio <- sumB / sumA
    return(ratio)
  } else {
    return(NA)
  }
}


calculate_ratio2 <- function(a, b, c, d, e, f) {
  if (!is.na(a) && !is.na(b) && !is.na(c) && !is.na(d) && !is.na(e) && !is.na(f)) {
    sumA <- sum(a, b, c)
    sumB <- sum(d, e, f)
    ratio <- sumB / sumA
    return(ratio)
  } else {
    return(NA)
  }
}

df_ratio1 = df_org1
df_ratio1$ratio1 = apply(df_ratio1[,c("int1", "int7")], 1, function(x) calculate_ratio_twonum(x[1], x[2]))
df_ratio1$ratio2 = apply(df_ratio1[,c("int2", "int8")], 1, function(x) calculate_ratio_twonum(x[1], x[2]))
df_ratio1$ratio3 = apply(df_ratio1[,c("int3", "int9")], 1, function(x) calculate_ratio_twonum(x[1], x[2]))
df_ratio1$ratio4 = apply(df_ratio1[, c("int1","int2", "int7","int8")], 1, function(x) calculate_ratio1(x[1], x[2], x[3], x[4]))
df_ratio1$ratio5 = apply(df_ratio1[, c("int1","int2", "int3","int7","int8", "int9")], 1, function(x) calculate_ratio2(x[1], x[2], x[3], x[4], x[5], x[6]))

df_ratio1 = df_ratio1 %>% mutate(nRatio1 = sum(!is.na(ratio1)), nRatio2 = sum(!is.na(ratio2)), nRatio3 = sum(!is.na(ratio3)), nRatio4 = sum(!is.na(ratio4)), nRatio5 = sum(!is.na(ratio5)))
write.xlsx(df_ratio1, file = paste(mainDir, subDir, "ratios.xlsx", sep = "/"))

df_plot = df_org1 %>% mutate(mz1.1 = int1 / int1, mz1.2 = int2 / int1, mz1.3 = int3 / int1, mz1.4 = int4 / int1, mz1.5 = int5 / int1, mz1.6 = int6 / int1, mz2.1 = int7 / int1, mz2.2 = int8 / int1, mz2.3 = int9 / int1, mz2.4 = int10 / int1, mz2.5 = int11 / int1, mz2.6 = int12 / int1) %>% select(mz1.1, mz1.2, mz1.3, mz1.4, mz1.5, mz1.6, mz2.1, mz2.2, mz2.3, mz2.4, mz2.5, mz2.6) 

df_plot1 = df_plot %>% pivot_longer(c("mz1.1":"mz2.6"), names_to = "mz", values_to = "ratio")

write.xlsx(df_plot, file = paste(mainDir, subDir, "ratio_plot.xlsx", sep = "/"))

num1 = round(m, 4)
num2 = round(m + 1*(1.00335/z), 4)
num3 = round(m + 2*(1.00335/z), 4)
num4 = round(m + 3*(1.00335/z), 4)
num5 = round(m + 4*(1.00335/z), 4)
num6 = round(m + 5*(1.00335/z), 4)
num7 = round(m + diff/z, 4)
num8 = round(m + diff/z + 1*(1.00335/z), 4)
num9 = round(m + diff/z + 2*(1.00335/z), 4)
num10 = round(m + diff/z + 3*(1.00335/z), 4)
num11 = round(m + diff/z + 4*(1.00335/z), 4)
num12 = round(m + diff/z + 5*(1.00335/z), 4)

library(ggplot2)
if (nrow(df_plot1) == 0) {
  next
} else {
ggplot(df_plot1, aes(x=mz, y=ratio)) + 
  geom_jitter(position = position_jitter(0.2), color = "darkgray")+
  stat_summary(fun.data="mean_sdl", fun.args = list(mult=1), geom="crossbar", width=0.5) +
  ylim(0,2) +
  xlab("m/z") +
  ylab("relative intensity")+
  scale_x_discrete(breaks = c("mz1.1", "mz1.2", "mz1.3", "mz1.4", "mz1.5", "mz1.6", "mz2.1", "mz2.2", "mz2.3", "mz2.4", "mz2.5", "mz2.6"), labels = c(`num1`, `num2`, `num3`,`num4`,`num5`,`num6`,`num7`,`num8`,`num9`,`num10`,`num11`,`num12`))+
  theme_classic() +
  theme(axis.text.x = element_text(angle = 90)) +
  ggtitle(paste(plotname))
  
ggsave(filename = paste(mainDir, subDir, "mzratioPlot.pdf", sep = '/'))
}

tmp = df_org1 %>% select(rt) %>% inner_join(., df, by = "rt") 
rts = split(tmp, f = tmp$rt)
mx = length(rts)
rg_min = round(m - 5, 0)
rg_max = round(m + diff/z + 5, 0)

t_ms1 = c(rbind(df_org1$mz1, df_org1$mz2, df_org1$mz3, df_org1$mz4,df_org1$mz5,df_org1$mz6))
t_ms2 = c(rbind(df_org1$mz7, df_org1$mz8, df_org1$mz9, df_org1$mz10, df_org1$mz11, df_org1$mz12))

library(ggrepel)

if (mx == 0) {
  next
} else {
for (j in 1:mx)
{
  dfa = rts[[j]] %>% filter(mz > rg_min & mz < rg_max) %>% mutate(color = case_when(
    mz %in% t_ms1 ~ 'red',
    mz %in% t_ms2 ~ 'green',
    TRUE ~ 'black'
  ))
  standard_int = dfa %>% filter(!(color == 'black')) %>% select(int) %>% max(.)
  dfb = dfa %>% mutate(int_norm = int / standard_int * 100) %>% mutate(int_fi = ifelse(int_norm > 150, 150, int_norm)) %>% mutate(combo = paste(round(mz,4), round(int,0), sep = ", I:"))
  
  library(ggrepel)
  ggplot(dfb %>% filter(color == "black"))+
  geom_bar(aes(x=mz, y=int_fi), stat = "identity", color = "#000000", width = 0.01)+
  geom_bar(data = dfb %>% filter(color == "red"), aes(x=mz, y=int_fi), stat = "identity", color = "red", width = 0.01)+
  geom_bar(data = dfb %>% filter(color == "green"), aes(x=mz, y=int_fi), stat = "identity", color = "green", width = 0.01)+
  geom_label_repel(data = dfb %>% filter(color != "black"), aes(x=mz, y=int_fi, label = combo), na.rm = T, segment.color = "#C55A11", color = "#C55A11", segment.size = 0.1)+
  scale_x_continuous(expand = expansion(mult = c(0, 0)), breaks = seq(rg_min, rg_max, 5)) + 
  scale_y_continuous(expand = expansion(mult = c(0, 0)), breaks = seq(0, 150, 10))+
  coord_cartesian(xlim=c(rg_min, rg_max), ylim = c(0, 150))+
  theme_classic()+
  theme(legend.position = "none")+
  theme(axis.line = element_line(size = 0.1, colour = "black"))+
  xlab("m/z")+
  ylab("relative intensity")+
  ggtitle(paste(plotname, round(as.numeric(names(rts)[j]), digits = 4), "min", sep = "_"))
  
  filepath = paste(mainDir, subDir, sep = '/')
  ggsave(paste(filepath, "/scan_", j, ".pdf",sep=""))
  #dev.off()
}
}
}

```


```{r 6mergePDF}
library(qpdf)

# 1 ratioPlot
#pdfFiles <- dir("./results/individual", recursive=TRUE, full.names=TRUE, pattern="(mzratioPlot)+(.pdf)")
pdfFiles <- dir("./results/individual", recursive=TRUE, full.names=TRUE, pattern="(mzratioPlot)+(.pdf)")

n = length(pdfFiles)

if (n %% 500 == 0) {
  round = n %/%500
} else {
  round = n %/%500 + 1
}

for (r in 1:round) {
  left = (r-1)*500 + 1
  right = r*500
  if (right > n) {
    right = n
  }
  qpdf::pdf_combine(input = pdfFiles[left:right], output = paste("./output1_ratios", r, ".pdf", sep = ""))
}

# 2 first_ms1_example
ms1Files <- dir("./results/individual", recursive=TRUE, full.names=TRUE, pattern="(scan_1)+(.pdf)")

n = length(ms1Files)

if (n %% 500 == 0) {
  round = n %/%500
} else {
  round = n %/%500 + 1
}

for (r in 1:round) {
  left = (r-1)*500 + 1
  right = r*500
  if (right > n) {
    right = n
  }
  qpdf::pdf_combine(input = ms1Files[left:right], output = paste("./output4_ms1example", r, ".pdf", sep = ""))
}

# 3 Scans
ScanFiles <- dir("./results/individual", recursive=TRUE, full.names=TRUE, pattern="scan_\\d{1}|d{2}|d{3}\\.pdf") 

n = length(ScanFiles)

if (n %% 500 == 0) {
  round = n %/%500
} else {
  round = n %/%500 + 1
}

for (r in 1:round) {
  left = (r-1)*500 + 1
  right = r*500
  if (right > n) {
    right = n
  }
  qpdf::pdf_combine(input = ScanFiles[left:right], output = paste("./results/scans/output2_500scans", r, ".pdf", sep = ""))
}


# 4 ratio+scans
AllFiles <- dir("./results/individual", recursive=TRUE, full.names=TRUE, pattern="\\.pdf$")

n = length(AllFiles)

if (n %% 500 == 0) {
  round = n %/%500
} else {
  round = n %/%500 + 1
}

for (r in 1:round) {
  left = (r-1)*500 + 1
  right = r*500
  if (right > n) {
    right = n
  }
  qpdf::pdf_combine(input = AllFiles[left:right], output = paste("./results/combined/output3_500combined", r, ".pdf", sep = ""))
}

```


```{r 7pdf_statistics}
pdf = as_tibble(pdfFiles) 
library(dplyr)

tmp = pdf %>% separate(value, into = c(NA,NA,NA,"file",NA), sep = "[/]") %>% separate(file, into = c("pep1", "other"), sep = "_z_|_(?<=\\d)") %>% separate(other, into = c("z", "raw"), sep = "_", extra = "merge", remove = FALSE) %>% select(-other) %>% mutate(MS1pair = "Yes") %>% mutate(z = as.numeric(z)) %>% mutate(raw = sub("_row_\\d+$", "",raw)) %>% unique (.)

final = left_join(source_temp4.2, tmp, by = c("raw", "pep1", "z.x" = "z")) #%>% mutate(MS1pair = replace_na(MS1pair, "No")) 
#write.xlsx(final, "MS1pair.xlsx")

tmp2 = pdf %>% separate(value, into = c(NA,NA,NA,"file",NA), sep = "[/]") %>% mutate(f0 = "./results/individual/", f8 = "/ratios.xlsx") %>% mutate(excel = paste0(f0, file, f8))
len1 = nrow(tmp2)
lst = vector('list', len1)

for (i in 1:len1) {
  data = read_excel(tmp2$excel[i]) 
  ratio1_avg = list()
  ratio2_avg = list()
  ratio3_avg = list()
  ratio4_avg = list()
  ratio5_avg = list()
  ratio1_avg = mean(data$ratio1, na.rm = TRUE)
  ratio2_avg = mean(data$ratio2, na.rm = TRUE)
  ratio3_avg = mean(data$ratio3, na.rm = TRUE)
  ratio4_avg = mean(data$ratio4, na.rm = TRUE)
  ratio5_avg = mean(data$ratio5, na.rm = TRUE)
  ratio1_sd = list()
  ratio2_sd = list()
  ratio3_sd = list()
  ratio4_sd = list()
  ratio5_sd = list()
  ratio1_sd = sd(data$ratio1, na.rm = TRUE)
  ratio2_sd = sd(data$ratio2, na.rm = TRUE)
  ratio3_sd = sd(data$ratio3, na.rm = TRUE)
  ratio4_sd = sd(data$ratio4, na.rm = TRUE)
  ratio5_sd = sd(data$ratio5, na.rm = TRUE)
  ratio1_number = list()
  ratio2_number = list()
  ratio3_number = list()
  ratio4_number = list()
  ratio5_number = list()
  ratio1_number = data$nRatio1[1]
  ratio2_number = data$nRatio2[1]
  ratio3_number = data$nRatio3[1]
  ratio4_number = data$nRatio4[1]
  ratio5_number = data$nRatio5[1]
  lst[[i]] = data.frame(tmp2$excel[i], ratio1_avg, ratio1_sd, ratio1_number, ratio2_avg, ratio2_sd, ratio2_number, ratio3_avg, ratio3_sd, ratio3_number, ratio4_avg, ratio4_sd, ratio4_number, ratio5_avg, ratio5_sd, ratio5_number)
}

df_ratio <- do.call("rbind", lst) %>% separate(tmp2.excel.i., into = c(NA,NA,NA,"file",NA), sep = "[/]") %>% separate(file, into = c("pep1", "other"), sep = "_z_|_(?<=\\d)") %>% separate(other, into = c("z", "raw"), sep = "_", extra = "merge", remove = FALSE) %>% select(-other) %>% mutate(z = as.numeric(z)) %>% mutate(raw = sub("_row_\\d+$", "",raw))
#df_ratio$raw = gsub("_row", "", df_ratio$raw)
#write.xlsx(df_ratio, "ratio_summary.xlsx")

#df_ratio1 = df_ratio %>% select("ratioAverage","sd","number","pep1","z.x","raw") %>% unique(.)
sum_temp1 = left_join(final, df_ratio, by = c("pep1", "z.x"= "z", "raw")) %>% mutate(HL_MS2 = "Yes") %>% unique(.)
colnames(sum_temp1) <-  c("Protein", "peptide_unmod","StartingPosition", "Peptide.H","Peptide.L","Modification.H","Modification.L","z","raw","maxScoreH","maxScoreL","MS1pair","HL_ratio_M","sd_ratio_M","n_M","HL_ratio_M+1","sd_M+1","n_M+1","HL_ratio_M+2","sd_M+2","n_M+2","HL_ratio_quad","sd_quad","n_quad","HL_ratio_sextet","sd_sextet","n_sextet", "HL_MS2") 

sum_temp2 = sum_temp1 %>% select("Protein", "peptide_unmod","StartingPosition", "Peptide.H","Peptide.L","z","raw","maxScoreH","maxScoreL", "HL_MS2", "MS1pair","HL_ratio_M","HL_ratio_M+1","HL_ratio_M+2","HL_ratio_quad","HL_ratio_sextet","n_M","n_M+1","n_M+2","n_quad","n_sextet","sd_ratio_M","sd_M+1","sd_M+2","sd_quad","sd_sextet") %>% unique(.)


write.xlsx(sum_temp2, "output_summary.xlsx", keepNA = TRUE, na.string = "NA")

```

```{r 8check}

library(tidyverse)
library(readxl)
library(openxlsx)
library(stringr)
library(dplyr)
library(magrittr)
library(tidyr)
files <- dir(".", recursive=TRUE, full.names=TRUE, pattern="\\output_summary.xlsx$")

result <- files %>% lapply(read_excel, sheet = 1, col_types = "text") %>% bind_rows %>% mutate(maxScoreH = as.numeric(maxScoreH)) %>% mutate(maxScoreL = as.numeric(maxScoreL))
result <- files %>%
  lapply(read_excel, sheet = 1, col_types = "text") %>%
  bind_rows() %>%
  mutate(maxScoreH = as.numeric(maxScoreH), maxScoreL = as.numeric(maxScoreL), HL_ratio_M = as.numeric(HL_ratio_M)) %>%
  filter(!(maxScoreH < 300 & maxScoreL < 300), !(HL_ratio_M > 6), MS1pair != "No", !grepl("Contaminant|TRYP|Keratin|ATE1|SYRC", Protein)) %>% mutate(values = str_extract(Protein, "(?<=\\|)[A-Za-z0-9_]+(?=_)"), site = str_extract(peptide_unmod, "^[A-Za-z]"))

result$Numbers <- str_extract(result$Peptide.L, "\\d+\\.\\d+")
result <- result %>%
  mutate(Numbers = case_when(
    Numbers == "156.101" ~ "R",
    Numbers == "204.086" ~ "RO3",
    Numbers == "157.085" ~ "R_deami",
    Numbers == "188.091" ~ "RO2",
    TRUE ~ as.character(Numbers)
  ))
result1 <- result %>% mutate(symbol = values, AA = site, StartingPosition1 = StartingPosition, mod_type = Numbers) %>% unite(protein_site_mod, values, site, StartingPosition, Numbers, sep = " ") %>% mutate(protein_site_mod = str_replace(protein_site_mod, "(\\S+)\\s(\\S+)(\\S+)", "\\1\\2\\3"))

result2 <- result1 %>% mutate(symbol1 = symbol, AA1 = AA, StartingPosition2 = StartingPosition1) %>% unite(protein_site, symbol, AA, StartingPosition1, sep = " ") %>% mutate(protein_site = str_replace(protein_site, "(\\S+)\\s(\\S+)(\\S+)", "\\1\\2\\3"))

result3 <- result2 %>% mutate(peptide_unmod1 = peptide_unmod, mod_type1 = mod_type) %>% unite(R_sequence, mod_type, peptide_unmod, sep = "_") %>% mutate(R_sequence = str_replace_all(R_sequence, "_", "")) 

colnames(result3) <-  c("Protein","R_sequence","protein_site_mod", "Peptide.H", "Peptide.L", "z", "raw", "maxScoreH", "maxScoreL", "HL_MS2", "MS1pair","HL_ratio_M", "HL_ratio_M+1", "HL_ratio_M+2", "HL_ratio_quad", "HL_ratio_sextet", "n_M", "n_M+1", "n_M+2",  "n_quad", "n_sextet",  "sd_ratio_M","sd_M+1", "sd_M+2", "sd_quad","sd_sextet","protein_site", "symbol","amino acid", "StartingPosition", "peptide_unmod", "mod_type")
result3 <- result3 %>% arrange(protein_site_mod)
#generate summary table
write.xlsx(result3, "1summary_sites.xlsx")
data_read <- read_excel("1summary_sites.xlsx") %>% select("protein_site_mod") %>% unique(.)
#generate information for website
data_website <- read_excel("1summary_sites.xlsx") %>% mutate(z = as.double(z)) %>% unique(.)


source_final <- read_excel("HLmatch_3min.xlsx") %>% unique(.)
data_website1 = left_join (data_website, source_final, by = c("peptide_unmod"="sub_pep", "z" ="Z", "raw"="raw")) %>% distinct(R_sequence, z, raw, .keep_all = TRUE)
#write.xlsx(data_website1, "1website_information.xlsx")
#data_read <- read_excel("1website_information.xlsx")
data_web <- data_website1 %>% mutate(Sample = ifelse(grepl("lin_1_29_4and5_pair", raw), "Standard_peptide", NA))
write.xlsx(data_web, "1website_information.xlsx")
```

```{r, 9extract MS1, MS2H, MS2L}
library(mzR)
#MS2 heavy scans
match_heavy1 = match_heavy %>% separate(Comment, into=c("raw","other"), sep = "[.]", remove = FALSE, extra = "drop") %>% mutate(z = as.numeric(z), Calcmass = as.character(Calcmass)) %>% mutate(scan_no = as.numeric(gsub(".*scan=", "", Scannum)))
match_heavy1$Peptide <- gsub("U", "\\.\\-", match_heavy1$Peptide) #

result31 = result3 %>% mutate(z = as.numeric(z))
heavy_ms2list = inner_join(match_heavy1, result31, by = c("Peptide"="Peptide.H", "z" ="z", "raw"="raw")) %>% select(raw, Peptide, R_sequence, z, scan_no, ScanTime, Calcmass)%>% unique(.) # %>% mutate(Calcmass = as.numeric(Calcmass)) 

total_row = nrow(heavy_ms2list)
mainDir = './website'

for (i in 1:total_row) # check rows.
{
subDir = paste(heavy_ms2list$raw[i], heavy_ms2list$z[i], heavy_ms2list$R_sequence[i], sep = '_')
plotname = paste(heavy_ms2list$raw[i], heavy_ms2list$z[i], heavy_ms2list$R_sequence[i],heavy_ms2list$scan_no[i], sep = '_')
dir.create(file.path(mainDir, subDir))

raw = paste(heavy_ms2list$raw[i], '.mzXML', sep = '')
msdata <- grabMSdata(files = paste('./mzxml/',raw, sep = ''), grab_what = c("MS2"))
msdata_org = msdata$MS2 %>% select(rt, fragmz, int) %>% mutate(rt = round(rt, digits = 4))
rti = round(heavy_ms2list$ScanTime[i], 4)
ms2data = msdata_org %>% filter(rt == rti)
write.xlsx(ms2data, file = paste(paste(mainDir, subDir, sep = "/"), '/ms2H_', plotname, '.xlsx', sep = ''))

}

#MS2 light scans
match_light1 = match_light %>% separate(Comment, into=c("raw","other"), sep = "[.]", remove = FALSE, extra = "drop") %>% mutate(z = as.numeric(z), Calcmass = as.character(Calcmass)) %>% mutate(scan_no = as.numeric(gsub(".*scan=", "", Scannum)))
match_light1$Peptide <- gsub("U", "\\.\\-", match_light1$Peptide)

result31 = result3 %>% mutate(z = as.numeric(z))
light_ms2list = inner_join(match_light1, result31, by = c("Peptide"="Peptide.L", "z" ="z", "raw"="raw")) %>% select(raw, Peptide, R_sequence, z, scan_no, ScanTime, Calcmass) %>% unique(.)

total_row = nrow(light_ms2list)
mainDir = './website'

for (i in 1:total_row) # check rows.
{
subDir = paste(light_ms2list$raw[i], light_ms2list$z[i], light_ms2list$R_sequence[i], sep = '_')
plotname = paste(light_ms2list$raw[i], light_ms2list$z[i], light_ms2list$R_sequence[i],light_ms2list$scan_no[i], sep = '_')
dir.create(file.path(mainDir, subDir))

raw = paste(light_ms2list$raw[i], '.mzXML', sep = '')
msdata <- grabMSdata(files = paste('./mzxml/',raw, sep = ''), grab_what = c("MS2"))
msdata_org = msdata$MS2 %>% select(rt, fragmz, int) %>% mutate(rt = round(rt, digits = 4))
rti = round(light_ms2list$ScanTime[i], 4)
ms2data = msdata_org %>% filter(rt == rti)
write.xlsx(ms2data, file = paste(paste(mainDir, subDir, sep = "/"), '/ms2L_', plotname, '.xlsx', sep = ''))

}
```

```{r 10 ms1 scans}
# MS1 scans

source_final_rt = source_final %>% mutate(Calcmass.y = as.character(Calcmass.y)) 
light_ms2list_rt = inner_join (light_ms2list, source_final_rt, by = c("Calcmass"="Calcmass.y", "z" ="Z", "raw"="raw")) %>% select (raw,Peptide,R_sequence,z,Calcmass,sub_pep, ScanTimeMean) %>% unique(.)
light_ms2list2 = light_ms2list_rt %>% select (raw, R_sequence, z, Calcmass, ScanTimeMean) %>% unique(.)

total_row = nrow(light_ms2list2)

for (i in 1:total_row) # check total_row.
{
raw = paste(light_ms2list2$raw[i], '.mzXML', sep = '')
subDir = paste(light_ms2list2$raw[i], light_ms2list2$z[i], light_ms2list2$R_sequence[i], sep = '_')
msdata <- grabMSdata(files = paste('./mzxml/',raw, sep = ''), grab_what = c("MS1"))
calculatedmass = as.numeric(light_ms2list2$Calcmass[i])# [M+H]+
z = light_ms2list2$z[i]
m = (calculatedmass - 1.00728 + z*1.00728)/z # L m/z
retentiontime =  light_ms2list2$ScanTimeMean[i]

diff = 10.008269

df = msdata$MS1 %>% filter(abs(rt-retentiontime) < 1.25) %>% select(-filename) 

df1 = df %>% filter(abs((mz - m)/m*10^6) <= 10) 
df7 = df %>% filter(abs((mz - (m + diff/z))/(m + diff/z)*10^6) <= 10) 

df_org1 = inner_join(df1, df7, by = c("rt")) 
colnames(df_org1) <- c("rt", "mz1", "int1", "mz7", "int7") 
tmp = df_org1 %>% select(rt) %>% inner_join(., df, by = "rt") 
rts = split(tmp, f = tmp$rt)
mx = length(rts)
rg_min = round(m - 5, 0)
rg_max = round(m + diff/z + 5, 0)

for (j in 1:mx) #mx
{
  dfa = rts[[j]] %>% filter(mz > rg_min & mz < rg_max)
  write.xlsx(dfa, file = paste(paste(mainDir, subDir, sep = "/"), '/ms1_', subDir, '_', round(rts[[j]]$rt[1], 4), '.xlsx', sep = ''))
}

}


```
````
