## Supplemental Dataset 1 for "An Unbiased Proteomic Platform for ATE1-based Arginylation Profiling": instructions to users.docx

**Instructions to run ArginylomePlot**

1. Install R to your computer.
2. Install RStudio to your computer.
3. Un-zip Supplementary dataset 1.
4. If you run the software for the first time, please ignore this step. Subfolders in “results” and folder “website” are empty by default. Before re-run “ArginylomePlot_v12”,
   4.1 Empty “combined”, “individual”, “scans”, and “website” folders.
   4.2 Delete Excel and PDF files in the dataset folder.
5. Open “ArginylomePlot_v12.rmd” by RStudio.
6. Click the green “run” button 10 times for each of the 10 R markdown sections (see below for section 1 as an example). User can click all 10 in order when the software is still running, or click next after previous section is done.

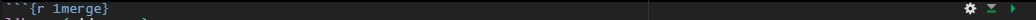

7. Upon completion, exported results on Heavy/light ratios and MS1 scans of each peptide with unique charge and raw are displayed in the “individual” folder. Paired MS1 and MS2 scans in excel format are displayed in the "website” folder for arginylation database indexing.

Note: information in the “website” folder is the same as downloaded from [www.arginylation.com](http://www.arginylation.com).
