## Supplemental Dataset 4 for "An Unbiased Proteomic Platform for ATE1-based Arginylation Profiling"

**Supplementary dataset 4**. **Representative MS2 spectra of 235 arginylation sites modified by Arg10 and Arg0 in human proteomes.** N-terminal arginylation (R: +156 Da), midchain arginylation (R: +156 Da) on D and E, Cys tri-oxidation and arginylation (RO3: +204 Da), Cys di-oxidation and arginylation (RO2: +188 Da), N/Q arginylation after deamidation (R_deami: +157 Da).

Spectra and sites are displayed in **alphabetic order** of protein symbols.

For all MS1 and MS2 spectra visualization and download, please visit [www.arginylation.com](http://www.arginylation.com).

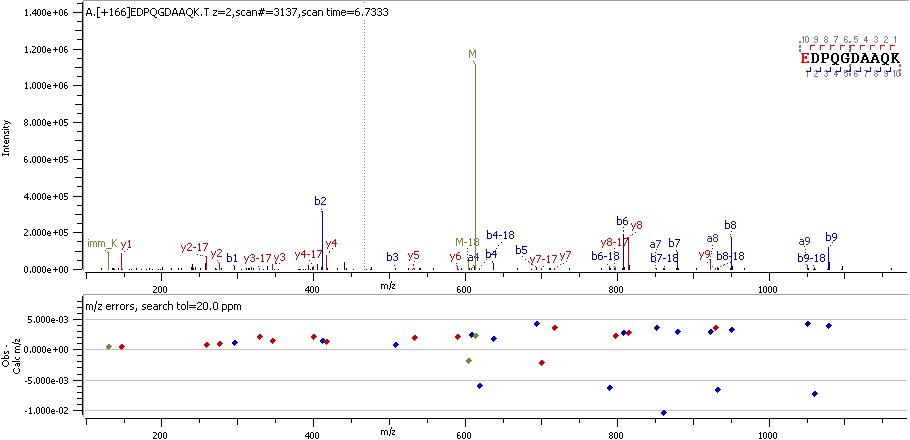

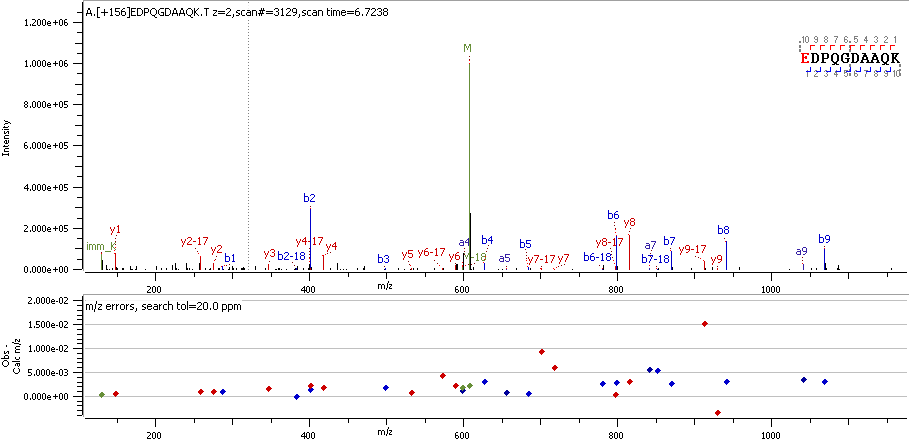

A1AT E25 R

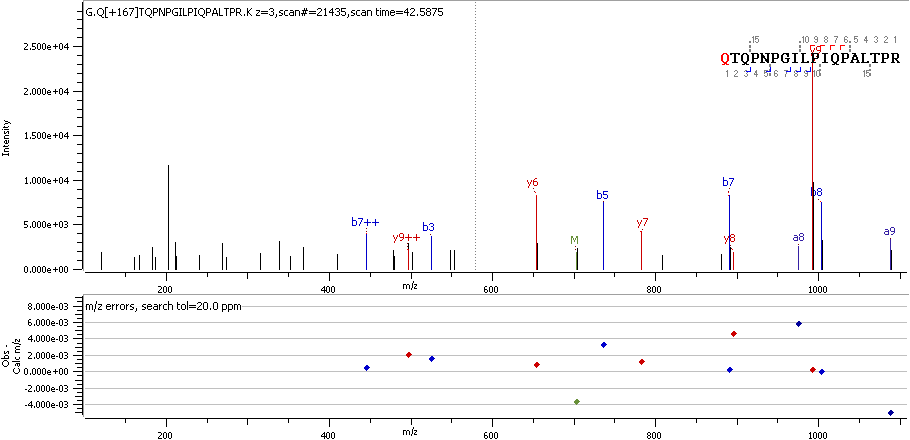

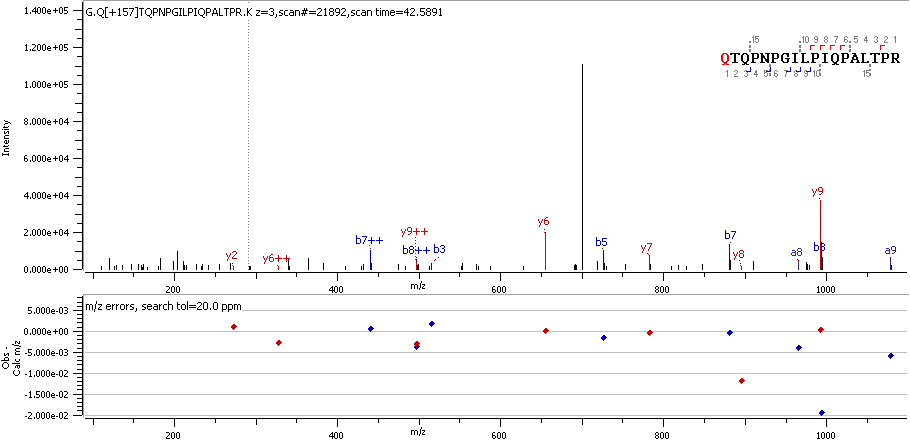

AAK1 Q374 R_deami

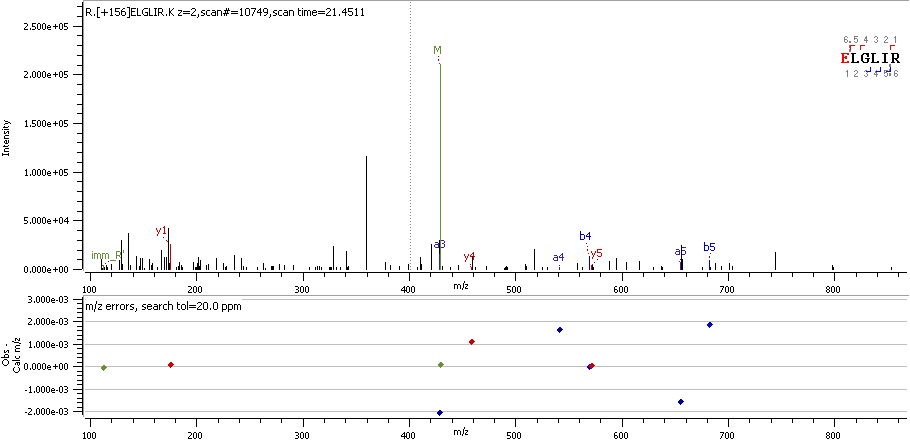

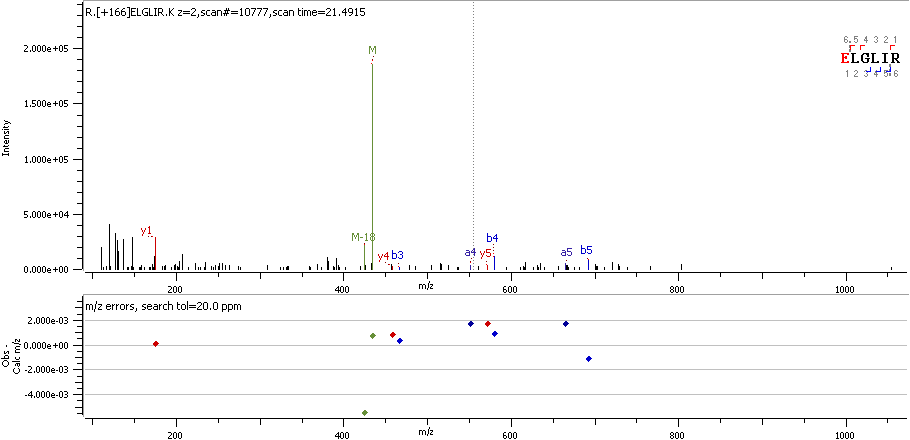

ACLY E830 R

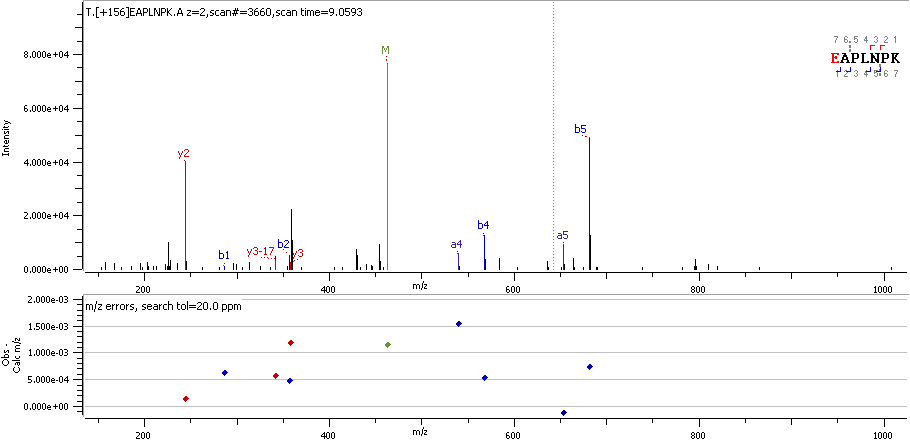

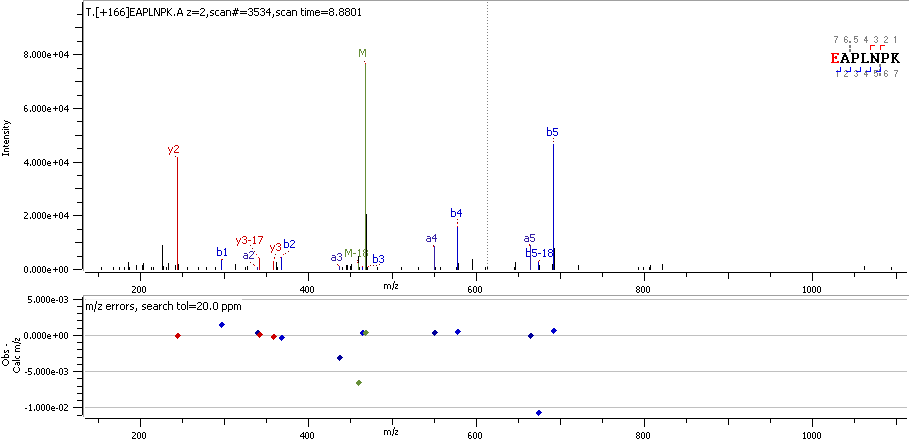

ACTA E109 R

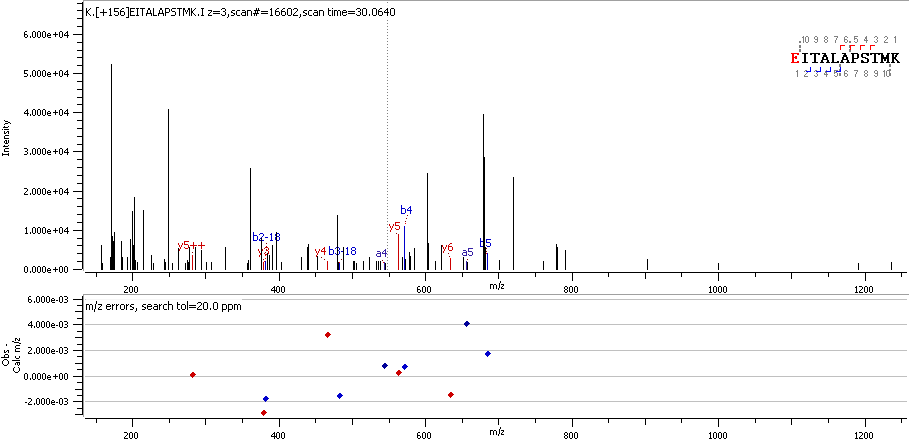

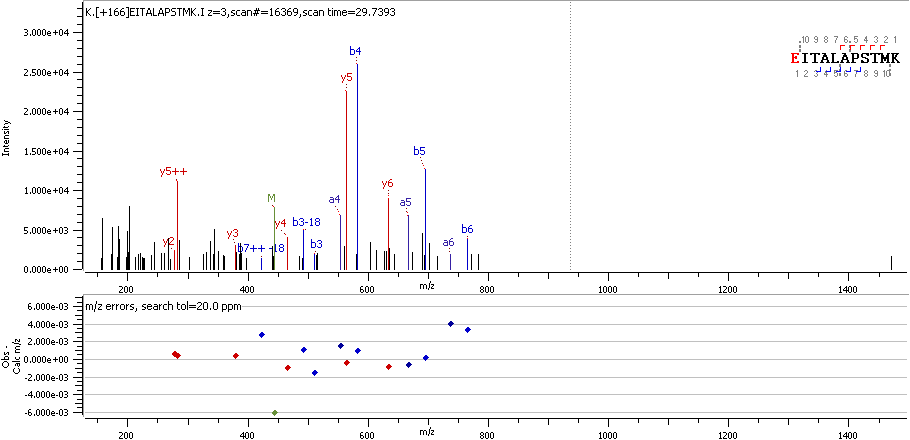

ACTA E318 R

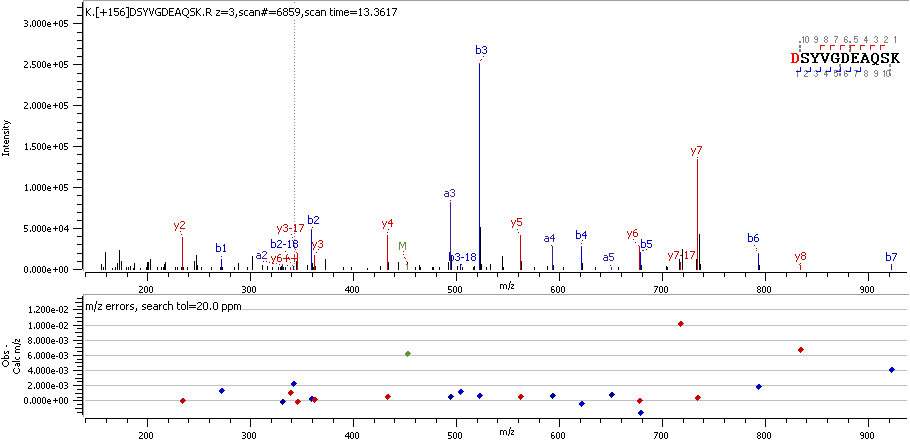

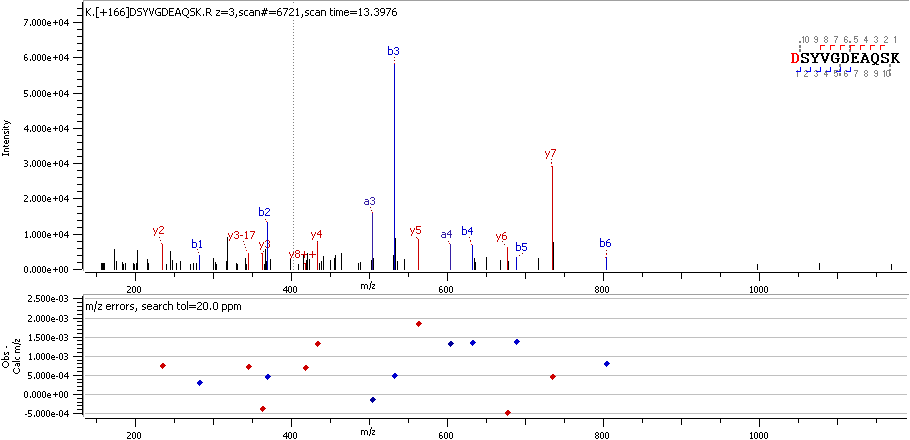

ACTB D51 R

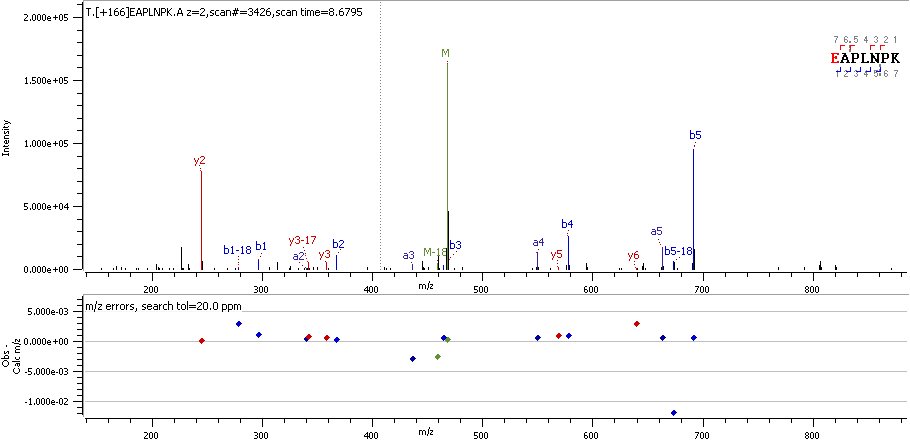

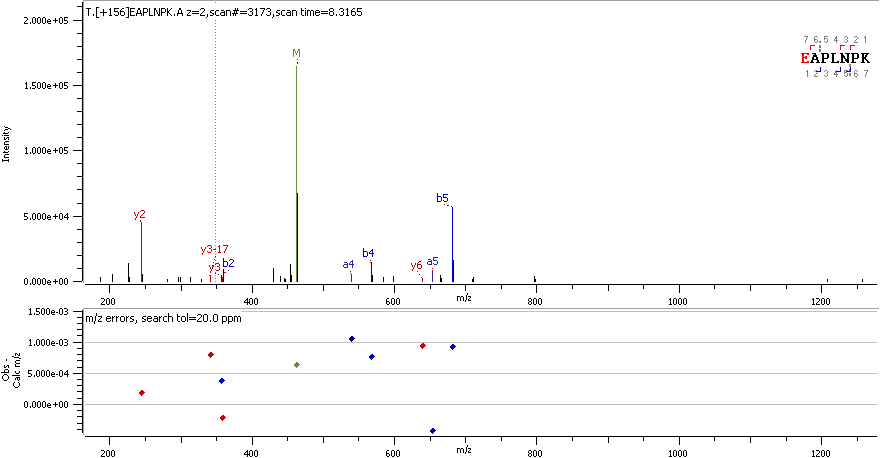

ACTB E107 R

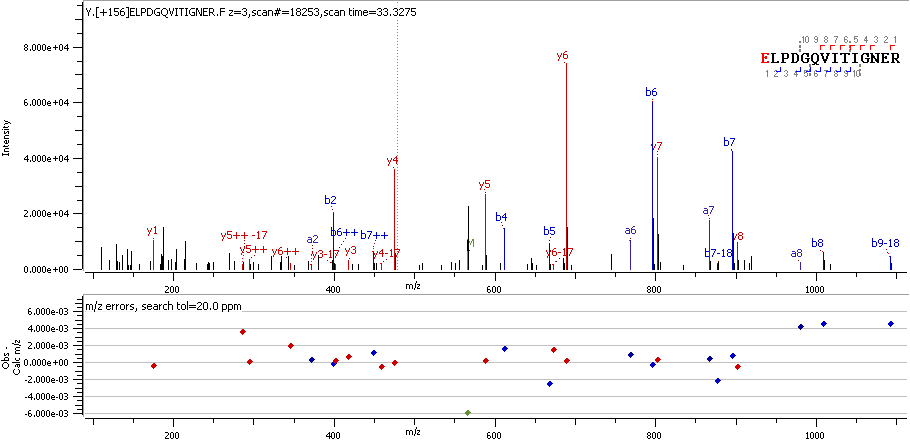

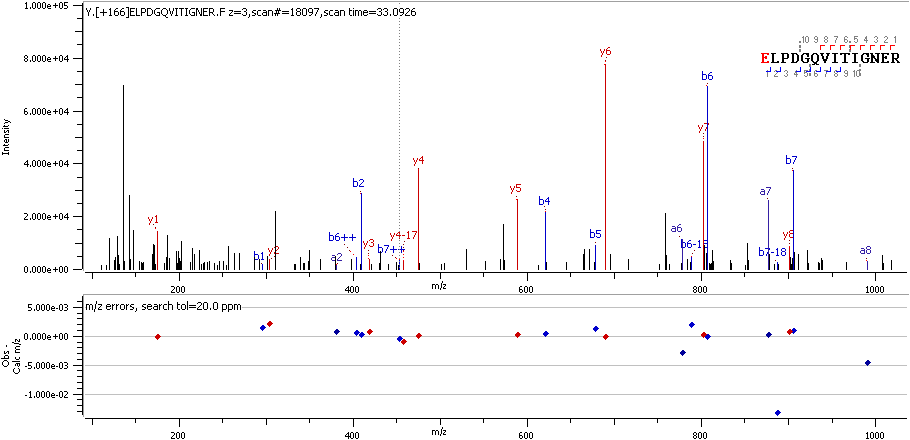

ACTB E241 R

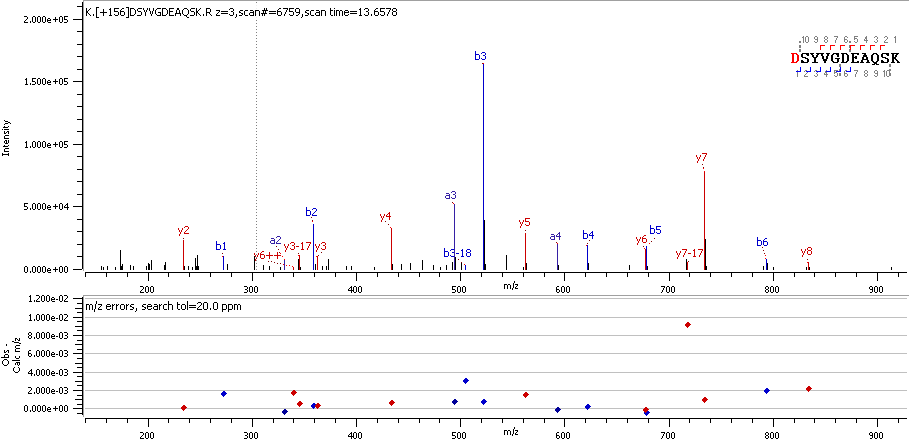

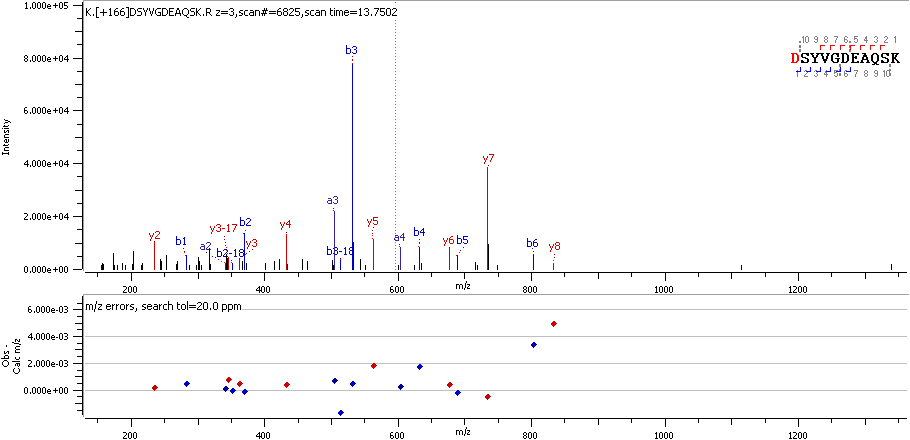

ACTC D53 R

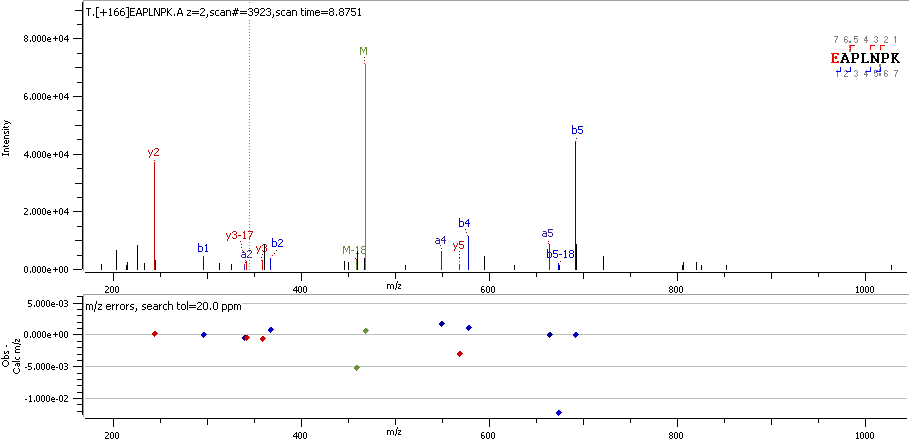

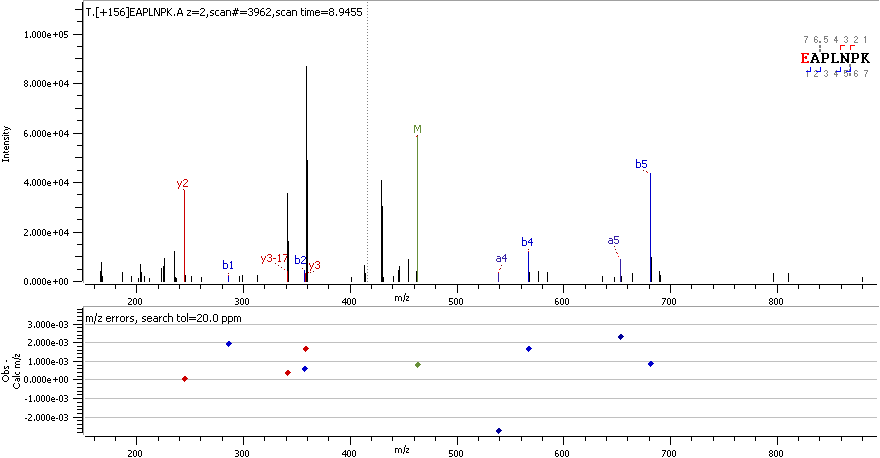

ACTC E109 R

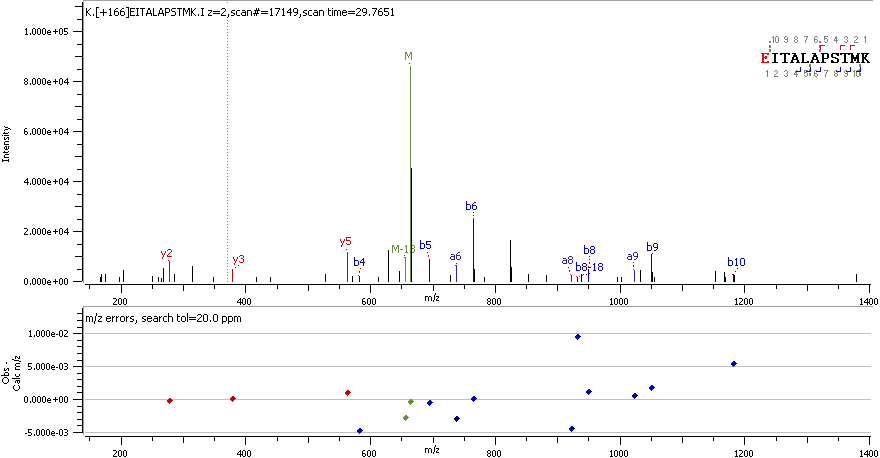

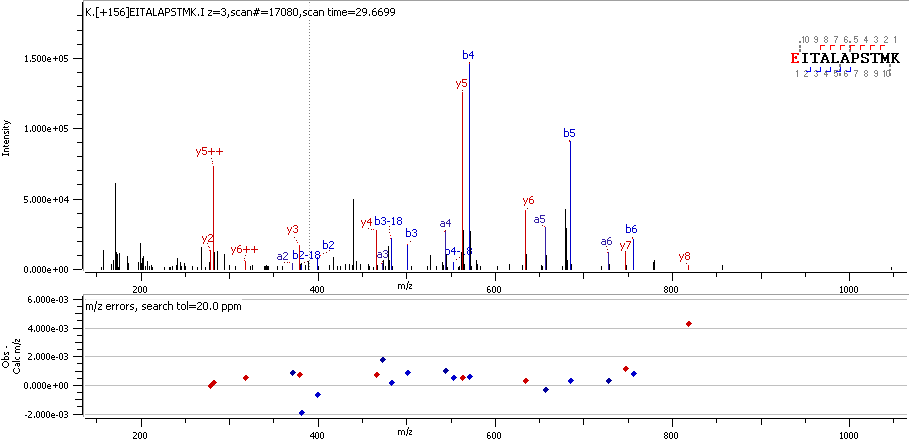

ACTC E318 R

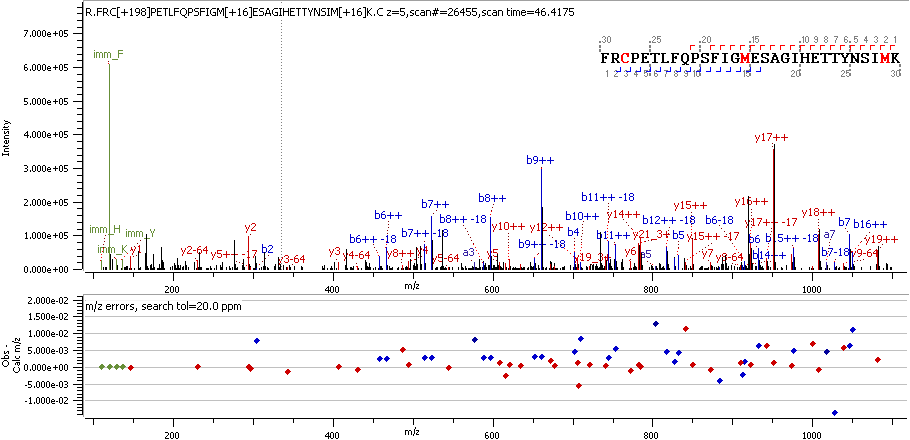

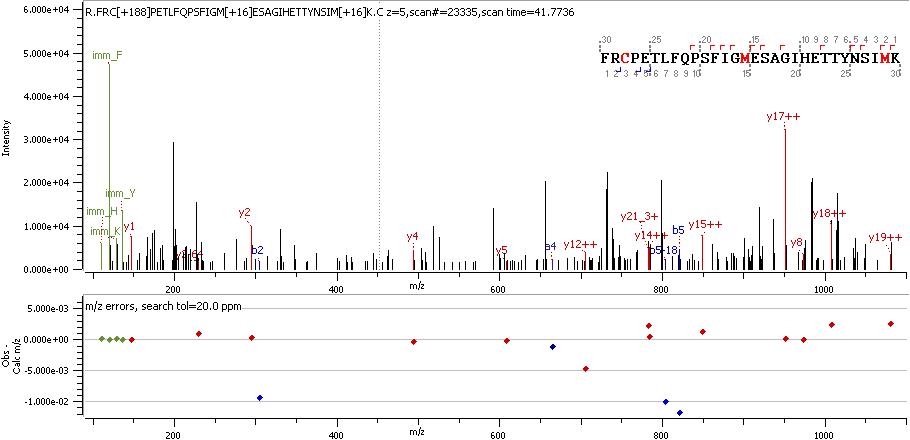
ACTC C259 RO2

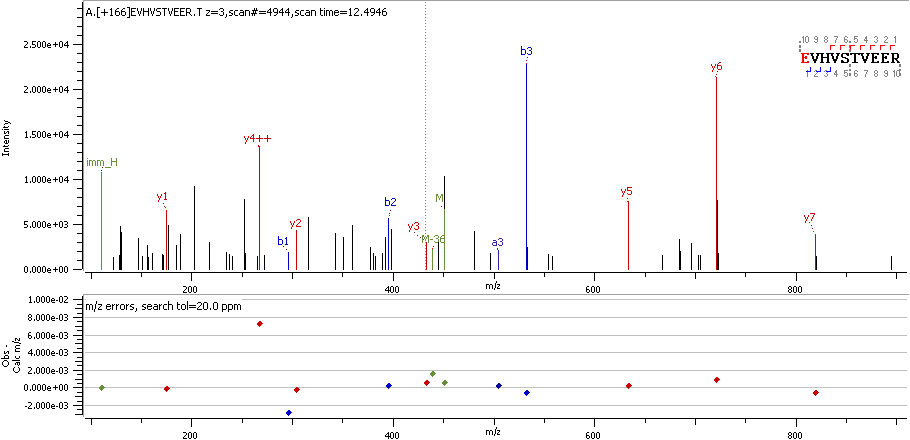

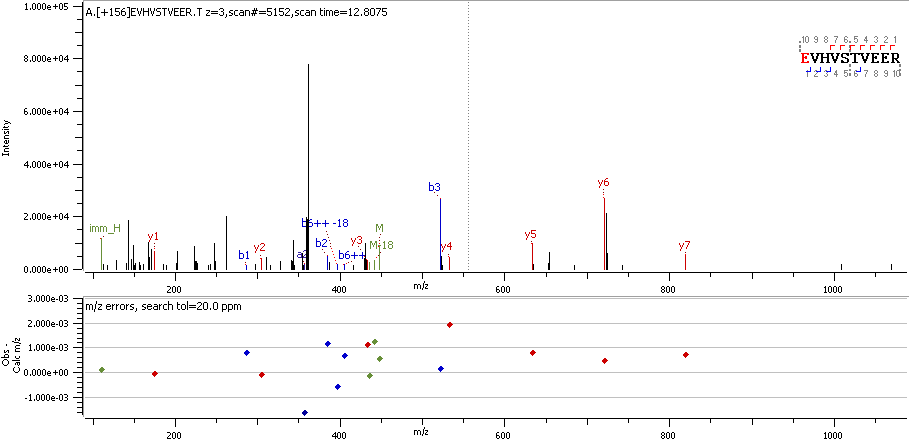

AKA12 E418 R

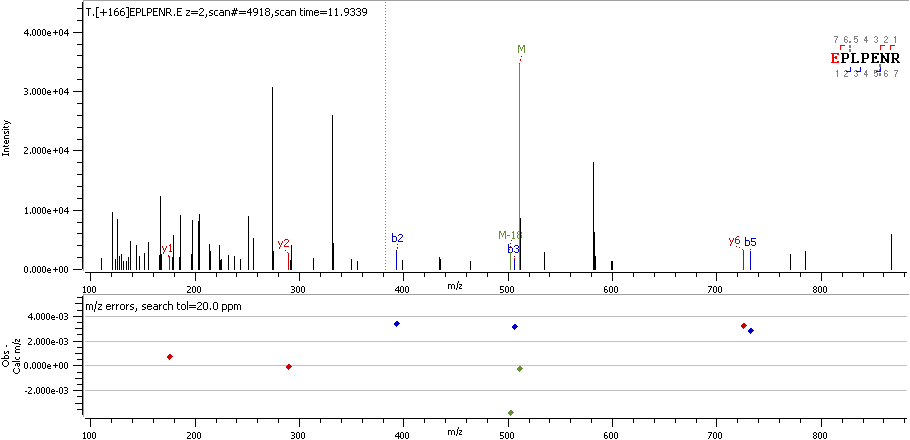

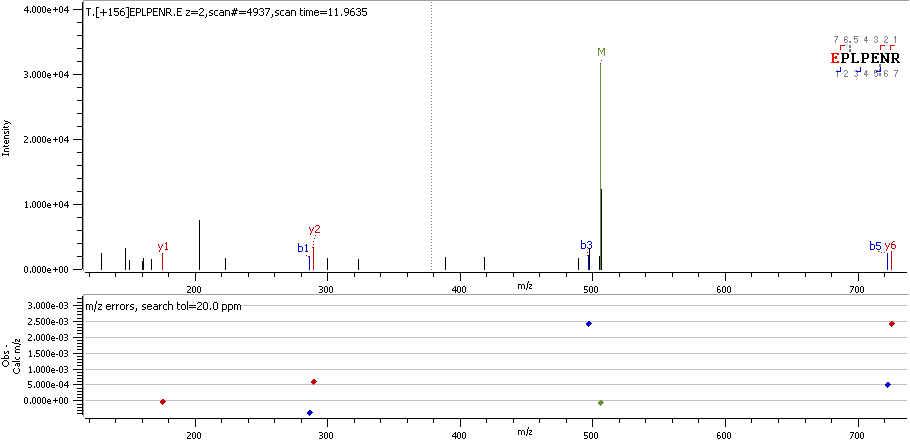

AKA12 E952 R

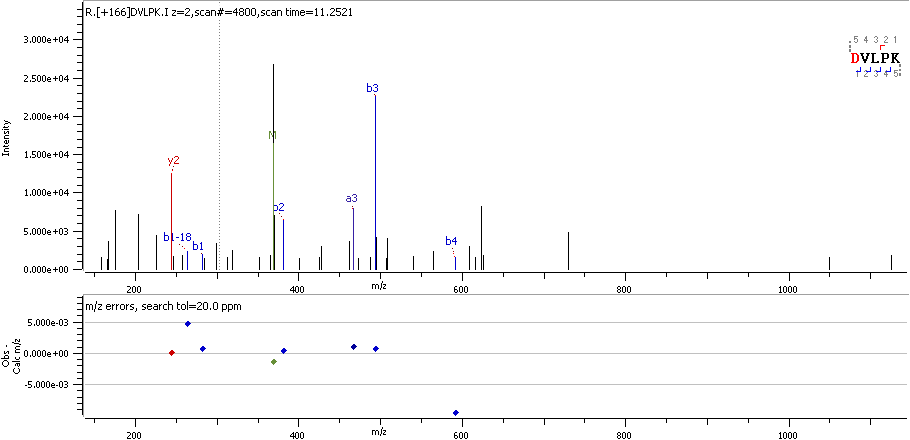

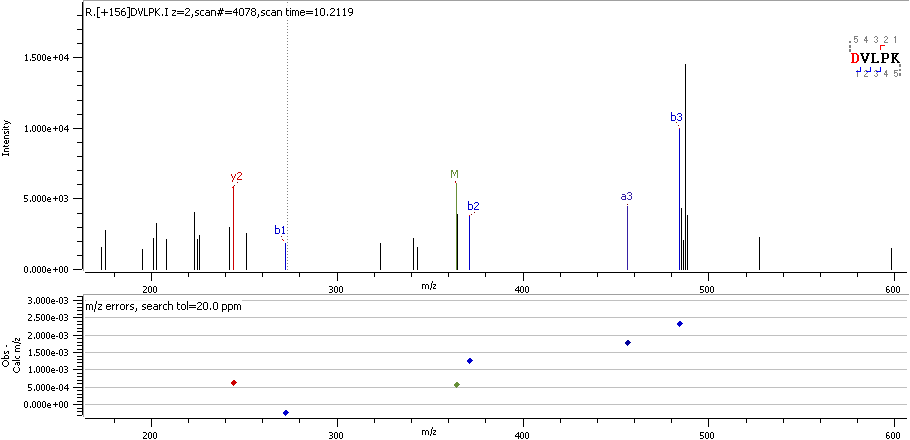

AKAP2 D638 R

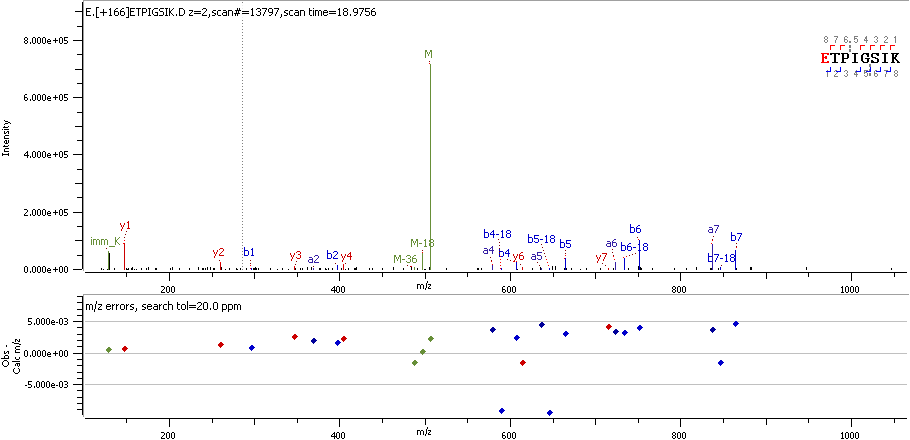

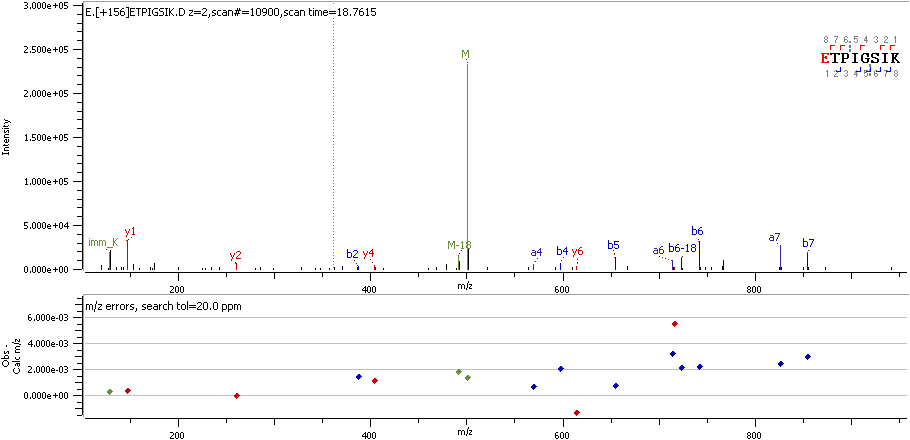

ANK2 E1760 R

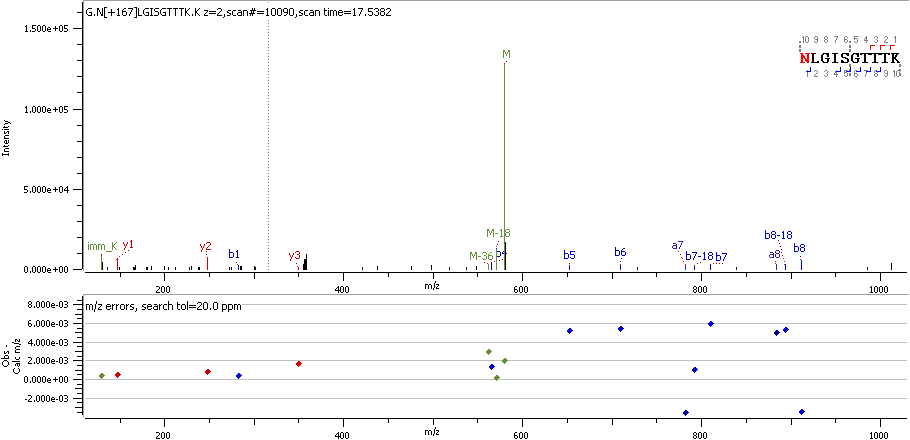

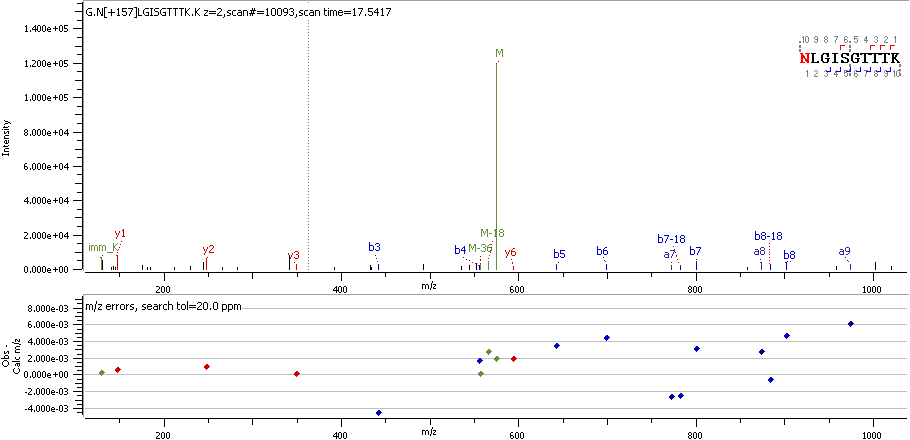

AP180 N767 R_deami

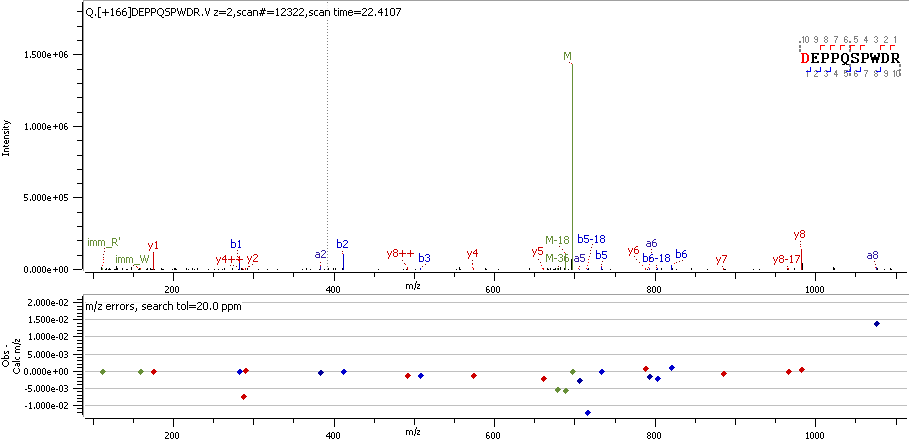

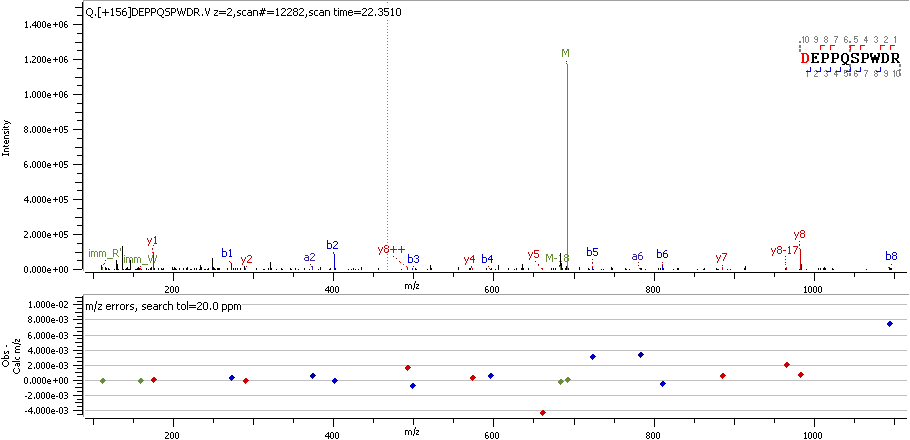

APOA1 D25 R

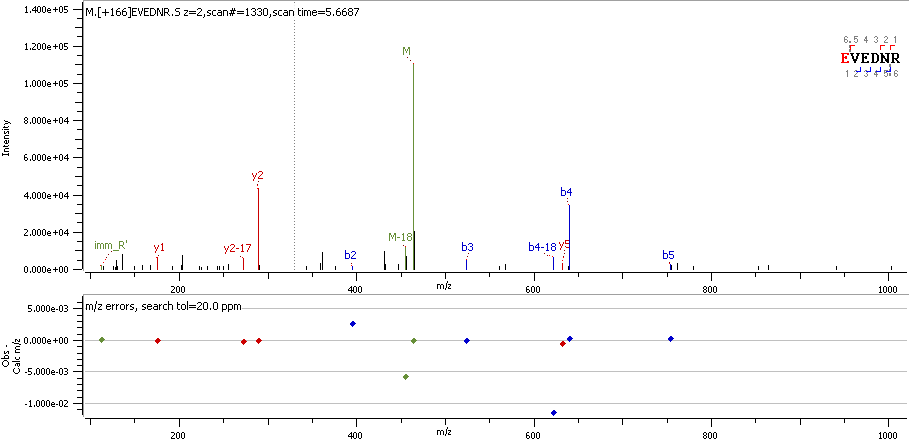

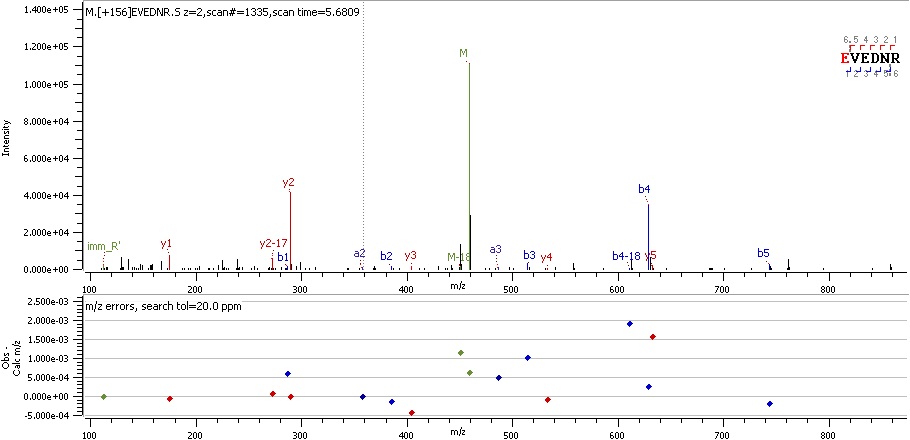

AQP4 E279 R

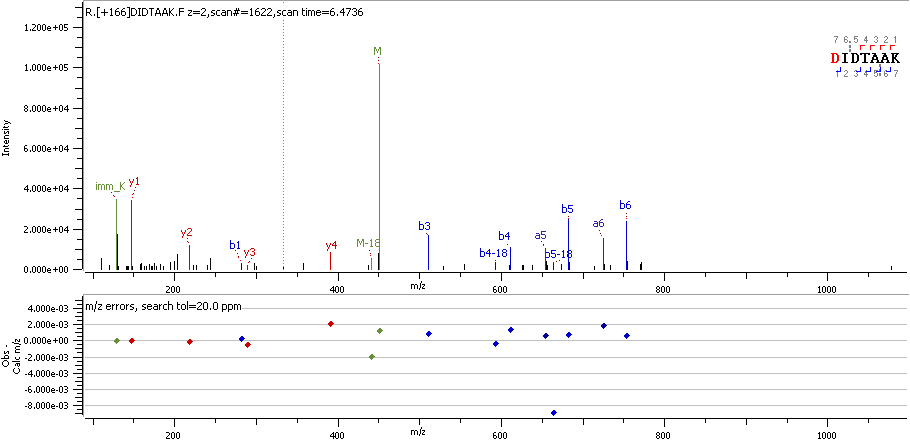

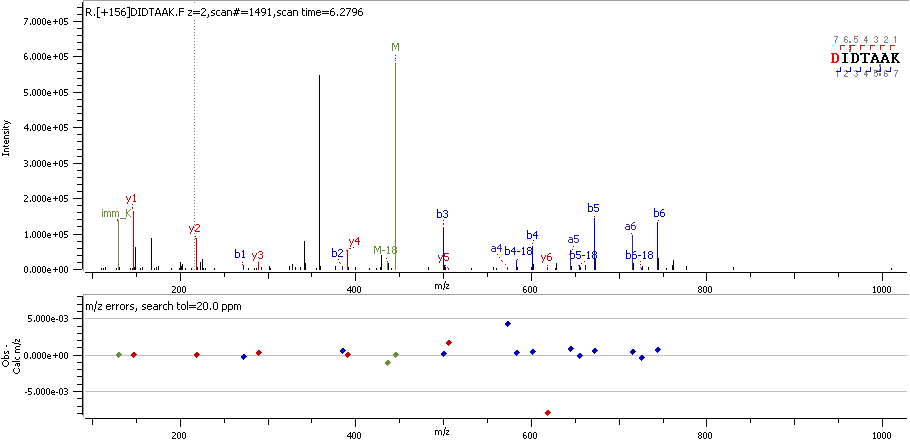

AT5G1 D62 R

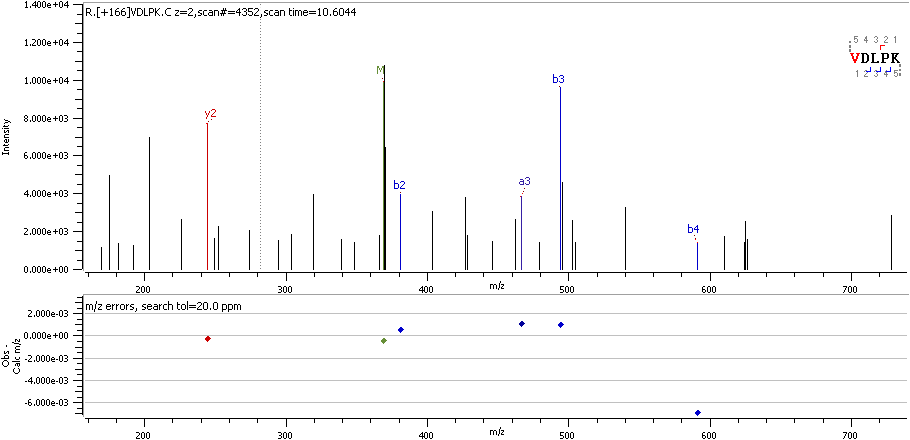

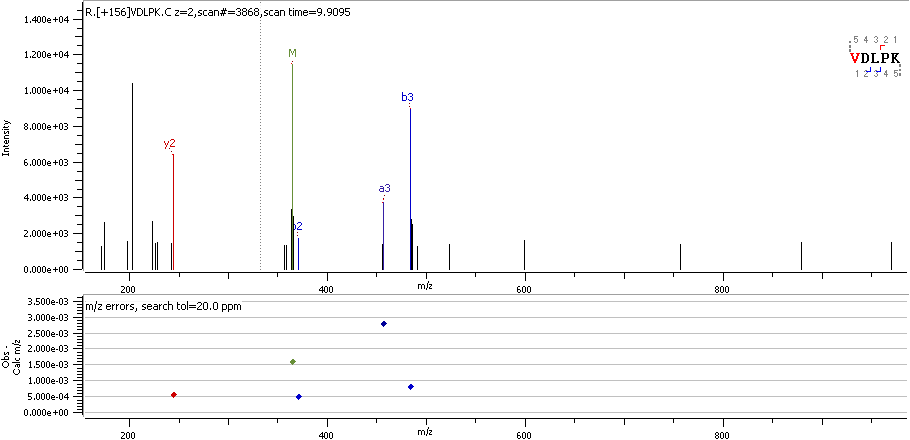

ATAD5 V1449 R

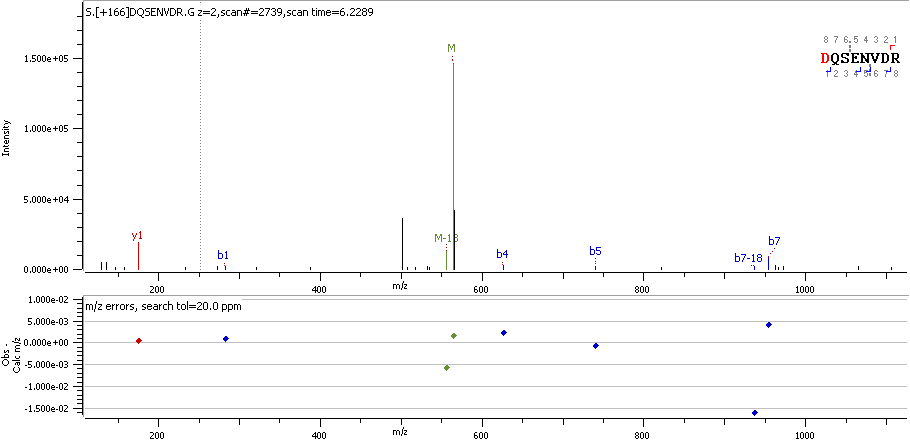

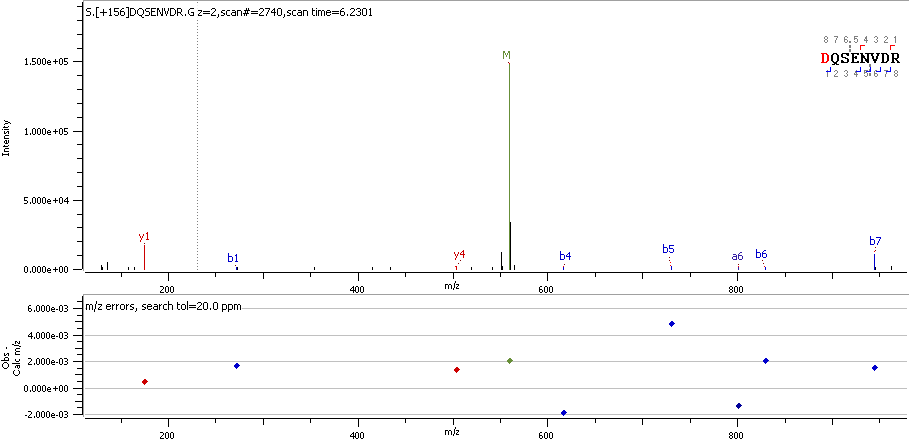

ATIF1 D28 R

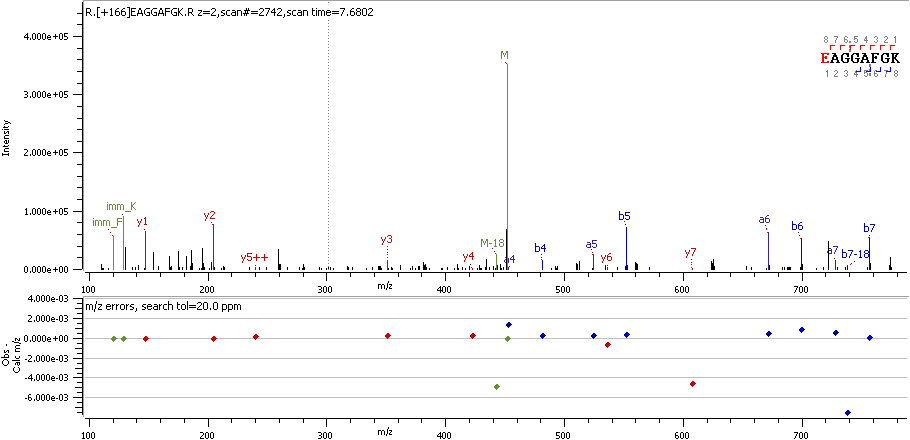

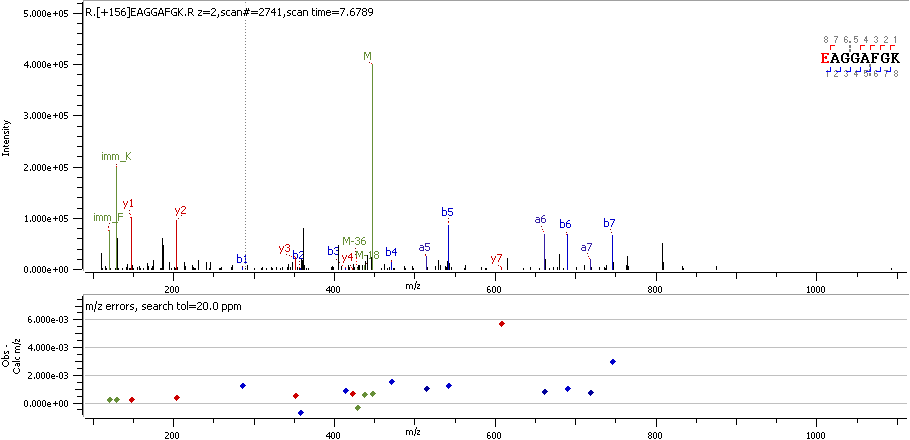

ATIF1 E42 R

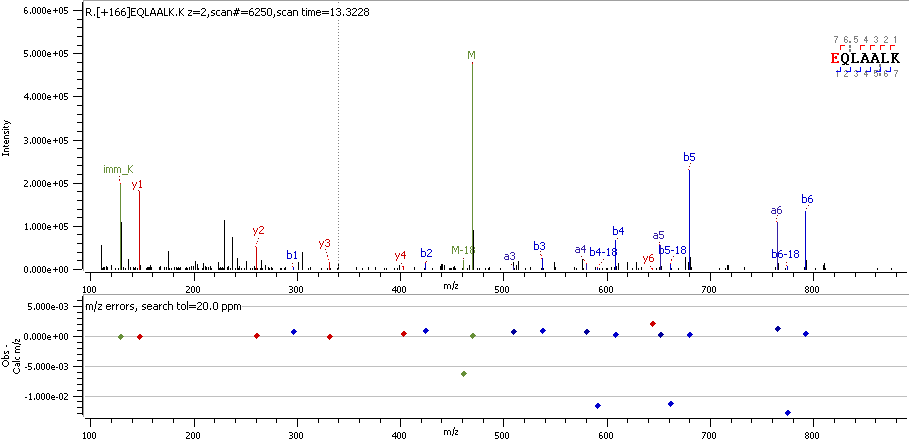

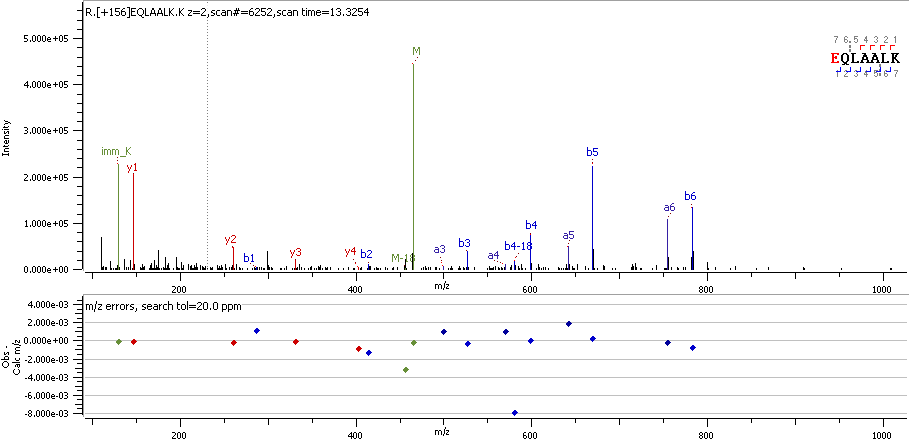

ATIF1 E65 R

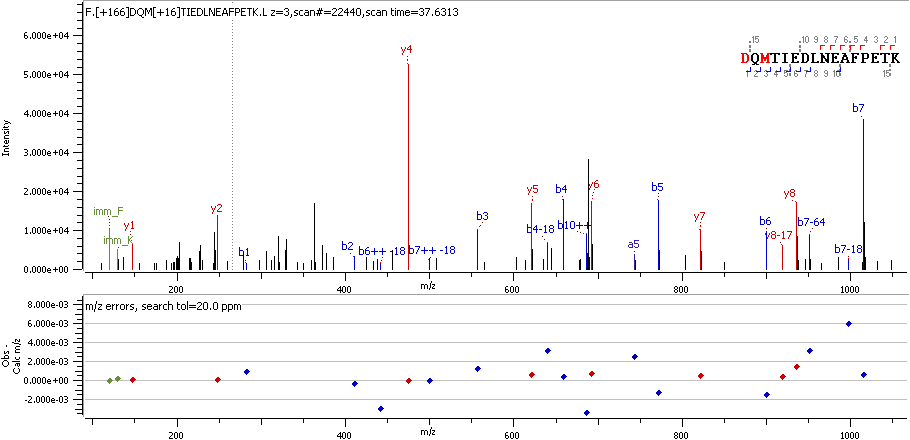

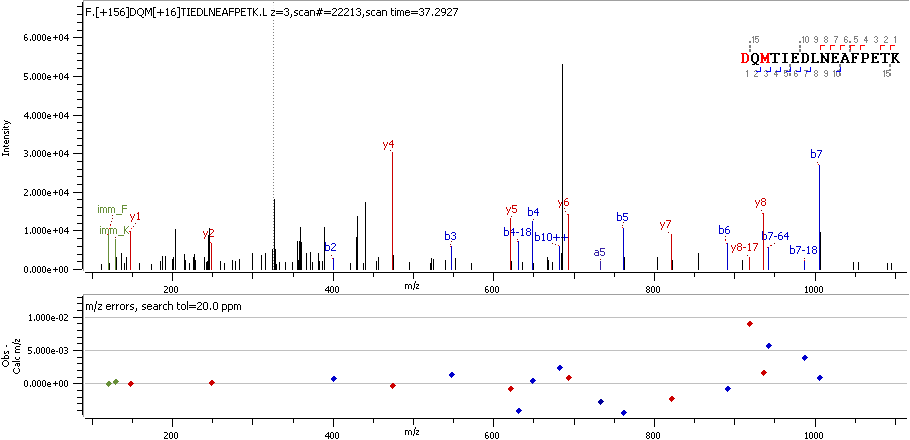

ATP5H D129 R

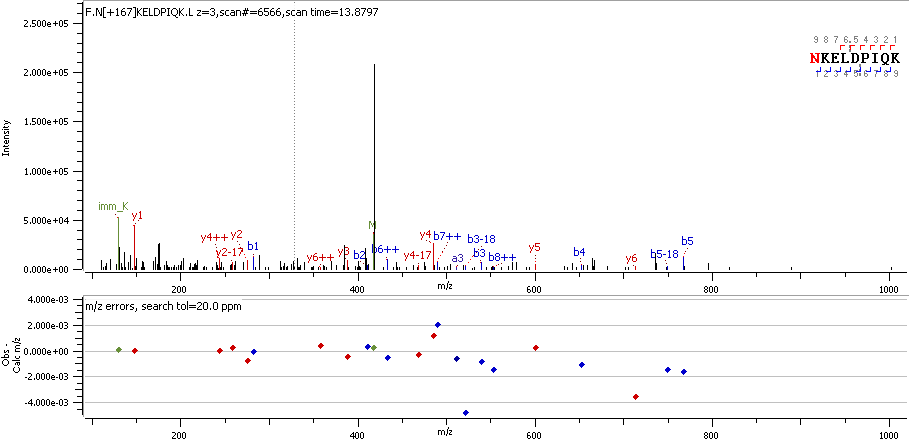

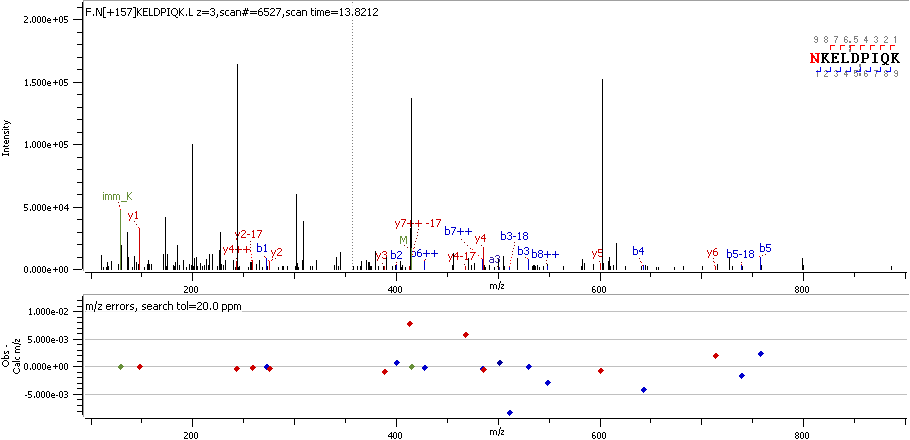

ATP5J N33 R_deami

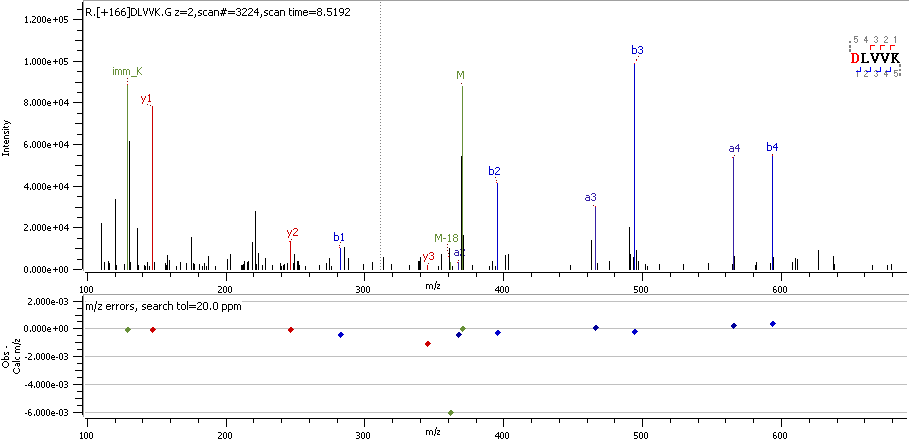

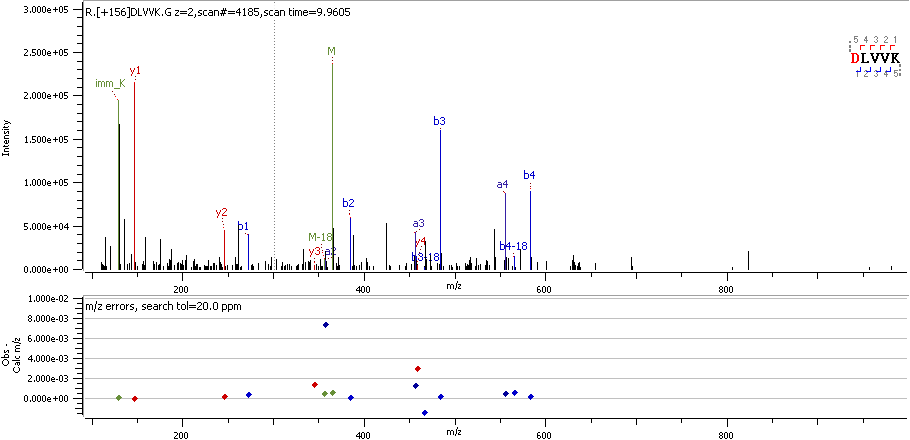

ATS15 D749 R

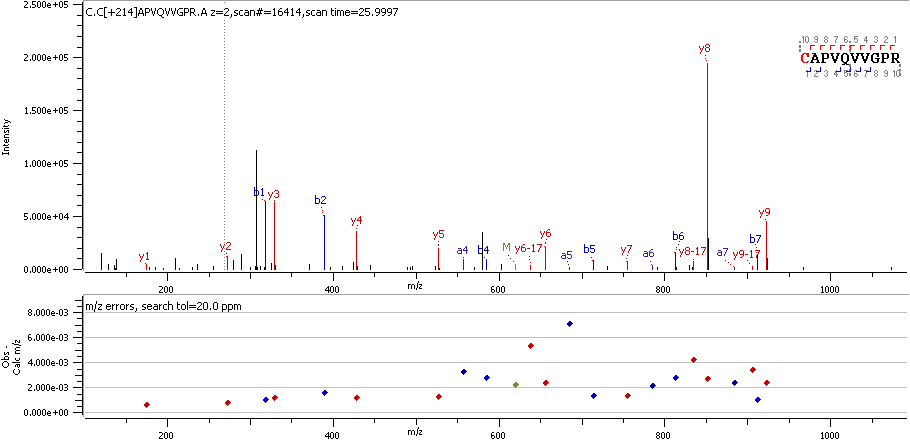

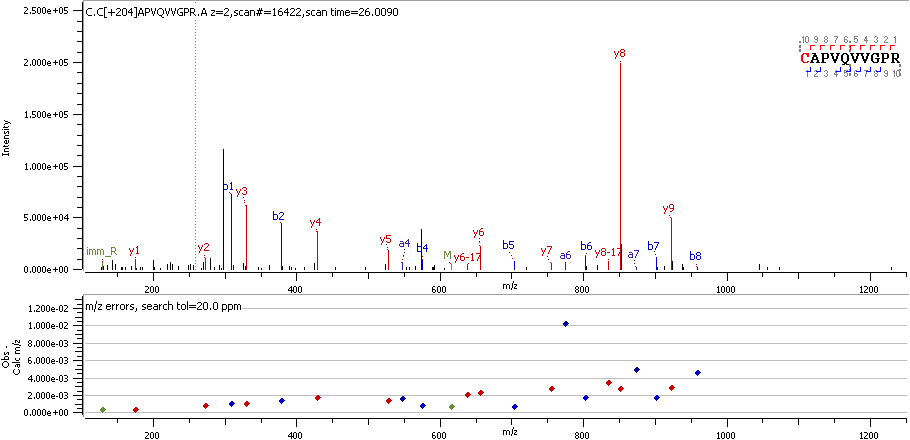

BACH C36 RO3

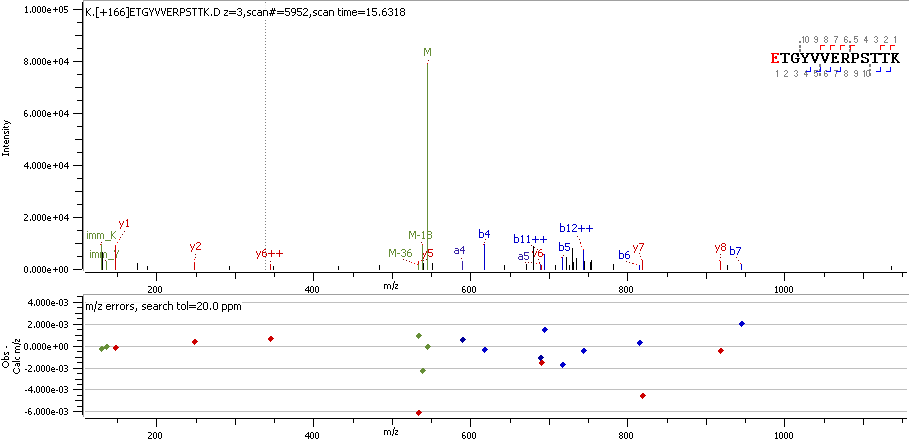

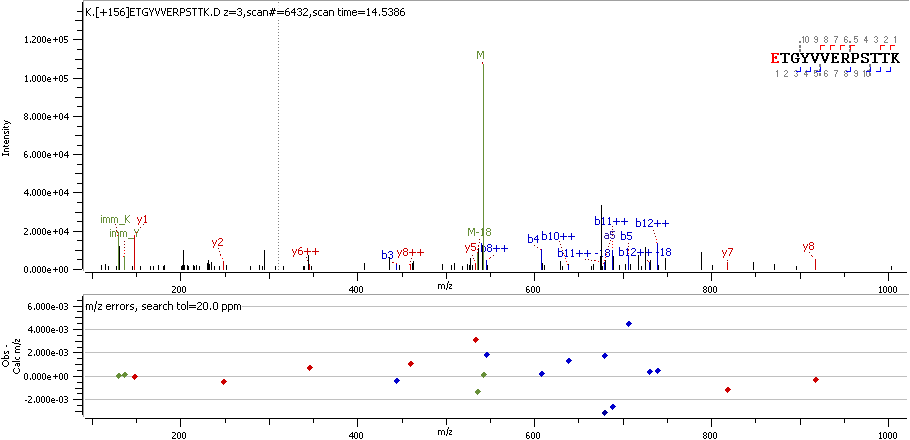

BCLF1 E463 R

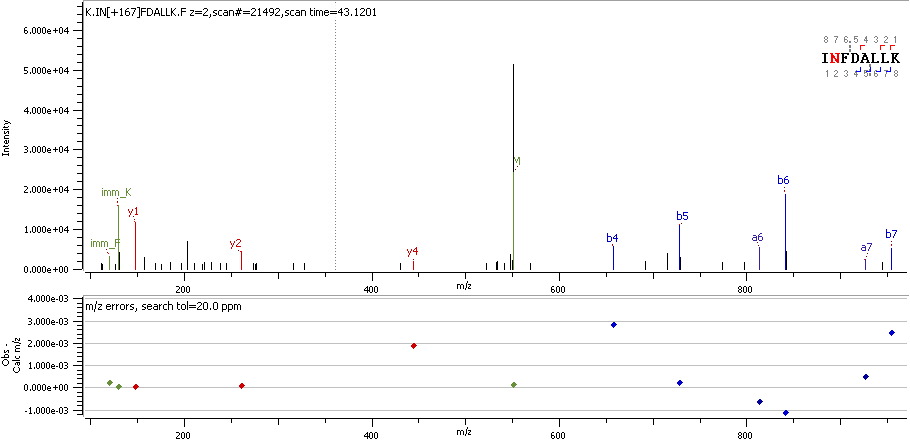

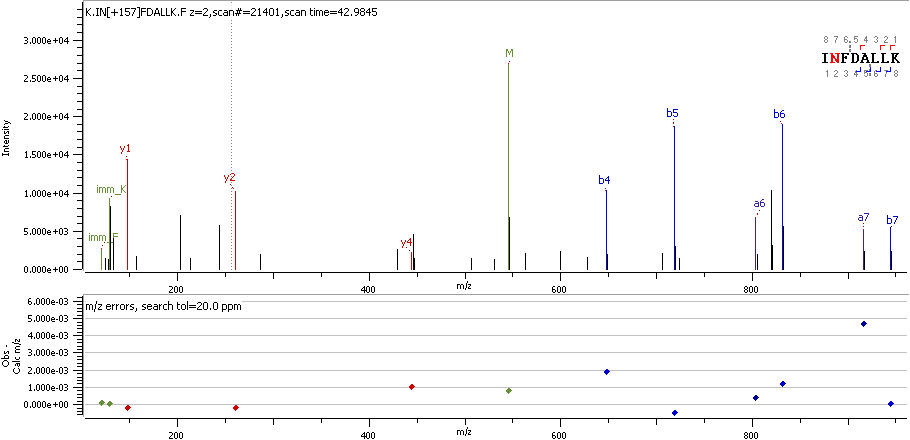

BI1 I 8R_deami

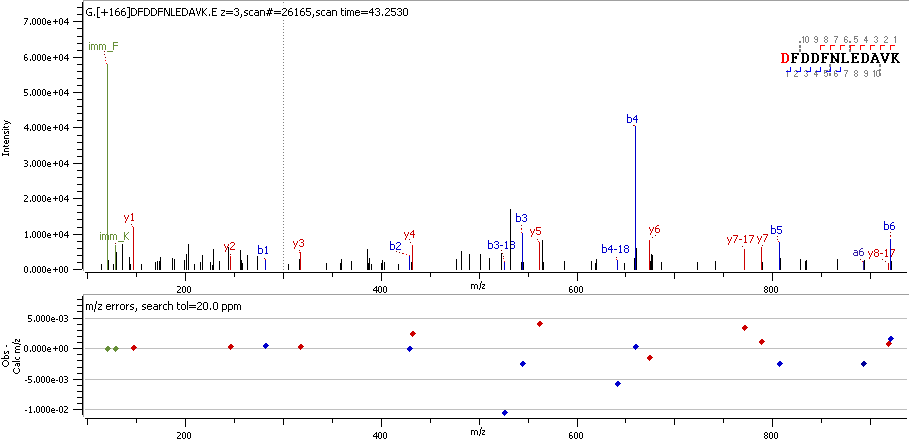

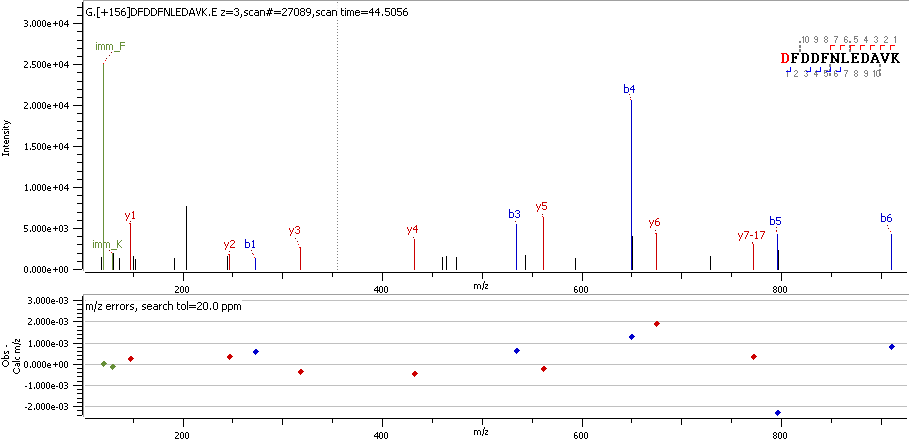

C99L2 D26 R

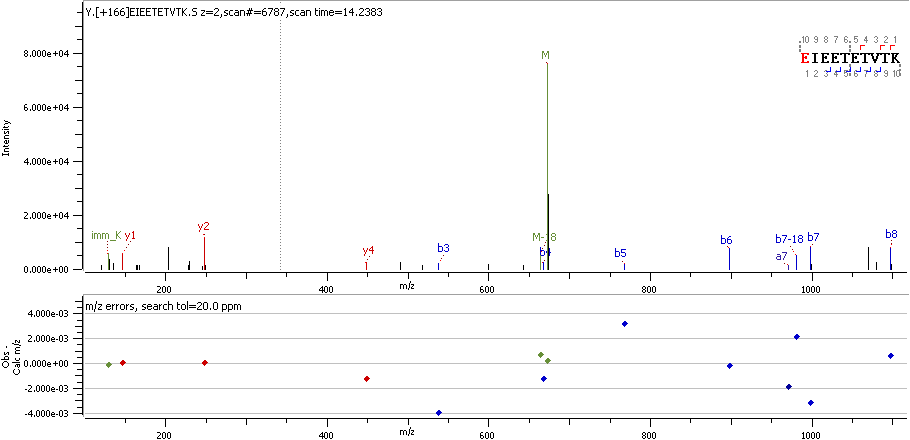

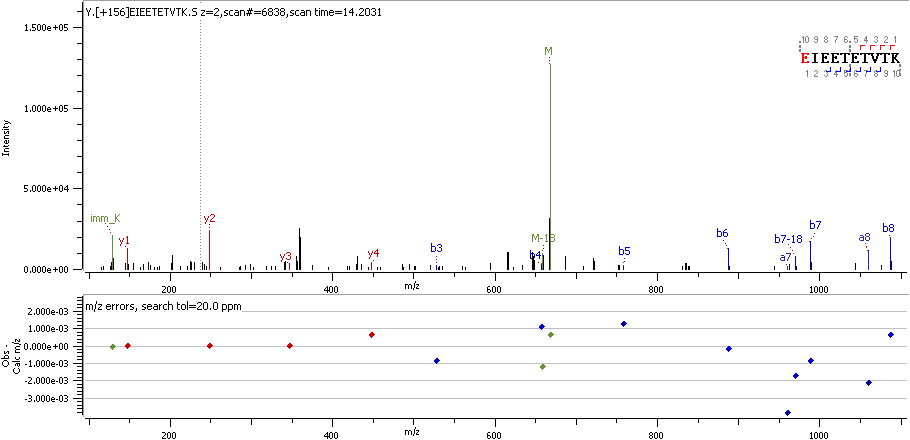

CALD1 E161 R

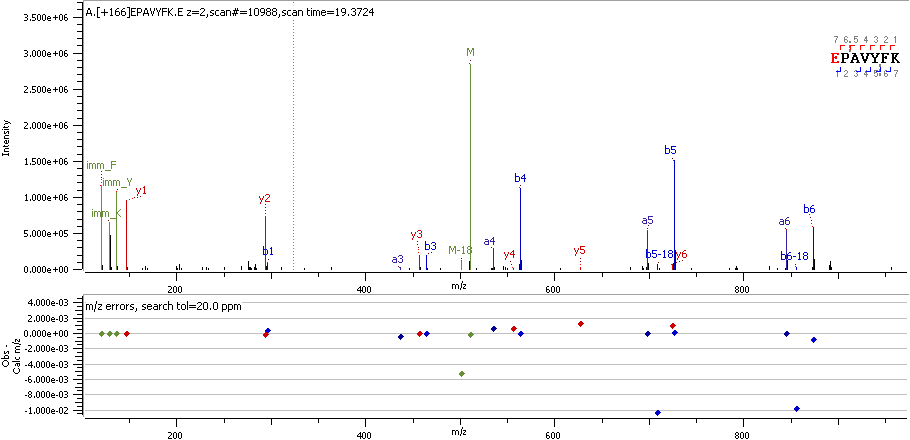

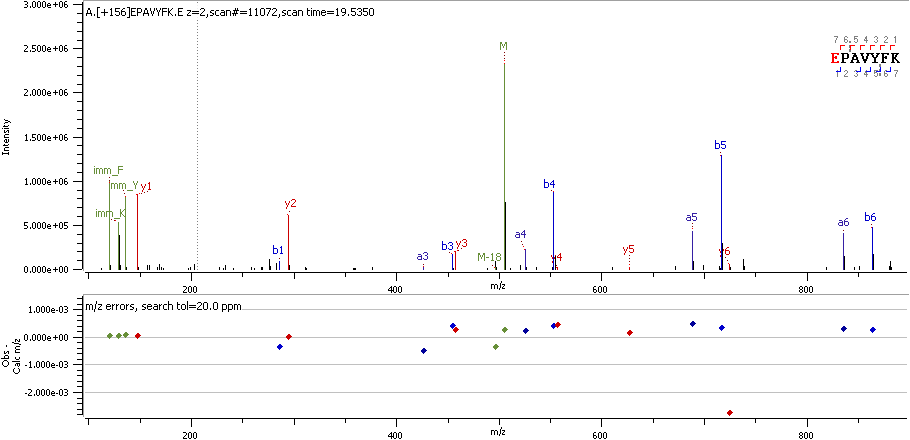

CALR E18 R

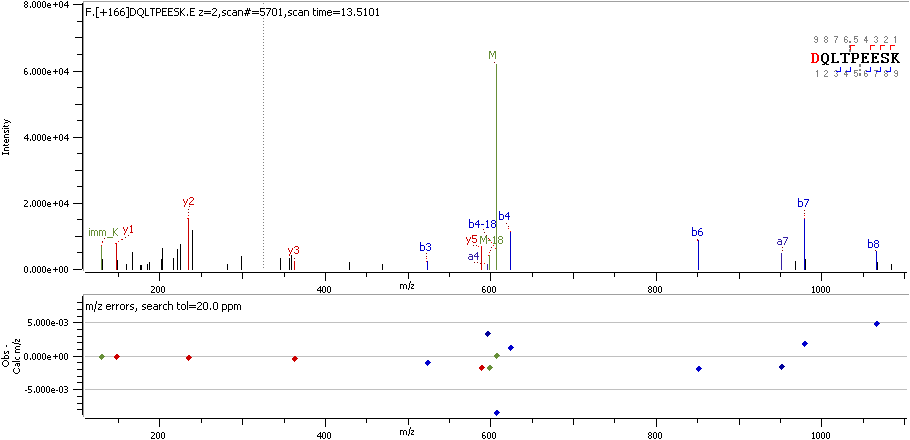

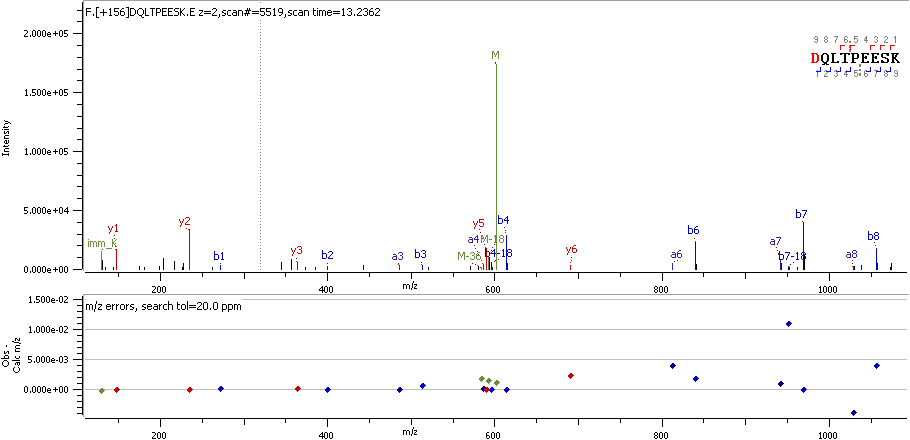

CALU D62 R

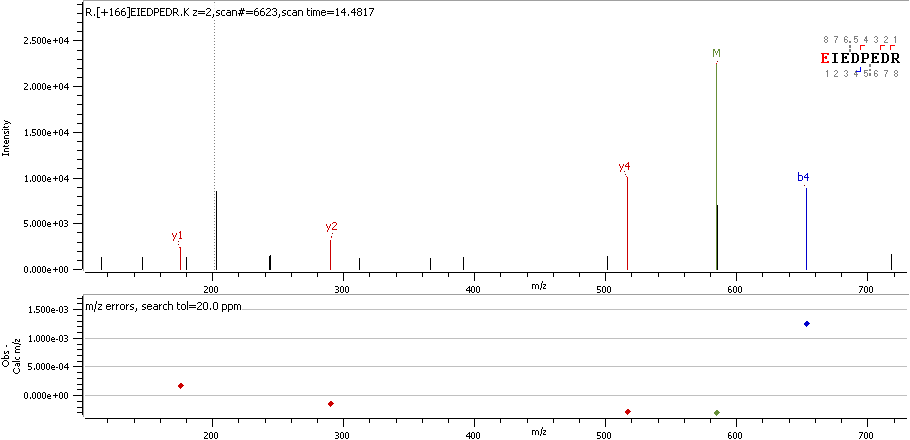

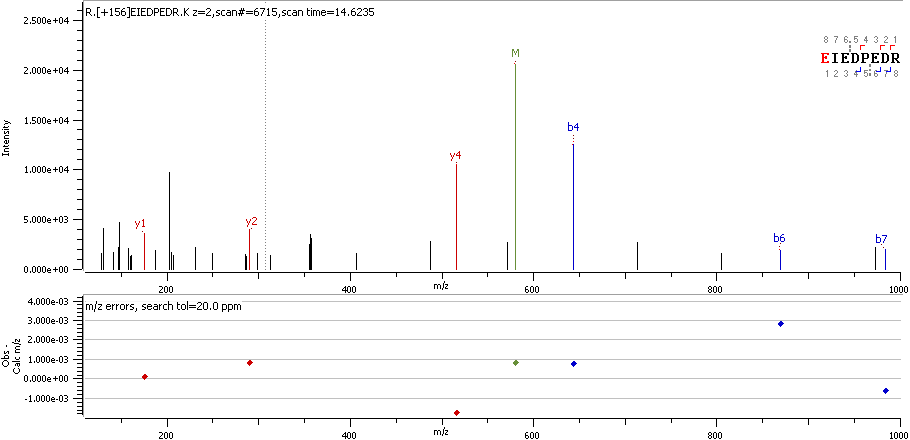

CALX E275 R

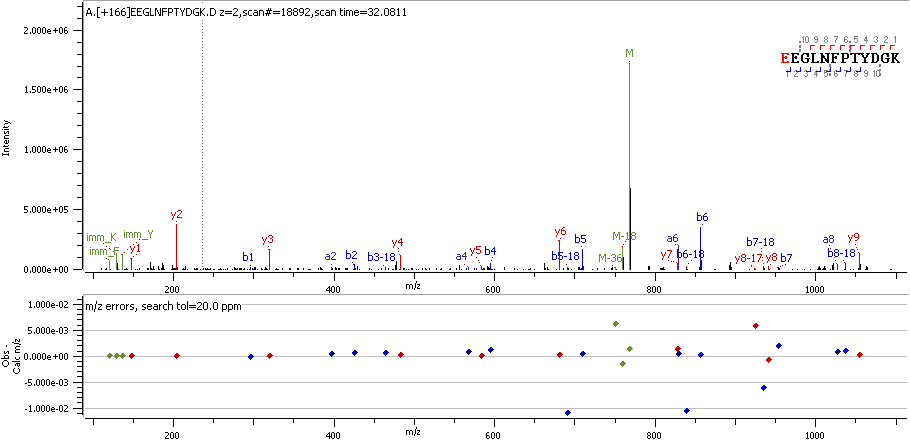

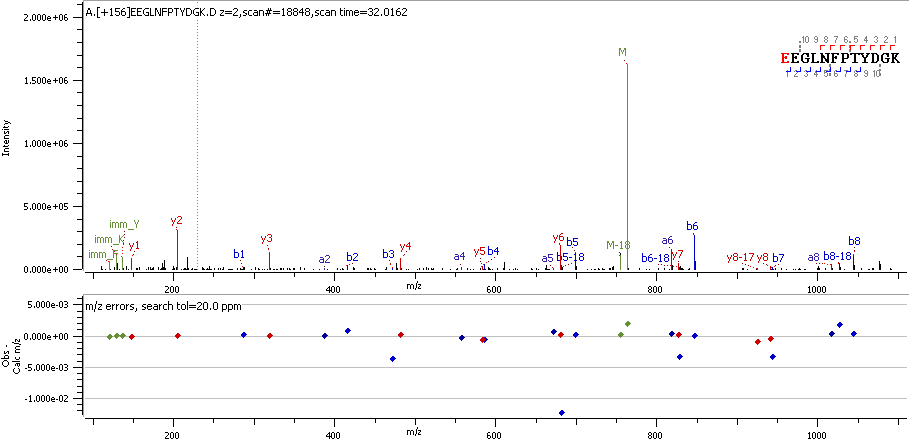

CASQ2 E20 R

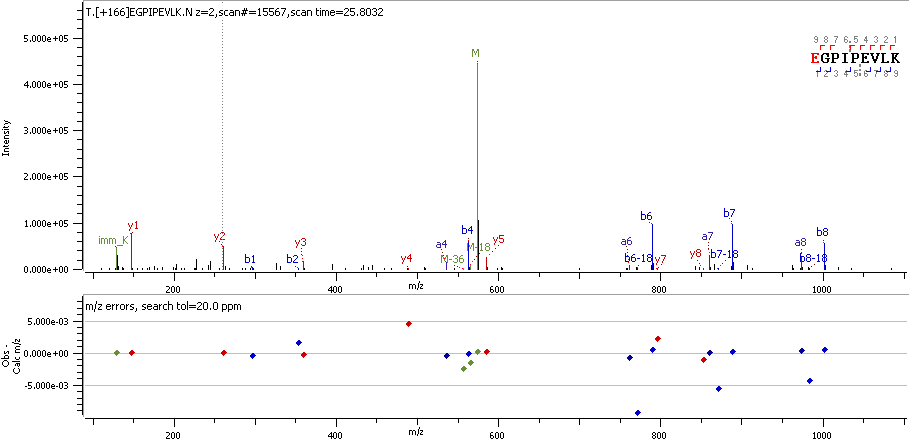

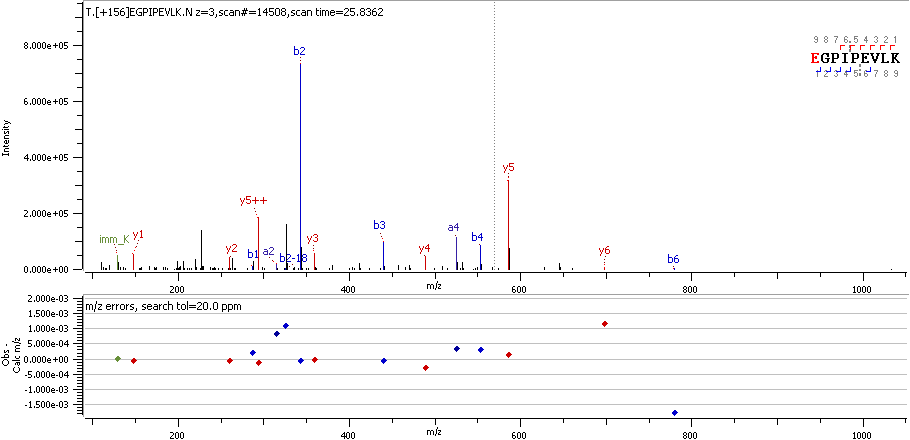

CATD E64 R

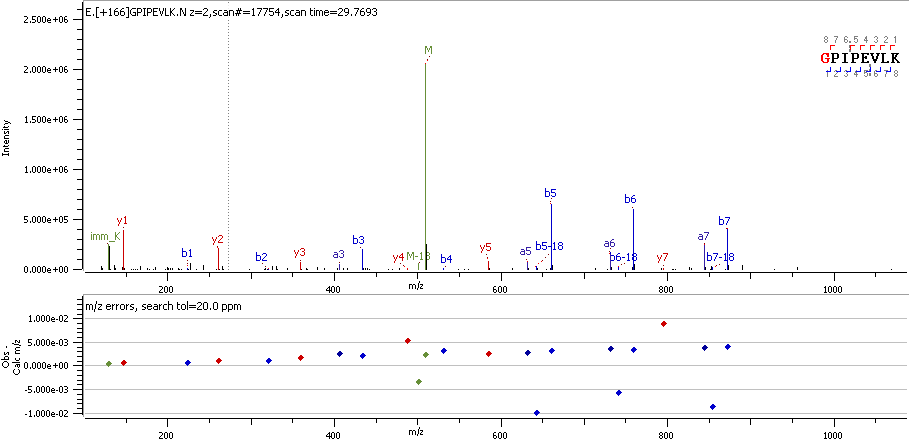

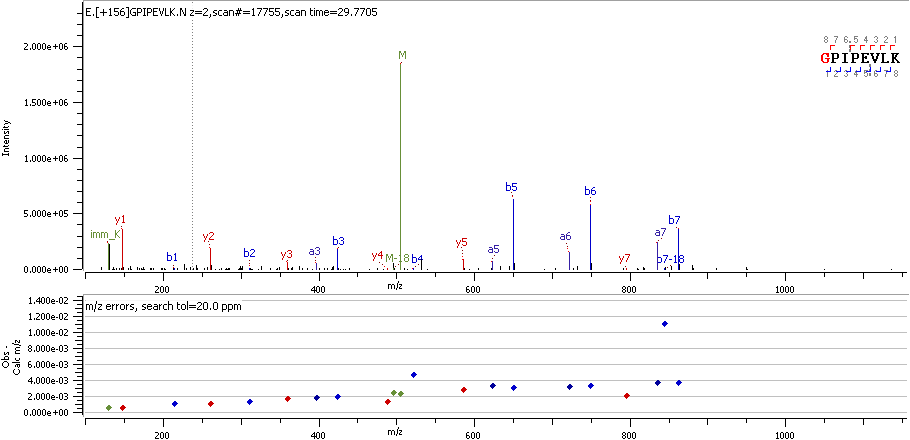

CATD G65 R

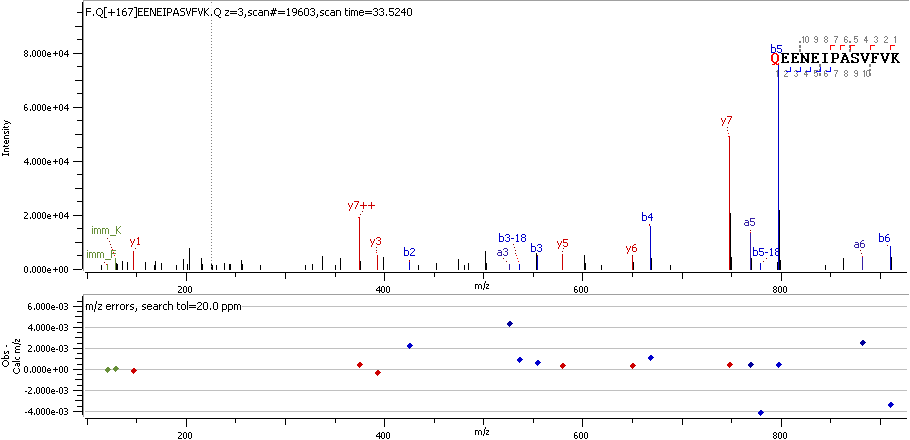

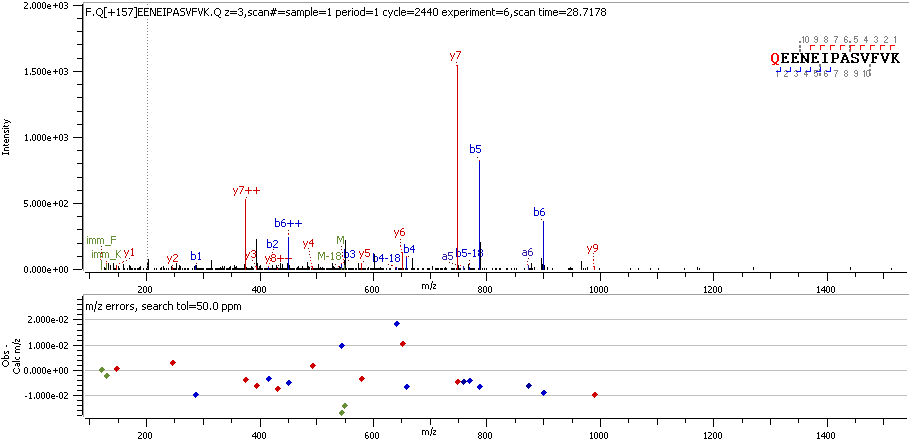

CAVN2 Q161 R_deami

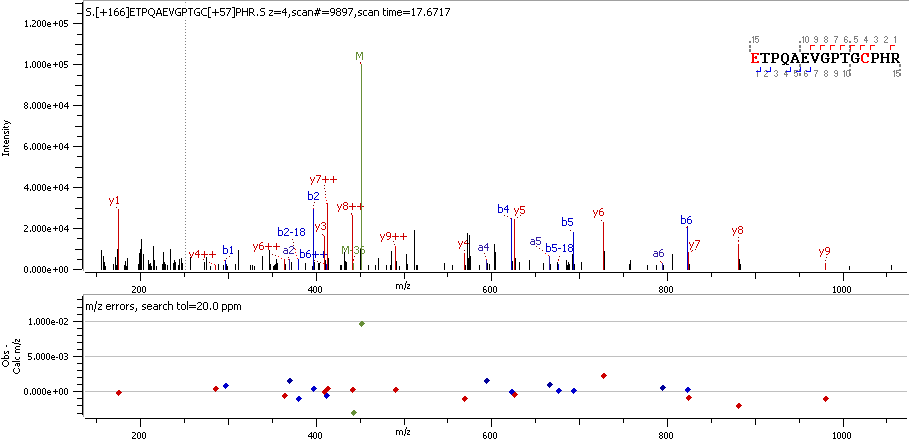

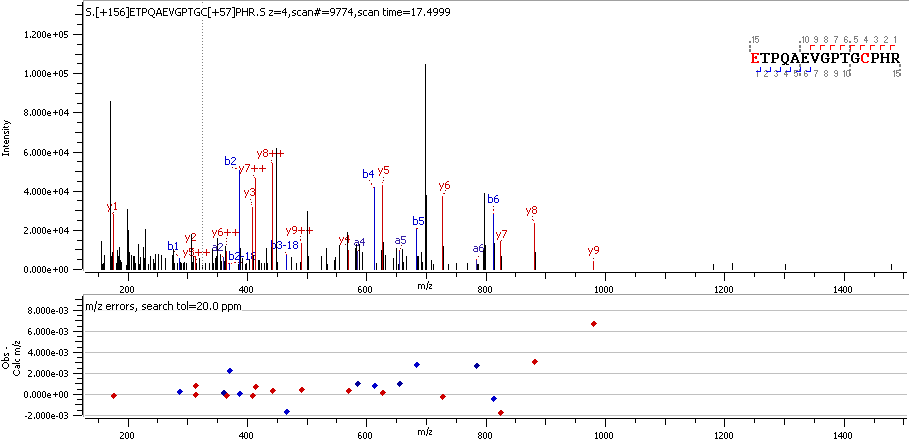

CBS E 4 R

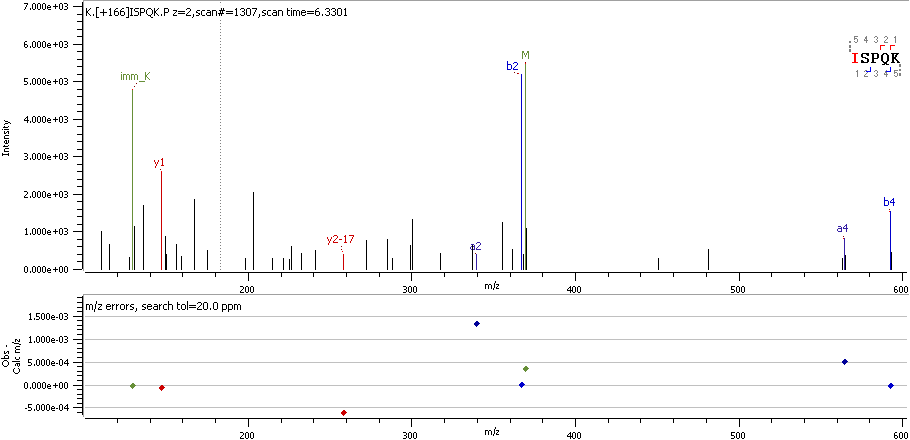

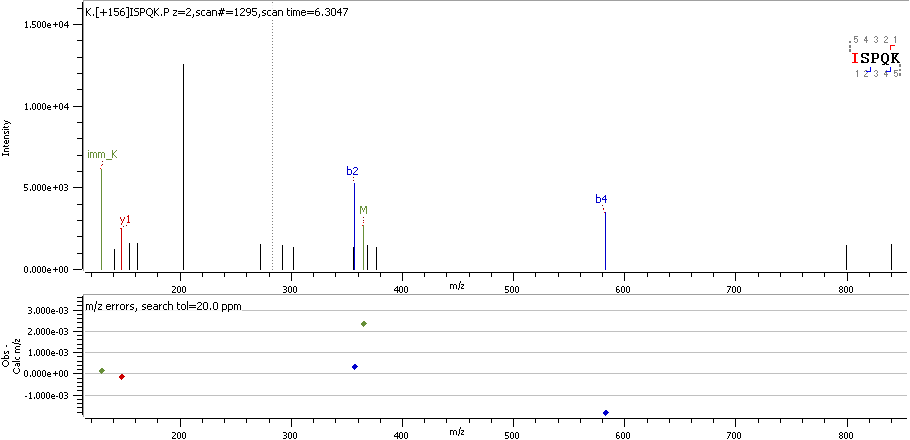

CC148 I574 R

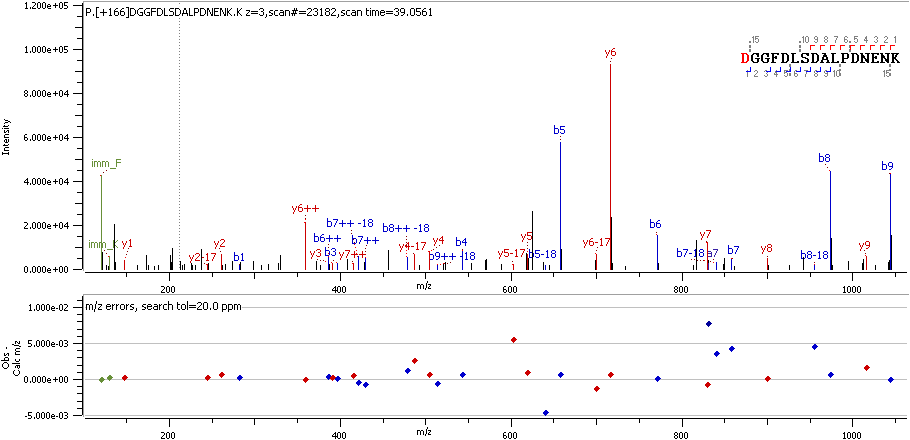

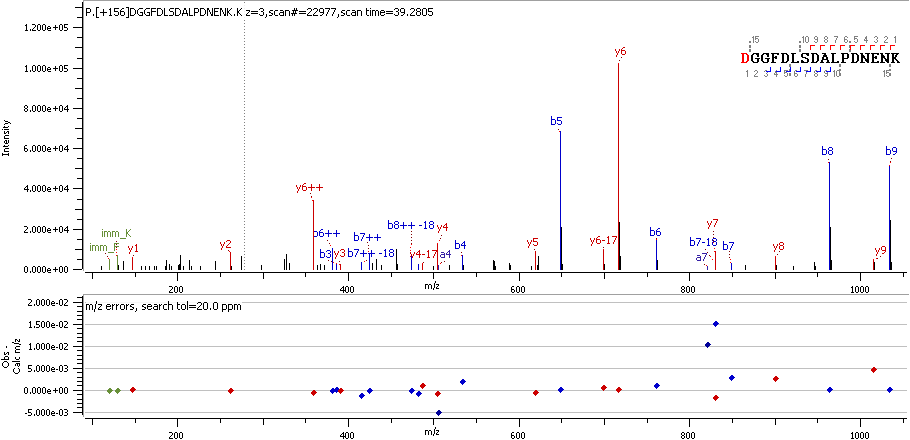

CD99 D23 R

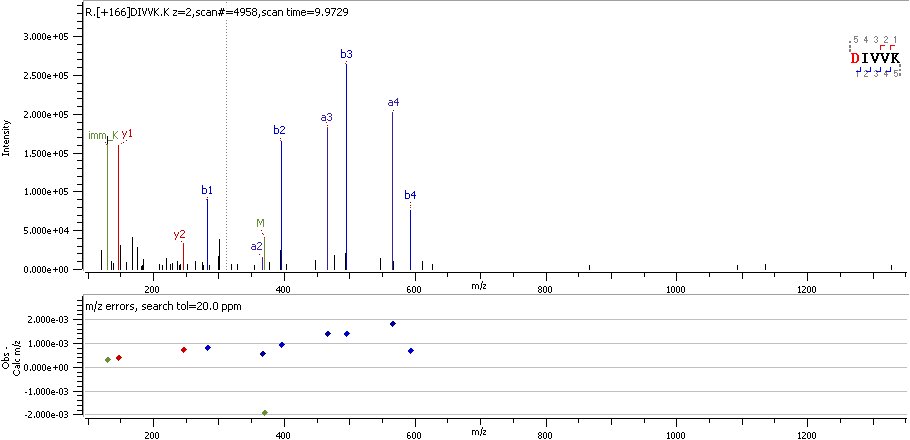

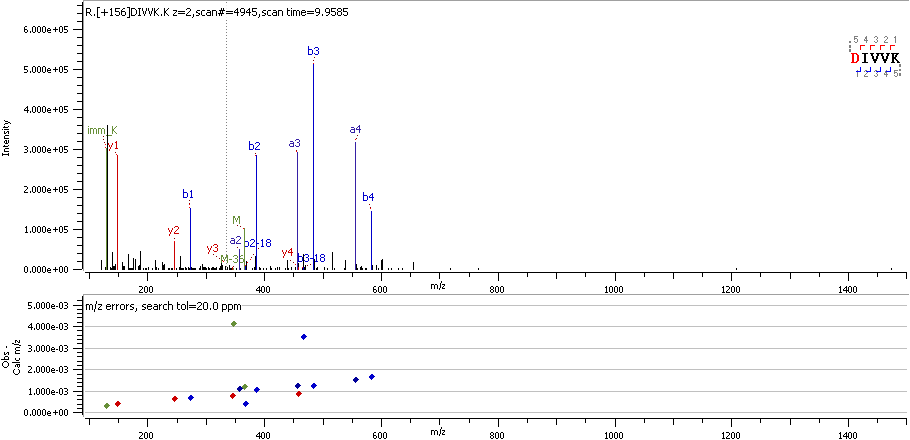

CDY1 D285 R

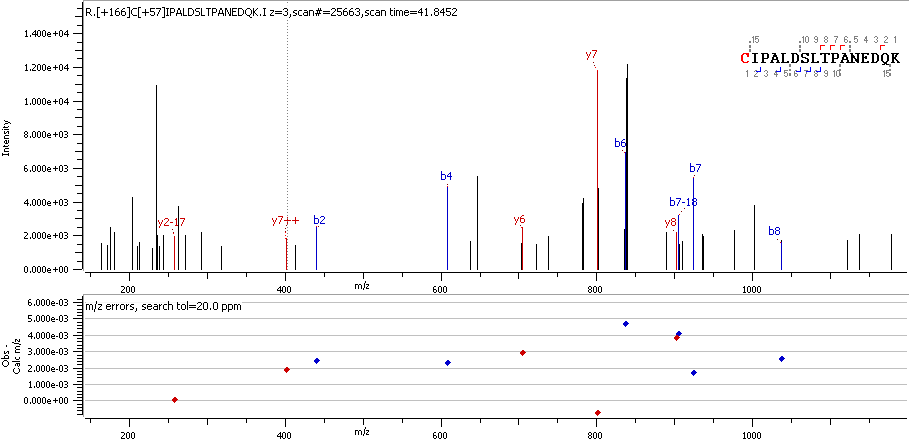

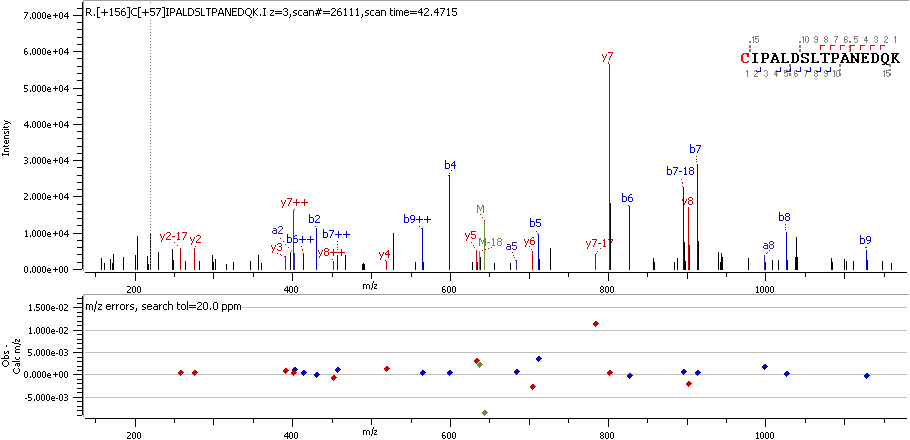

CH60 C447 R

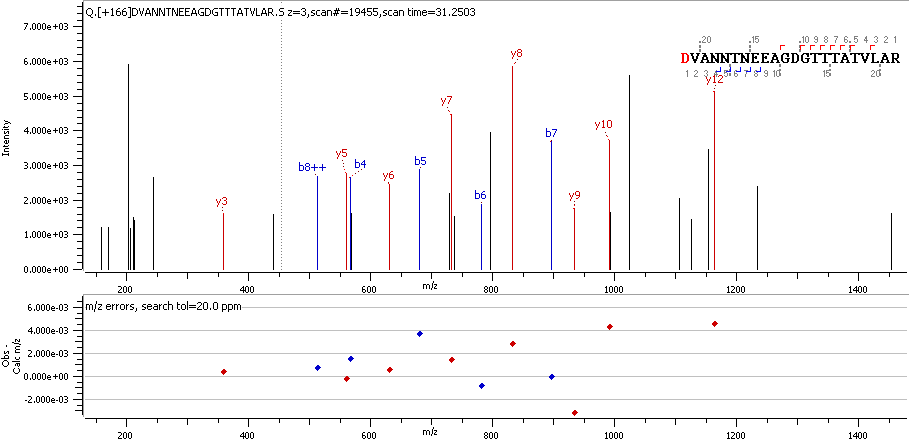

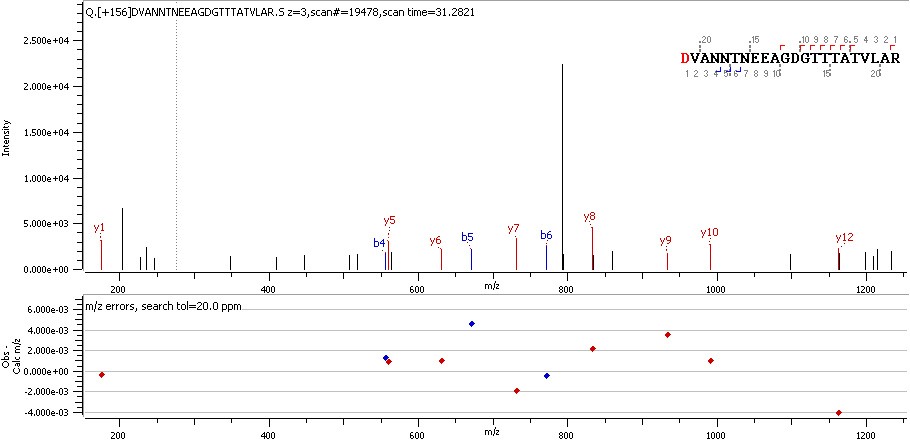

CH60 D100 R

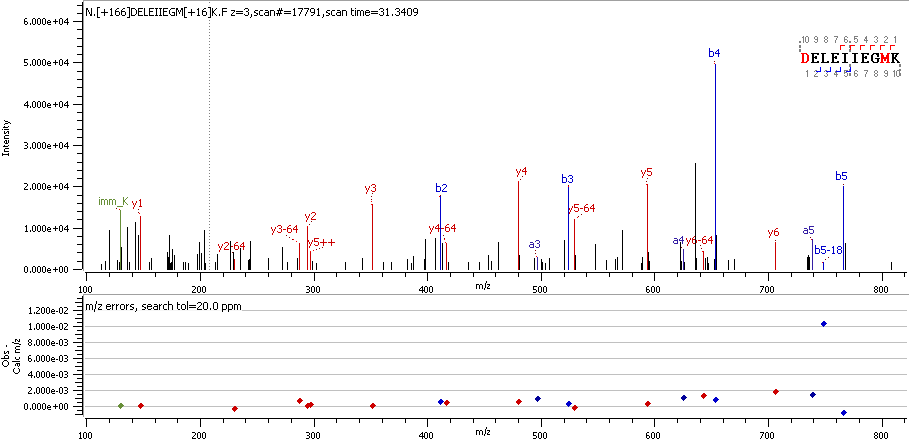

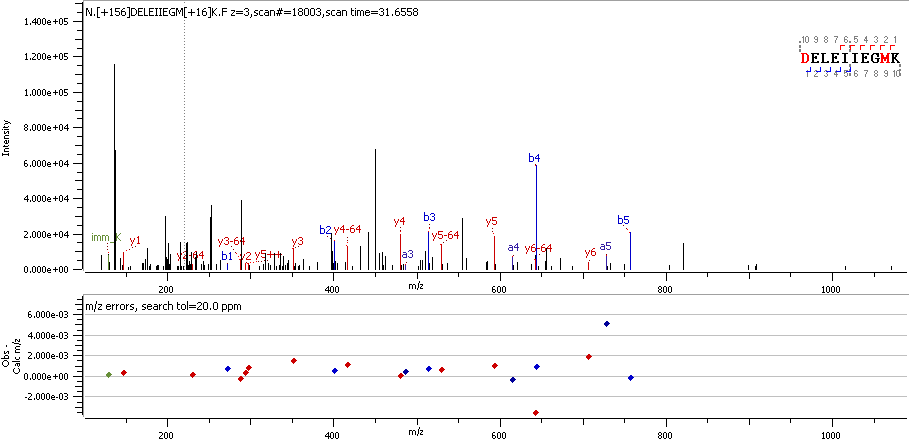

CH60 D209 R

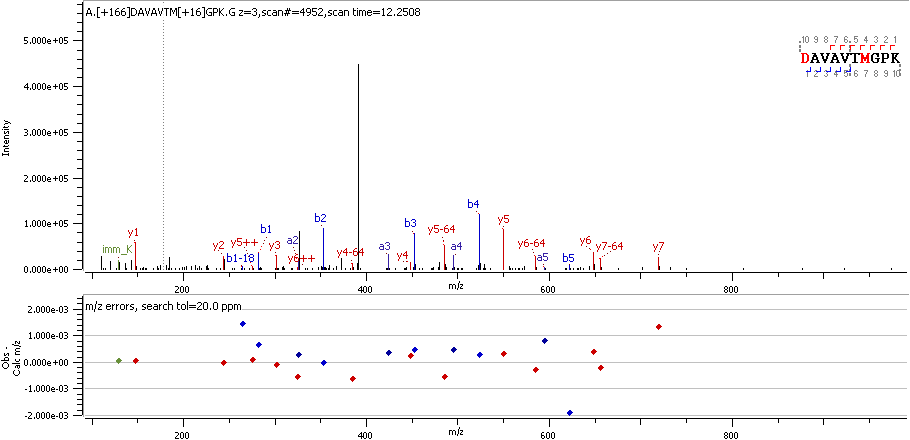

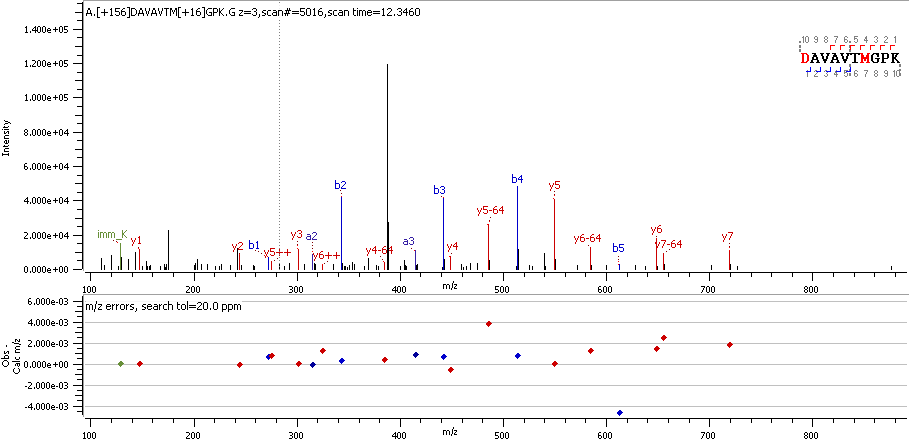

CH60 D49 R

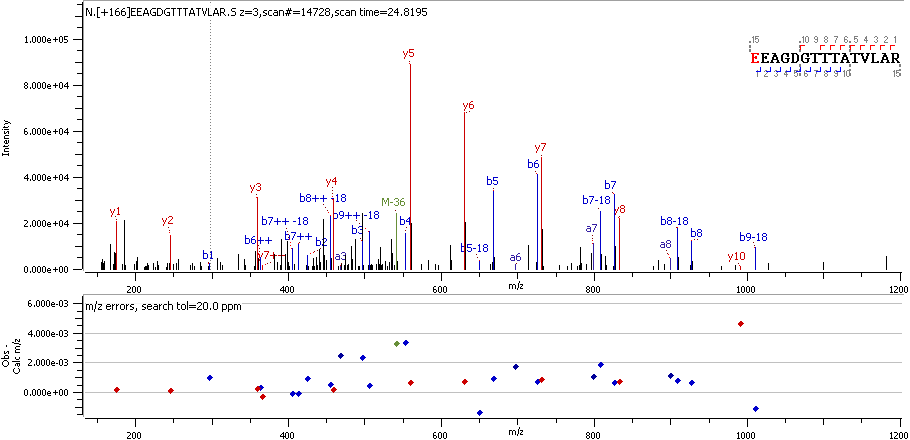

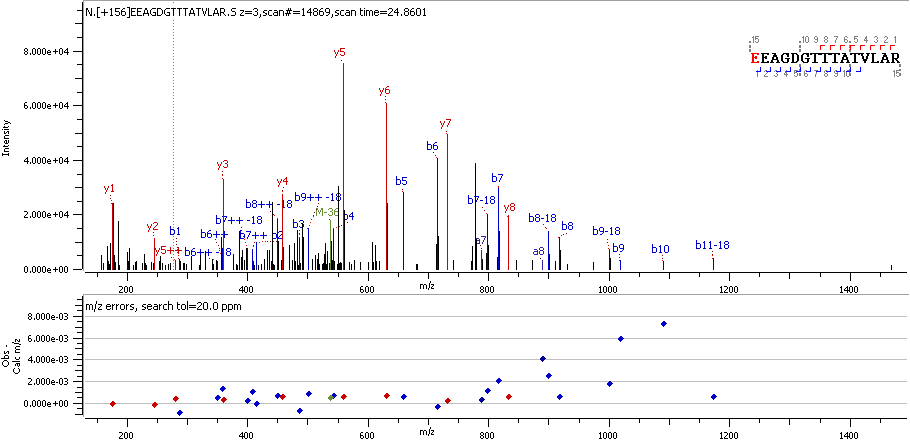

CH60 E107 R

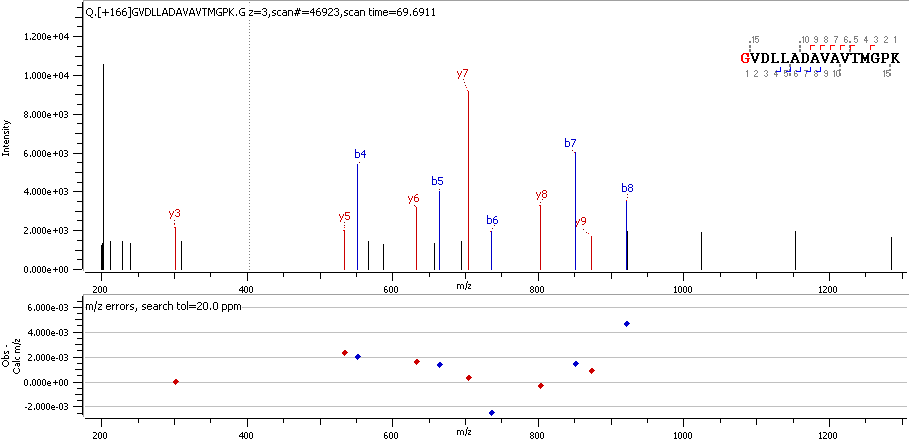

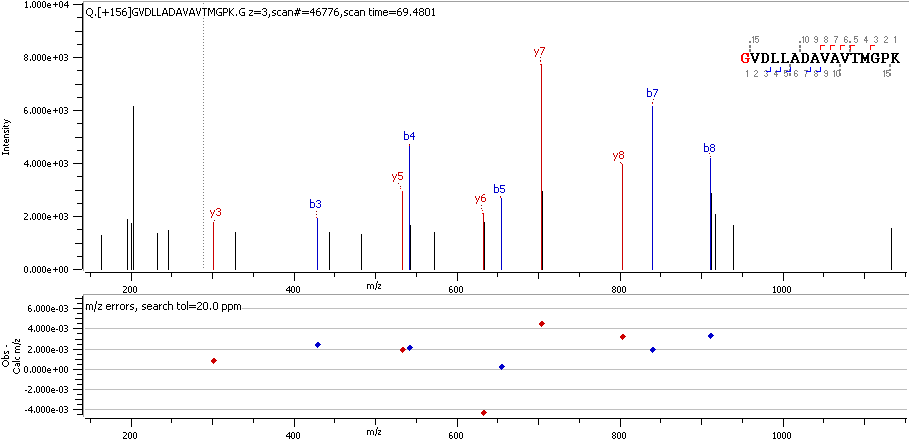

CH60 G43 R

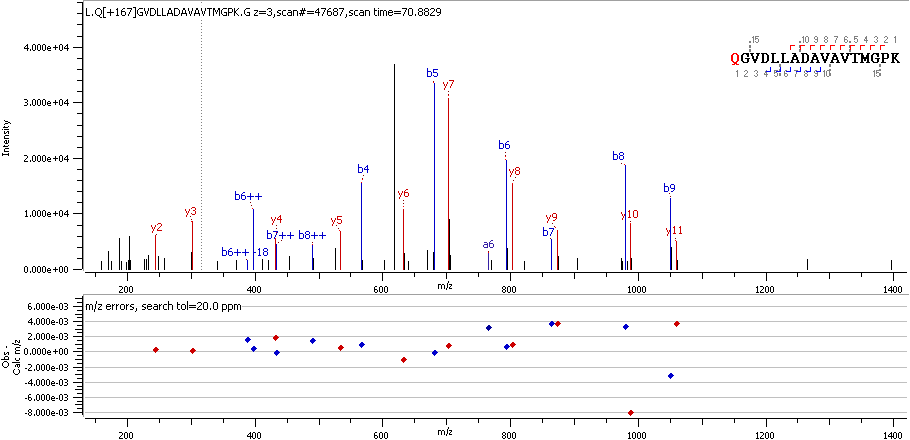

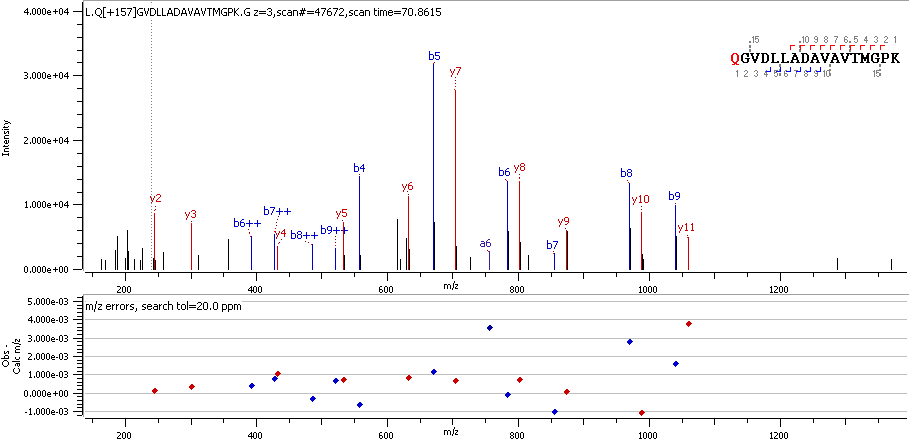

CH60 Q42 R_deami

CLUS D23 R

CNBP Q44 R_deami

COX17 D 6 R

COX41 D43 R

CWC15 Q45 R_deami

DJC10 D33 R

DPEP3 D150 R

DREB D634 R

DREB D636 R

DZI1L Q132 R

E2F7 D10 R

EF1D E27 R

EFHD2 Q67 R_deami

ENPL D22 R

ERO1A E24 R

F162A C41 RO3

F205A E1023 R

FCN1 D120 R

FUBP1 N369 R_deami

G3P D296 R

G3P D315 R

GATB E42 R

GFAP D417 R

GFAP E391 R

GLYM Q27 R_deami

H15 E 8 R

H2B1K E94 R

H4 D25 R

HACD3 P100 R

HBA D48 R

HBA D65 R_deami

HMGA1 E47 R

HMSD Q43 R_deami

HNRPC D123 R

HNRPK D279 R

HNRPK N306 R

HNRPK Q358 R_deami

HNRPL D284 R

HNRPU E259 R

HOOK1 D659 R

HS90B E539 R

HSP7C G372 R

IF4H D196 R

ILF2 D347 R

JTB E31 R

KAD4 G82 R

KS6A1 D37 R

KTN1 D238 R

LDB3 E257 R

LDB3 N197 R_deami

LETM1 D116 R

LMAN2 D45 R

LN28A E18 R

MAP1A E1580 R

MAP1A E663 R

MAP1B Q907 R_deami

MBP D168 R

MBP D216 R

MBP D279 R

MBP E217 R

MBP G277 R

MBP N226 R

MBP N226 R_deami

MBP T200 R

MCFD2 E27 R

MESD E35 R

MITOK C37 RO3

MLRV E22 R

MLRV N154 R_deami

MPPB Q46 R_deami

MTAP2 Q431 R_deami

MTDC E36 R

MYH6 E73 R

MYH7 D1008 R

MYH7 D1229 R

MYH7 E1376 R

MYH7 E1461 R

MYH7 E1489 R

MYH7 E1821 R

MYH7 E1902 R

MYH7 E73 R

MYH7 E927 R

MYH9 D1136 R

MYH9 D1846 R

MYL3 E131 R

MYOZ2 D255 R

MYPT2 D816 R

NCAM1 E833 R

NCAN E23 R

NDKM K91 R

NECP1 C162 RO3

NEUG C 9RO3

NEUM E89 R

NFH E469 R

NFL D468 R

NFL E453 R

NFL E469 R

NFL E507 R

NFL E509 R

NFL Q75 R_deami

NFM E454 R

NFM E455 R

NFM E470 R

NFM E496 R

NFM E497 R

NFM E599 R

NFM E763 R

NFM E884 R

NHRF1 E121 R

NOP16 E77 R

NPC2 E20 R

NPM D178 R

NPM D179 R

NPM D180 R

NT5D3 C44 RO3

NUCL E628 R

ODO2 D68 R

PACN1 N330 R_deami

PCDH1 D944 R

PDIA1 D18 R

PDIA4 E25 R

PDLI5 E215 R

PDLI5 E297 R

PELP1 E1008 R

PGCB D23 R

PGCB E26 R

PGCB E412 R

PGRC1 D49 R

PLEC E4050 R

PSD12 G331 R

PSMD6 L84 R

PTBP1 D 7 R

PTBP1 G306 R

PTGDS E25 R

RD23B E136 R

RL1D1 D319 R

RL23A E 8 R

RLA1 D18 R

RLA1 D19 R

RLA1 N62 R_deami

RM12 C45 R

RM12 C45 RO2

RM12 C45 RO3

ROA1 E93 R

ROA2 D161 R

RON D1232 R

RS27 D 6 R

RS7 E188 R

RT18A E37 R

S10AE C74 RO3

SAP D405 R

SAP G407 R

SCG1 D88 R

SCMC1 Q16 R_deami

SCN1A V1579 RO2

SF3B1 D93 R

SFPQ E696 R

SNX17 E463 R

SPTB2 E1469 R

SPTN1 E1124 R

SRCA E21 R

SRRT D866 R

SRRT D868 R

SSBP E17 R

TAU N644 R_deami

TBA1B D218 R

TBA1B D251 R

TBA1B D327 R

TBA1B D69 R

TBA4A D69 R

TBB4A D67 R

TBB5 E 3 R

TCAF1 V114 R

TFF1 E25 R

TMM11 D163 R

TNNI3 E165 R

TNNT2 D108 R

TXND5 E40 R

TYB4 D 6 R

UTP18 E374 R

VATD E242 R

VIME D451 R

VIME D90 R

VIME E425 R

XRCC5 E583 R
